# Flanking sequences regulate the toxicity, non-ATG translation, and aggregation of RNA with tandem CAG Repeats

**DOI:** 10.1101/2021.08.30.458106

**Authors:** Michael R. Das, Yeonji Chang, Reuben Saunders, Nan Zhang, Colson Tomberlin, Ronald D. Vale, Ankur Jain

## Abstract

Nucleotide repeat-expansions cause several neurodegenerative disorders, including Huntington’s disease and spinocerebellar ataxia. The expanded repeat-containing RNA transcribed from the affected loci agglomerate in the nucleus as pathogenic foci. Here we demonstrate that depending on their surrounding sequence context, RNAs with expanded CAG repeats can also undergo nuclear export and aggregate in the cytoplasm. Cytoplasmic aggregation of repeat-containing RNA coincides with several disease hallmarks, including repeat-associated non-AUG (RAN) translation, mislocalization of RNA binding proteins, and cell toxicity. Interestingly, the repeat-containing RNA co-aggregate with RAN translation products. Inhibition of RAN translation prevents cytoplasmic RNA aggregation and also alleviates cell toxicity. Our findings provide a cogent explanation for aberrant cytoplasmic localization of RNA binding proteins and implicate cis-acting flanking sequences in mediating RAN translation and disease.

## Introduction

Over 40 genetic diseases are traced to the aberrant expansion of short nucleotide repeats (Khristich and Mirkin, 2020). This family of diseases, collectively referred to as nucleotide repeat expansion disorders, includes Huntington’s disease, several spinocerebellar ataxias, myotonic dystrophy, and certain forms of amyotrophic lateral sclerosis (ALS). Disease-causing repeats are found in both protein-coding as well as non-coding regions. For example, Huntington’s disease results from a CAG trinucleotide repeat expansion in exon 1 of the huntingtin gene, whereas myotonic dystrophy is caused by a CTG repeat expansion in the 3’ untranslated region of *DMPK*. Disease usually manifests when the number of repeats exceeds a critical threshold (Khristich and Mirkin, 2020).

Although repeat expansion diseases are monogenic disorders, the disease mechanisms are more complex than the simple loss of function of the mutated gene (Paulson, 2018). In particular, the RNA transcripts harboring pathogenic repeat expansions are suggested to promote pathogenesis via at least two routes. First, the repeat-containing RNAs can form higher-order assemblies and accumulate as ‘foci’ in the nucleus. These nuclear RNA foci sequester various RNA-binding proteins and result in transcriptome-wide RNA processing (Jiang et al., 2004; Kanadia et al., 2006; Wojciechowska and Krzyzosiak, 2011). Second, the disease-associated repeat-containing RNA can be translated without a bona fide AUG-start codon in multiple reading frames: a process referred to as repeat-associated non-AUG or RAN translation (Zu et al., 2011). The homopolymeric peptides produced upon RAN translation disrupt cellular functions and are potentially lethal to the cell (Kwon et al., 2014; Lee et al., 2016; Zhang et al., 2014).

Retention of RNA at nuclear foci could potentially limit its access to the translation apparatus in the cytoplasm. The interplay between these two RNA gain-of-function routes and their relative contribution to cell toxicity remains to be established. In addition to RNA foci and RAN translation, a common theme in repeat expansion diseases is the aberrant cytoplasmic localization of several RNA binding proteins such as TDP-43 and FUS (DeJesus-Hernandez et al., 2011; Deng et al., 2010; Schwab et al., 2008). Mechanistically, how the repeat expansion mutation results in cytoplasmic recruitment of otherwise nuclear proteins is unknown. Another peculiar feature of repeat expansion disorders is that the sequences surrounding the repeat can also influence cellular toxicity and disease (Cleary and Pearson, 2003), but the molecular basis for this phenomenon is not clear.

Here, we examined how the surrounding sequence context influences RNA toxicity. We demonstrate that RNA with expanded CAG repeats exhibit markedly different sub-cellular localization depending on their upstream flanking sequences. In most cases, incorporation of expanded CAG-repeats results in retention of the RNA at nuclear foci. However, certain upstream flanking sequences facilitate nuclear export and RAN translation of the repeat-containing RNA. RAN translation products co-aggregate with the repeat-containing RNA and accumulate in perinuclear cytoplasmic inclusions. These RNA-RAN protein inclusions sequester TDP-43, an RNA binding protein frequently mislocalized in neurodegenerative disease, as well as p62, a marker for protein aggregation. RNA with CAG repeats in surrounding sequence contexts that result in cytoplasmic aggregates are substantially more toxic than ones that only result in nuclear foci. RAN translation is required for cytoplasmic RNA aggregation, and inhibition of RAN translation rescues the cells from repeat RNA mediated toxicity. Our findings establish a direct regulatory role of flanking sequences in determining the localization of repeat-containing RNA, causing RAN translation, and mediating cell toxicity.

## Results

### Flanking sequences influence the toxicity and sub-cellular localization of CAG repeat-containing RNA

To study the effect of the surrounding sequence context on repeat-containing RNA, we developed a minimal reporter system. We generated 12 synthetic sequences (250 bp each, GC-content 30-79%, sequences in Supp. Table 1) and introduced them upstream of CAG-repeats (Fig. 1A). Multiple stop codons were incorporated in each reading frame between the test sequences and the CAG repeats to prevent readthrough translation from potential start codons in the upstream flanking sequences. Downstream of the CAG repeats, we incorporated twelve MS2 hairpins, which allowed live-cell RNA visualization via co-expression of a YFP-tagged MS2 coat protein (MS2CP-YFP) (Bertrand et al., 1998). The repeat-containing RNAs were expressed under the control of a doxycycline-inducible promoter.

**Figure 1.**
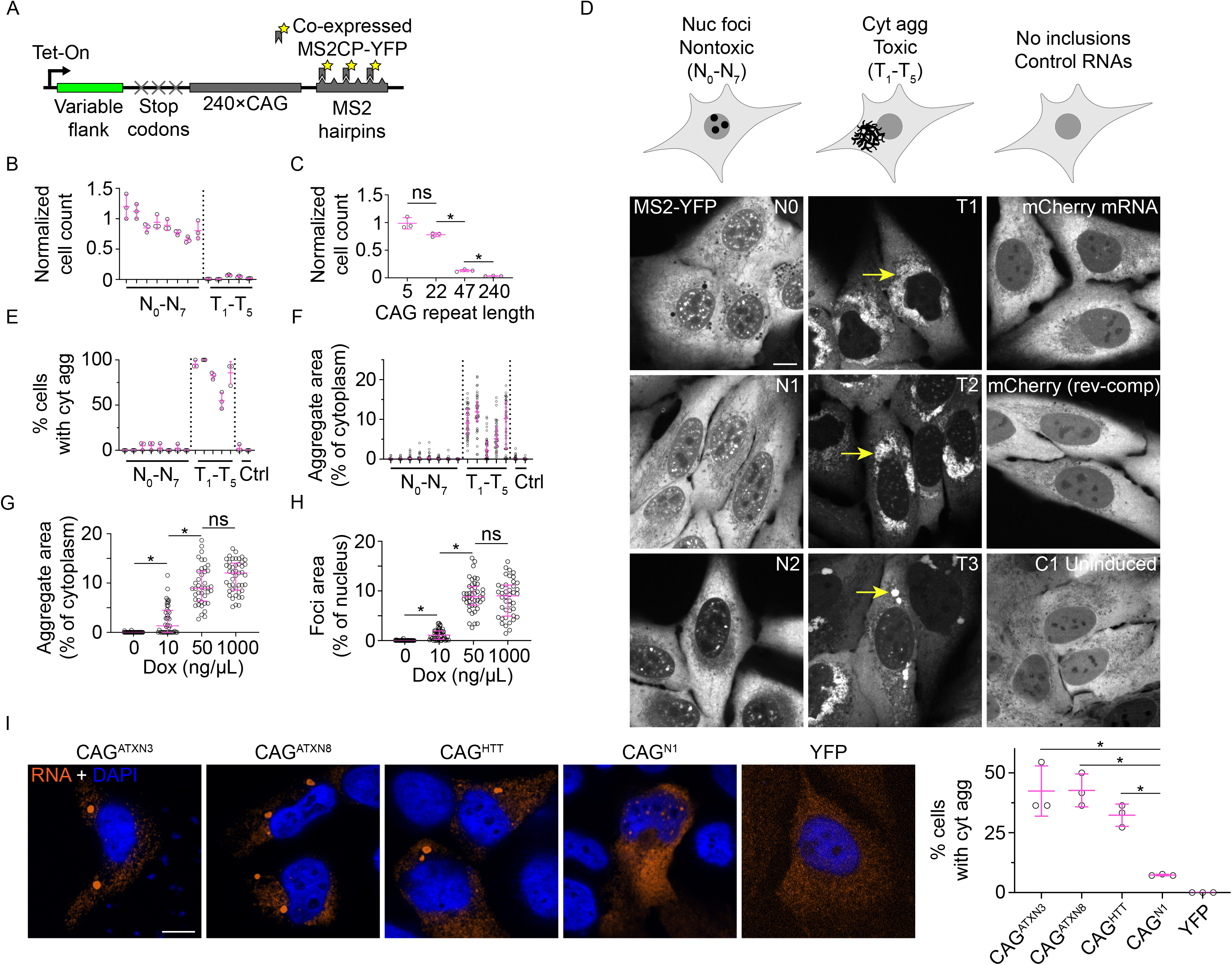
Flanking sequences modulate the localization of CAG repeat-containing RNA **A,** Schematic of repeat-containing constructs. **B, C**, Quantification of cell toxicity in lines expressing CAG repeats with various flanking sequences (B) or as a function of CAG repeat number for the same flanking sequence as in T1 (C). Each data point represents one experiment; toxicity was measured 5 days post induction. Data are summarized as mean ± s.d. Asterisks denote significance by Student’s t-test. **D,** Schematic depicting the observed localization of CAG repeat-containing RNA (top). Representative fluorescence micrographs 24 h post-induction (bottom). Arrows denote representative cytoplasmic aggregates. Micrographs are representative of ≥ 40 cells from ≥ 3 independent experiments. **E,** Quantification of the percent of cells containing cytoplasmic inclusions. Data are representative of at least three experiments. Control cells express mCherry mRNAs (left control on each graph) or the reverse-complement of mCherry RNAs (right control on each graph), each fused to MS2 hairpins. **F,** Quantification of aggregate area. Each point represents a single cell. Data are summarized as median ± interquartile range. Control cells are the same as in E. **G, H,** Quantification of the area of cytoplasmic aggregates (G) and nuclear foci (H) at varying doxycycline concentrations which results in varying levels of RNA expression. Each point represents a single cell; data are summarized as median ± interquartile range. Asterisks denote significance by Mann-Whitney test. **I,** Representative fluorescence micrographs 48 h after transfection with indicated constructs (left). Quantification of percent of cells with cytoplasmic aggregates (right). Each point represents ≥ 10 cells. Data are representative of at least 3 independent experiments. Asterisks denote significance by Student’s t-test. All scale bars, 10 µm. *: p < 0.01, n.s.: not significant (p > 0.01).

After prolonged (5 days) expression of the repeat-containing RNA, we observed two broad classes of cellular phenotypes. The majority of constructs (7 out of 12 sequences tested) did not produce a discernable difference in cellular viability as compared to uninduced controls (constructs N1-N7 for nontoxic CAG repeats, Fig. 1B). This result is similar to our previous report that the expression of RNA with expanded CAG repeat near the 5’ end of the transcript (construct N0) does not produce cell growth defects over a 2-month period (Jain and Vale, 2017). Interestingly, however, five constructs induced marked cell death (constructs T1-T5 for toxicity-inducing CAG repeats). The 5’ flanking sequences that produced cytotoxicity had a higher GC-content compared to the flanking sequences that did not induce toxicity (Supp. Fig 1A, Supp. Table 1) but beyond this modest correlation with GC-content, we have not found other sequence motifs in the toxicity inducing flanking sequences. Cellular toxicity did not correlate with RNA expression levels (Supp. Fig. 1B-C) but was dependent on the number of CAG repeats: for a construct that induced cell death (T1), decreasing the number of CAG repeats progressively diminished the toxicity associated with RNA expression (Fig. 1C, Supp. Fig. 1D). RNA with 5×CAG repeats cloned downstream of T1 flanking sequence did not produce discernible defects in cell viability as compared to uninduced controls, indicating that cell toxicity was not trivially imparted by the flanking sequence alone. In summary, our findings reveal that the sequences upstream of CAG repeats substantially impact the cytotoxicity resulting from the expression of expanded repeat-containing RNAs.

We next examined the localization of the repeat-containing RNAs at early time points (24 h after induction) before cell toxicity was evident (Supp. Fig. 1E). In the constructs that did not induce toxicity, the repeat-containing RNA accumulated at nuclear foci (Fig. 1D, Supp. Fig. 1F-I), similar to our previous observation with transcripts harboring CAG-repeats near the 5’ end of the RNA (construct N0) (Jain and Vale, 2017). Surprisingly, we discovered that in the constructs that produced cellular toxicity, the repeat-containing RNA accumulated in peri-nuclear cytoplasmic inclusions (hereafter, called cytoplasmic RNA aggregates; Fig. 1D-F, Supp. Fig. 1J). Reducing the RNA expression level down to a few hundred copies per cell (as assessed by RNA FISH) did not affect the cytoplasmic RNA localization pattern, although the size of the inclusions decreased (Fig. 1G-H; Supp. Fig. 1K). Fluorescence in situ hybridization (FISH) using a probe against CAG repeats or against MS2-hairpins confirmed the accumulation of cytoplasmic RNA aggregates in cells expressing constructs that induced toxicity (Supp. Fig. 1L). Control RNAs without CAG repeats, encoding a fluorescent protein or its reverse complement, that were tagged with MS2 hairpins, did not produce detectable inclusions in the nucleus or the cytoplasm (Fig. 1D-F). Deletion of the MS2-hairpin tag did not appreciably affect RNA localization (Supp. Fig. 1M), indicating that the tag was not the cause of cytoplasmic RNA accumulation and aggregation (see also, Supp. Discussion). Cytoplasmic aggregates formed whether the toxicity-inducing repeat sequence (T1) was introduced through lentiviral transductions, by piggyBac transposase-mediated integration, or via transient transfections (Supp. Fig. 1N). These results indicate that upstream flanking sequences can profoundly alter the sub-cellular localization of the expanded CAG repeat-containing RNAs, and cell toxicity correlates with the accumulation of RNA in cytoplasmic aggregates.

We examined how the native sequences that occur upstream of CAG repeats in *HTT*, *ATXN3* and *ATXN8* genes, associated with Huntington’s disease, spinocerebellar ataxia type 3, and spinocerebellar ataxia type 8 respectively, affect the localization of the CAG repeat-containing RNA. These native sequences were placed upstream of expanded CAG repeats and separated from the repeats by multiple stop codons in each frame, as previously described (Zu et al., 2011). Previous work has also shown that the expression of these repeat-containing constructs triggers cell apoptosis (Zu et al., 2011). In all three cases, we observed that a substantial fraction of cells that expressed these repeats exhibited cytoplasmic RNA aggregates whereas cells transfected with control plasmids did not (Fig 1I). Thus, disease-causing repeats occur in sequence contexts that promote nuclear export and cytoplasmic RNA aggregation.

### Cytoplasmic RNA aggregates are gel-like and deform the nuclear envelope

We previously reported that the nuclear foci formed by CAG repeat-containing RNA are liquid-like (Jain and Vale, 2017). Consistent with liquid-like behavior, RNA at the nuclear foci is mobile, and exhibited rapid fluorescence recovery after photobleaching (characteristic recovery time, τ = 60 ± 28 s, n = 23 foci across constructs N1 and N2; Fig. 2A-B; Supp. Movie 1). In contrast to the globular nuclear foci, cytoplasmic RNA aggregates have a dendritic morphology (Fig 1D). The RNA clustered in the cytoplasmic aggregates was immobile, and upon fluorescence photobleaching, we did not observe appreciable recovery over a 5 min period (<15% recovery, n = 24 aggregates across constructs T1 and T2, Fig. 2A-B, Supp. Movie 2). Cells with cytoplasmic RNA aggregates often displayed a deformed nuclear morphology (Fig. 2C-D), and in numerous cases, the nuclear envelope was indented in the regions adjoining the RNA aggregates (Fig. 2E). Thus, nuclear and cytoplasmic inclusions formed by CAG repeat-containing RNA exhibit markedly distinct biophysical properties, with the cytoplasmic aggregates being solid-like and capable of deforming the nucleus.

**Figure 2.**
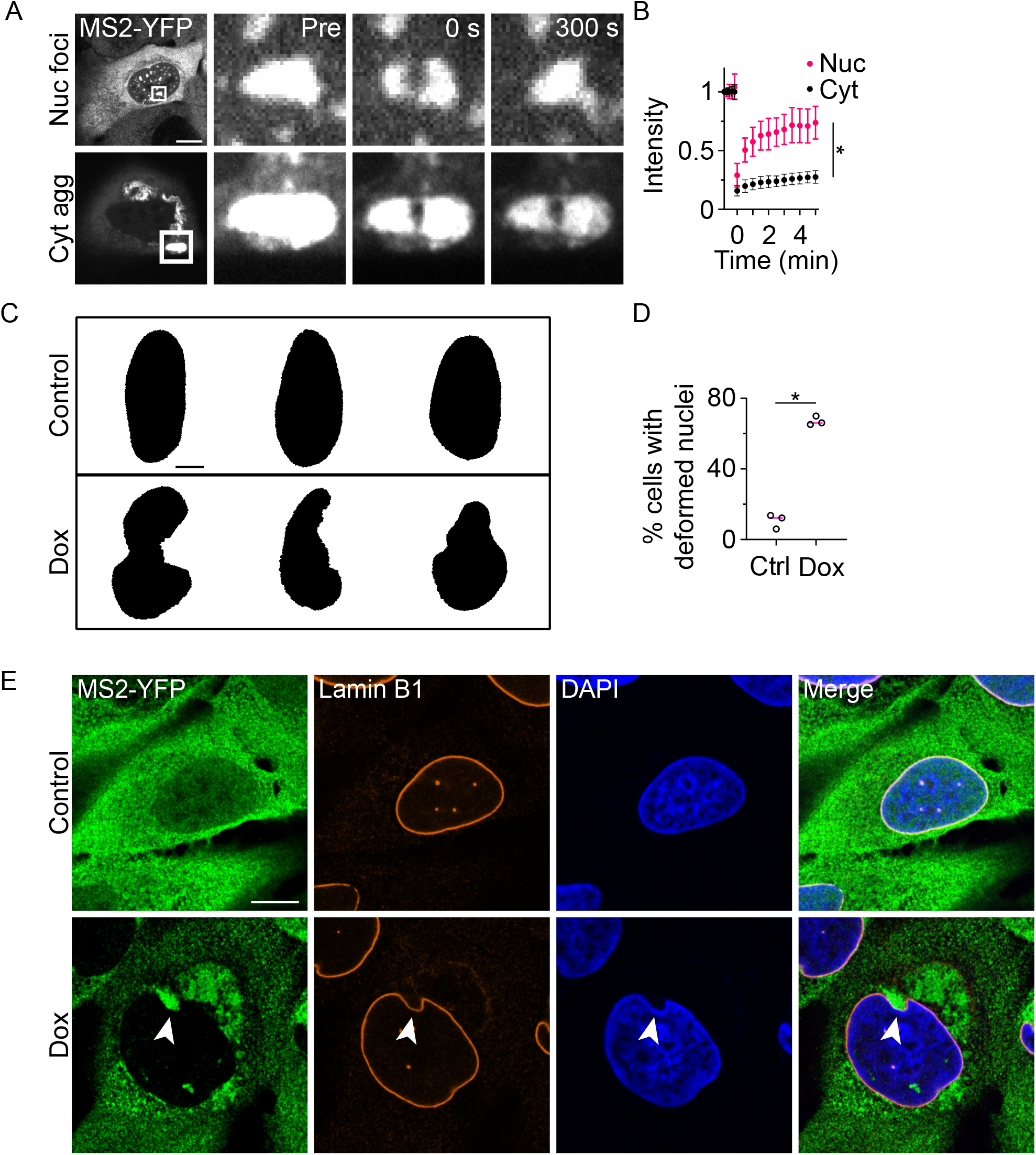
Cytoplasmic RNA inclusions are solid-like and deform the nucleus **A,** Representative micrographs of a nuclear (top) and cytoplasmic (bottom) inclusion before (pre) and at the stated timepoints after photobleaching. **B,** FRAP trajectories are shown for each inclusion. Data are summarized as mean ± s.d., n ≥ 20 events. Asterisks denote significance by Wilcoxon signed-rank test. **C,** Representative masks of nuclei depicting nuclear envelope deformation in cells with cytoplasmic RNA aggregates. **D,** Quantification of the percent of cells with deformed nuclei. Each data point represents an independent experiment with each experiment consisting of ≥ 50 cells. Asterisks denote significance by Student’s t-test. **E,** Immunofluorescence micrographs showing localization of 240×CAG RNA (MS2-YFP) and lamin B1. Arrowheads denote representative point of nuclear deformation. Immunofluorescence data are representative of 3 independent experiments, each of ≥ 30 cells. Scale bars, 10 µm for A, E and 5 µm for C. *: p < 0.01, n.s.: not significant (p > 0.01).

### Cytoplasmic RNA aggregation induces mislocalization of TDP-43, FUS and p62

We next evaluated the localization of cytoplasmic RNA aggregates in comparison to markers for known cytoplasmic RNA granules. Canonical markers for stress granules (G3BP1) (Tourrière et al., 2003), P-bodies (EDC4) (Yu et al., 2005), and TIS granules (TIS11b) (Ma and Mayr, 2018) did not appreciably enrich at the cytoplasmic RNA aggregates (Fig. 3A-B, Supp. Fig. 2A), suggesting that the aggregates of CAG repeat-containing RNA are distinct from these RNA granules.

**Figure 3.**
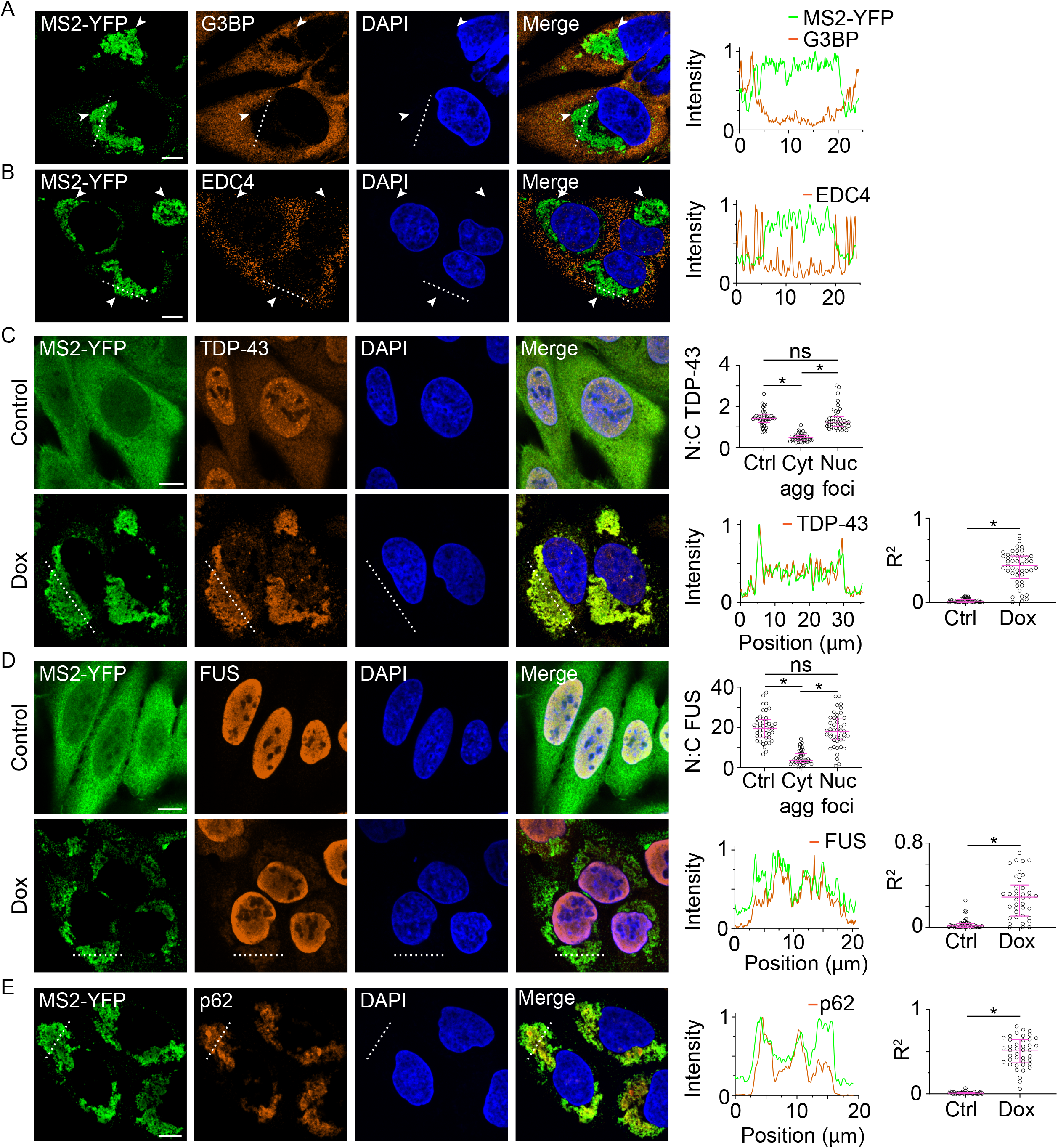
Cytoplasmic RNA inclusions co-localize with RNA binding proteins. **A-B,** Immunofluorescence micrographs (left) and representative line profiles (right) showing localization of 240×CAG RNA (MS2-YFP) and G3BP1/2 (A) and EDC4 (B) in cells with cytoplasmic inclusions. Arrowheads are used as visual guides to indicate representative RNA aggregates. Nuclei were counterstained with DAPI (blue). **C, D,** Same as A-B except showing TDP-43 (C) and FUS (D) in induced and uninduced controls. Ratio of nuclear:cytoplasmic TDP-43 (C) and FUS (D) as calculated from immunofluorescence micrographs (right, top). Controls are uninduced cells that contain constructs capable of forming cytoplasmic aggregates. Each point represents a single cell and data are summarized as median ± interquartile range. Asterisks denote significance by Mann-Whitney test. Line profiles as in A-B (right, bottom). Correlation coefficient, R^2^, for colocalization between MS2-YFP and TDP-43 (C) or FUS (D) (far right). Each point represents a single cell and data are summarized as median ± interquartile range. Asterisks denote significance by Mann-Whitney test. **E,** Same as A-B except showing localization of p62 in cells with cytoplasmic inclusions. R^2^ values (far right) as in C-D. All immunofluorescence data are representative of ≥ 2 independent experiments, each of ≥ 40 cells. All scale bars, 10 µm. *: p < 0.01, n.s.: not significant (p > 0.01).

The morphology and the localization of the RNA aggregates were similar to those of cytoplasmic inclusions that are observed in neurodegenerative diseases including Huntington’s disease and ALS (Gomez-Deza et al., 2015; Liu et al., 2015). TDP-43, normally a predominantly nuclear RNA binding protein, mislocalizes to the cytoplasm in several neurodegenerative diseases (Amador-Ortiz et al., 2007; Neumann et al., 2006; Schwab et al., 2008). We found that cytoplasmic levels of TDP-43 were significantly higher in cells with RNA aggregates than in uninduced controls (Fig. 3C; nuclear to cytoplasmic TDP-43 ratio 0.49 ± 0.2 compared to 1.4 ± 0.4 in uninduced controls, p < 0.0001 by Mann-Whitney test, n ≥ 40 cells). Further, TDP-43 was enriched at the cytoplasmic RNA aggregates (Fig. 3C; R^2^ = 0.45, compared to R^2^ = 0.01 for uninduced cells where R is the correlation coefficient between the two images, p < 0.0001 by Mann-Whitney test, n ≥ 40 cells). FUS, another RNA binding protein that forms aggregates in several repeat expansion disorders (Deng et al., 2010), also mislocalized to the cytoplasm and was sequestered at the RNA aggregates (Fig. 3D; nuclear to cytoplasmic ratio 3.6 ± 3.5 compared to 19.7 ± 6.9 in uninduced controls, p < 0.0001 by Mann-Whitney test, n ≥ 40 cells; R^2^ = 0.29, compared to R^2^ = 0.01 for uninduced controls, p < 0.0001 by Mann-Whitney test, n ≥ 38 cells). Neither TDP-43 nor FUS mislocalized in cells with nuclear foci only, and the nuclear to cytoplasmic ratio of these proteins in cells with RNA foci was comparable to that observed in uninduced controls (Fig. 3C-D, Supp. Fig. 2B). We also found that p62, a canonical biomarker for protein aggregation and autophagy (Zatloukal et al., 2002), was substantially enriched at the RNA aggregates (Fig. 3E; R^2^ = 0.52, compared to R^2^ = 0.01 for uninduced controls, p < 0.0001 by Mann-Whitney test, n ≥ 40 cells). In summary, cytoplasmic aggregates of CAG repeat-containing RNA co-localize with proteins that are frequently mislocalized in repeat expansion diseases.

### Cytoplasmic RNA aggregation coincides with RAN translation of the repeat-containing RNA

RNA with expanded CAG-repeats has the potential to undergo non-canonical RAN translation (Cleary and Ranum, 2013). The repetitive proteins produced upon translation of repeat-expansion RNAs are aggregation-prone, and these aggregates recruit biomarkers for protein aggregation such as p62 (Donaldson et al., 2003; Mori et al., 2013; Nagaoka et al., 2004). We therefore tested whether cells with cytoplasmic RNA inclusions produce RAN translation products. The stop codons upstream of repeats should preclude translation readthrough from the flanking sequences and any polyglutamine production (encoded in the CAG frame) likely results from non-canonical translation of the repeats. Western blots using an antibody targeting polyglutamine revealed an ∼8-fold increase in polyglutamine-containing proteins in cells with cytoplasmic aggregates compared to uninduced controls (Fig. 4A). In contrast, cells with only nuclear RNA foci did not exhibit significant polyglutamine production (Fig. 4A). Immunostaining using a polyglutamine antibody confirmed an increase in production of polyglutamine-containing proteins in cells with cytoplasmic aggregates. Interestingly, immunolabeling also revealed that polyglutamine staining co-localized with the CAG repeat-containing RNA (Fig. 4B-C, Supp. Fig. 3A; R^2^ = 0.35 compared to R^2^ = 0.01 for uninduced cells, p < 0.0001 by Mann-Whitney test, n ≥ 40 cells). This result suggests that RAN translation products are co-aggregating with CAG repeat-containing RNA.

**Figure 4.**
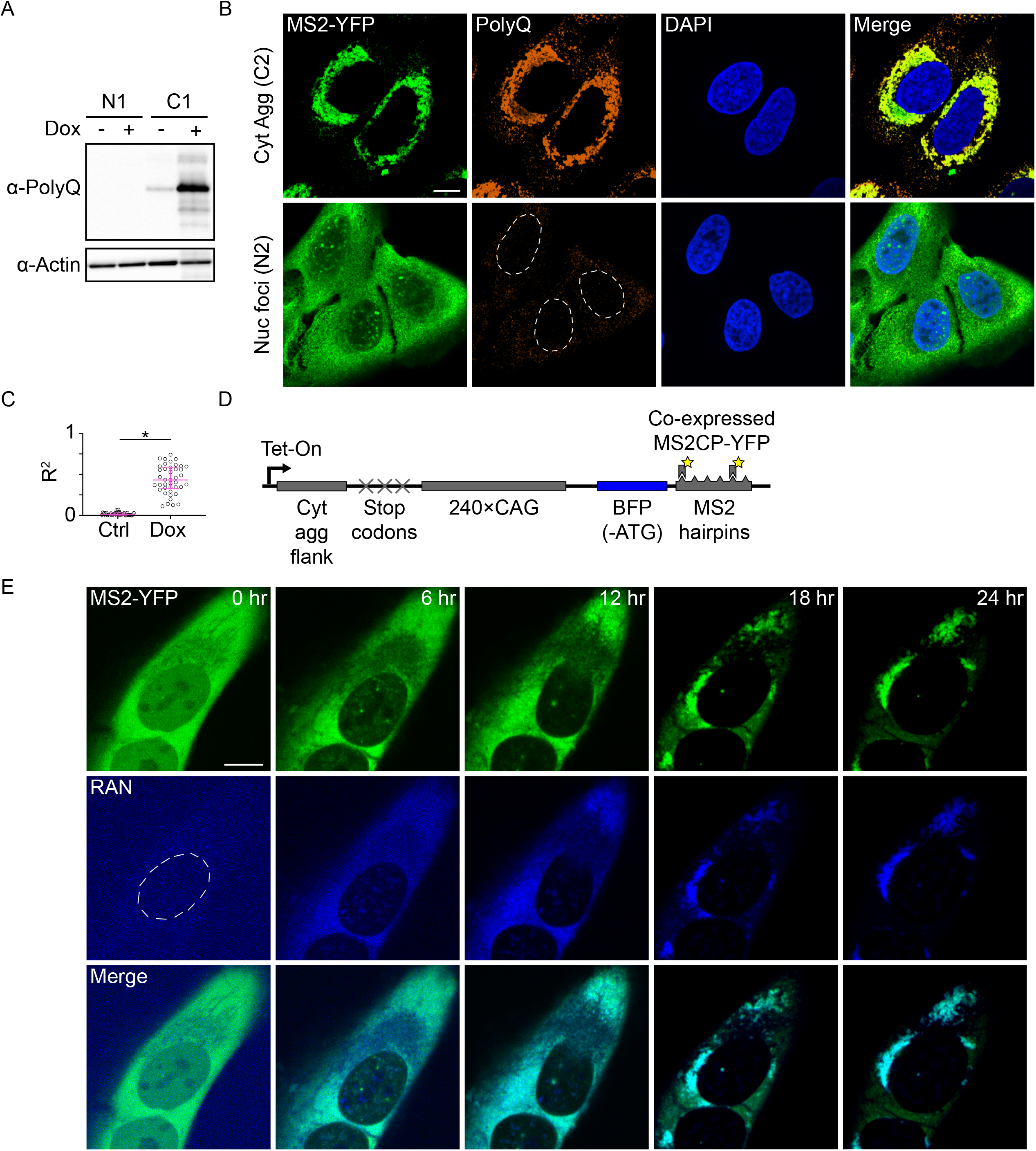
CAG repeat-containing RNA that accumulate at cytoplasmic inclusions undergo RAN translation and co-aggregate with their RAN translation products. **A,** Immunoblot with an anti-polyglutamine antibody, with actin as a loading control. **B,** Immunofluorescence micrographs showing localization of 240×CAG RNA (MS2-YFP) and polyglutamine-containing proteins (PolyQ). Nuclei were counterstained with DAPI. PolyQ channels are equally scaled. **C,** Quantification of R^2^ values for colocalization between MS2-YFP and polyglutamine. Each point represents a single cell and data are summarized as median ± interquartile range. Asterisks denote significance by Mann-Whitney test. **D,** Schematic of RAN translation reporter. **E,** Fluorescence micrographs of RNA (MS2-YFP) and RAN translation products (RAN) at the stated timepoints after induction. All scale bars, 10 µm. Data are representative of two experiments for western blots and at least three experiments for immunofluorescence and live imaging experiments.

Large cytoplasmic inclusions of RAN-translation products may promiscuously sequester cellular macromolecules. To evaluate the progression of RAN translation with respect to RNA aggregation, we developed a live-cell reporter of RAN translation by placing a blue fluorescent protein, EBFP2, lacking an ATG-start codon, immediately downstream of the CAG-repeats (Fig 4D). We chose EBFP2 as our reporter of choice as it does not contain another in-frame methionine in the first 78 amino acids and translation initiation at this residue should not result in a fluorescent product (Heim et al., 1994). RAN translation, which likely initiates within or upstream of the repeats, will read through the downstream EBFP2, resulting in fluorescent protein production in one of the frames. Cells transduced with this construct exhibited BFP fluorescence upon induction (Fig. 4E). Incorporation of stop codons between the CAG-repeats and the EBFP2 led to a near complete loss of BFP fluorescence (Supp. Fig. 3B) validating that the translation of EBFP2 initiates within or upstream of the CAG-repeats.

Using this dual-reporter cell line, we monitored repeat RNA localization and RAN translation in real-time. Upon transcription induction, no appreciable BFP fluorescence or RNA aggregation was observed at early time points (Fig. 4E). After ∼12 h, we observed an increase in BFP fluorescence (Fig. 4E, Supp. Fig. 3C, Supp. Movie 3). Intriguingly, BFP fluorescence appeared prior to any apparent RNA aggregation. Over time, the BFP-tagged RAN translation products co-aggregated with the RNA throughout the cytoplasm, and these RNA-protein co-clusters gradually coalesced into larger perinuclear aggregates. Given that the appearance of BFP fluorescence occurs ∼30 min after translation (due to folding and maturation (Balleza et al., 2018)), these data suggest that RAN translation starts prior to observable RNA aggregation, and that the RAN-translation products likely aggregate co-translationally or soon after translation with the repeat-containing RNA.

### Inhibition of RAN translation abrogates RNA aggregation and rescues cytotoxicity

Finally, we investigated whether translation of CAG-repeats was required for RNA aggregation. Treatment with a translation elongation inhibitor, cycloheximide (10 µg/mL), reduced the levels of both RAN translation products as well as RNA aggregates by >90% compared to untreated controls (Fig. 5A-B, Supp. Fig. 4A; p < 0.0001 by Mann-Whitney test for each, n ≥ 40 cells). Cycloheximide did not substantially affect RNA stability (Supp. Fig. 4B). The repeat-containing RNA was still exported to the cytoplasm in cycloheximide-treated cells but did not form any observable inclusions (Supp. Fig. 4B).

**Figure 5.**
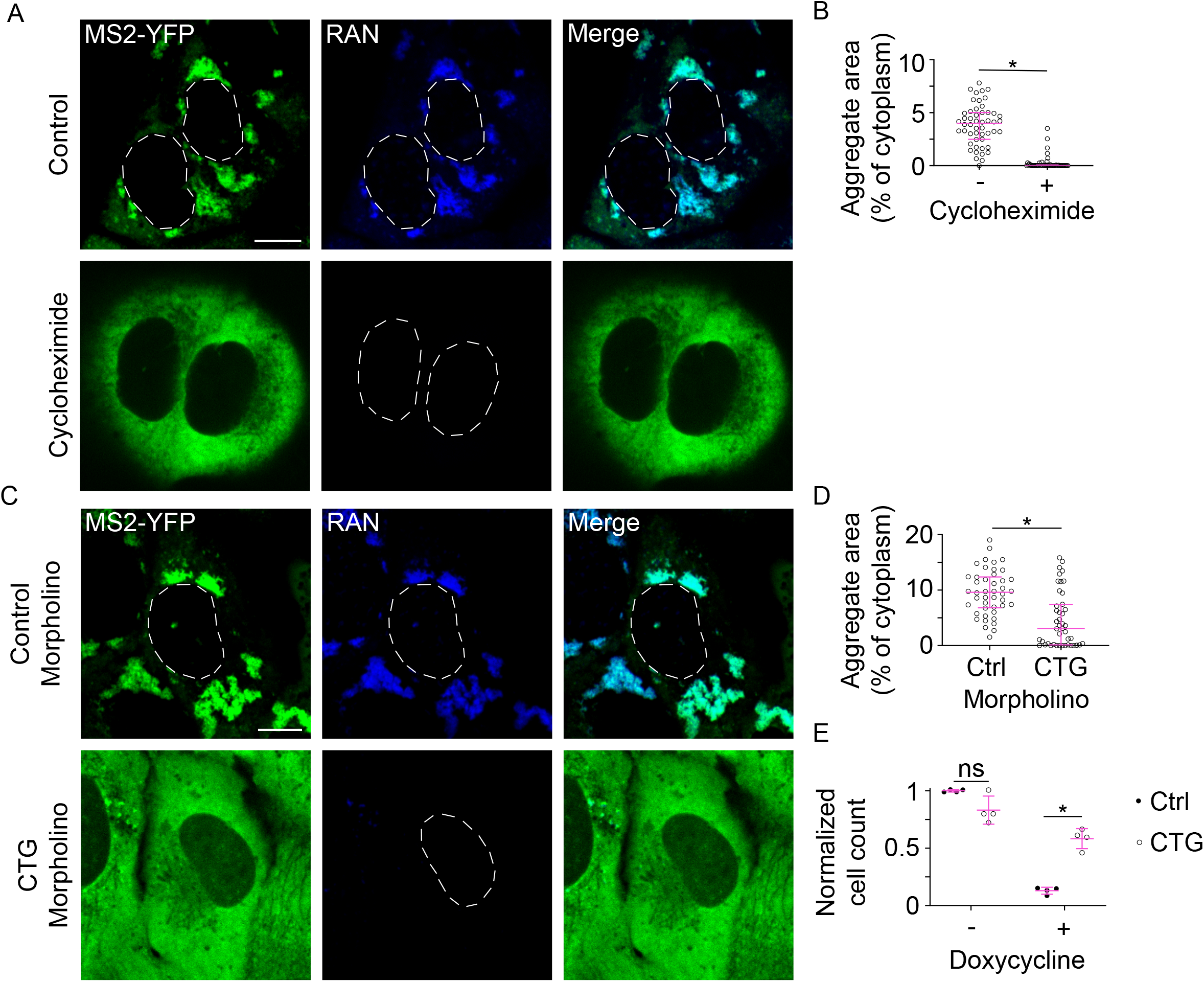
Translation is required for CAG repeat RNA aggregation and toxicity. **A,** Micrographs of cells expressing the RAN translation reporter construct with doxycycline alone (Control) or with doxycycline plus cycloheximide (Cycloheximide). Micrographs depict RNA (MS2-YFP) and RAN translation products (RAN). **B,** Quantification of aggregate area in control and cycloheximide treated cells. Each data point represents a single cell; data are summarized as median ± interquartile range. Asterisks denote significance by Mann-Whitney test. **C-D,** Same as A-B except showing cells treated with doxycycline and either a control morpholino or 8×CTG morpholino. **E,** Quantification of cell survival after four days of induction with either a control morpholino or 8×CTG morpholino. Each data point represents one experiment, and data are summarized as the mean ± s.d. Asterisks denote significance by Student’s t-test. Data are representative of two experiments for cycloheximide treatments and three experiments for morpholino treatments, each with ≥ 40 cells. All scale bars, 10 µm. *: p < 0.01, n.s.: not significant (p > 0.01).

Since global translation inhibition itself exerts substantial toxicity, we tested whether targeted translation inhibition of CAG-repeat-containing RNA could prevent aggregation and toxicity. We designed a phosphorodiamidate backbone 8×CTG morpholino that can hybridize with the CAG-repeats. Morpholinos sterically inhibit protein translation without affecting the stability of the target RNA (Summerton, 2007). Treatment with 8×CTG morpholino suppressed RAN translation and RNA aggregation (reduction by 43% and 71%, respectively compared to treatment with control morpholinos, p < 0.001 by Mann-Whitney test, n ≥ 40 cells) while only modestly affecting RNA levels (Fig 5C-D, Supp. Fig. 4C-D; 29% reduction in RNA compared to cells treated with control morpholino, n ≥ 40 cells). 8×CTG morpholino treatment also rescued cell viability. The expression of toxic CAG-repeat RNAs led to >85% cell death after 4 days in cells treated with a control morpholino, compared to <35% cell death upon treatment with the 8×CTG morpholino (Fig. 5E). In summary, inhibition of RAN translation is sufficient to mitigate both cytoplasmic RNA aggregation and cytotoxicity associated with CAG repeat RNA.

## Discussion

Nuclear foci of repeat-containing RNA were first observed in myotonic dystrophy (Taneja et al., 1995) and have been subsequently described in nearly all repeat expansion diseases (Wojciechowska and Krzyzosiak, 2011). We demonstrate that, in addition to aggregating in the nucleus, repeat-containing RNA also aggregate in the cytoplasm. The formation of cytoplasmic RNA aggregates coincides with RAN translation, and these aggregates are the sites for the accumulation of several RNA binding proteins as well as markers for protein aggregation. We propose that this cytoplasmic aggregation of repeat-containing RNA may provide a mechanistic explanation for the aberrant cytoplasmic localization of RNA binding proteins such as TDP-43 which is commonplace in repeat expansion disorders (Goodwin and Swanson, 2014; Todd and Paulson, 2010).

Why have cytoplasmic RNA aggregates not been detected in previous RNA localization studies in repeat expansion diseases? We found that commonly used chemical cross-linking fixatives, such as formaldehyde, reduced the detection of RNA at cytoplasmic aggregates via FISH. While both nuclear foci and cytoplasmic aggregates of RNA were observed in methanol-fixed cells, only nuclear foci were stained in cells fixed with formaldehyde (Supp. Fig 5A). Even in cells where RNA detection by MS2CP-YFP revealed abundant cytoplasmic inclusions, RNA aggregates were not readily observed via FISH after formaldehyde fixation (Supp. Fig. 5A). Chemical cross-linking of proteins and RNA may prevent probes from accessing the aggregated RNA. Formaldehyde treatment can also result in the addition of mono-methylol groups to RNA bases – particularly adenine and cytosine residues which are enriched in CAG repeat RNA – and chemically cross-link RNA molecules (Masuda et al., 1999). These findings suggest that optimization of FISH protocols may allow for better detection of cytoplasmic RNA inclusions. Consistent with this notion, a recent study reported a substantially greater number of cytoplasmic inclusions of GGGGCC- and CGG-repeat-containing RNAs when using hybridization chain reaction to amplify RNA FISH signal compared to the traditional methods (Glineburg et al., 2021).

Previous studies have described peri-nuclear cytoplasmic protein inclusions in a variety of CAG repeat expansion disorders such as Huntington’s disease and spinocerebellar ataxia 8 (Ayhan et al., 2018; Waelter et al., 2001). Our data suggest that these inclusions likely contain the repeat-containing RNA which may be revealed by using milder fixatives. Co-aggregation of RNA may also be a common feature of protein aggregation disorders that do not involve repeat-expansions. Early studies on amyloid beta plaques in Alzheimer’s disease showed that these aggregates stain with RNA-binding dyes (Ginsberg et al., 1997), and a recent study found that tau aggregates contain RNA (Lester et al., 2021). Modifications in tissue storage and fixation methods to retain integrity and advancements in RNA detection technologies may allow one to determine the true prevalence of RNA aggregates in neurodegenerative disease.

Our data directly demonstrate that RAN translation is more toxic to cells than the accumulation of repeat-containing RNAs at nuclear foci. RNAs that coalesced exclusively at nuclear foci did not induce substantial cytotoxicity in our cell culture model, and inhibiting RAN translation in cells where repeat-containing RNA was exported to the cytoplasm, was sufficient to mitigate cell death. Genetic screens have identified several RNA export-factors such as NXF1 and NXT1 as well as nuclear pore components as potential regulators of repeat expansion induced toxicity (Cheng et al., 2019). Previous work from the Ranum lab showed that over-expression of certain RNA binding proteins like MBNL1 may increase retention of RNA in the nucleus and consequently inhibit RAN translation and cell death (Zu et al., 2017). Our findings, based upon a tissue culture model, raise the possibility that sequestration of RNA in the nucleus at foci may be protective to cells, and if so, may provide a therapeutic intervention strategy to reduce disease burden. However, we cannot exclude the possibility that prolonged accumulation of repeat-containing RNA in nuclear foci also interferes with cell function, such as in myotonic dystrophy where foci formation is associated with various splicing defects (Jiang et al., 2004).

Our results also underscore the role of cis-acting sequences in mediating disease. Diseases caused by CAG trinucleotide repeat expansion share several pathobiological features. However, at a given repeat number, these diseases differ considerably in the age of disease onset and severity of symptoms (Gusella and MacDonald, 2009). Previous studies have shown that the surrounding sequence context may influence repeat instability (Brock et al., 1999). Likewise, when the repeats are translated, the sequence context influences the aggregation propensity of resultant polyglutamine-containing protein (Duennwald et al., 2006). Our results unequivocally show that besides affecting repeat instability and protein aggregation, flanking sequences, in combination with repeat length, modulate RNA toxicity. We found that GC-rich flanking sequences were more likely to induce RNA aggregation and RAN translation. Interestingly, recent genome-wide screens for modulators of RAN translation have identified DDX3X, a dead-box helicase that preferentially binds GC-rich UTRs as a key modulator of RAN translation (Cheng et al., 2019; Linsalata et al., 2019). The assays that we have developed in this study may provide means to screen for sequence motifs that modulate RNA export and RAN translation.

Our work provides a new lens to examine the role of repeat-containing RNA in the cellular pathology of repeat expansion diseases. Disease-associated repeat expansions in RNA provide sites for multivalent intermolecular base-pairing. Such multivalent interactions promote RNA oligomerization and its accumulation in the nucleus at foci (Jain and Vale, 2017). The sequences flanking the repeats may potentially provide binding sites for yet-unknown RNA helicases and export factors. In the cytoplasm, the repeat-containing RNA encounters the translation machinery and may undergo non-canonical RAN translation. Interestingly the cytoplasmic RNA does not appear to form aggregates until it is RAN translated, suggesting that the repeat-containing RNA in the cytoplasm is initially in a state that inhibits extensive inter-molecular RNA-RNA interactions. The process of RAN translation may relieve this repression. The RAN peptides are also aggregation-prone and another possibility is that the repeat-containing RNA may co-translationally aggregate with the RAN peptides. The smaller RNA-RAN protein aggregates are deposited, possibly by the minus-end directed motor dynein, into large perinuclear, aggresome-like inclusions (Johnston et al., 1998). These cytoplasmic inclusions sequester various RNA binding proteins, disrupt nuclear morphology, and over time, kill cells. Future work may reveal the factors involved in repeat-RNA export, RAN translation, and the mechanism of RNA-RAN-protein co-aggregation.

## Supporting information

Key Resources Table

Supplemental Legends and Discussion

## Author Contributions

M.D., Y.C., R.V., and A.J. designed research; M.D., Y.C., R.S., N.Z., C.T., and A.J. performed research; M.D. and A.J. analyzed data; M.D., R.V., and A.J. wrote the paper.

## Acknowledgements

We thank members of the Jain lab for helpful discussions. Plasmids containing CAG repeats with flanking sequences from *ATXN3*, *ATXN8*, and *HTT* were a gift from Laura Ranum. This work was supported by grants from the National Institutes of Health (R00AG053434), the David and Lucile Packard Foundation, and the Smith Family Awards Program. Y.C. was supported by a fellowship from the National Research Foundation of Korea. R.V. is an investigator with the Howard Hughes Medical Institute.

## STAR Methods

### Resource availability

#### Lead contact

Further information and requests for resources and reagents should be directed to and will be fulfilled by the lead contact, Ankur Jain.

#### Materials availability

Plasmids generated in this study are available upon request to the lead contact.

#### Data and code availability

All data reported in this paper will be shared by the lead contact upon request. This paper does not report original code. Any additional information required to reanalyze the data reported in this paper is available from the lead contact upon request.

### Experimental model and subject details

Immortalized female human cell lines were used in this study. All experiments were performed in U-2 OS cells (ATCC, HTB-96) stably expressing a TetOn3G transactivator protein and an MS2 hairpin binding protein fused to YFP was previously described (Jain and Vale, 2017). HEK293T (ATCC, CRL-3216) cells were used for generating lentivirus. ATCC authenticated cell lines by STR profiling. Cells were grown in DMEM (Life Technologies, 11965-126) supplemented with 10% (v/v) fetal bovine serum (Gibco, 26140-079) and 1× penicillin, streptomycin, and glutamine (Gibco, 10378016). Transgene expression was induced by adding 1 µg mL^-1^ doxycycline (Sigma-Aldrich, D9891) for 24 hours except where otherwise specified. Cells were maintained in a humidified incubator at 37 C with 5% CO2. Representative characterization of cytoplasmic inclusions was conducted in cells expressing construct T1 or T2; representative characterization of nuclear inclusions was conducted in cells expressing either construct N1 or N2.

### Method details

#### Cloning and plasmid preparation

Full-length plasmid sequences are provided in Supp. Table 2. Lentiviral transfer plasmids with CAG-repeats and MS2-hairpins were previously described (Jain and Vale, 2017). The upstream flanking sequences as well as the downstream EBFP2 tag, were obtained as double stranded DNA fragments (Quintara Biosciences or IDT). The library of 12 upstream flanking sequences were inserted via restriction digestion cloning in constructs with 47× or 240× CAG repeats (construct N1 or equivalent) between *Eco*RI and *Mlu*I sites. 5× or 22× CAG repeats were inserted as annealed primers/ultramers (IDT) between *Eco*RI and *Sgr*DI sites downstream of the T1 flanking sequence. EBFP2 was cloned between *Bam*HI and *Not*I sites. All cloning and plasmid preparations were performed in Stbl3 *E. coli* cells (Invitrogen, C7373-03) grown at 30°C. In constructs with 240× CAG repeats, Sanger sequencing did not provide sufficient read length to unambiguously determine the number of repeats, so repeat length was first validated via restriction digestion of the insert and gel electrophoresis. To sequence the repeat tract, we optimized a Sanger sequencing protocol (in collaboration with Quintara Biosciences) which used betaine and 7-deaza-dGTP. This optimized sequencing protocol provided ∼800 base long reads from both ends. Sanger sequencing revealed eight interruptions in the CAG repeat track in constructs with 240× CAG repeats (sequences in Supp. Table 2). These sites contained a deletion of G nucleotide and the first interruption occurred at 42 bases from the start of the repeat tract. These interruptions were present in our parent plasmid (N0), and were common to the constructs with various flanking sequences (N1-N7, T1-T5). We also observed similar cellular phenotypes (*e.g.*, toxicity, cytoplasmic RNA aggregation, and RAN translation) in constructs containing either these repeat interruptions or with pure expanded ≥ 47×CAG repeat tracks.

Lentiviral packaging and envelope constructs were obtained from addgene: pCMV-VSV-G was a gift from Bob Weinberg (Addgene plasmid # 8454; http://n2t.net/addgene:8454; RRID: Addgene_8454) (Stewart et al., 2003); psPAX2 was a gift from Didier Trono (Addgene plasmid # 12260; http://n2t.net/addgene:12260; RRID: Addgene_12260). Plasmids containing CAG repeats with endogenous flanking sequences from *ATXN3*, *ATXN8*, and *HTT* were generously provided by Laura Ranum (Zu et al., 2011).

#### Transfections and lentiviral transductions

Lentivirus was generated using second generation lentiviral packaging system in HEK293T cells. In brief, ∼500,000 HEK293T cells were plated in a 6-well plate until they reached 60-80% confluence. Each well was transfected with 0.5 µg of envelope plasmid (Addgene #8454), 1 µg of packaging plasmid (Addgene #12260), and 2 µg of the transfer plasmid with 8 µL of Lipofectamine LTX (Invitrogen, 15338-100) in 500 µL of Opti-MEM reduced serum media (Gibco, 31985-070). This solution was incubated for 5 min before pipetting onto HEK293T cells in IMDM (Gibco, 12440-053) + 10% FBS + 1× PSG. After three days of incubation, virus was collected by spinning the supernatant for 5 min at 20,000 x g. U-2OS cells were transduced with varying titers of virus in 10 µg/mL polybrene (Millipore, TR-1003-G).

For transient transfections in U-2OS cells, 500 ng of plasmid was mixed with Opti-MEM reduced serum media and 1.5 µL Viafect transfection reagent (Promega, E4981) for a total volume of 50 µL. 10 µL of this mixture was added per well of U-2OS cells in a 96-well glass bottom plate in 100 µL of freshly changed DMEM +FBS + PSG media.

#### Transgene copy quantification

Genomic DNA and total RNA were collected from cells using PureLink Genomic DNA mini kits (Invitrogen, K1820-01) and PureLink RNA mini kits (Invitrogen, 12183018A), respectively, according to the manufacturer’s protocols. RNA was converted to cDNA using SuperScript III reverse transcriptase (LifeTechnologies, 11754050). Transgene genomic DNA or cDNA copies were quantified using real-time quantitative PCR (RT-qPCR) with SYBR green reagents (Applied Biosystems, 4309155) using primers targeting the TetPromoter for DNA and targeting the WPRE for cDNA (Key Resource Table). Transgene copies were normalized to copies of *ACTB* (primers listed in Key Resource Table).

#### Toxicity assays

Cell toxicity was quantified using two methods: (1) counting the number of adhered cells and (2) exclusion of trypan blue dye. (1) Cells were plated on a six well plate at 10,000 cells per well. The following day, cells were induced with 1 µg mL^-1^ doxycycline or left as uninduced controls in a total volume of 2.5 mL. Each cell line was compared to a corresponding uninduced control to account for any differences in plating density between cell lines. Five days post induction (unless otherwise specified), cells were washed five times with DPBS to remove dead cells from the plate, dissociated from the plate using trypsin, and neutralized with growth medium (DMEM + 10% (v/v) FBS) two times in order to collect all cells from the plate. The cells were then pelleted at 500 x g for 3 min and resuspended in growth medium. Cells were counted using a Countess II FL automated cell counter (Invitrogen, AMQAF1000) and disposable cell counting chamber slides (Invitrogen, C10283). At least two technical replicate counts were performed for each sample.

(2) Trypan blue assays were adapted from published protocols (Strober, 2001). Cells were plated on a six well plate and induced with doxycycline the following day. Five days after induction, the supernatant which contains any dead or detached cells was collected. Cells were washed with DPBS and this wash solution was added to the supernatant. After trypsinization, growth medium was added to neutralize trypsin and the collected cells were added to the supernatant/wash solutions. Cells were pelleted at 500 x g for 3 min and pellets were resuspended in 100 uL of DPBS. This resuspension was mixed 1:1 with 0.4% trypan blue (Invitrogen, T10282) for three minutes and cell counts were performed using a Countess II FL automated cell counter as described above. At least two technical replicate counts were performed for each sample.

#### Fluorescence microscopy

Cells were plated in a glass-bottom 96-well plate (Brooks, MGB096-1-2-LG-L) and imaged using a Dragonfly 505 spinning-disk confocal microscope (Andor Technologies) equipped with a piezo Z-stage (ASI) and an iXon Ultra 888 EMCCD camera. Pin-hole size was kept at 40 microns. Z-stacks were acquired with a step size of 0.3 to 0.5 µm. Live cells were imaged in a humidified chamber (OKO labs) maintained at 37° C and 5% (v/v) CO_2_ using a 100x oil immersion objective NA 1.45 (Nikon, MRD01905) (pixel size 121 nm × 121 nm). Fixed cells were imaged at room temperature. BFP and DAPI were excited with a 405 nm laser and fluorescence was collected using a 445/46 bandpass filter. YFP and Alexa488 were imaged using a 488 nm laser and corresponding 521/38 nm band pass emission filter. Cy3 or mCherry labeled samples were imaged using a 561 nm excitation and a 594/43 emission filter. Last, Cy5 or Atto647N labeled samples were imaged using 640 nm laser line and a 698/77 bandpass emission filter. At least 40 cells were imaged in at least two independent experiments.

#### Image analysis and aggregate quantification

All image analysis was performed in FIJI. Background fluorescence was determined from the average signal of ten ROIs (approximately 50 square microns in size) that did not contain any cells. This background signal was determined for each channel in each experiment and subtracted in all analyses.

Cells were manually segmented to analyze signal in the cytoplasm or in the nucleus. To identify cytoplasmic or nuclear inclusions, we empirically determined the intensity and size threshold that adequately recapitulated the features as observed by eye. Cytoplasmic inclusions for MS2-YFP images were identified as objects with 2× the mean intensity of the cytoplasm and a size threshold of 0.1 square microns. A similar thresholding was used for identifying cytoplasmic inclusions by RNA FISH, after appropriate adjustments to the background fluorescence. We classified cells with at least 2% of their cytoplasm occupied by inclusions as having cytoplasmic aggregates. Likewise, nuclear foci were identified as having intensity 1.5-fold the median nucleoplasm intensity and a size threshold at 0.14 square microns. We classified cells with at least 1% of their nucleoplasm occupied by inclusions as having nuclear foci.

In cases where transient transfection was used, we set the size threshold for aggregates as 0.1 square microns and set an intensity threshold at 4× the mean cytoplasmic intensity after background subtraction. We classified cells with at least 0.2% of their cytoplasm occupied by inclusions as having cytoplasmic aggregates. We tested several thresholding parameters, and obtained comparable results (Supp. Fig. 5B).

To determine the nuclear or cytoplasmic immunofluorescence, we manually segmented the cells (based on DAPI and total immunofluorescence signal from the cell), and determined the total integrated fluorescence in nucleus or cytoplasm after subtracting the background. Similar trends in nuclear to cytoplasmic fluorescence were observed when we compared the mean pixel intensities in the two compartments.

For co-localization analysis, line-profiles were obtained using the ‘Plot profile’ plugin in FIJI. Background signal was subtracted from all points and the intensity of each channel was normalized to the maximum intensity along the line. For calculating correlation coefficients, we empirically determined the intensity threshold to classify a pixel as background or foreground. The thresholded images were used to determine per-pixel intensity correlation coefficients.

Thresholds were determined using the sample image, and identical settings were used for the corresponding controls.

To quantify total RNA levels by FISH, average background signal was measured in the nucleus and cytoplasm of uninduced cells. The integrated intensity of each compartment was calculated and summed to find the relative transgene RNA levels in each cell. Similar results were found in confocal and widefield images.

Movies were stabilized using the Image Stabilizer plugin in FIJI (“Kang Li @ CMU - Image Stabilizer Plugin for ImageJ,” n.d.).

#### FRAP imaging and analysis

Regions approximately one square micron in size within cytoplasmic or nuclear inclusions were photobleached using a MicroPoint FRAP module (Andor Technologies) using a 405 nm wavelength photoablation laser. Cells were imaged for 5 frames prior to bleaching, immediately after the bleach (within two seconds), and then in 30 second intervals for five minutes after bleaching. After background subtraction, fluorescence of the bleached region at each timepoint was normalized to the fluorescence of the bleached region prior to photobleaching to determine fluorescence recovery. To correct for photobleaching over the course of imaging, fluorescence of the bleached region was also normalized to the fluorescence of an unbleached region of the aggregate at each timepoint. Unbleached foci were used as a control region to correct for photobleaching due to imaging. FRAP was performed at 24 hours (± 3 hours) after induction with doxycycline. At least 20 FRAP events from two independent experiments were performed for each case. Time constants were determined by fitting the fluorescence recovery to the equation: I = A − I_0_ exp(−t/τ).

#### RNA FISH

Cells were fixed with a solution of 75% (v/v) methanol, 25% (v/v) acetic acid for 10 minutes at 4°C. Fixed cells were washed three times with a wash solution (PBS or nuclease-free water with 300 mM NaCl, 30 mM sodium citrate, 10% (v/v) formamide, and 0.1% (v/v) NP-40 substitute) at room temperature (23°C). Hybridization was performed at 37°C for three hours with 200 nM Cy3-conjugated probes targeting either the CAG repeats or the MS2 hairpin region (probe sequences Key Resource Table). The probes were dissolved in hybridization buffer (100 mg mL^-1^ dextran sulfate, 10% formamide, 300 mM NaCl, and 30 mM sodium citrate in nuclease-free water). After hybridization, cells were washed for 30 minutes with wash solution and counterstained for 30 minutes with wash solution containing 0.5 µg mL^-1^ DAPI; both this wash step and DAPI staining were performed at 37°C. Cells were then washed three times with PBS, and kept in PBS for imaging.

Where indicated, cells were fixed with 2% (w/v) formaldehyde for 10 minutes, followed by a similar protocol to that described above, except that the probes were hybridized for 16 hours. Detection of cytoplasmic inclusions was substantially impeded after formaldehyde treatment (see Discussion).

#### Immunofluorescence

Cells were fixed with 2% formaldehyde in PBS for 40 minutes, washed four times with PBS, and then permeabilized with 0.1% (v/v) Triton-X-100 in PBS for 10 minutes at room temperature. Cells were then blocked in 0.45 µm-filtered 3% (w/v) bovine serum albumin (BSA; Sigma-Aldrich, A7906) in PBS. Primary antibodies (Key Resource Table) were diluted 1:100 in 1% (w/v) BSA in PBS and incubated with cells for one hour at room temperature. Cells were washed three times with PBS and then incubated with the appropriate secondary antibody (Key Resource Table) diluted 1:2000 in 1% (w/v) BSA (in PBS) for one hour at room temperature. After three washes with PBS, cells were counterstained with DAPI solution (PBS containing 0.5 µg mL^-1^ DAPI) for three minutes, washed three times with PBS again, and then imaged as previously described. To enhance MS2CP-YFP signal after fixation, cells were incubated with an anti-GFP antibody and a subsequent Alexa488-conjugated secondary antibody. These incubations were performed along with other antibodies (*i.e.*, cells were simultaneously incubated with anti-G3BP and anti-GFP primary antibodies). At least two independent experiments were performed for all immunofluorescence experiments.

#### Western blots

Cells were washed with PBS and then lysed with RIPA lysis buffer (25 mM Tris-HCl pH=7.5, 150 mM NaCl, 1% (v/v) NP-40, 1% (w/v) sodium deoxycholate, 0.1% (w/v) SDS) with protease and phosphatase inhibitors (Thermo Scientific, 78442). The lysate was homogenized by passing through a 22-gauge syringe 10 times and then incubated on ice for 30 min with vortexing every 5 min. Cell debris was removed by centrifugation at 500 x g for 10 min at 4℃ and lysates were mixed with 4× Bolt™ LDS sample buffer (Invitrogen, B0007) with 50 mM DTT and heated at 70℃ for 10 min. Samples were loaded onto a Bolt™ 4-12% Bis-Tris polyacrylamide gel (Invitrogen, NW04122) and transferred to a PVDF membrane using iBlot 2 dry blotting system (Invitrogen, IB21001). Membranes were blocked in 5% (w/v) skim milk in TBST (tris-buffered saline with 0.1% (v/v) Tween-20) for one hour at room temperature. Membranes were then incubated with primary antibodies (Key Resource Table) that were diluted 1:1000 in 1% (w/v) skim milk in TBST at 4°C overnight. After three TBST washes, membranes were incubated with horseradish peroxidase-conjugated secondary antibodies at a 1:10,000 dilution in 1% (w/v) skim milk in TBST for one hour at room temperature (Key Resource Table). Membranes were washed three times with TBST and chemiluminescence signals were detected using SuperSignal West Femto Maximum sensitivity substrate (Thermo Scientific, 34095) on a ChemiDoc XRS+ imager (Bio-Rad). Band intensities were quantified using FIJI after subtracting background signal.

#### Translation inhibition

Where indicated, cells were treated with 10 µg mL^-1^ cycloheximide (Sigma-Aldrich, C1998). 1 µg mL^-1^ doxycycline was concurrently added to induce transgene expression. Cells were then imaged as previously described 14 h post treatment. Cells transduced with a high viral titer were used in these experiments so that aggregates began to form prior to substantial cell death due to the general toxicity of cycloheximide. Two independent experiments were performed. Targeted translation inhibition was achieved through addition of 8×CTG morpholinos (GeneTools). Control experiments were performed using a standard intron control morpholino (CCTCTTACCTCAGTTACAATTTATA; GeneTools), targeting a mutated intron of the human beta-globin gene. In each case, 50 µM morpholino were delivered with 6 µM EndoPorter peptide (GeneTools, OT-EP-PEG-1) for 24 h prior to induction with doxycycline. Morpholinos were kept in the media after doxycycline addition giving a total treatment time of 48 h with morpholinos and 24 h of treatment with doxycycline. For toxicity assays, cells were plated at 5,000 cells/well in 24-well plates in a total volume of 300 µL. Cells were treated with 50 µM 8×CTG or control morpholinos for five days and with 10 ng/mL doxycycline for four days.

### Quantification and statistical analyses

Cell viability, qPCR, and quantifications of percent of cells exhibiting a phenotype (*e.g.*, nuclear deformation or the presence of inclusions) were analyzed by two-tailed unpaired Student’s t-tests. Morpholino rescue of cell viability was analyzed by two-way ANOVA in addition to t-tests. FRAP experiments were analyzed using Wilcoxon signed-rank test. All other experiments were analyzed by Mann-Whitney U-tests. All statistical tests are described in figure legends including the test used, minimum values of n, definitions of center, and precision measurements.

