## Supplementary figures and images for "Flanking sequences regulate the toxicity, non-ATG translation, and aggregation of RNA with tandem CAG Repeats"

### Supp. Fig. 1

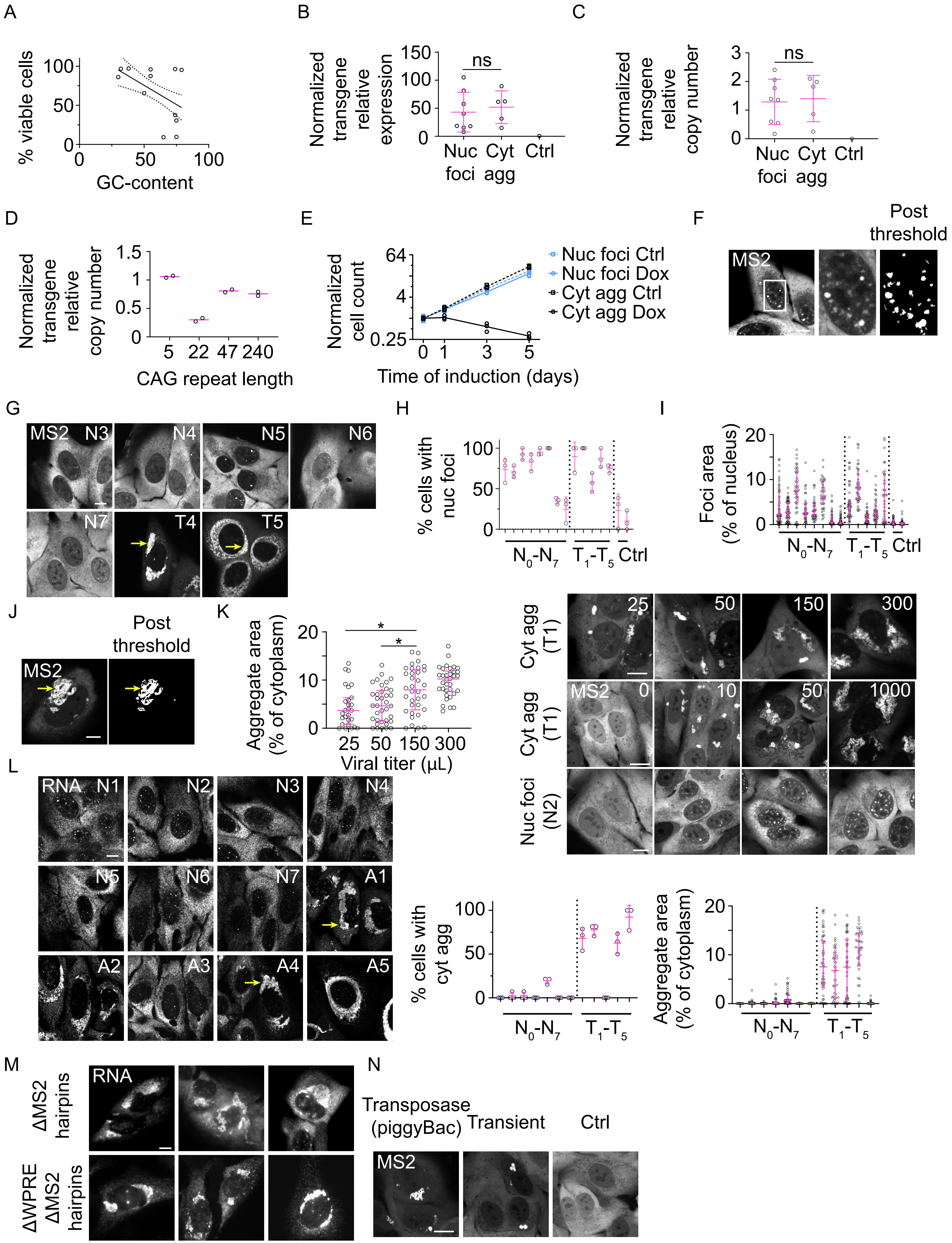

### Supp. Fig. 2

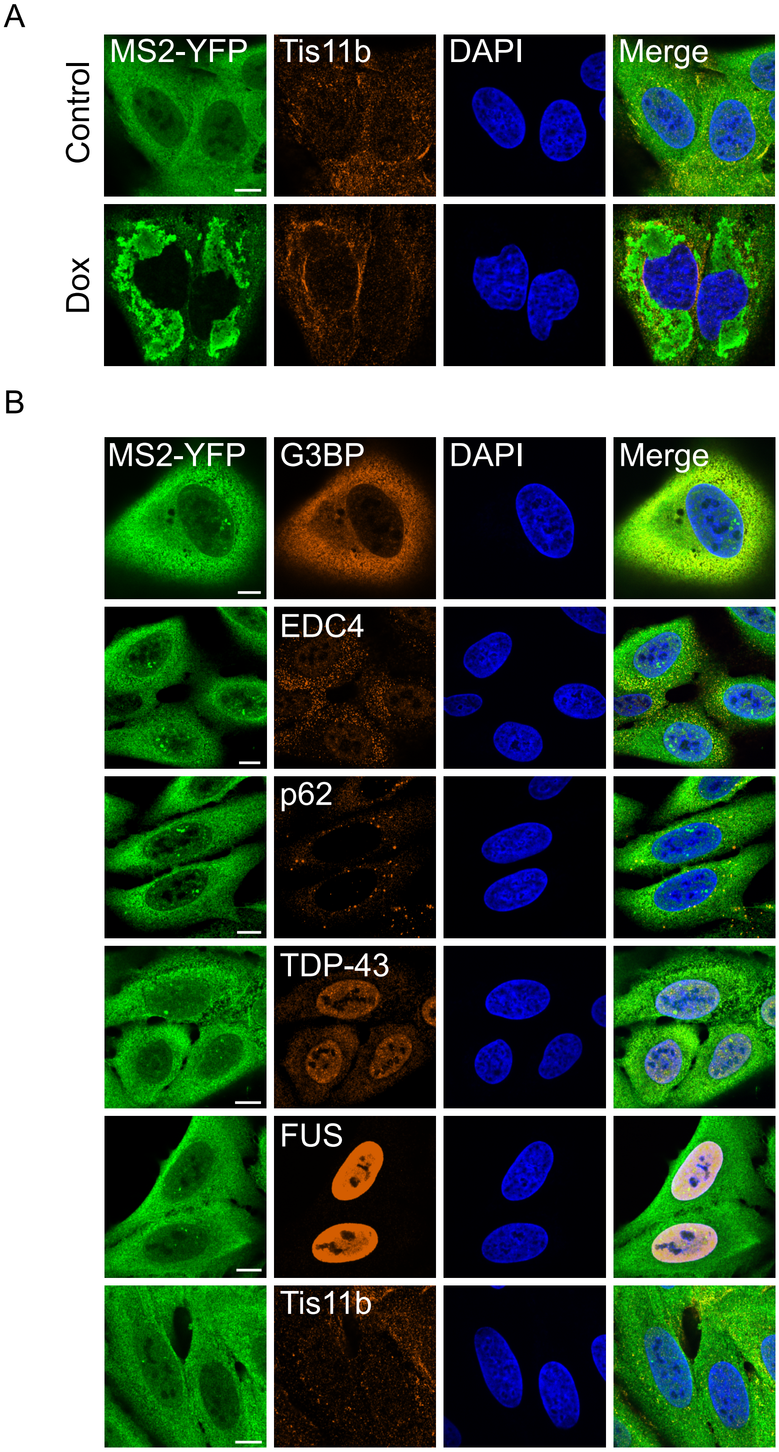

### Supp. Fig. 3

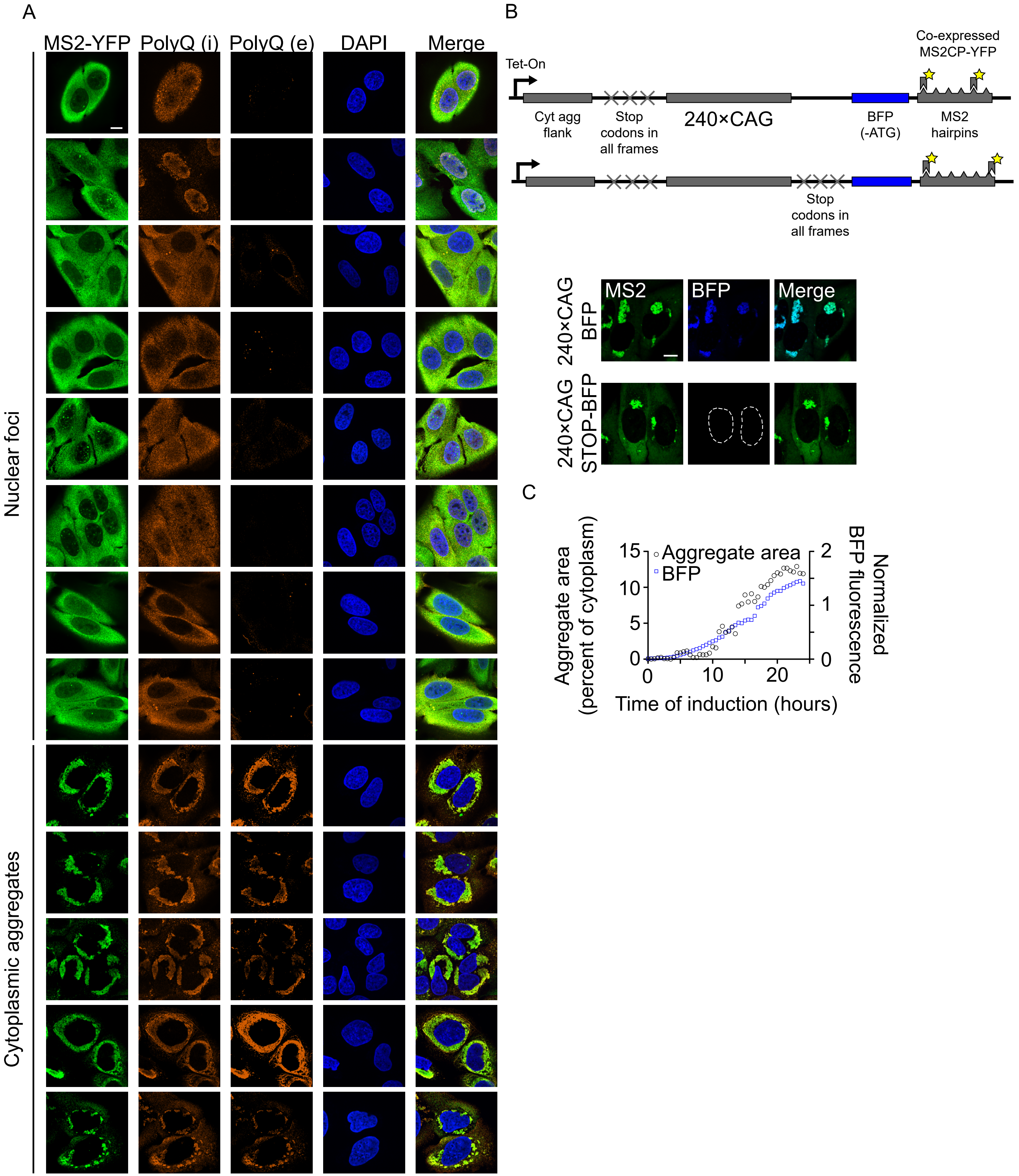

### Supp. Fig. 4

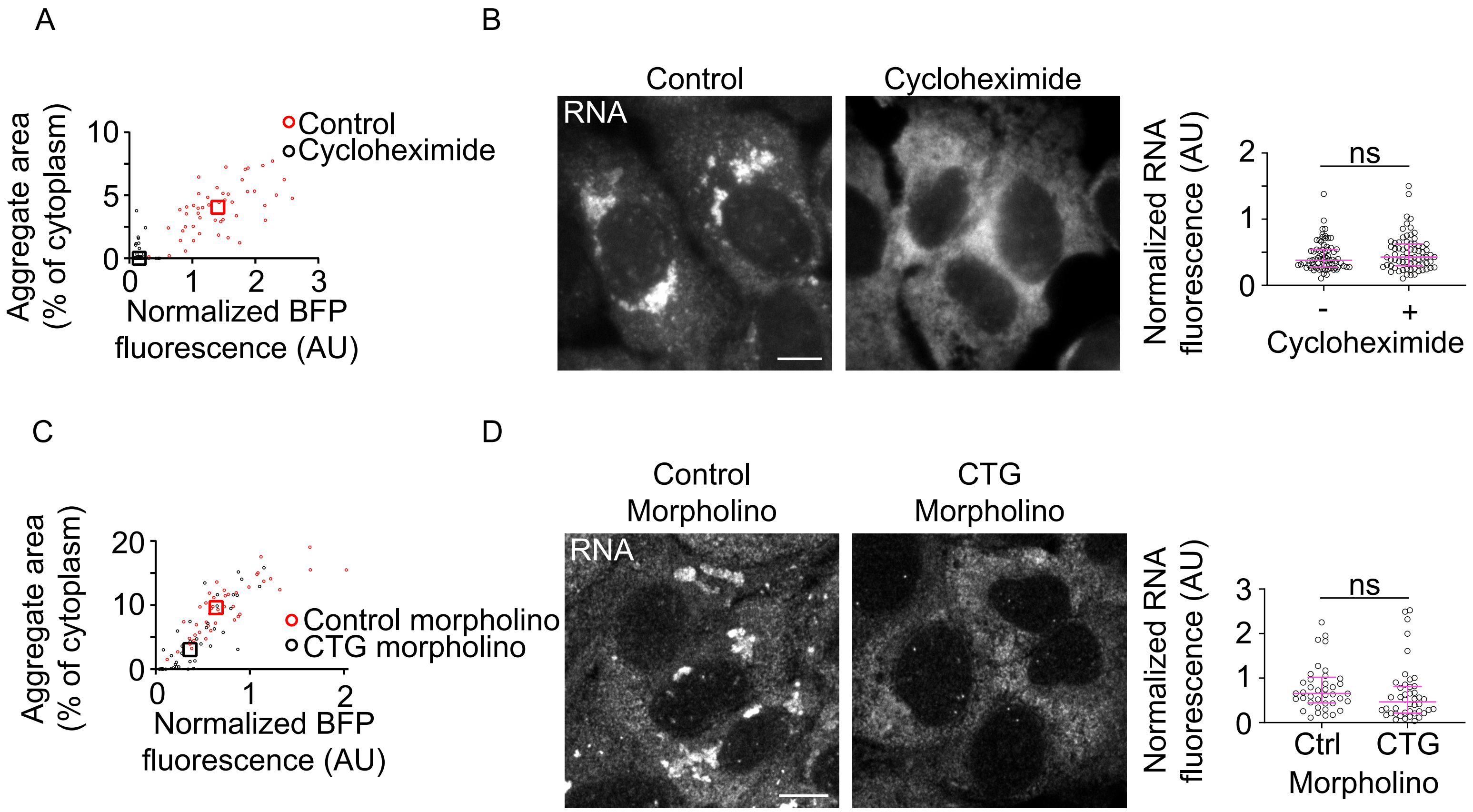

### Supp. Fig. 5

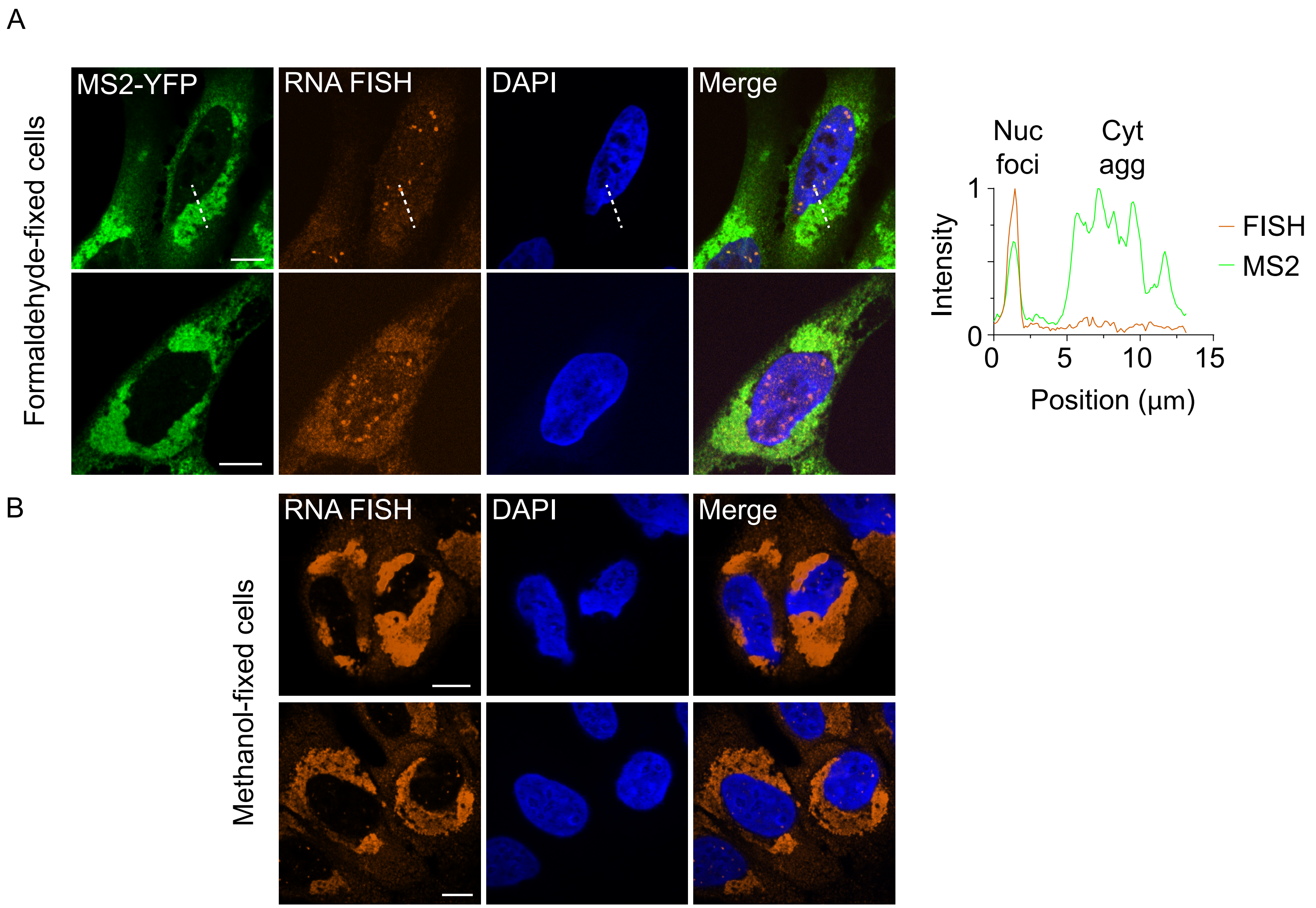
