## Supplementary material for "Flanking sequences regulate the toxicity, non-ATG translation, and aggregation of RNA with tandem CAG Repeats": Key Resources Table

| REAGENT or RESOURCE | SOURCE | IDENTIFIER |
|---|---|---|
| Antibodies | | |
| Rabbit anti-lamin B1 | Abcam | Cat#ab16048 |
| Mouse anti-G3BP | Abcam | Cat#ab56574 |
| Rabbit anti-EDC4 | Abcam | Cat#ab72408 |
| Mouse anti-p62/SQSTM1 | Abcam | Cat#ab56416 |
| Rabbit anti-TDP-43 | Proteintech | Cat#10782-2-AP |
| Rabbit anti-FUS | Invitrogen | Cat#PA552610 |
| Rabbit anti-TIS11b/ZFP36L1 | Abcam | Cat#ab230507 |
| Goat anti-GFP | Rockland | Cat#600-101-215 |
| Mouse anti-polyglutamine | Millipore-Sigma | Cat#MAB1574 |
| Alexa488-conjugated donkey anti-goat | Invitrogen | Cat#A11055 |
| Cy3-conjugated donkey anti-mouse | Jackson ImmunoResearch | Cat#715-165-151 |
| HRP-conjugated rabbit anti-mouse | Sigma-Aldrich | Cat#A9044-2ML |
| Cy3-conjugated donkey anti-rabbit | Jackson ImmunoResearch | Cat#711-165-152 |
| Bacterial and virus strains | | |
| Stbl3 *E. coli* | Invitrogen | Cat#C7373-03 |
| Biological samples | | |
| Chemicals, peptides, and recombinant proteins | | |
| EndoPorter peptide | GeneTools | SKU: OT-EP-PEG-1 |
| Methanol | Millipore-Sigma | Cat#MX0475-1 |
| Acetic acid | Sigma-Aldrich | Cat#695092 |
| UltaPure 20X SSC Buffer | Invitrogen | Cat#15557-044 |
| NP-40 substitute | Fisher Scientific | Cat#AAJ19628AP |
| Formamide | Sigma-Aldrich | Cat#47671 |

| | | |
|---|---|---|
| 16% Formaldehyde solution (w/v) Methanol-free | Thermo Scientific | Cat#28906 |
| Sodium citrate | Calbiochem | Cat#567446 |
| Dextran sulfate | Sigma-Aldrich | Cat#D8906 |
| Triton-X-100 | Sigma-Aldrich | Cat#T8787 |
| 5M Sodium chloride | Promega | Cat#V4221 |
| Sodium deoxycholate | Sigma-Aldrich | Cat#D6750 |
| SDS | Bio-Rad | Cat#1610302 |
| Tween-20 | FisherBiotech | Cat#BP337 |
| DTT | Thermo Scientific | Cat#R0861 |
| Skim milk powder | BD Biosciences | Cat#232100 |
| DAPI | Sigma-Aldrich | Cat#D9542 |
| Doxycycline | Sigma-Aldrich | Cat#D9891 |
| Cycloheximide | Sigma-Aldrich | Cat#C1998 |
| Polybrene | Millipore-Sigma | Cat#TR-1003-G |
| Lipofectamine LTX | Invitrogen | Cat#15338-100 |
| Viafect transfection reagent | Promega | Cat#E4981 |
| Trypan-blue 0.4% | Invitrogen | Cat#T10282 |
| Critical commercial assays | | |
| PureLink Genomic DNA Mini kit | Invitrogen | Cat#K1820-01 |
| PureLink RNA Mini kit | Invitrogen | Cat#12183018A |
| SuperScript III Reverse transcriptase | LifeTechnologies | Cat#11754050 |
| SYBR Green PCR Master Mix | AppliedBiosystems | Cat#4309155 |
| SuperSginal West Femto Maximum Sensitivity Substrate | Thermo Scientific | Cat#34095 |
| Protease and phosphatase inhibitors | Thermo Scientific | Cat#78442 |
| Deposited data | | |
| Experimental models: Cell lines | | |

| | | |
|---|---|---|
| Human: U-2 OS cells | ATCC | HTB-96 |
| Human: HEK293T cells | ATCC | CRL-3216 |
| Experimental models: Organisms/strains | | |
| Oligonucleotides | | |
| qPCR primer WPRE_F:<br><br>TGTCGGGGAAATCATCGTCC | IDT | N/A |
| qPCR primer WPRE_R:<br><br>AAGGAAGGTCCGCTGGATTG | IDT | N/A |
| qPCR primer TetPromoter_F:<br><br>AACGTATGTCGAGGTAGGCG | IDT | N/A |
| qPCR primer TetPromoter_R:<br><br>ATTGCTCCAGGCGATCTGAC | IDT | N/A |
| qPCR primer Actin_F:<br><br>GCTACGAGCTGCCTGACG | IDT | N/A |
| qPCR primer Actin_R:<br><br>GGCTGGAAGAGTGCCTCA | IDT | N/A |
| 7xCTG FISH probe:<br>Cy3/CTGCTGCTGCTGCTGCTGCTG | IDT | N/A |
| MS2 FISH probe:<br><br>Cy3/TTCTAGAGTCGACCTGCAG | IDT | N/A |
| 8xCTG morpholino:<br><br>CTGCTGCTGCTGCTGCTGCTGCTG | GeneTools | N/A |
| Standard intron control morpholino:<br><br>CCTCTTACCTCAGTTACAATTTATA | GeneTools | SKU: PCO-StandardControl-300 |
| Recombinant DNA | | |
| N0: pHR Tre3G 240xCAG 12xMS2 WPRE | This study | Supp. Table 2 |
| N1: pHR Tre3G Flank N1 240xCAG 12xMS2 WPRE | This study | Supp. Table 2 |

| | | |
|---|---|---|
| N2: pHR Tre3G Flank N2 240xCAG 12xMS2 WPRE | This study | Supp. Table 2 |
| N3: pHR Tre3G Flank N3 240xCAG 12xMS2 WPRE | This study | Supp. Table 2 |
| N4: pHR Tre3G Flank N4 240xCAG 12xMS2 WPRE | This study | Supp. Table 2 |
| N5: pHR Tre3G Flank N5 240xCAG 12xMS2 WPRE | This study | Supp. Table 2 |
| N6: pHR Tre3G Flank N6 240xCAG 12xMS2 WPRE | This study | Supp. Table 2 |
| N7: pHR Tre3G Flank N7 240xCAG 12xMS2 WPRE | This study | Supp. Table 2 |
| C1: pHR Tre3G Flank C1 240xCAG 12xMS2 WPRE | This study | Supp. Table 2 |
| C2: pHR Tre3G Flank C2 240xCAG 12xMS2 WPRE | This study | Supp. Table 2 |
| C3: pHR Tre3G Flank C3 240xCAG 12xMS2 WPRE | This study | Supp. Table 2 |
| C4: pHR Tre3G Flank C4 240xCAG 12xMS2 WPRE | This study | Supp. Table 2 |
| C5: pHR Tre3G Flank C5 240xCAG 12xMS2 WPRE | This study | Supp. Table 2 |
| ΔMS2: pHR Tre3G Flank C1 240xCAG WPRE | This study | Supp. Table 2 |
| ΔMS2, ΔWPRE: pHR Tre3G Flank C1 240xCAG | This study | Supp. Table 2 |
| mCherry: pHR Tre3G mCherry 12xMS2 | (Jain and Vale, 2017) | N/A |
| mCherry-revComp:<br>pHR Tre3G mCherryReverseComplement 12xMS2 | (Jain and Vale, 2017) | N/A |
| PiggyBac:<br>pPB Tre3G FlankC1 240xCAG 12xMS2 | This study | Supp. Table 2 |
| PiggyBac Transposase:<br>pJM134_piggybac_Transposase | Systems Biosciences | Cat# PB210PA-1 |
| CAG repeats with *ATXN8* flanking sequence:<br>pcDNA3.1 ATXN8 KKQ | (Zu et al., 2011) | N/A |
| CAG repeats with *HTT* flanking sequence:<br>pcDNA3.1 HTT | (Zu et al., 2011) | N/A |
| CAG repeats with *ATXN3* flanking sequence:<br>pcDNA3.1 ATXN3 | (Zu et al., 2011) | N/A |

| | | |
|---|---|---|
| MS2 coat protein – YFP fusion:<br><br>pHR-tdMS2CP-YFP-WPRE | (Jain and Vale, 2017) | N/A |
| RAN translation reporter:<br><br>pHR Tre3G Flank C1 240xCAG BFP 12xMS2 | This study | Supp. Table 2 |
| RAN translation reporter with stop:<br><br>pHR Tre3G Flank C1 240xCAG STOP BFP 12xMS2 | This study | Supp. Table 2 |
| Lentiviral envelope:<br><br>pCMV-VSV-G | (Stewart et al., 2003) | Addgene plasmid<br><br>#8454 |
| Lentiviral packaging:<br><br>psPAX2 | Gift from Didier Trono | Addgene plasmid<br><br>#12260 |
| Software and algorithms | | |
| ImageJ | (Schneider et al., 2012) | https://imagej.nih.gov/ij/ |
| Fusion | Oxford Instruments Andor | https://andor.oxinst.com/downloads/view/fusion-release-2.3 |
| Other | | |
| Dragonfly spinning disk confocal microscope | Oxford Instruments<br><br>Andor | Model#Dragonfly505 |
| iXon Ultra 888 EMCCD camera | Oxford Instruments<br><br>Andor | Model#DU-888U3-CS0-#BV |
| 100X Oil Immersion Objective, NA 1.45 | Nikon | CAT# MRD01905 |
| 96-well glass bottom plates | Brooks | CAT# MGB096-1-2-LG-L |
| iBlot 2 | Invitrogen | CAT# IB21001 |
| Countess II FL automated cell counter | Invitrogen | CAT# AMQAF1000 |
| Countess II disposable cell counting chamber slides | Invitrogen | CAT# C10283 |

| | | |
|---|---|---|
| DMEM | Gibco | CAT# 11965-126 |
| Fetal bovine serum | Gibco | CAT# 26140-079 |
| Penicillin-streptomycin-glutamine 100X | Gibco | CAT# 10378016 |
| Opti-MEM | Gibco | CAT# 31985-070 |
| IMDM | Gibco | CAT# 12440-053 |
| DPBS | Gibco | CAT# 14190-144 |
| Trypsin-EDTA 0.25% | Gibco | CAT# 25200-072 |
| PBS, pH=7.2 | Gibco | CAT# 20012-027 |
| Nuclease-free water | Invitrogen | CAT# AM9932 |
| Bovine serum albumin (BSA) | Sigma-Aldrich | CAT# A7906 |
| 4xBolt LDS sample buffer | Invitrogen | CAT# B0007 |
| 4-12% Bis-tris polyacrylamide gel | Invitrogen | CAT# NW04122 |
