## Supplementary material for "Flanking sequences regulate the toxicity, non-ATG translation, and aggregation of RNA with tandem CAG Repeats": Supp. Table 1

| Flank | GC-content | Source | |
|---|---|---|---|
| N1 | 32 | Synthetic | GTTGAATACTCATACTCTTCCTTTTTCAATATTATTGAAGCATTTATCA GGGTTATTGTCTCATGAGCGGATACATATTTGAATGTATTTAGAAAA ATAAACAAATAGGGGTTCCGCGCACATTTCCCCGAAAAGTGCCACCT AAATTGTAAGCGTTAATATTTTGTTAAAATTCGCGTTAAATTTTTGTT AAATCAGCTCATTTTTTAACCAATAGGCCGAAATCGGCAAAATCCCTT ATAAATCAAAAG |
| N2 | 74 | Upstream flank of *DMPK* | CGCCGCCCAGGAGCCGCCCGCGCTCCCTGAACCCTAGAACTGTCTTC GACTCCGGGGCCCCGTTGGAAGACTGAGTGCCCGGGGCACGGCAC AGAAGCCGCGCCCACCGCCTGCCAGTTCACAACCGCTCCGAGCGTG GGTCTCCGCCCAGCTCCAGTCCTGTGATCCGGGCCCGCCCCCTAGCG GCCGGGGAGGGAGGGGCCGGGTCCGCGGCCGGCGAACGGGGCTC GAAGGGTCCTTGTAGCCGGGA |
| N3 | 55 | Synthetic | GCTCCGGATCCCTTCGATCCACGACCCATCCTGGACGACTACCCGGG TATTTCTTCGGTATTGAGTATACGACCGCGTAGTGGACCGATACCTA GGCGGCAGAACTATCTTATACAAATAACACGAGGCCTAGTGGCTGA TCTAAGCAGCAAGTCATGGGCCCAAGCACTACGGACAAAATGGCCT GGGGTTGGGAGCAGTAGGTACGGTACCCCGTATAGTGCGCAGATCT AGTTCGCGGGATCGGTCG |
| N4 | 55 | Synthetic | GTCCACTATTAAAGAACGTGGACTCCAACGTCAAAGGGCGAAAAAC CGTCTATCAGGGCGATGGCCCACTACGTGAACCATCACCCTAATCAA GTTTTTTGGGGTCGAGGTGCCGTAAAGCACTAAATCGGAACCCTAAA GGGAGCCCCCGATTTAGAGCTTGACGGGGAAAGCCGGCGAACGTG GCGAGAAAGGAAGGGAAGAAAGCGAAAGGAGCGGGCGCTAGGGC GCTGGCAAGTGTAGCGGTCAC |
| N5 | 79 | Downstream flank of *HTT* | CCGCCACCGCCGCCGCCGCCGCCGCCGCCTCCTCAGCTTCCTCAGCC GCCGCCGCAGGCACAGCCGCTGCTGCCTCAGCCGCAGCCGCCCCCG CCGCCACCCGGCCCGGCTGTGGCTGAGGAGCCGCTGCACCGACCGT GAGTTTGGGCCCGCTGCAGCTCCCTGTCCCGGCGGGTCCCAGGCTAC GGCGGGGATGGCGGTAACCCTGCAGCCTGCGGGCCGGCGACACGA ACCCCCG |
| N6 | 30 | Synthetic | CATAGTGTCTAAGAAGTAATCCTAAATTAATCTATGAACAAGAGCGT ATTAAAATTATAAATGGCTCATAATTTTCTCTACGAGTGCAAACATAT ATAAAACATAAGGGTCTATGGATTCTGAGTAAGTAACATACTTCTTA TTATAATCTTCCGATGGTTCAAGCAAGAAGAAGTTTTCTCAACTGAA AGGAAACTTTATTAATTATCTTAATGAGGAGTGAGGTACAGGGGAG ATAATAATAATGAAA |
| N7 | 38 | Upstream flank of *CNBP* | TCACCAGGCAAGTGCTGCAGTATAACTAGGTACTACGTCAGGTGCTA AGGTTAAGAGAGTATTTTCCTTCACTGACTCCTCACTCCGAGAATCCA TTTTACAGCTTCATTGGTTTGGGTTATTCCAATTTTTTGATGTGAGTA AATAAATGACTTCTATTTGCCCAAAATAAAGCTTATATAGGCCTTATA ACCATGCAAATGTGTCCATTAAGTTGGACTTGGAATGAGTGAATGA GTATTACTGCCAG |
| C1 | 74 | Synthetic | GGTTCCTGGCCACCGTCGGCGTCTCGCCCGACCACCAGGGCAAGGG TCTGGGCAGCGCCGTCGTGCTCCCCGGAGTGGAGGCGGCCGAGCGC |

| | | | |
|---|---|---|---|
| | | | GCCGGGGTGCCCGCCTTCCTGGAGACCTCCGCGCCCCGCAACCTCCC CTTCTACGAGCGGCTCGGCTTCACCGTCACCGCCGACGTCGAGGTGC CCGAAGGACCGCGCACCTGGTGCAcGACCCGCAAGCCCGGTGCCGG ATCGGGAGAGGGCAGAGG |
| C2 | 65 | Synthetic | TTACAGACTATCCGGGTCGCTAGCCGGTACCAGGCACTTCGTGGGAT CGCCTTGCGAGCGGCACGCGGAGGTCCACCCCTCATGTCGATGTACC GACGCAGCTCCCGATGTTCTGCCCAGAATATTGCAGGGGGTCCGAA CTCCACGCGGACTAGGCAGAGGGCCTGGAATGCGGCCGGCGCATTC CCGGGGGTGGTCCTGAGGCTGCGGACGGCGGGGTATTGGCGAATC GGCGAGGCGGATTGTGAGA |
| C3 | 50 | Synthetic | TTGGGTCTGGGCTCATAGGGTGTTCACATCTGGGGAGTATGGTCGG ATCTAATACGTCGTTTGGGAAATCAAGAGTCGGCAGCGCAGGTCCC ACCAACTACTCATGTGATTCGCACTTACTATTCCTACTCGATGGAGAG GATCTAGGGTTAGCAGCCAAGCCGGAGCCTCACATATCACCGTCACT AGTCGGTAATATCCTGCCTATGCAAAGCACGATGAAGCATAGATTCA TTCTCGTCATTCGGGA |
| C4 | 75 | Synthetic | CACGGGGTACCGGCCGTCGGACGCCGCGTTCGCCCAACTGGCACAG TACCCCGGCACTGCCCGCGCAGGCCTCGCGCGCGCCAGGACCTGCC CTGCGCGAGCGGGTACCGCGTACGATGCGGCCTCGGCGCCCAAGCG CGCACGGGCCCGTGGCCCAATTCCAGCGTGGGGTGAGCGAGAGGC GGGACCCGCCGCTCTGCCTAGGCCACCTCGACGCGCAAACGTTGGT GGTCGCGGCTACCTCCGTCCC |
| C5 | 75 | Synthetic | TGGCACCGCGAGGTGGCACCCCGGCGCCGAGACCCTCGTGTCTCAG TCCTGGCCCGCAACACGCCCTGCAAGTGCGCGCGATCAGGAACCGA CCCCCGCGAGCCCCGCTGAGCCCGTACTCCGGCCCACCCCGCGTGGC CCCGTCCCCTATCGCGGTGGGTGGGCGCCACAGGCGAGTCCAGGGG GGTGGTGGGCTGTTCGGCTGGCCCGGGCCTTGGTGGGGCCGAGGG AGCCGGCTGATGGTCGAACC |
