## Supplementary material for "Flanking sequences regulate the toxicity, non-ATG translation, and aggregation of RNA with tandem CAG Repeats": Supp. Table 2

| | |
|---|---|
| N0: pHR Tre3G 240xCAG 12xMS2 WPRE | ctatagtgtcacctaaatcgtatgtgtatgatacataaggttatgtattaattgtagccgcgttctaacgacaatatgtacaagccta<br>attgtgtagcatctggcttactgaagcagaccctatcatctctctcgtaaactgccgtcagagtcggtttggttggacgaaccttctg<br>agtttctggtaacgccgtcccgcacccggaaatggtcagcgaaccaatcagcagggtcatcgctagccagatcctctacgccgga<br>cgcatcgtggccggcatcaccggcgccacaggtgcggttgctggcgcctatatcgccgacatcaccgatggggaagatcgggctc<br>gccacttcgggctcatgagcgcttgtttcggcgtgggtatggtggcaggccccgtggccgggggactgttgggcgccatctccttgc<br>atgcaccattccttgcggcggcggtgctcaacggcctcaacctactactgggctgcttcctaatgcaggagtcgcataagggagag<br>cgtcgaatggtgcactctcagtacaatctgctctgatgccgcatagttaagccagccccgacacccgccaacacccgctgacgcgc<br>cctgacgggcttgtctgctcccggcatccgcttacagacaagctgtgaccgtctccgggagctgcatgtgtcagaggttttcaccgt<br>catcaccgaaacgcgcgagacgaaagggcctcgtgatacgcctatttttataggttaatgtcatgataataatggtttcttagacgt<br>caggtggcacttttcggggaaatgtgcgcggaacccctatttgtttatttttctaaatacattcaaatatgtatccgctcatgagaca<br>ataaccctgataaatgcttcaataatattgaaaaaggaagagtatgagtattcaacatttccgtgtcgcccttattcccttttttgcg<br>gcattttgccttcctgtttttgctcacccagaaacgctggtgaaagtaaaagatgctgaagatcagttgggtgcacgagtgggttac<br>atcgaactggatctcaacagcggtaagatccttgagagttttcgccccgaagaacgttttccaatgatgagcacttttaaagttctg<br>ctatgtggcgcggtattatcccgtattgacgccgggcaagagcaactcggtcgccgcatacactattctcagaatgacttggttga<br>gtactcaccagtcacagaaaagcatcttacggatggcatgacagtaagagaattatgcagtgctgccataaccatgagtgataac<br>actgcggccaacttacttctgacaacgatcggaggaccgaaggagctaaccgcttttttgcacaacatgggggatcatgtaactcg<br>ccttgatcgttgggaaccggagctgaatgaagccataccaaacgacgagcgtgacaccacgatgcctgtagcaatggcaacaac<br>gttgcgcaaactattaactggcgaactacttactctagcttcccggcaacaattaatagactggatggaggcggataaagttgcag<br>gaccacttctgcgctcggcccttccggctggctggtttattgctgataaatctggagccggtgagcgtgggtctcgcggtatcattgc<br>agcactggggccagatggtaagccctcccgtatcgtagttatctacacgacggggagtcaggcaactatggatgaacgaaatag<br>acagatcgctgagataggtgcctcactgattaagcattggtaactgtcagaccaagtttactcatatatactttagattgatttaaa<br>acttcatttttaatttaaaaggatctaggtgaagatcctttttgataatctcatgaccaaaatcccttaacgtgagttttcgttccact<br>gagcgtcagaccccgtagaaaagatcaaaggatcttcttgagatcctttttttctgcgcgtaatctgctgcttgcaaacaaaaaaa<br>ccaccgctaccagcggtggtttgtttgccggatcaagagctaccaactctttttccgaaggtaactggcttcagcagagcgcagat<br>accaaatactgtctttctagtgtagccgtagttaggccaccacttcaagaactctgtagcaccgcctacatacctcgctctgctaatc<br>ctgttaccagtggctgctgccagtggcgataagtcgtgtcttaccgggttggactcaagacgatagttaccggataaggcgcagcg<br>gtcgggctgaacggggggttcgtgcacacagcccagcttggagcgaacgacctacaccgaactgagatacctacagcgtgagct<br>atgagaaagcgccacgcttcccgaagggagaaaggcggacaggtatccggtaagcggcagggtcggaacaggagagcgcacg<br>agggagcttccagggggaaacgcctggtatctttatagtcctgtcgggtttcgccacctctgacttgagcgtcgatttttgtgatgct<br>cgtcaggggggcggagcctatggaaaaacgccagcaacgcggcctttttacggttcctggccttttgctggccttttgctcacatgt<br>tctttcctgcgttatcccctgattctgtggataaccgtattaccgcctttgagtgagctgataccgctcgccgcagccgaacgaccga<br>gcgcagcgagtcagtgagcgaggaagcggaagagcgcccaatacgcaaaccgcctctccccgcgcgttggccgattcattaatg<br>cagctgtggaatgtgtgtcagttagggtgtggaaagtccccaggctccccagcaggcagaagtatgcaaagcatgcatctcaatt<br>agtcagcaaccaggtgtggaaagtccccaggctccccagcaggcagaagtatgcaaagcatgcatctcaattagtcagcaacca<br>tagtcccgcccctaactccgcccatcccgcccctaactccgcccagttccgcccattctccgccccatggctgactaattttttttattt<br>atgcagaggccgaggccgcctcggcctctgagctattccagaagtagtgaggaggcttttttggaggcctaggcttttgcaaaaag<br>cttggacacaagacaggcttgcgagatatgtttgagaataccactttatcccgcgtcagggagaggcagtgcgtaaaaagacgc<br>ggactcatgtgaaatactggtttttagtgcgccagatctctataatctcgcgcaacctattttcccctcgaacactttttaagccgtag<br>ataaacaggctgggacacttcacatgagcgaaaaatacatcgtcacctgggacatgttgcagatccatgcacgtaaactcgcaa<br>gccgactgatgccttctgaacaatggaaaggcattattgccgtaagccgtggcggtctgtaccgggtgcgttactggcgcgtgaac<br>tgggtattcgtcatgtcgataccgtttgtatttccagctacgatcacgacaaccagcgcgagcttaaagtgctgaaacgcgcagaa<br>ggcgatggcgaaggcttcatcgttattgatgacctggtggataccggtggtactgcggttgcgattcgtgaaatgtatccaaaagc<br>gcactttgtcaccatcttcgcaaaaccggctggtcgtccgctggttgatgactatgttgttgatatcccgcaagatacctggattgaa<br>cagccgtgggatatgggcgtcgtattcgtcccgccaatctccggtcgctaatcttttcaacgcctggcactgccgggcgttgttctttt<br>taacttcaggcgggttacaatagtttccagtaagtattctggaggctgcatccatgacacaggcaaacctgagcgaaaccctgttc<br>aaaccccgctttaaacatcctgaaacctcgacgctagtccgccgctttaatcacggcgcacaaccgcctgtgcagtcggcccttga<br>tggtaaaaccatccctcactggtatcgcatgattaaccgtctgatgtggatctggcgcggcattgacccacgcgaaatcctcgacg<br>tccaggcacgtattgtgatgagcgatgccgaacgtaccgacgatgatttatacgatacggtgattggctaccgtggcggcaactgg |

| | |
|---|---|
| | atttatgagtgggccccggatctttgtgaaggaaccttacttctgtggtgtgacataattggacaaactacctacagagatttaaag<br>ctctaaggtaaatataaaatttttaagtgtataatgtgttaaactactgattctaattgtttgtgtattttagattccaacctatggaa<br>ctgatgaatgggagcagtggtggaatgcctttaatgaggaaaacctgttttgctcagaagaaatgccatctagtgatgatgaggct<br>actgctgactctcaacattctactcctccaaaaaagaagagaaaggtagaagaccccaaggactttccttcagaattgctaagttt<br>tttgagtcatgctgtgtttagtaatagaactcttgcttgctttgctatttacaccacaaaggaaaaagctgcactgctatacaagaa<br>aattatggaaaaatattctgtaacctttataagtaggcataacagttataatcataacatactgttttttcttactccacacaggcat<br>agagtgtctgctattaataactatgctcaaaaattgtgtacctttagctttttaatttgtaaaggggttaataaggaatatttgatgta<br>tagtgccttgactagagatcataatcagccataccacatttgtagaggttttacttgctttaaaaaacctcccacacctccccctga<br>acctgaaacataaaatgaatgcaattgttgttgttaacttgtttattgcagcttataatggttacaaataaagcaatagcatcacaa<br>atttcacaaataaagcatttttttcactgcattctagttgtggtttgtccaaactcatcaatgtatcttatcatgtctggatcaactgga<br>taactcaagctaaccaaaatcatcccaaacttcccaccccataccctattaccactgccaattacctagtggtttcatttactctaaa<br>cctgtgattcctctgaattattttcattttaaagaaattgtatttgttaaatatgtactacaaacttagtagttggaagggctaattca<br>ctcccaaagaagacaagatatccttgatctgtggatctaccacacacaaggctacttccctgattagcagaactacacaccaggg<br>ccaggggtcagatatccactgacctttggatggtgctacaagctagtaccagttgagccagataaggtagaagaggccaataaa<br>ggagagaacaccagcttgttacaccctgtgagcctgcatgggatggatgacccggagagagaagtgttagagtggaggtttgac<br>agccgcctagcatttcatcacgtggcccgagagctgcatccggagtacttcaagaactgctgatatcgagcttgctacaagggact<br>ttccgctggggactttccagggaggcgtggcctgggcgggactggggagtggcgagccctcagatcctgcatataagcagctgct<br>ttttgcctgtactgggtctctctggttagaccagatctgagcctgggagctctctggctaactagggaacccactgcttaagcctcaa<br>taaagcttgccttgagtgcttcaagtagtgtgtgcccgtctgttgtgtgactctggtaactagagatccctcagacccttttagtcagt<br>gtggaaaatctctagcagtggcgcccgaacagggacttgaaagcgaaagggaaaccagaggagctctctcgacgcaggactcg<br>gcttgctgaagcgcgcacggcaagaggcgaggggcggcgactggtgagtacgccaaaaattttgactagcggaggctagaagg<br>agagagatgggtgcgagagcgtcagtattaagcgggggagaattagatcgcgatgggaaaaaattcggttaaggccagggga<br>aagaaaaaatataaattaaaacatatagtatgggcaagcagggagctagaacgattcgcagttaatcctggcctgttagaaaca<br>tcagaaggctgtagacaaatactgggacagctacaaccatcccttcagacaggatcagaagaacttagatcattatataatacag<br>tagcaaccctctattgtgtgcatcaaaggatagagataaaagacaccaaggaagctttagacaagatagaggaagagcaaaac<br>aaaagtaagaccaccgcacagcaagcggccggccgctgatcttcagacctggaggaggagatatgagggacaattggagaagt<br>gaattatataaatataaagtagtaaaaattgaaccattaggagtagcacccaccaaggcaaagagaagagtggtgcagagaga<br>aaaaagagcagtgggaataggagctttgttccttgggttcttgggagcagcaggaagcactatgggcgcagcgtcaatgacgctg<br>acggtacaggccagacaattattgtctggtatagtgcagcagcagaacaatttgctgagggctattgaggcgcaacagcatctgtt<br>gcaactcacagtctggggcatcaagcagctccaggcaagaatcctggctgtgaaagatacctaaaggatcaacagctcctgggg<br>atttggggttgctctggaaaactcatttgcaccactgctgtgccttggaatgctagttggagtaataaatctctggaacagatttgg<br>aatcacacgacctggatggagtgggacagagaaattaacaattacacaagcttaatacactccttaattgaagaatcgcaaaac<br>cagcaagaaaagaatgaacaagaattattggaattagataaatgggcaagtttgtggaattggtttaacataacaaattggctgt<br>ggtatataaaattattcataatgatagtaggaggcttggtaggtttaagaatagtttttgctgtactttctatagtgaatagagttag<br>gcagggatattcaccattatcgtttcagacccacctcccaaccccgaggggacccgacaggcccgaaggaatagaagaagaag<br>gtggagagagagacagagacagatccattcgattagtgaacggatctcgacggtatcgccaaatggcagtattcatccacaattt<br>taaaagaaaaggggggattggggggtacagtgcaggggaaagaatagtagacataatagcaacagacatacaaactaaagaa<br>ttacaaaaacaaattacaaaaattcaaaattttcgggtttattacagggacagcagagatccagtttatcgatgaggccctttcgt<br>cttcactcgagtttactccctatcagtgatagagaacgtatgaagagtttactccctatcagtgatagagaacgtatgcagacttta<br>ctccctatcagtgatagagaacgtataaggagtttactccctatcagtgatagagaacgtatgaccagtttactccctatcagtgat<br>agagaacgtatctacagtttactccctatcagtgatagagaacgtatatccagtttactccctatcagtgatagagaacgtatgtcg<br>aggtaggcgtgtacggtgggcgcctataaaagcagagctcgtttagtgaaccgtcagatcgcctggagcaattccacaacactttt<br>gtcttatacttACGCGTGTCGACcatatgACTAGTatgGGTAATTATAACCCGGGCCCTATATATTAGGCTA<br>ACTAGCTAACTAGGAATTCggtctcaCAGCAGCAGCAGCAGCAGCAGCAGCAGCAGCAGCAGC<br>AGCAGCAGCAGCAGCAGCAGCAGCAGCAGCAGCAGCAGCAGCAGCAGCAGCAGCAGC<br>AGCAGCAGCAGCAGCAGCAGCAGCAGCAGCAGCAGCAGCAGCAGCAGCAGCAGCAGC<br>AGCAGCAGCAGCAGCAGCAGCAGCAGCAGCAGCAGCAGCAGCAGCAGCAGCAGCAGC<br>AGCAGCAGCAGCAGCAGCAGCAGCAGCAGCAGCAGCAGCAGCAGCAGCAGCAGCAGC |

| | |
|---|---|
| | AGCAGCAGCAGCAGCAGCAGCAGCAGCAGCAGCAGCAGCAGCAGCAGCAGCAGCAGCAGC<br>AGCAGCAGCAGCAGCAGCAGCAGCAGCAGCAGCAGCAGCAGCAGCAGCAGCAGCAGCAGC<br>AGCAGCAGCAGCAGCAGCAGCAGCAGCAGCAGCAGCAGCAGCAGCAGCAGCAGCAGCAGC<br>AGCAGCAGCAGCAGCAGCAGCAGCAGCAGCAGCAGCAGCAGCAGCAGCAGCAGCAGCAGC<br>AGCAGCAGCAGCAGCAGCAGCAGCAGCAGCAGCAGCAGCAGCAGCAGCAGCAGCAGCAGC<br>AGCAGCAGCAGCAGCAGCAGCAGCAGCAGCAGCAGCAGCAGCAGCAGCAGCAGCAGCAGC<br>AGCAGCAGCAGCAGCAGCAGCAGCAGCAGCAGCAGCAGCAGCAGCAGCAGCAGCAGCAGC<br>AGCAGCAgagacgGAGCTCgtcgacGCggccgcaagcttggtaccgagctcGGATCCTAAGGTACCTAATTGC<br>CTAGAAAACATGAGGATCACCCATGTCTGCAGGTCGACTCTAGAAAACATGAGGATCACCCATGTC<br>TGCAGTATTCCCGGGTTCATTAGATCCTAAGGTACCTAATTGCCTAGAAAACATGAGGATCACCCA<br>TGTCTGCAGGTCGACTCTAGAAAACATGAGGATCACCCATGTCTGCAGTATTCCCGGGTTCATTAG<br>ATCCTAAGGTACCTAATTGCCTAGAAAACATGAGGATCACCCATGTCTGCAGGTCGACTCTAGAAA<br>ACATGAGGATCACCCATGTCTGCAGTATTCCCGGGTTCATTAGATCCTAAGGTACCTAATTGCCTAG<br>AAAACATGAGGATCACCCATGTCTGCAGGTCGACTCTAGAAAACATGAGGATCACCCATGTCTGCA<br>GTATTCCCGGGTTCATTAGATCCTAAGGTACCTAATTGCCTAGAAAACATGAGGATCACCCATGTCT<br>GCAGGTCGACTCTAGAAAACATGAGGATCACCCATGTCTGCAGTATTCCCGGGTTCATTAGATCCT<br>AAGGTACCTAATTGCCTAGAAAACATGAGGATCACCCATGTCTGCAGGTCGACTCTAGAAAACATG<br>AGGATCACCCATGTCTGCAGTATTCCCGGGTTCATTAGATCTGCGCGCGATCGATATCAGCGCTTT<br>AAATTTGCGCATGCTAGTgTAAggcatgcaagcttgatatcaagcttatcgataatcaacctctggattacaaaatttgtg<br>aaagattgactggtattcttaactatgttgctccttttacgctatgtggatacgctgctttaatgcctttgtatcatgctattgcttccc<br>gtatggctttcattttctcctccttgtataaatcctggttgctgtctctttatgaggagttgtggcccgttgtcaggcaacgtggcgtgg<br>tgtgcactgtgtttgctgacgcaacccccactggttggggcattgccaccacctgtcagctcctttccgggactttcgctttccccctc<br>cctattgccacggcggaactcatcgccgcctgccttgcccgctgctggacaggggctcggctgttgggcactgacaattccgtggt<br>gttgtcggggaaatcatcgtcctttccttggctgctcgcctgtgttgccacctggattctgcgcgggacgtccttctgctacgtccctt<br>cggccctcaatccagcggaccttccttcccgcggcctgctgccggctctgcggcctcttccgcgtcttcgccttcgccctcagacgag<br>tcggatctccctttgggccgcctccccgcatcgataccgtcgacctcgagggaattaattcgagctcggtacctttaagaccaatga<br>cttacaaggcagctgtagatcttagccactttttaaaagaaaaggggggactggaagggctaattcactcccaacgaagacaag<br>atctgctttttgcttgtactgggtctctctggttagaccagatctgagcctgggagctctctggctaactagggaacccactgcttaa<br>gcctcaataaagcttgccttgagtgcttcaagtagtgtgtgcccgtctgttgtgtgactctggtaactagagatccctcagaccctttt<br>tagtcagtgtggaaaatctctagcagcatctagaattaattccgtgtatt |
| N1: pHR Tre3G Flank N1 240xCAG 12xMS2 WPRE | ctatagtgtcacctaaatcgtatgtgtatgatacataaggttatgtattaattgtagccgcgttctaacgacaatatgtacaagccta<br>attgtgtagcatctggcttactgaagcagaccctatcatctctctcgtaaactgccgtcagagtcggtttggttggacgaaccttctg<br>agtttctggtaacgccgtcccgcacccggaaatggtcagcgaaccaatcagcagggtcatcgctagccagatcctctacgccgga<br>cgcatcgtggccggcatcaccggcgccacaggtgcggttgctggcgcctatatcgccgacatcaccgatggggaagatcgggctc<br>gccacttcgggctcatgagcgcttgtttcggcgtgggtatggtggcaggccccgtggccgggggactgttgggcgccatctccttgc<br>atgcaccattccttgcggcggcggtgctcaacggcctcaacctactactgggctgcttcctaatgcaggagtcgcataagggagag<br>cgtcgaatggtgcactctcagtacaatctgctctgatgccgcatagttaagccagccccgacacccgccaacacccgctgacgcgc<br>cctgacgggcttgtctgctcccggcatccgcttacagacaagctgtgaccgtctccgggagctgcatgtgtcagaggttttcaccgt<br>catcaccgaaacgcgcgagacgaaagggcctcgtgatacgcctatttttataggttaatgtcatgataataatggtttcttagacgt<br>caggtggcacttttcggggaaatgtgcgcggaacccctatttgtttatttttctaaatacattcaaatatgtatccgctcatgagaca<br>ataaccctgataaatgcttcaataatattgaaaaaggaagagtatgagtattcaacatttccgtgtcgcccttattcccttttttgcg<br>gcattttgccttcctgtttttgctcacccagaaacgctggtgaaagtaaaagatgctgaagatcagttgggtgcacgagtgggttac<br>atcgaactggatctcaacagcggtaagatccttgagagttttcgccccgaagaacgttttccaatgatgagcacttttaaagttctg<br>ctatgtggcgcggtattatcccgtattgacgccgggcaagagcaactcggtcgccgcatacactattctcagaatgacttggttga<br>gtactcaccagtcacagaaaagcatcttacggatggcatgacagtaagagaattatgcagtgctgccataaccatgagtgataac<br>actgcggccaacttacttctgacaacgatcggaggaccgaaggagctaaccgcttttttgcacaacatgggggatcatgtaactcg<br>ccttgatcgttgggaaccggagctgaatgaagccataccaaacgacgagcgtgacaccacgatgcctgtagcaatggcaacaac |

| | |
|---|---|
| | gttgcgcaaactattaactggcgaactacttactctagcttcccggcaacaattaatagactggatggaggcggataaagttgcag<br>gaccacttctgcgctcggcccttccggctggctggtttattgctgataaatctggagccggtgagcgtgggtctcgcggtatcattgc<br>agcactggggccagatggtaagccctcccgtatcgtagttatctacacgacggggagtcaggcaactatggatgaacgaaatag<br>acagatcgctgagataggtgcctcactgattaagcattggtaactgtcagaccaagtttactcatatatactttagattgatttaaa<br>acttcatttttaatttaaaaggatctaggtgaagatcctttttgataatctcatgaccaaaatcccttaacgtgagttttcgttccact<br>gagcgtcagaccccgtagaaaagatcaaaggatcttcttgagatcctttttttctgcgcgtaatctgctgcttgcaaacaaaaaaa<br>ccaccgctaccagcggtggtttgtttgccggatcaagagctaccaactctttttccgaaggtaactggcttcagcagagcgcagat<br>accaaatactgtctttctagtgtagccgtagttaggccaccacttcaagaactctgtagcaccgcctacatacctcgctctgctaatc<br>ctgttaccagtggctgctgccagtggcgataagtcgtgtcttaccgggttggactcaagacgatagttaccggataaggcgcagcg<br>gtcgggctgaacggggggttcgtgcacacagcccagcttggagcgaacgacctacaccgaactgagatacctacagcgtgagct<br>atgagaaagcgccacgcttcccgaagggagaaaggcggacaggtatccggtaagcggcagggtcggaacaggagagcgcacg<br>agggagcttccagggggaaacgcctggtatctttatagtcctgtcgggtttcgccacctctgacttgagcgtcgatttttgtgatgct<br>cgtcaggggggcggagcctatggaaaaacgccagcaacgcggcctttttacggttcctggccttttgctggccttttgctcacatgt<br>tctttcctgcgttatcccctgattctgtggataaccgtattaccgcctttgagtgagctgataccgctcgccgcagccgaacgaccga<br>gcgcagcgagtcagtgagcgaggaagcggaagagcgcccaatacgcaaaccgcctctccccgcgcgttggccgattcattaatg<br>cagctgtggaatgtgtgtcagttagggtgtggaaagtccccaggctccccagcaggcagaagtatgcaaagcatgcatctcaatt<br>agtcagcaaccaggtgtggaaagtccccaggctccccagcaggcagaagtatgcaaagcatgcatctcaattagtcagcaacca<br>tagtcccgcccctaactccgcccatcccgcccctaactccgcccagttccgcccattctccgccccatggctgactaattttttttattt<br>atgcagaggccgaggccgcctcggcctctgagctattccagaagtagtgaggaggcttttttggaggcctaggcttttgcaaaaag<br>cttggacacaagacaggcttgcgagatatgtttgagaataccactttatcccgcgtcagggagaggcagtgcgtaaaaagacgc<br>ggactcatgtgaaatactggtttttagtgcgccagatctctataatctcgcgcaacctattttcccctcgaacactttttaagccgtag<br>ataaacaggctgggacacttcacatgagcgaaaaatacatcgtcacctgggacatgttgcagatccatgcacgtaaactcgcaa<br>gccgactgatgccttctgaacaatggaaaggcattattgccgtaagccgtggcggtctgtaccgggtgcgttactggcgcgtgaac<br>tgggtattcgtcatgtcgataccgtttgtatttccagctacgatcacgacaaccagcgcgagcttaaagtgctgaaacgcgcagaa<br>ggcgatggcgaaggcttcatcgttattgatgacctggtggataccggtggtactgcggttgcgattcgtgaaatgtatccaaaagc<br>gcactttgtcaccatcttcgcaaaaccggctggtcgtccgctggttgatgactatgttgttgatatcccgcaagatacctggattgaa<br>cagccgtgggatatgggcgtcgtattcgtcccgccaatctccggtcgctaatcttttcaacgcctggcactgccgggcgttgttctttt<br>taacttcaggcgggttacaatagtttccagtaagtattctggaggctgcatccatgacacaggcaaacctgagcgaaaccctgttc<br>aaaccccgctttaaacatcctgaaacctcgacgctagtccgccgctttaatcacggcgcacaaccgcctgtgcagtcggcccttga<br>tggtaaaaccatccctcactggtatcgcatgattaaccgtctgatgtggatctggcgcggcattgacccacgcgaaatcctcgacg<br>tccaggcacgtattgtgatgagcgatgccgaacgtaccgacgatgatttatacgatacggtgattggctaccgtggcggcaactgg<br>atttatgagtgggccccggatctttgtgaaggaaccttacttctgtggtgtgacataattggacaaactacctacagagatttaaag<br>ctctaaggtaaatataaaatttttaagtgtataatgtgttaaactactgattctaattgtttgtgtattttagattccaacctatggaa<br>ctgatgaatgggagcagtggtggaatgcctttaatgaggaaaacctgttttgctcagaagaaatgccatctagtgatgatgaggct<br>actgctgactctcaacattctactcctccaaaaaagaagagaaaggtagaagaccccaaggactttccttcagaattgctaagttt<br>tttgagtcatgctgtgtttagtaatagaactcttgcttgctttgctatttacaccacaaaggaaaaagctgcactgctatacaagaa<br>aattatggaaaaatattctgtaacctttataagtaggcataacagttataatcataacatactgttttttcttactccacacaggcat<br>agagtgtctgctattaataactatgctcaaaaattgtgtacctttagctttttaatttgtaaaggggttaataaggaatatttgatgta<br>tagtgccttgactagagatcataatcagccataccacatttgtagaggttttacttgctttaaaaaacctcccacacctccccctga<br>acctgaaacataaaatgaatgcaattgttgttgttaacttgtttattgcagcttataatggttacaaataaagcaatagcatcacaa<br>atttcacaaataaagcatttttttcactgcattctagttgtggtttgtccaaactcatcaatgtatcttatcatgtctggatcaactgga<br>taactcaagctaaccaaaatcatcccaaacttcccaccccataccctattaccactgccaattacctagtggtttcatttactctaaa<br>cctgtgattcctctgaattattttcattttaaagaaattgtatttgttaaatatgtactacaaacttagtagttggaagggctaattca<br>ctcccaaagaagacaagatatccttgatctgtggatctaccacacacaaggctacttccctgattagcagaactacacaccaggg<br>ccaggggtcagatatccactgacctttggatggtgctacaagctagtaccagttgagccagataaggtagaagaggccaataaa<br>ggagagaacaccagcttgttacaccctgtgagcctgcatgggatggatgacccggagagagaagtgttagagtggaggtttgac<br>agccgcctagcatttcatcacgtggcccgagagctgcatccggagtacttcaagaactgctgatatcgagcttgctacaagggact<br>ttccgctggggactttccagggaggcgtggcctgggcgggactggggagtggcgagccctcagatcctgcatataagcagctgct |

| | |
|---|---|
| | ttttgcctgtactgggtctctctggttagaccagatctgagcctgggagctctctggctaactagggaacccactgcttaagcctcaa<br>taaagcttgccttgagtgcttcaagtagtgtgtgcccgtctgttgtgtgactctggtaactagagatccctcagacccttttagtcagt<br>gtggaaaatctctagcagtggcgcccgaacagggacttgaaagcgaaagggaaaccagaggagctctctcgacgcaggactcg<br>gcttgctgaagcgcgcacggcaagaggcgaggggcggcgactggtgagtacgccaaaaattttgactagcggaggctagaagg<br>agagagatgggtgcgagagcgtcagtattaagcgggggagaattagatcgcgatgggaaaaaattcggttaaggccaggggga<br>aagaaaaaatataaattaaaacatatagtatgggcaagcagggagctagaacgattcgcagttaatcctggcctgttagaaaca<br>tcagaaggctgtagacaaatactgggacagctacaaccatcccttcagacaggatcagaagaacttagatcattatataatacag<br>tagcaaccctctattgtgtgcatcaaaggatagagataaaagacaccaaggaagctttagacaagatagaggaagagcaaaac<br>aaaagtaagaccaccgcacagcaagcggccggccgctgatcttcagacctggaggaggagatatgagggacaattggagaagt<br>gaattatataaatataaagtagtaaaaattgaaccattaggagtagcacccaccaaggcaaagagaagagtggtgcagagaga<br>aaaaagagcagtgggaataggagctttgttccttgggttcttgggagcagcaggaagcactatgggcgcagcgtcaatgacgctg<br>acggtacaggccagacaattattgtctggtatagtgcagcagcagaacaatttgctgagggctattgaggcgcaacagcatctgtt<br>gcaactcacagtctggggcatcaagcagctccaggcaagaatcctggctgtgaaagatacctaaaggatcaacagctcctgggg<br>atttggggttgctctggaaaactcatttgcaccactgctgtgccttggaatgctagttggagtaataaatctctggaacagatttgg<br>aatcacacgacctggatggagtgggacagagaaattaacaattacacaagcttaatacactccttaattgaagaatcgcaaaac<br>cagcaagaaaagaatgaacaagaattattggaattagataaatgggcaagtttgtggaattggtttaacataacaaattggctgt<br>ggtatataaaattattcataatgatagtaggaggcttggtaggtttaagaatagtttttgctgtactttctatagtgaatagagttag<br>gcagggatattcaccattatcgtttcagacccacctcccaaccccgaggggacccgacaggcccgaaggaatagaagaagaag<br>gtggagagagagacagagacagatccattcgattagtgaacggatctcgacggtatcgccaaatggcagtattcatccacaattt<br>taaaagaaaaggggggattggggggtacagtgcaggggaaagaatagtagacataatagcaacagacatacaaactaaagaa<br>ttacaaaaacaaattacaaaaattcaaaattttcgggtttattacagggacagcagagatccagtttatcgatgaggccctttcgt<br>cttcactcgagtttactccctatcagtgatagagaacgtatgaagagtttactccctatcagtgatagagaacgtatgcagacttta<br>ctccctatcagtgatagagaacgtataaggagtttactccctatcagtgatagagaacgtatgaccagtttactccctatcagtgat<br>agagaacgtatctacagtttactccctatcagtgatagagaacgtatatccagtttactccctatcagtgatagagaacgtatgtcg<br>aggtaggcgtgtacggtgggcgcctataaaagcagagctcgtttagtgaaccgtcagatcgcctggagcaattccacaacactttt<br>gtcttatacttACGCGTGTCGACcatatgAGTTGAATACTCATACTCTTCCTTTTTCAATATTATTGAAGCA<br>TTTATCAGGGTTATTGTCTCATGAGCGGATACATATTTGAATGTATTTAGAAAAATAAACAAATAG<br>GGGTTCCGCGCACATTTCCCCGAAAAGTGCCACCTAAATTGTAAGCGTTAATATTTTGTTAAAATTC<br>GCGTTAAATTTTTGTTAAATCAGCTCATTTTTTAACCAATAGGCCGAAATCGGCAAAATCCCTTATA<br>AATCAAAAGTatgGGTAATTATAACCCGGGCCCTATATATTAGGCTAACTAGCTAACTAGGAATTCg<br>gtctcaCAGCAGCAGCAGCAGCAGCAGCAGCAGCAGCAGCAGCAGCACAGCAGCAGCAGCAG<br>CAGCAGCAGCAGCAGCAGCAGCAGCAGCAGCAGCAGCAGCAGCAGCAGCAGCAGCAGCAG<br>CAGCAGCAGCAGCAGCACAGCAGCAGCAGCAGCAGCAGCAGCAGCAGCAGCAGCAGCAGCAGC<br>AGCAGCAGCAGCAGCAGCAGCAGCAGCAGCAGCAGCAGCAGCAGCACAGCAGCAGCAGCA<br>GCAGCAGCAGCAGCAGCAGCAGCAGCAGCAGCAGCAGCAGCAGCAGCAGCAGCAGCAGCA<br>GCAGCAGCAGCAGCACAGCAGCAGCAGCAGCAGCAGCAGCAGCAGCAGCAGCAGCAGCAG<br>CAGCAGCAGCAGCAGCAGCAGCAGCAGCAGCAGCAGCAGCAGCACAGCAGCAGCAGCAGC<br>AGCAGCAGCAGCAGCAGCAGCAGCAGCAGCAGCAGCAGCAGCAGCAGCAGCAGCAGCAGC<br>AGCAGCAGCAGCAGCACAGCAGCAGCAGCAGCAGCAGCAGCAGCAGCAGCAGCAGCAGCA<br>GCAGCAGCAGCAGCAGCAGCAGCAGCACAGCAGCAGCAGCAGCAGCAGCAGCAGCAGCAG<br>CAGCAGCAGCAGCAGCAGCAGCAGCAGCAGCAGCAGCAGCAGCAGCAGCAGCAGCACAGC<br>AGCAGCAGCAGCAGCAGCAGCAGCAGCAGCAGCAGCAgagacgGAGCTCgtcgacGCggccgcaa<br>gcttggtaccgagctcGGATCCTAAGGTACCTAATTGCCTAGAAAACATGAGGATCACCCATGTCTGCAG<br>GTCGACTCTAGAAAACATGAGGATCACCCATGTCTGCAGTATTCCCGGGTTCATTAGATCCTAAGG<br>TACCTAATTGCCTAGAAAACATGAGGATCACCCATGTCTGCAGGTCGACTCTAGAAAACATGAGGA<br>TCACCCATGTCTGCAGTATTCCCGGGTTCATTAGATCCTAAGGTACCTAATTGCCTAGAAAACATGA<br>GGATCACCCATGTCTGCAGGTCGACTCTAGAAAACATGAGGATCACCCATGTCTGCAGTATTCCCG<br>GGTTCATTAGATCCTAAGGTACCTAATTGCCTAGAAAACATGAGGATCACCCATGTCTGCAGGTCG |

| | |
|---|---|
| | ACTCTAGAAAACATGAGGATCACCCATGTCTGCAGTATTCCCGGGTTCATTAGATCCTAAGGTACCT<br>AATTGCCTAGAAAACATGAGGATCACCCATGTCTGCAGGTCGACTCTAGAAAACATGAGGATCACC<br>CATGTCTGCAGTATTCCCGGGTTCATTAGATCCTAAGGTACCTAATTGCCTAGAAAACATGAGGAT<br>CACCCATGTCTGCAGGTCGACTCTAGAAAACATGAGGATCACCCATGTCTGCAGTATTCCCGGGTT<br>CATTAGATCTGCGCGCGATCGATATCAGCGCTTTAAATTTGCGCATGCTAGTgTAAggcatgcaagcttg<br>atatcaagcttatcgataatcaacctctggattacaaaatttgtgaaagattgactggtattcttaactatgttgctccttttacgcta<br>tgtggatacgctgctttaatgcctttgtatcatgctattgcttcccgtatggctttcattttctcctccttgtataaatcctggttgctgtc<br>tctttatgaggagttgtggcccgttgtcaggcaacgtggcgtggtgtgcactgtgtttgctgacgcaacccccactggttggggcatt<br>gccaccacctgtcagctcctttccgggactttcgctttccccctccctattgccacggcggaactcatcgccgcctgccttgcccgct<br>gctggacaggggctcggctgttgggcactgacaattccgtggtgttgtcggggaaatcatcgtcctttccttggctgctcgcctgtgt<br>tgccacctggattctgcgcgggacgtccttctgctacgtcccttcggccctcaatccagcggaccttccttcccgcggcctgctgccg<br>gctctgcggcctcttccgcgtcttcgccttcgccctcagacgagtcggatctccctttgggccgcctccccgcatcgataccgtcgac<br>ctcgagggaattaattcgagctcggtacctttaagaccaatgacttacaaggcagctgtagatcttagccactttttaaaagaaaa<br>ggggggactggaagggctaattcactcccaacgaagacaagatctgctttttgcttgtactgggtctctctggttagaccagatctg<br>agcctgggagctctctggctaactagggaacccactgcttaagcctcaataaagcttgccttgagtgcttcaagtagtgtgtgcccg<br>tctgttgtgtgactctggtaactagagatccctcagacccttttagtcagtgtggaaaatctctagcagcatctagaattaattccgt<br>gtatt |
| N2: pHR Tre3G Flank N2 240xCAG 12xMS2 WPRE | ctatagtgtcacctaaatcgtatgtgtatgatacataaggttatgtattaattgtagccgcgttctaacgacaatatgtacaagccta<br>attgtgtagcatctggcttactgaagcagaccctatcatctctctcgtaaactgccgtcagagtcggtttggttggacgaaccttctg<br>agtttctggtaacgccgtcccgcacccggaaatggtcagcgaaccaatcagcagggtcatcgctagccagatcctctacgccgga<br>cgcatcgtggccggcatcaccggcgccacaggtgcggttgctggcgcctatatcgccgacatcaccgatggggaagatcgggctc<br>gccacttcgggctcatgagcgcttgtttcggcgtgggtatggtggcaggccccgtggccgggggactgttgggcgccatctccttgc<br>atgcaccattccttgcggcggcggtgctcaacggcctcaacctactactgggctgcttcctaatgcaggagtcgcataagggagag<br>cgtcgaatggtgcactctcagtacaatctgctctgatgccgcatagttaagccagccccgacacccgccaacacccgctgacgcgc<br>cctgacgggcttgtctgctcccggcatccgcttacagacaagctgtgaccgtctccgggagctgcatgtgtcagaggttttcaccgt<br>catcaccgaaacgcgcgagacgaaagggcctcgtgatacgcctatttttataggttaatgtcatgataataatggtttcttagacgt<br>caggtggcacttttcggggaaatgtgcgcggaacccctatttgtttatttttctaaatacattcaaatatgtatccgctcatgagaca<br>ataaccctgataaatgcttcaataatattgaaaaaggaagagtatgagtattcaacatttccgtgtcgcccttattcccttttttgcg<br>gcattttgccttcctgtttttgctcacccagaaacgctggtgaaagtaaaagatgctgaagatcagttgggtgcacgagtgggttac<br>atcgaactggatctcaacagcggtaagatccttgagagttttcgccccgaagaacgttttccaatgatgagcacttttaaagttctg<br>ctatgtggcgcggtattatcccgtattgacgccgggcaagagcaactcggtcgccgcatacactattctcagaatgacttggttga<br>gtactcaccagtcacagaaaagcatcttacggatggcatgacagtaagagaattatgcagtgctgccataaccatgagtgataac<br>actgcggccaacttacttctgacaacgatcggaggaccgaaggagctaaccgcttttttgcacaacatgggggatcatgtaactcg<br>ccttgatcgttgggaaccggagctgaatgaagccataccaaacgacgagcgtgacaccacgatgcctgtagcaatggcaacaac<br>gttgcgcaaactattaactggcgaactacttactctagcttcccggcaacaattaatagactggatggaggcggataaagttgcag<br>gaccacttctgcgctcggcccttccggctggctggtttattgctgataaatctggagccggtgagcgtgggtctcgcggtatcattgc<br>agcactggggccagatggtaagccctcccgtatcgtagttatctacacgacggggagtcaggcaactatggatgaacgaaatag<br>acagatcgctgagataggtgcctcactgattaagcattggtaactgtcagaccaagtttactcatatatactttagattgatttaaa<br>acttcatttttaatttaaaaggatctaggtgaagatcctttttgataatctcatgaccaaaatcccttaacgtgagttttcgttccact<br>gagcgtcagaccccgtagaaaagatcaaaggatcttcttgagatcctttttttctgcgcgtaatctgctgcttgcaaacaaaaaaa<br>ccaccgctaccagcggtggtttgtttgccggatcaagagctaccaactctttttccgaaggtaactggcttcagcagagcgcagat<br>accaaatactgtctttctagtgtagccgtagttaggccaccacttcaagaactctgtagcaccgcctacatacctcgctctgctaatc<br>ctgttaccagtggctgctgccagtggcgataagtcgtgtcttaccgggttggactcaagacgatagttaccggataaggcgcagcg<br>gtcgggctgaacggggggttcgtgcacacagcccagcttggagcgaacgacctacaccgaactgagatacctacagcgtgagct<br>atgagaaagcgccacgcttcccgaagggagaaaggcggacaggtatccggtaagcggcagggtcggaacaggagagcgcacg<br>agggagcttccagggggaaacgcctggtatctttatagtcctgtcgggtttcgccacctctgacttgagcgtcgatttttgtgatgct<br>cgtcaggggggcggagcctatggaaaaacgccagcaacgcggcctttttacggttcctggccttttgctggccttttgctcacatgt |

| | |
|---|---|
| | tctttcctgcgttatcccctgattctgtggataaccgtattaccgcctttgagtgagctgataccgctcgccgcagccgaacgaccga<br>gcgcagcgagtcagtgagcgaggaagcggaagagcgcccaatacgcaaaccgcctctccccgcgcgttggccgattcattaatg<br>cagctgtggaatgtgtgtcagttagggtgtggaaagtccccaggctccccagcaggcagaagtatgcaaagcatgcatctcaatt<br>agtcagcaaccaggtgtggaaagtccccaggctccccagcaggcagaagtatgcaaagcatgcatctcaattagtcagcaacca<br>tagtcccgcccctaactccgcccatcccgcccctaactccgcccagttccgcccattctccgccccatggctgactaattttttttattt<br>atgcagaggccgaggccgcctcggcctctgagctattccagaagtagtgaggaggcttttttggaggcctaggcttttgcaaaaag<br>cttggacacaagacaggcttgcgagatatgtttgagaataccactttatcccgcgtcagggagaggcagtgcgtaaaaagacgc<br>ggactcatgtgaaatactggtttttagtgcgccagatctctataatctcgcgcaacctattttcccctcgaacactttttaagccgtag<br>ataaacaggctgggacacttcacatgagcgaaaaatacatcgtcacctgggacatgttgcagatccatgcacgtaaactcgcaa<br>gccgactgatgccttctgaacaatggaaaggcattattgccgtaagccgtggcggtctgtaccgggtgcgttactggcgcgtgaac<br>tgggtattcgtcatgtcgataccgtttgtatttccagctacgatcacgacaaccagcgcgagcttaaagtgctgaaacgcgcagaa<br>ggcgatggcgaaggcttcatcgttattgatgacctggtggataccggtggtactgcggttgcgattcgtgaaatgtatccaaaagc<br>gcactttgtcaccatcttcgcaaaaccggctggtcgtccgctggttgatgactatgttgttgatatcccgcaagatacctggattgaa<br>cagccgtgggatatgggcgtcgtattcgtcccgccaatctccggtcgctaatcttttcaacgcctggcactgccgggcgttgttctttt<br>taacttcaggcgggttacaatagtttccagtaagtattctggaggctgcatccatgacacaggcaaacctgagcgaaaccctgttc<br>aaaccccgctttaaacatcctgaaacctcgacgctagtccgccgctttaatcacggcgcacaaccgcctgtgcagtcggcccttga<br>tggtaaaaccatccctcactggtatcgcatgattaaccgtctgatgtggatctggcgcggcattgacccacgcgaaatcctcgacg<br>tccaggcacgtattgtgatgagcgatgccgaacgtaccgacgatgatttatacgatacggtgattggctaccgtggcggcaactgg<br>atttatgagtgggccccggatctttgtgaaggaaccttacttctgtggtgtgacataattggacaaactacctacagagatttaaag<br>ctctaaggtaaatataaaatttttaagtgtataatgtgttaaactactgattctaattgtttgtgtattttagattccaacctatggaa<br>ctgatgaatgggagcagtggtggaatgcctttaatgaggaaaacctgttttgctcagaagaaatgccatctagtgatgatgaggct<br>actgctgactctcaacattctactcctccaaaaaagaagagaaaggtagaagaccccaaggactttccttcagaattgctaagttt<br>tttgagtcatgctgtgtttagtaatagaactcttgcttgctttgctatttacaccacaaaggaaaaagctgcactgctatacaagaa<br>aattatggaaaaatattctgtaacctttataagtaggcataacagttataatcataacatactgttttttcttactccacacaggcat<br>agagtgtctgctattaataactatgctcaaaaattgtgtacctttagctttttaatttgtaaaggggttaataaggaatatttgatgta<br>tagtgccttgactagagatcataatcagccataccacatttgtagaggttttacttgctttaaaaaacctcccacacctccccctga<br>acctgaaacataaaatgaatgcaattgttgttgttaacttgtttattgcagcttataatggttacaaataaagcaatagcatcacaa<br>atttcacaaataaagcatttttttcactgcattctagttgtggtttgtccaaactcatcaatgtatcttatcatgtctggatcaactgga<br>taactcaagctaaccaaaatcatcccaaacttcccaccccataccctattaccactgccaattacctagtggtttcatttactctaaa<br>cctgtgattcctctgaattattttcattttaaagaaattgtatttgttaaatatgtactacaaacttagtagttggaagggctaattca<br>ctcccaaagaagacaagatatccttgatctgtggatctaccacacacaaggctacttccctgattagcagaactacacaccaggg<br>ccaggggtcagatatccactgacctttggatggtgctacaagctagtaccagttgagccagataaggtagaagaggccaataaa<br>ggagagaacaccagcttgttacaccctgtgagcctgcatgggatggatgacccggagagagaagtgttagagtggaggtttgac<br>agccgcctagcatttcatcacgtggcccgagagctgcatccggagtacttcaagaactgctgatatcgagcttgctacaagggact<br>ttccgctggggactttccagggaggcgtggcctgggcgggactggggagtggcgagccctcagatcctgcatataagcagctgct<br>ttttgcctgtactgggtctctctggttagaccagatctgagcctgggagctctctggctaactagggaacccactgcttaagcctcaa<br>taaagcttgccttgagtgcttcaagtagtgtgtgcccgtctgttgtgtgactctggtaactagagatccctcagacccttttagtcagt<br>gtggaaaatctctagcagtggcgcccgaacagggacttgaaagcgaaagggaaaccagaggagctctctcgacgcaggactcg<br>gcttgctgaagcgcgcacggcaagaggcgaggggcggcgactggtgagtacgccaaaaattttgactagcggaggctagaagg<br>agagagatgggtgcgagagcgtcagtattaagcgggggagaattagatcgcgatgggaaaaaattcggttaaggccaggggga<br>aagaaaaaatataaattaaaacatatagtatgggcaagcagggagctagaacgattcgcagttaatcctggcctgttagaaaca<br>tcagaaggctgtagacaaatactgggacagctacaaccatcccttcagacaggatcagaagaacttagatcattatataatacag<br>tagcaaccctctattgtgtgcatcaaaggatagagataaaagacaccaaggaagctttagacaagatagaggaagagcaaaac<br>aaaagtaagaccaccgcacagcaagcggccggccgctgatcttcagacctggaggaggagatatgagggacaattggagaagt<br>gaattatataaatataaagtagtaaaaattgaaccattaggagtagcacccaccaaggcaaagagaagagtggtgcagagaga<br>aaaaagagcagtgggaataggagctttgttccttgggttcttgggagcagcaggaagcactatgggcgcagcgtcaatgacgctg<br>acggtacaggccagacaattattgtctggtatagtgcagcagcagaacaatttgctgagggctattgaggcgcaacagcatctgtt<br>gcaactcacagtctggggcatcaagcagctccaggcaagaatcctggctgtgaaagatacctaaaggatcaacagctcctgggg |

| | |
|---|---|
| | atttggggttgctctggaaaactcatttgcaccactgctgtgccttggaatgctagttggagtaataaatctctggaacagatttgg<br>aatcacacgacctggatggagtgggacagagaaattaacaattacacaagcttaatacactccttaattgaagaatcgcaaaac<br>cagcaagaaaagaatgaacaagaattattggaattagataaatgggcaagtttgtggaattggtttaacataacaaattggctgt<br>ggtatataaaattattcataatgatagtaggaggcttggtaggtttaagaatagtttttgctgtactttctatagtgaatagagttag<br>gcagggatattcaccattatcgtttcagacccacctcccaaccccgaggggacccgacaggcccgaaggaatagaagaagaag<br>gtggagagagagacagagacagatccattcgattagtgaacggatctcgacggtatcgccaaatggcagtattcatccacaattt<br>taaaagaaaaggggggattggggggtacagtgcaggggaaagaatagtagacataatagcaacagacatacaaactaaagaa<br>ttacaaaaacaaattacaaaaattcaaaattttcgggtttattacagggacagcagagatccagtttatcgatgaggccctttcgt<br>cttcactcgagtttactccctatcagtgatagagaacgtatgaagagtttactccctatcagtgatagagaacgtatgcagacttta<br>ctccctatcagtgatagagaacgtataaggagtttactccctatcagtgatagagaacgtatgaccagtttactccctatcagtgat<br>agagaacgtatctacagtttactccctatcagtgatagagaacgtatatccagtttactccctatcagtgatagagaacgtatgtcg<br>aggtaggcgtgtacggtgggcgcctataaaagcagagctcgtttagtgaaccgtcagatcgcctggagcaattccacaacactttt<br>gtcttatacttACGCGTGTCGACcatatgACGCCGCCCAGGAGCCGCCCGCGCTCCCTGAACCCTAGAAC<br>TGTCTTCGACTCCGGGGCCCCGTTGGAAGACTGAGTGCCCGGGGCACGGCACAGAAGCCGCGCCC<br>ACCGCCTGCCAGTTCACAACCGCTCCGAGCGTGGGTCTCCGCCCAGCTCCAGTCCTGTGATCCGGG<br>CCCGCCCCCTAGCGGCCGGGGAGGGAGGGGCCGGGTCCGCGGCCGGCGAACGGGGCTCGAAGG<br>GTCCTTGTAGCCGGGATatgGGTAATTATAACCCGGGCCCTATATATTAGGCTAACTAGCTAACTAG<br>GAATTCggtctcaCAGCAGCAGCAGCAGCAGCAGCAGCAGCAGCAGCAGCAGCACAGCAGCAG<br>CAGCAGCAGCAGCAGCAGCAGCAGCAGCAGCAGCAGCAGCAGCAGCAGCAGCAGCAGCAG<br>CAGCAGCAGCAGCAGCAGCAGCACAGCAGCAGCAGCAGCAGCAGCAGCAGCAGCAGCAGC<br>AGCAGCAGCAGCAGCAGCAGCAGCAGCAGCAGCAGCAGCAGCAGCAGCAGCACAGCAGCA<br>GCAGCAGCAGCAGCAGCAGCAGCAGCAGCAGCAGCAGCAGCAGCAGCAGCAGCAGCAGCA<br>GCAGCAGCAGCAGCAGCAGCACAGCAGCAGCAGCAGCAGCAGCAGCAGCAGCAGCAGCAG<br>CAGCAGCAGCAGCAGCAGCAGCAGCAGCAGCAGCAGCAGCAGCAGCAGCACAGCAGCAGC<br>AGCAGCAGCAGCAGCAGCAGCAGCAGCAGCAGCAGCAGCAGCAGCAGCAGCAGCAGCAGC<br>AGCAGCAGCAGCAGCAGCAGCACAGCAGCAGCAGCAGCAGCAGCAGCAGCAGCAGCAGCA<br>GCAGCAGCAGCAGCAGCAGCAGCAGCAGCAGCACAGCAGCAGCAGCAGCAGCAGCAGCAG<br>CAGCAGCAGCAGCAGCAGCAGCAGCAGCAGCAGCAGCAGCAGCAGCAGCAGCAGCAGCAG<br>CACAGCAGCAGCAGCAGCAGCAGCAGCAGCAGCAGCAGCAGCAgagacgGAGCTCgtcgacGC<br>ggccgcaagcttggtaccgagctcGGATCCTAAGGTACCTAATTGCCTAGAAAACATGAGGATCACCCATG<br>TCTGCAGGTCGACTCTAGAAAACATGAGGATCACCCATGTCTGCAGTATTCCCGGGTTCATTAGAT<br>CCTAAGGTACCTAATTGCCTAGAAAACATGAGGATCACCCATGTCTGCAGGTCGACTCTAGAAAAC<br>ATGAGGATCACCCATGTCTGCAGTATTCCCGGGTTCATTAGATCCTAAGGTACCTAATTGCCTAGAA<br>AACATGAGGATCACCCATGTCTGCAGGTCGACTCTAGAAAACATGAGGATCACCCATGTCTGCAGT<br>ATTCCCGGGTTCATTAGATCCTAAGGTACCTAATTGCCTAGAAAACATGAGGATCACCCATGTCTGC<br>AGGTCGACTCTAGAAAACATGAGGATCACCCATGTCTGCAGTATTCCCGGGTTCATTAGATCCTAA<br>GGTACCTAATTGCCTAGAAAACATGAGGATCACCCATGTCTGCAGGTCGACTCTAGAAAACATGAG<br>GATCACCCATGTCTGCAGTATTCCCGGGTTCATTAGATCCTAAGGTACCTAATTGCCTAGAAAACAT<br>GAGGATCACCCATGTCTGCAGGTCGACTCTAGAAAACATGAGGATCACCCATGTCTGCAGTATTCC<br>CGGGTTCATTAGATCTGCGCGCGATCGATATCAGCGCTTTAAATTTGCGCATGCTAGTgTAAggcatg<br>caagcttgatatcaagcttatcgataatcaacctctggattacaaaatttgtgaaagattgactggtattcttaactatgttgctcctt<br>ttacgctatgtggatacgctgctttaatgcctttgtatcatgctattgcttcccgtatggctttcattttctcctccttgtataaatcctg<br>gttgctgtctctttatgaggagttgtggcccgttgtcaggcaacgtggcgtggtgtgcactgtgtttgctgacgcaacccccactggt<br>tggggcattgccaccacctgtcagctcctttccgggactttcgctttccccctccctattgccacggcggaactcatcgccgcctgcc<br>ttgcccgctgctggacaggggctcggctgttgggcactgacaattccgtggtgttgtcggggaaatcatcgtcctttccttggctgct<br>cgcctgtgttgccacctggattctgcgcgggacgtccttctgctacgtcccttcggccctcaatccagcggaccttccttcccgcggc<br>ctgctgccggctctgcggcctcttccgcgtcttcgccttcgccctcagacgagtcggatctccctttgggccgcctccccgcatcgat<br>accgtcgacctcgagggaattaattcgagctcggtacctttaagaccaatgacttacaaggcagctgtagatcttagccactttta |

| | |
|---|---|
| | aaagaaaagggggggactggaagggctaattcactcccaacgaagacaagatctgctttttgcttgtactgggtctctctggttaga<br>ccagatctgagcctgggagctctctggctaactagggaacccactgcttaagcctcaataaagcttgccttgagtgcttcaagtagt<br>gtgtgcccgtctgttgtgtgactctggtaactagagatccctcagacccttttagtcagtgtggaaaatctctagcagcatctagaat<br>taattccgtgtatt |
| N3: pHR Tre3G Flank N3 240xCAG 12xMS2 WPRE | ctatagtgtcacctaaatcgtatgtgtatgatacataaggttatgtattaattgtagccgcgttctaacgacaatatgtacaagccta<br>attgtgtagcatctggcttactgaagcagaccctatcatctctctcgtaaactgccgtcagagtcggtttggttggacgaaccttctg<br>agtttctggtaacgccgtcccgcacccggaaatggtcagcgaaccaatcagcagggtcatcgctagccagatcctctacgccgga<br>cgcatcgtggccggcatcaccggcgccacaggtgcggttgctggcgcctatatcgccgacatcaccgatggggaagatcgggctc<br>gccacttcgggctcatgagcgcttgtttcggcgtgggtatggtggcaggccccgtggccgggggactgttgggcgccatctccttgc<br>atgcaccattccttgcggcggcggtgctcaacggcctcaacctactactgggctgcttcctaatgcaggagtcgcataagggagag<br>cgtcgaatggtgcactctcagtacaatctgctctgatgccgcatagttaagccagccccgacacccgccaacacccgctgacgcgc<br>cctgacgggcttgtctgctcccggcatccgcttacagacaagctgtgaccgtctccgggagctgcatgtgtcagaggttttcaccgt<br>catcaccgaaacgcgcgagacgaaagggcctcgtgatacgcctatttttataggttaatgtcatgataataatggtttcttagacgt<br>caggtggcacttttcggggaaatgtgcgcggaacccctatttgtttatttttctaaatacattcaaatatgtatccgctcatgagaca<br>ataaccctgataaatgcttcaataatattgaaaaaggaagagtatgagtattcaacatttccgtgtcgcccttattcccttttttgcg<br>gcattttgccttcctgtttttgctcacccagaaacgctggtgaaagtaaaagatgctgaagatcagttgggtgcacgagtgggttac<br>atcgaactggatctcaacagcggtaagatccttgagagttttcgccccgaagaacgttttccaatgatgagcacttttaaagttctg<br>ctatgtggcgcggtattatcccgtattgacgccgggcaagagcaactcggtcgccgcatacactattctcagaatgacttggttga<br>gtactcaccagtcacagaaaagcatcttacggatggcatgacagtaagagaattatgcagtgctgccataaccatgagtgataac<br>actgcggccaacttacttctgacaacgatcggaggaccgaaggagctaaccgcttttttgcacaacatgggggatcatgtaactcg<br>ccttgatcgttgggaaccggagctgaatgaagccataccaaacgacgagcgtgacaccacgatgcctgtagcaatggcaacaac<br>gttgcgcaaactattaactggcgaactacttactctagcttcccggcaacaattaatagactggatggaggcggataaagttgcag<br>gaccacttctgcgctcggcccttccggctggctggtttattgctgataaatctggagccggtgagcgtgggtctcgcggtatcattgc<br>agcactggggccagatggtaagccctcccgtatcgtagttatctacacgacggggagtcaggcaactatggatgaacgaaatag<br>acagatcgctgagataggtgcctcactgattaagcattggtaactgtcagaccaagtttactcatatatactttagattgatttaaa<br>acttcatttttaatttaaaaggatctaggtgaagatcctttttgataatctcatgaccaaaatcccttaacgtgagttttcgttccact<br>gagcgtcagaccccgtagaaaagatcaaaggatcttcttgagatcctttttttctgcgcgtaatctgctgcttgcaaacaaaaaaa<br>ccaccgctaccagcggtggtttgtttgccggatcaagagctaccaactctttttccgaaggtaactggcttcagcagagcgcagat<br>accaaatactgtctttctagtgtagccgtagttaggccaccacttcaagaactctgtagcaccgcctacatacctcgctctgctaatc<br>ctgttaccagtggctgctgccagtggcgataagtcgtgtcttaccgggttggactcaagacgatagttaccggataaggcgcagcg<br>gtcgggctgaacggggggttcgtgcacacagcccagcttggagcgaacgacctacaccgaactgagatacctacagcgtgagct<br>atgagaaagcgccacgcttcccgaagggagaaaggcggacaggtatccggtaagcggcagggtcggaacaggagagcgcacg<br>agggagcttccagggggaaacgcctggtatctttatagtcctgtcgggtttcgccacctctgacttgagcgtcgatttttgtgatgct<br>cgtcaggggggcggagcctatggaaaaacgccagcaacgcggcctttttacggttcctggccttttgctggccttttgctcacatgt<br>tctttcctgcgttatcccctgattctgtggataaccgtattaccgcctttgagtgagctgataccgctcgccgcagccgaacgaccga<br>gcgcagcgagtcagtgagcgaggaagcggaagagcgcccaatacgcaaaccgcctctccccgcgcgttggccgattcattaatg<br>cagctgtggaatgtgtgtcagttagggtgtggaaagtccccaggctccccagcaggcagaagtatgcaaagcatgcatctcaatt<br>agtcagcaaccaggtgtggaaagtccccaggctccccagcaggcagaagtatgcaaagcatgcatctcaattagtcagcaacca<br>tagtcccgcccctaactccgcccatcccgcccctaactccgcccagttccgcccattctccgccccatggctgactaattttttttattt<br>atgcagaggccgaggccgcctcggcctctgagctattccagaagtagtgaggaggcttttttggaggcctaggcttttgcaaaaag<br>cttggacacaagacaggcttgcgagatatgtttgagaataccactttatcccgcgtcagggagaggcagtgcgtaaaaagacgc<br>ggactcatgtgaaatactggtttttagtgcgccagatctctataatctcgcgcaacctattttcccctcgaacactttttaagccgtag<br>ataaacaggctgggacacttcacatgagcgaaaaatacatcgtcacctgggacatgttgcagatccatgcacgtaaactcgcaa<br>gccgactgatgccttctgaacaatggaaaggcattattgccgtaagccgtggcggtctgtaccgggtgcgttactggcgcgtgaac<br>tgggtattcgtcatgtcgataccgtttgtatttccagctacgatcacgacaaccagcgcgagcttaaagtgctgaaacgcgcagaa<br>ggcgatggcgaaggcttcatcgttattgatgacctggtggataccggtggtactgcggttgcgattcgtgaaatgtatccaaaagc<br>gcactttgtcaccatcttcgcaaaaccggctggtcgtccgctggttgatgactatgttgttgatatcccgcaagatacctggattgaa |

| | |
|---|---|
| | cagccgtgggatatgggcgtcgtattcgtcccgccaatctccggtcgctaatcttttcaacgcctggcactgccgggcgttgttctttt<br>taacttcaggcgggttacaatagtttccagtaagtattctggaggctgcatccatgacacaggcaaacctgagcgaaaccctgttc<br>aaaccccgctttaaacatcctgaaacctcgacgctagtccgccgctttaatcacggcgcacaaccgcctgtgcagtcggcccttga<br>tggtaaaaccatccctcactggtatcgcatgattaaccgtctgatgtggatctggcgcggcattgacccacgcgaaatcctcgacg<br>tccaggcacgtattgtgatgagcgatgccgaacgtaccgacgatgatttatacgatacggtgattggctaccgtggcggcaactgg<br>atttatgagtgggccccggatctttgtgaaggaaccttacttctgtggtgtgacataattggacaaactacctacagagatttaaag<br>ctctaaggtaaatataaaatttttaagtgtataatgtgttaaactactgattctaattgtttgtgtattttagattccaacctatggaa<br>ctgatgaatgggagcagtggtggaatgcctttaatgaggaaaacctgttttgctcagaagaaatgccatctagtgatgatgaggct<br>actgctgactctcaacattctactcctccaaaaaagaagagaaaggtagaagaccccaaggactttccttcagaattgctaagttt<br>tttgagtcatgctgtgtttagtaatagaactcttgcttgctttgctatttacaccacaaaggaaaaagctgcactgctatacaagaa<br>aattatggaaaaatattctgtaacctttataagtaggcataacagttataatcataacatactgttttttcttactccacacaggcat<br>agagtgtctgctattaataactatgctcaaaaattgtgtacctttagctttttaatttgtaaaggggttaataaggaatatttgatgta<br>tagtgccttgactagagatcataatcagccataccacatttgtagaggttttacttgctttaaaaaacctcccacacctccccctga<br>acctgaaacataaaatgaatgcaattgttgttgttaacttgtttattgcagcttataatggttacaaataaagcaatagcatcacaa<br>atttcacaaataaagcatttttttcactgcattctagttgtggtttgtccaaactcatcaatgtatcttatcatgtctggatcaactgga<br>taactcaagctaaccaaaatcatcccaaacttcccaccccataccctattaccactgccaattacctagtggtttcatttactctaaa<br>cctgtgattcctctgaattattttcattttaaagaaattgtatttgttaaatatgtactacaaacttagtagttggaagggctaattca<br>ctcccaaagaagacaagatatccttgatctgtggatctaccacacacaaggctacttccctgattagcagaactacacaccaggg<br>ccaggggtcagatatccactgacctttggatggtgctacaagctagtaccagttgagccagataaggtagaagaggccaataaa<br>ggagagaacaccagcttgttacaccctgtgagcctgcatgggatggatgacccggagagagaagtgttagagtggaggtttgac<br>agccgcctagcatttcatcacgtggcccgagagctgcatccggagtacttcaagaactgctgatatcgagcttgctacaagggact<br>ttccgctggggactttccagggaggcgtggcctgggcgggactggggagtggcgagccctcagatcctgcatataagcagctgct<br>ttttgcctgtactgggtctctctggttagaccagatctgagcctgggagctctctggctaactagggaacccactgcttaagcctcaa<br>taaagcttgccttgagtgcttcaagtagtgtgtgcccgtctgttgtgtgactctggtaactagagatccctcagacccttttagtcagt<br>gtggaaaatctctagcagtggcgcccgaacagggacttgaaagcgaaagggaaaccagaggagctctctcgacgcaggactcg<br>gcttgctgaagcgcgcacggcaagaggcgaggggcggcgactggtgagtacgccaaaaattttgactagcggaggctagaagg<br>agagagatgggtgcgagagcgtcagtattaagcgggggagaattagatcgcgatgggaaaaaattcggttaaggccagggggа<br>aagaaaaaatataaattaaaacatatagtatgggcaagcagggagctagaacgattcgcagttaatcctggcctgttagaaaca<br>tcagaaggctgtagacaaatactgggacagctacaaccatcccttcagacaggatcagaagaacttagatcattatataatacag<br>tagcaaccctctattgtgtgcatcaaaggatagagataaaagacaccaaggaagctttagacaagatagaggaagagcaaaac<br>aaaagtaagaccaccgcacagcaagcggccggccgctgatcttcagacctggaggaggagatatgagggacaattggagaagt<br>gaattatataaatataaagtagtaaaaattgaaccattaggagtagcacccaccaaggcaaagagaagagtggtgcagagaga<br>aaaaagagcagtgggaataggagctttgttccttgggttcttgggagcagcaggaagcactatgggcgcagcgtcaatgacgctg<br>acggtacaggccagacaattattgtctggtatagtgcagcagcagaacaatttgctgagggctattgaggcgcaacagcatctgtt<br>gcaactcacagtctggggcatcaagcagctccaggcaagaatcctggctgtgaaagatacctaaaggatcaacagctcctgggg<br>atttggggttgctctggaaaactcatttgcaccactgctgtgccttggaatgctagttggagtaataaatctctggaacagatttgg<br>aatcacacgacctggatggagtgggacagagaaattaacaattacacaagcttaatacactccttaattgaagaatcgcaaaac<br>cagcaagaaaagaatgaacaagaattattggaattagataaatgggcaagtttgtggaattggtttaacataacaaattggctgt<br>ggtatataaaattattcataatgatagtaggaggcttggtaggtttaagaatagtttttgctgtactttctatagtgaatagagttag<br>gcagggatattcaccattatcgtttcagacccacctcccaaccccgaggggacccgacaggcccgaaggaatagaagaagaag<br>gtggagagagagacagagacagatccattcgattagtgaacggatctcgacggtatcgccaaatggcagtattcatccacaattt<br>taaaagaaaaggggggattggggggtacagtgcaggggaaagaatagtagacataatagcaacagacatacaaactaaagaa<br>ttacaaaaacaaattacaaaaattcaaaattttcgggtttattacagggacagcagagatccagtttatcgatgaggccctttcgt<br>cttcactcgagtttactccctatcagtgatagagaacgtatgaagagtttactccctatcagtgatagagaacgtatgcagacttta<br>ctccctatcagtgatagagaacgtataaggagtttactccctatcagtgatagagaacgtatgaccagtttactccctatcagtgat<br>agagaacgtatctacagtttactccctatcagtgatagagaacgtatatccagtttactccctatcagtgatagagaacgtatgtcg<br>aggtaggcgtgtacggtgggcgcctataaaagcagagctcgtttagtgaaccgtcagatcgcctggagcaattccacaacactttt<br>gtcttatacttACGCGTGTCGACcatatgAGCTCCGGATCCCTTCGATCCACGACCCATCCTGGACGACTA |

| | |
|---|---|
| | CCCGGGTATTTCTTCGGTATTGAGTATACGACCGCGTAGTGGACCGATACCTAGGCGGCAGAACTA<br>TCTTATACAAATAACACGAGGCCTAGTGGCTGATCTAAGCAGCAAGTCATGGGCCCAAGCACTACG<br>GACAAAATGGCCTGGGGTTGGGAGCAGTAGGTACGGTACCCCGTATAGTGCGCAGATCTAGTTCG<br>CGGGATCGGTCGTatgGGTAATTATAACCCGGGCCCTATATATTAGGCTAACTAGCTAACTAGGAAT<br>TCggtctcaCAGCAGCAGCAGCAGCAGCAGCAGCAGCAGCAGCAGCAGCACAGCAGCAGCAGC<br>AGCAGCAGCAGCAGCAGCAGCAGCAGCAGCAGCAGCAGCAGCAGCAGCAGCAGCAGCAGC<br>AGCAGCAGCAGCAGCAGCACAGCAGCAGCAGCAGCAGCAGCAGCAGCAGCAGCAGCAGCA<br>GCAGCAGCAGCAGCAGCAGCAGCAGCAGCAGCAGCAGCAGCAGCAGCACAGCAGCAGCAG<br>CAGCAGCAGCAGCAGCAGCAGCAGCAGCAGCAGCAGCAGCAGCAGCAGCAGCAGCAGCAG<br>CAGCAGCAGCAGCAGCACAGCAGCAGCAGCAGCAGCAGCAGCAGCAGCAGCAGCAGCAGC<br>AGCAGCAGCAGCAGCAGCAGCAGCAGCAGCAGCAGCAGCAGCAGCACAGCAGCAGCAGCA<br>GCAGCAGCAGCAGCAGCAGCAGCAGCAGCAGCAGCAGCAGCAGCAGCAGCAGCAGCAGCA<br>GCAGCAGCAGCAGCAGCACAGCAGCAGCAGCAGCAGCAGCAGCAGCAGCAGCAGCAGCAG<br>CAGCAGCAGCAGCAGCAGCAGCAGCAGCACAGCAGCAGCAGCAGCAGCAGCAGCAGCAGC<br>AGCAGCAGCAGCAGCAGCAGCAGCAGCAGCAGCAGCAGCAGCAGCAGCAGCAGCAGCACA<br>GCAGCAGCAGCAGCAGCAGCAGCAGCAGCAGCAGCAGCAgagacgGAGCTCgtcgacGCggccgc<br>aagcttggtaccgagctcGGATCCTAAGGTACCTAATTGCCTAGAAAACATGAGGATCACCCATGTCTGC<br>AGGTCGACTCTAGAAAACATGAGGATCACCCATGTCTGCAGTATTCCCGGGTTCATTAGATCCTAA<br>GGTACCTAATTGCCTAGAAAACATGAGGATCACCCATGTCTGCAGGTCGACTCTAGAAAACATGAG<br>GATCACCCATGTCTGCAGTATTCCCGGGTTCATTAGATCCTAAGGTACCTAATTGCCTAGAAAACAT<br>GAGGATCACCCATGTCTGCAGGTCGACTCTAGAAAACATGAGGATCACCCATGTCTGCAGTATTCC<br>CGGGTTCATTAGATCCTAAGGTACCTAATTGCCTAGAAAACATGAGGATCACCCATGTCTGCAGGT<br>CGACTCTAGAAAACATGAGGATCACCCATGTCTGCAGTATTCCCGGGTTCATTAGATCCTAAGGTA<br>CCTAATTGCCTAGAAAACATGAGGATCACCCATGTCTGCAGGTCGACTCTAGAAAACATGAGGATC<br>ACCCATGTCTGCAGTATTCCCGGGTTCATTAGATCCTAAGGTACCTAATTGCCTAGAAAACATGAG<br>GATCACCCATGTCTGCAGGTCGACTCTAGAAAACATGAGGATCACCCATGTCTGCAGTATTCCCGG<br>GTTCATTAGATCTGCGCGCGATCGATATCAGCGCTTTAAATTTGCGCATGCTAGTgTAAggcatgcaag<br>cttgatatcaagcttatcgataatcaacctctggattacaaaatttgtgaaagattgactggtattcttaactatgttgctccttttac<br>gctatgtggatacgctgctttaatgcctttgtatcatgctattgcttcccgtatggctttcattttctcctccttgtataaatcctggttg<br>ctgtctctttatgaggagttgtggcccgttgtcaggcaacgtggcgtggtgtgcactgtgtttgctgacgcaacccccactggttggg<br>gcattgccaccacctgtcagctcctttccgggactttcgctttccccctccctattgccacggcggaactcatcgccgcctgccttgcc<br>cgctgctggacaggggctcggctgttgggcactgacaattccgtggtgttgtcggggaaatcatcgtcctttccttggctgctcgcct<br>gtgttgccacctggattctgcgcgggacgtccttctgctacgtcccttcggccctcaatccagcggaccttccttcccgcggcctgct<br>gccggctctgcggcctcttccgcgtcttcgccttcgccctcagacgagtcggatctccctttgggccgcctccccgcatcgataccgt<br>cgacctcgagggaattaattcgagctcggtacctttaagaccaatgacttacaaggcagctgtagatcttagccactttttaaaag<br>aaaaggggggactggaagggctaattcactcccaacgaagacaagatctgctttttgcttgtactgggtctctctggttagaccag<br>atctgagcctgggagctctctggctaactagggaacccactgcttaagcctcaataaagcttgccttgagtgcttcaagtagtgtgt<br>gcccgtctgttgtgtgactctggtaactagagatccctcagacccttttagtcagtgtggaaaatctctagcagcatctagaattaat<br>tccgtgtatt |
| N4: pHR Tre3G Flank N4 240xCAG 12xMS2 WPRE | ctatagtgtcacctaaatcgtatgtgtatgatacataaggttatgtattaattgtagccgcgttctaacgacaatatgtacaagccta<br>attgtgtagcatctggcttactgaagcagaccctatcatctctctcgtaaactgccgtcagagtcggtttggttggacgaaccttctg<br>agtttctggtaacgccgtcccgcacccggaaatggtcagcgaaccaatcagcagggtcatcgctagccagatcctctacgccgga<br>cgcatcgtggccggcatcaccggcgccacaggtgcggttgctggcgcctatatcgccgacatcaccgatggggaagatcgggctc<br>gccacttcgggctcatgagcgcttgtttcggcgtgggtatggtggcaggccccgtggccgggggactgttgggcgccatctccttgc<br>atgcaccattccttgcggcggcggtgctcaacggcctcaacctactactgggctgcttcctaatgcaggagtcgcataagggagag<br>cgtcgaatggtgcactctcagtacaatctgctctgatgccgcatagttaagccagccccgacacccgccaacacccgctgacgcgc<br>cctgacgggcttgtctgctcccggcatccgcttacagacaagctgtgaccgtctccgggagctgcatgtgtcagaggttttcaccgt |

| | |
|---|---|
| | catcaccgaaacgcgcgagacgaaagggcctcgtgatacgcctatttttataggttaatgtcatgataataatggtttcttagacgt<br>caggtggcacttttcggggaaatgtgcgcggaacccctatttgtttatttttctaaatacattcaaatatgtatccgctcatgagaca<br>ataaccctgataaatgcttcaataatattgaaaaaggaagagtatgagtattcaacatttccgtgtcgcccttattcccttttttgcg<br>gcattttgccttcctgtttttgctcacccagaaacgctggtgaaagtaaaagatgctgaagatcagttgggtgcacgagtgggttac<br>atcgaactggatctcaacagcggtaagatccttgagagttttcgccccgaagaacgttttccaatgatgagcacttttaaagttctg<br>ctatgtggcgcggtattatcccgtattgacgccgggcaagagcaactcggtcgccgcatacactattctcagaatgacttggttga<br>gtactcaccagtcacagaaaagcatcttacggatggcatgacagtaagagaattatgcagtgctgccataaccatgagtgataac<br>actgcggccaacttacttctgacaacgatcggaggaccgaaggagctaaccgcttttttgcacaacatgggggatcatgtaactcg<br>ccttgatcgttgggaaccggagctgaatgaagccataccaaacgacgagcgtgacaccacgatgcctgtagcaatggcaacaac<br>gttgcgcaaactattaactggcgaactacttactctagcttcccggcaacaattaatagactggatggaggcggataaagttgcag<br>gaccacttctgcgctcggcccttccggctggctggtttattgctgataaatctggagccggtgagcgtgggtctcgcggtatcattgc<br>agcactggggccagatggtaagccctcccgtatcgtagttatctacacgacggggagtcaggcaactatggatgaacgaaatag<br>acagatcgctgagataggtgcctcactgattaagcattggtaactgtcagaccaagtttactcatatatactttagattgatttaaa<br>acttcatttttaatttaaaaggatctaggtgaagatcctttttgataatctcatgaccaaaatcccttaacgtgagttttcgttccact<br>gagcgtcagaccccgtagaaaagatcaaaggatcttcttgagatcctttttttctgcgcgtaatctgctgcttgcaaacaaaaaaa<br>ccaccgctaccagcggtggtttgtttgccggatcaagagctaccaactctttttccgaaggtaactggcttcagcagagcgcagat<br>accaaatactgtctttctagtgtagccgtagttaggccaccacttcaagaactctgtagcaccgcctacatacctcgctctgctaatc<br>ctgttaccagtggctgctgccagtggcgataagtcgtgtcttaccgggttggactcaagacgatagttaccggataaggcgcagcg<br>gtcgggctgaacggggggttcgtgcacacagcccagcttggagcgaacgacctacaccgaactgagatacctacagcgtgagct<br>atgagaaagcgccacgcttcccgaagggagaaaggcggacaggtatccggtaagcggcagggtcggaacaggagagcgcacg<br>agggagcttccagggggaaacgcctggtatctttatagtcctgtcgggtttcgccacctctgacttgagcgtcgatttttgtgatgct<br>cgtcaggggggcggagcctatggaaaaacgccagcaacgcggcctttttacggttcctggccttttgctggccttttgctcacatgt<br>tctttcctgcgttatcccctgattctgtggataaccgtattaccgcctttgagtgagctgataccgctcgccgcagccgaacgaccga<br>gcgcagcgagtcagtgagcgaggaagcggaagagcgcccaatacgcaaaccgcctctccccgcgcgttggccgattcattaatg<br>cagctgtggaatgtgtgtcagttagggtgtggaaagtccccaggctccccagcaggcagaagtatgcaaagcatgcatctcaatt<br>agtcagcaaccaggtgtggaaagtccccaggctccccagcaggcagaagtatgcaaagcatgcatctcaattagtcagcaacca<br>tagtcccgcccctaactccgcccatcccgcccctaactccgcccagttccgcccattctccgccccatggctgactaattttttttattt<br>atgcagaggccgaggccgcctcggcctctgagctattccagaagtagtgaggaggcttttttggaggcctaggcttttgcaaaaag<br>cttggacacaagacaggcttgcgagatatgtttgagaataccactttatcccgcgtcagggagaggcagtgcgtaaaaagacgc<br>ggactcatgtgaaatactggtttttagtgcgccagatctctataatctcgcgcaacctattttcccctcgaacactttttaagccgtag<br>ataaacaggctgggacacttcacatgagcgaaaaatacatcgtcacctgggacatgttgcagatccatgcacgtaaactcgcaa<br>gccgactgatgccttctgaacaatggaaaggcattattgccgtaagccgtggcggtctgtaccgggtgcgttactggcgcgtgaac<br>tgggtattcgtcatgtcgataccgtttgtatttccagctacgatcacgacaaccagcgcgagcttaaagtgctgaaacgcgcagaa<br>ggcgatggcgaaggcttcatcgttattgatgacctggtggataccggtggtactgcggttgcgattcgtgaaatgtatccaaaagc<br>gcactttgtcaccatcttcgcaaaaccggctggtcgtccgctggttgatgactatgttgttgatatcccgcaagatacctggattgaa<br>cagccgtgggatatgggcgtcgtattcgtcccgccaatctccggtcgctaatcttttcaacgcctggcactgccgggcgttgttctttt<br>taacttcaggcgggttacaatagtttccagtaagtattctggaggctgcatccatgacacaggcaaacctgagcgaaaccctgttc<br>aaaccccgctttaaacatcctgaaacctcgacgctagtccgccgctttaatcacggcgcacaaccgcctgtgcagtcggcccttga<br>tggtaaaaccatccctcactggtatcgcatgattaaccgtctgatgtggatctggcgcggcattgacccacgcgaaatcctcgacg<br>tccaggcacgtattgtgatgagcgatgccgaacgtaccgacgatgatttatacgatacggtgattggctaccgtggcggcaactgg<br>atttatgagtgggccccggatctttgtgaaggaaccttacttctgtggtgtgacataattggacaaactacctacagagatttaaag<br>ctctaaggtaaatataaaatttttaagtgtataatgtgttaaactactgattctaattgtttgtgtattttagattccaacctatggaa<br>ctgatgaatgggagcagtggtggaatgcctttaatgaggaaaacctgttttgctcagaagaaatgccatctagtgatgatgaggct<br>actgctgactctcaacattctactcctccaaaaaagaagagaaaggtagaagaccccaaggactttccttcagaattgctaagttt<br>tttgagtcatgctgtgtttagtaatagaactcttgcttgctttgctatttacaccacaaaggaaaaagctgcactgctatacaagaa<br>aattatggaaaaatattctgtaacctttataagtaggcataacagttataatcataacatactgttttttcttactccacacaggcat<br>agagtgtctgctattaataactatgctcaaaaattgtgtacctttagctttttaatttgtaaaggggttaataaggaatatttgatgta<br>tagtgccttgactagagatcataatcagccataccacatttgtagaggttttacttgctttaaaaaacctcccacacctccccctga |

| | |
|---|---|
| | acctgaaacataaaatgaatgcaattgttgttgttaacttgtttattgcagcttataatggttacaaataaagcaatagcatcacaa<br>atttcacaaataaagcatttttttcactgcattctagttgtggtttgtccaaactcatcaatgtatcttatcatgtctggatcaactgga<br>taactcaagctaaccaaaatcatcccaaacttcccaccccataccctattaccactgccaattacctagtggtttcatttactctaaa<br>cctgtgattcctctgaattattttcattttaaagaaattgtatttgttaaatatgtactacaaacttagtagttggaagggctaattca<br>ctcccaaagaagacaagatatccttgatctgtggatctaccacacacaaggctacttccctgattagcagaactacacaccaggg<br>ccaggggtcagatatccactgacctttggatggtgctacaagctagtaccagttgagccagataaggtagaagaggccaataaa<br>ggagagaacaccagcttgttacaccctgtgagcctgcatgggatggatgacccggagagagaagtgttagagtggaggtttgac<br>agccgcctagcatttcatcacgtggcccgagagctgcatccggagtacttcaagaactgctgatatcgagcttgctacaagggact<br>ttccgctggggactttccagggaggcgtggcctgggcgggactggggagtggcgagccctcagatcctgcatataagcagctgct<br>ttttgcctgtactgggtctctctggttagaccagatctgagcctgggagctctctggctaactagggaacccactgcttaagcctcaa<br>taaagcttgccttgagtgcttcaagtagtgtgtgcccgtctgttgtgtgactctggtaactagagatccctcagacccttttagtcagt<br>gtggaaaatctctagcagtggcgcccgaacagggacttgaaagcgaaagggaaaccagaggagctctctcgacgcaggactcg<br>gcttgctgaagcgcgcacggcaagaggcgaggggcggcgactggtgagtacgccaaaaattttgactagcggaggctagaagg<br>agagagatgggtgcgagagcgtcagtattaagcgggggagaattagatcgcgatgggaaaaaattcggttaaggccagggggа<br>aagaaaaaatataaattaaaacatatagtatgggcaagcagggagctagaacgattcgcagttaatcctggcctgttagaaaca<br>tcagaaggctgtagacaaatactgggacagctacaaccatcccttcagacaggatcagaagaacttagatcattatataatacag<br>tagcaaccctctattgtgtgcatcaaaggatagagataaaagacaccaaggaagctttagacaagatagaggaagagcaaaac<br>aaaagtaagaccaccgcacagcaagcggccggccgctgatcttcagacctggaggaggagatatgagggacaattggagaagt<br>gaattatataaatataaagtagtaaaaattgaaccattaggagtagcacccaccaaggcaaagagaagagtggtgcagagaga<br>aaaaagagcagtgggaataggagctttgttccttgggttcttgggagcagcaggaagcactatgggcgcagcgtcaatgacgctg<br>acggtacaggccagacaattattgtctggtatagtgcagcagcagaacaatttgctgagggctattgaggcgcaacagcatctgtt<br>gcaactcacagtctggggcatcaagcagctccaggcaagaatcctggctgtgaaagatacctaaaggatcaacagctcctgggg<br>atttggggttgctctggaaaactcatttgcaccactgctgtgccttggaatgctagttggagtaataaatctctggaacagatttgg<br>aatcacacgacctggatggagtgggacagagaaattaacaattacacaagcttaatacactccttaattgaagaatcgcaaaac<br>cagcaagaaaagaatgaacaagaattattggaattagataaatgggcaagtttgtggaattggtttaacataacaaattggctgt<br>ggtatataaaattattcataatgatagtaggaggcttggtaggtttaagaatagtttttgctgtactttctatagtgaatagagttag<br>gcagggatattcaccattatcgtttcagacccacctcccaaccccgaggggacccgacaggcccgaaggaatagaagaagaag<br>gtggagagagagacagagacagatccattcgattagtgaacggatctcgacggtatcgccaaatggcagtattcatccacaattt<br>taaaagaaaaggggggattggggggtacagtgcaggggaaagaatagtagacataatagcaacagacatacaaactaaagaa<br>ttacaaaaacaaattacaaaaattcaaaattttcgggtttattacagggacagcagagatccagtttatcgatgaggccctttcgt<br>cttcactcgagtttactccctatcagtgatagagaacgtatgaagagtttactccctatcagtgatagagaacgtatgcagacttta<br>ctccctatcagtgatagagaacgtataaggagtttactccctatcagtgatagagaacgtatgaccagtttactccctatcagtgat<br>agagaacgtatctacagtttactccctatcagtgatagagaacgtatatccagtttactccctatcagtgatagagaacgtatgtcg<br>aggtaggcgtgtacggtgggcgcctataaaagcagagctcgtttagtgaaccgtcagatcgcctggagcaattccacaacactttt<br>gtcttatacttACGCGTGTCGACcatatgAGTCCACTATTAAAGAACGTGGACTCCAACGTCAAAGGGCG<br>AAAAACCGTCTATCAGGGCGATGGCCCACTACGTGAACCATCACCCTAATCAAGTTTTTTGGGGTC<br>GAGGTGCCGTAAAGCACTAAATCGGAACCCTAAAGGGAGCCCCCGATTTAGAGCTTGACGGGGA<br>AAGCCGGCGAACGTGGCGAGAAAGGAAGGGAAGAAAGCGAAAGGAGCGGGCGCTAGGGCGCT<br>GGCAAGTGTAGCGGTCACTatgGGTAATTATAACCCGGGCCCTATATATTAGGCTAACTAGCTAACT<br>AGGAATTCggtctcaCAGCAGCAGCAGCAGCAGCAGCAGCAGCAGCAGCAGCAGCACAGCAGC<br>AGCAGCAGCAGCAGCAGCAGCAGCAGCAGCAGCAGCAGCAGCAGCAGCAGCAGCAGCAGC<br>AGCAGCAGCAGCAGCAGCAGCAGCACAGCAGCAGCAGCAGCAGCAGCAGCAGCAGCAGCA<br>GCAGCAGCAGCAGCAGCAGCAGCAGCAGCAGCAGCAGCAGCAGCAGCAGCAGCACAGCAG<br>CAGCAGCAGCAGCAGCAGCAGCAGCAGCAGCAGCAGCAGCAGCAGCAGCAGCAGCAGCAG<br>CAGCAGCAGCAGCAGCAGCAGCACAGCAGCAGCAGCAGCAGCAGCAGCAGCAGCAGCAGC<br>AGCAGCAGCAGCAGCAGCAGCAGCAGCAGCAGCAGCAGCAGCAGCAGCAGCACAGCAGCA<br>GCAGCAGCAGCAGCAGCAGCAGCAGCAGCAGCAGCAGCAGCAGCAGCAGCAGCAGCAGCA<br>GCAGCAGCAGCAGCAGCAGCACAGCAGCAGCAGCAGCAGCAGCAGCAGCAGCAGCAG |

| | |
|---|---|
| | CAGCAGCAGCAGCAGCAGCAGCAGCAGCAGCAGCACAGCAGCAGCAGCAGCAGCAGCAGCAGC<br>AGCAGCAGCAGCAGCAGCAGCAGCAGCAGCAGCAGCAGCAGCAGCAGCAGCAGCAGCAGC<br>AGCACAGCAGCAGCAGCAGCAGCAGCAGCAGCAGCAGCAGCAGCAGCAgagacgGAGCTCgtcgac<br>GCggccgcaagcttggtaccgagctcGGATCCTAAGGTACCTAATTGCCTAGAAAACATGAGGATCACCCA<br>TGTCTGCAGGTCGACTCTAGAAAACATGAGGATCACCCATGTCTGCAGTATTCCCGGGTTCATTAG<br>ATCCTAAGGTACCTAATTGCCTAGAAAACATGAGGATCACCCATGTCTGCAGGTCGACTCTAGAAA<br>ACATGAGGATCACCCATGTCTGCAGTATTCCCGGGTTCATTAGATCCTAAGGTACCTAATTGCCTAG<br>AAAACATGAGGATCACCCATGTCTGCAGGTCGACTCTAGAAAACATGAGGATCACCCATGTCTGCA<br>GTATTCCCGGGTTCATTAGATCCTAAGGTACCTAATTGCCTAGAAAACATGAGGATCACCCATGTCT<br>GCAGGTCGACTCTAGAAAACATGAGGATCACCCATGTCTGCAGTATTCCCGGGTTCATTAGATCCT<br>AAGGTACCTAATTGCCTAGAAAACATGAGGATCACCCATGTCTGCAGGTCGACTCTAGAAAACATG<br>AGGATCACCCATGTCTGCAGTATTCCCGGGTTCATTAGATCCTAAGGTACCTAATTGCCTAGAAAAC<br>ATGAGGATCACCCATGTCTGCAGGTCGACTCTAGAAAACATGAGGATCACCCATGTCTGCAGTATT<br>CCCGGGTTCATTAGATCTGCGCGCGATCGATATCAGCGCTTTAAATTTGCGCATGCTAGTgTAAggc<br>atgcaagcttgatatcaagcttatcgataatcaacctctggattacaaaatttgtgaaagattgactggtattcttaactatgttgct<br>ccttttacgctatgtggatacgctgctttaatgcctttgtatcatgctattgcttcccgtatggctttcattttctcctccttgtataaatc<br>ctggttgctgtctctttatgaggagttgtggcccgttgtcaggcaacgtggcgtggtgtgcactgtgtttgctgacgcaacccccact<br>ggttggggcattgccaccacctgtcagctcctttccgggactttcgctttccccctccctattgccacggcggaactcatcgccgcct<br>gccttgcccgctgctggacaggggctcggctgttgggcactgacaattccgtggtgttgtcggggaaatcatcgtcctttccttggct<br>gctcgcctgtgttgccacctggattctgcgcgggacgtccttctgctacgtcccttcggccctcaatccagcggaccttccttcccgc<br>ggcctgctgccggctctgcggcctcttccgcgtcttcgccttcgccctcagacgagtcggatctccctttgggccgcctccccgcatc<br>gataccgtcgacctcgagggaattaattcgagctcggtacctttaagaccaatgacttacaaggcagctgtagatcttagccacttt<br>ttaaaagaaaaggggggactggaagggctaattcactcccaacgaagacaagatctgctttttgcttgtactgggtctctctggtt<br>agaccagatctgagcctgggagctctctggctaactagggaacccactgcttaagcctcaataaagcttgccttgagtgcttcaag<br>tagtgtgtgcccgtctgttgtgtgactctggtaactagagatccctcagacccttttagtcagtgtggaaaatctctagcagcatcta<br>gaattaattccgtgtatt |
| N5: pHR Tre3G Flank N5 240xCAG 12xMS2 WPRE | ctatagtgtcacctaaatcgtatgtgtatgatacataaggttatgtattaattgtagccgcgttctaacgacaatatgtacaagccta<br>attgtgtagcatctggcttactgaagcagaccctatcatctctctcgtaaactgccgtcagagtcggtttggttggacgaaccttctg<br>agtttctggtaacgccgtcccgcacccggaaatggtcagcgaaccaatcagcagggtcatcgctagccagatcctctacgccgga<br>cgcatcgtggccggcatcaccggcgccacaggtgcggttgctggcgcctatatcgccgacatcaccgatggggaagatcgggctc<br>gccacttcgggctcatgagcgcttgtttcggcgtgggtatggtggcaggccccgtggccgggggactgttgggcgccatctccttgc<br>atgcaccattccttgcggcggcggtgctcaacggcctcaacctactactgggctgcttcctaatgcaggagtcgcataagggagag<br>cgtcgaatggtgcactctcagtacaatctgctctgatgccgcatagttaagccagccccgacacccgccaacacccgctgacgcgc<br>cctgacgggcttgtctgctcccggcatccgcttacagacaagctgtgaccgtctccgggagctgcatgtgtcagaggttttcaccgt<br>catcaccgaaacgcgcgagacgaaagggcctcgtgatacgcctatttttataggttaatgtcatgataataatggtttcttagacgt<br>caggtggcacttttcggggaaatgtgcgcggaacccctatttgtttatttttctaaatacattcaaatatgtatccgctcatgagaca<br>ataaccctgataaatgcttcaataatattgaaaaaggaagagtatgagtattcaacatttccgtgtcgcccttattcccttttttgcg<br>gcattttgccttcctgtttttgctcacccagaaacgctggtgaaagtaaaagatgctgaagatcagttgggtgcacgagtgggttac<br>atcgaactggatctcaacagcggtaagatccttgagagttttcgccccgaagaacgttttccaatgatgagcacttttaaagttctg<br>ctatgtggcgcggtattatcccgtattgacgccgggcaagagcaactcggtcgccgcatacactattctcagaatgacttggttga<br>gtactcaccagtcacagaaaagcatcttacggatggcatgacagtaagagaattatgcagtgctgccataaccatgagtgataac<br>actgcggccaacttacttctgacaacgatcggaggaccgaaggagctaaccgcttttttgcacaacatgggggatcatgtaactcg<br>ccttgatcgttgggaaccggagctgaatgaagccataccaaacgacgagcgtgacaccacgatgcctgtagcaatggcaacaac<br>gttgcgcaaactattaactggcgaactacttactctagcttcccggcaacaattaatagactggatggaggcggataaagttgcag<br>gaccacttctgcgctcggcccttccggctggctggtttattgctgataaatctggagccggtgagcgtgggtctcgcggtatcattgc<br>agcactggggccagatggtaagccctcccgtatcgtagttatctacacgacggggagtcaggcaactatggatgaacgaaatag<br>acagatcgctgagataggtgcctcactgattaagcattggtaactgtcagaccaagtttactcatatatactttagattgatttaa |

| | |
|---|---|
| | acttcatttttaatttaaaaggatctaggtgaagatcctttttgataatctcatgaccaaaatcccttaacgtgagttttcgttccact<br>gagcgtcagaccccgtagaaaagatcaaaggatcttcttgagatcctttttttctgcgcgtaatctgctgcttgcaaacaaaaaaa<br>ccaccgctaccagcggtggtttgtttgccggatcaagagctaccaactctttttccgaaggtaactggcttcagcagagcgcagat<br>accaaatactgtctttctagtgtagccgtagttaggccaccacttcaagaactctgtagcaccgcctacatacctcgctctgctaatc<br>ctgttaccagtggctgctgccagtggcgataagtcgtgtcttaccgggttggactcaagacgatagttaccggataaggcgcagcg<br>gtcgggctgaacggggggttcgtgcacacagcccagcttggagcgaacgacctacaccgaactgagatacctacagcgtgagct<br>atgagaaagcgccacgcttcccgaagggagaaaggcggacaggtatccggtaagcggcagggtcggaacaggagagcgcacg<br>agggagcttccagggggaaacgcctggtatctttatagtcctgtcgggtttcgccacctctgacttgagcgtcgatttttgtgatgct<br>cgtcaggggggcggagcctatggaaaaacgccagcaacgcggcctttttacggttcctggccttttgctggccttttgctcacatgt<br>tctttcctgcgttatcccctgattctgtggataaccgtattaccgcctttgagtgagctgataccgctcgccgcagccgaacgaccga<br>gcgcagcgagtcagtgagcgaggaagcggaagagcgcccaatacgcaaaccgcctctccccgcgcgttggccgattcattaatg<br>cagctgtggaatgtgtgtcagttagggtgtggaaagtccccaggctccccagcaggcagaagtatgcaaagcatgcatctcaatt<br>agtcagcaaccaggtgtggaaagtccccaggctccccagcaggcagaagtatgcaaagcatgcatctcaattagtcagcaacca<br>tagtcccgcccctaactccgcccatcccgcccctaactccgcccagttccgcccattctccgccccatggctgactaattttttttattt<br>atgcagaggccgaggccgcctcggcctctgagctattccagaagtagtgaggaggcttttttggaggcctaggcttttgcaaaaag<br>cttggacacaagacaggcttgcgagatatgtttgagaataccactttatcccgcgtcagggagaggcagtgcgtaaaaagacgc<br>ggactcatgtgaaatactggtttttagtgcgccagatctctataatctcgcgcaacctattttcccctcgaacactttttaagccgtag<br>ataaacaggctgggacacttcacatgagcgaaaaatacatcgtcacctgggacatgttgcagatccatgcacgtaaactcgcaa<br>gccgactgatgccttctgaacaatggaaaggcattattgccgtaagccgtggcggtctgtaccgggtgcgttactggcgcgtgaac<br>tgggtattcgtcatgtcgataccgtttgtatttccagctacgatcacgacaaccagcgcgagcttaaagtgctgaaacgcgcagaa<br>ggcgatggcgaaggcttcatcgttattgatgacctggtggataccggtggtactgcggttgcgattcgtgaaatgtatccaaaagc<br>gcactttgtcaccatcttcgcaaaaccggctggtcgtccgctggttgatgactatgttgttgatatcccgcaagatacctggattgaa<br>cagccgtgggatatgggcgtcgtattcgtcccgccaatctccggtcgctaatcttttcaacgcctggcactgccgggcgttgttctttt<br>taacttcaggcgggttacaatagtttccagtaagtattctggaggctgcatccatgacacaggcaaacctgagcgaaaccctgttc<br>aaaccccgctttaaacatcctgaaacctcgacgctagtccgccgctttaatcacggcgcacaaccgcctgtgcagtcggcccttga<br>tggtaaaaccatccctcactggtatcgcatgattaaccgtctgatgtggatctggcgcggcattgacccacgcgaaatcctcgacg<br>tccaggcacgtattgtgatgagcgatgccgaacgtaccgacgatgatttatacgatacggtgattggctaccgtggcggcaactgg<br>atttatgagtgggccccggatctttgtgaaggaaccttacttctgtggtgtgacataattggacaaactacctacagagatttaaag<br>ctctaaggtaaatataaaatttttaagtgtataatgtgttaaactactgattctaattgtttgtgtattttagattccaacctatggaa<br>ctgatgaatgggagcagtggtggaatgcctttaatgaggaaaacctgttttgctcagaagaaatgccatctagtgatgatgaggct<br>actgctgactctcaacattctactcctccaaaaaagaagagaaaggtagaagaccccaaggactttccttcagaattgctaagttt<br>tttgagtcatgctgtgtttagtaatagaactcttgcttgctttgctatttacaccacaaaggaaaaagctgcactgctatacaagaa<br>aattatggaaaaatattctgtaacctttataagtaggcataacagttataatcataacatactgttttttcttactccacacaggcat<br>agagtgtctgctattaataactatgctcaaaaattgtgtacctttagctttttaatttgtaaaggggttaataaggaatatttgatgta<br>tagtgccttgactagagatcataatcagccataccacatttgtagaggttttacttgctttaaaaaacctcccacacctccccctga<br>acctgaaacataaaatgaatgcaattgttgttgttaacttgtttattgcagcttataatggttacaaataaagcaatagcatcacaa<br>atttcacaaataaagcatttttttcactgcattctagttgtggtttgtccaaactcatcaatgtatcttatcatgtctggatcaactgga<br>taactcaagctaaccaaaatcatcccaaacttcccaccccataccctattaccactgccaattacctagtggtttcatttactctaaa<br>cctgtgattcctctgaattattttcattttaaagaaattgtatttgttaaatatgtactacaaacttagtagttggaagggctaattca<br>ctcccaaagaagacaagatatccttgatctgtggatctaccacacacaaggctacttccctgattagcagaactacacaccaggg<br>ccaggggtcagatatccactgacctttggatggtgctacaagctagtaccagttgagccagataaggtagaagaggccaataaa<br>ggagagaacaccagcttgttacaccctgtgagcctgcatgggatggatgacccggagagagaagtgttagagtggaggtttgac<br>agccgcctagcatttcatcacgtggcccgagagctgcatccggagtacttcaagaactgctgatatcgagcttgctacaagggact<br>ttccgctggggactttccagggaggcgtggcctgggcgggactggggagtggcgagccctcagatcctgcatataagcagctgct<br>ttttgcctgtactgggtctctctggttagaccagatctgagcctgggagctctctggctaactagggaacccactgcttaagcctcaa<br>taaagcttgccttgagtgcttcaagtagtgtgtgcccgtctgttgtgtgactctggtaactagagatccctcagacccttttagtcagt<br>gtggaaaatctctagcagtggcgcccgaacagggacttgaaagcgaaagggaaaccagaggagctctctcgacgcaggactcg<br>gcttgctgaagcgcgcacggcaagaggcgaggggcggcgactggtgagtacgccaaaaattttgactagcggaggctagaagg |

| | |
|---|---|
| | agagagatgggtgcgagagcgtcagtattaagcgggggagaattagatcgcgatgggaaaaaattcggttaaggccaggggga<br>aagaaaaaatataaattaaaacatatagtatgggcaagcagggagctagaacgattcgcagttaatcctggcctgttagaaaca<br>tcagaaggctgtagacaaatactgggacagctacaaccatcccttcagacaggatcagaagaacttagatcattatataatacag<br>tagcaaccctctattgtgtgcatcaaaggatagagataaaagacaccaaggaagctttagacaagatagaggaagagcaaaac<br>aaaagtaagaccaccgcacagcaagcggccggccgctgatcttcagacctggaggaggagatatgagggacaattggagaagt<br>gaattatataaatataaagtagtaaaaattgaaccattaggagtagcacccaccaaggcaaagagaagagtggtgcagagaga<br>aaaaagagcagtgggaataggagctttgttccttgggttcttgggagcagcaggaagcactatgggcgcagcgtcaatgacgctg<br>acggtacaggccagacaattattgtctggtatagtgcagcagcagaacaatttgctgagggctattgaggcgcaacagcatctgtt<br>gcaactcacagtctggggcatcaagcagctccaggcaagaatcctggctgtgaaagatacctaaaggatcaacagctcctgggg<br>atttggggttgctctggaaaactcatttgcaccactgctgtgccttggaatgctagttggagtaataaatctctggaacagatttgg<br>aatcacacgacctggatggagtgggacagagaaattaacaattacacaagcttaatacactccttaattgaagaatcgcaaaac<br>cagcaagaaaagaatgaacaagaattattggaattagataaatgggcaagtttgtggaattggtttaacataacaaattggctgt<br>ggtatataaaattattcataatgatagtaggaggcttggtaggtttaagaatagtttttgctgtactttctatagtgaatagagttag<br>gcagggatattcaccattatcgtttcagacccacctcccaaccccgaggggacccgacaggcccgaaggaatagaagaagaag<br>gtggagagagagacagagacagatccattcgattagtgaacggatctcgacggtatcgccaaatggcagtattcatccacaattt<br>taaaagaaaagggggggattggggggtacagtgcaggggaaagaatagtagacataatagcaacagacatacaaactaaagaa<br>ttacaaaaacaaattacaaaaattcaaaattttcgggtttattacagggacagcagagatccagtttatcgatgaggccctttcgt<br>cttcactcgagtttactccctatcagtgatagagaacgtatgaagagtttactccctatcagtgatagagaacgtatgcagacttta<br>ctccctatcagtgatagagaacgtataaggagtttactccctatcagtgatagagaacgtatgaccagtttactccctatcagtgat<br>agagaacgtatctacagtttactccctatcagtgatagagaacgtatatccagtttactccctatcagtgatagagaacgtatgtcg<br>aggtaggcgtgtacggtgggcgcctataaaagcagagctcgtttagtgaaccgtcagatcgcctggagcaattccacaacactttt<br>gtcttatacttACGCGTGTCGACcatatgACCGCCACCGCCGCCGCCGCCGCCGCCTCCTCAGCTTCC<br>TCAGCCGCCGCCGCAGGCACAGCCGCTGCTGCCTCAGCCGCAGCCGCCCCCGCCGCCACCCGGCC<br>CGGCTGTGGCTGAGGAGCCGCTGCACCGACCGTGAGTTTGGGCCCGCTGCAGCTCCCTGTCCCGG<br>CGGGTCCCAGGCTACGGCGGGGATGGCGGTAACCCTGCAGCCTGCGGCCGGCGACACGAACCC<br>CCGTatgGGTAATTATAACCCGGGCCCTATATATTAGGCTAACTAGCTAACTAGGAATTCggtctcaCA<br>GCAGCAGCAGCAGCAGCAGCAGCAGCAGCAGCAGCAGCAGCACAGCAGCAGCAGCAGCAGCAG<br>CAGCAGCAGCAGCAGCAGCAGCAGCAGCAGCAGCAGCAGCAGCAGCAGCAGCAGCAGCAG<br>CAGCAGCAGCACAGCAGCAGCAGCAGCAGCAGCAGCAGCAGCAGCAGCAGCAGCAGCAGC<br>AGCAGCAGCAGCAGCAGCAGCAGCAGCAGCAGCAGCAGCAGCACAGCAGCAGCAGCAGCA<br>GCAGCAGCAGCAGCAGCAGCAGCAGCAGCAGCAGCAGCAGCAGCAGCAGCAGCAGCAGCA<br>GCAGCAGCACAGCAGCAGCAGCAGCAGCAGCAGCAGCAGCAGCAGCAGCAGCAGCAGCAG<br>CAGCAGCAGCAGCAGCAGCAGCAGCAGCAGCAGCAGCAGCACAGCAGCAGCAGCAGCAGC<br>AGCAGCAGCAGCAGCAGCAGCAGCAGCAGCAGCAGCAGCAGCAGCAGCAGCAGCAGCAGC<br>AGCAGCAGCACAGCAGCAGCAGCAGCAGCAGCAGCAGCAGCAGCAGCAGCAGCAGCAGCA<br>GCAGCAGCAGCAGCAGCAGCACAGCAGCAGCAGCAGCAGCAGCAGCAGCAGCAGCAGCAG<br>CAGCAGCAGCAGCAGCAGCAGCAGCAGCAGCAGCAGCAGCAGCAGCAGCACAGCAGCAGC<br>AGCAGCAGCAGCAGCAGCAGCAGCAGCAGCAGCAgagacgGAGCTCgtcgacGCggccgcaagcttggta<br>ccgagctcGGATCCTAAGGTACCTAATTGCCTAGAAAACATGAGGATCACCCATGTCTGCAGGTCGAC<br>TCTAGAAAACATGAGGATCACCCATGTCTGCAGTATTCCCGGGTTCATTAGATCCTAAGGTACCTA<br>ATTGCCTAGAAAACATGAGGATCACCCATGTCTGCAGGTCGACTCTAGAAAACATGAGGATCACCC<br>ATGTCTGCAGTATTCCCGGGTTCATTAGATCCTAAGGTACCTAATTGCCTAGAAAACATGAGGATC<br>ACCCATGTCTGCAGGTCGACTCTAGAAAACATGAGGATCACCCATGTCTGCAGTATTCCCGGGTTC<br>ATTAGATCCTAAGGTACCTAATTGCCTAGAAAACATGAGGATCACCCATGTCTGCAGGTCGACTCT<br>AGAAAACATGAGGATCACCCATGTCTGCAGTATTCCCGGGTTCATTAGATCCTAAGGTACCTAATT<br>GCCTAGAAAACATGAGGATCACCCATGTCTGCAGGTCGACTCTAGAAAACATGAGGATCACCCAT<br>GTCTGCAGTATTCCCGGGTTCATTAGATCCTAAGGTACCTAATTGCCTAGAAAACATGAGGATCAC<br>CCATGTCTGCAGGTCGACTCTAGAAAACATGAGGATCACCCATGTCTGCAGTATTCCCGGGTTCAT |

| | |
|---|---|
| | TAGATCTGCGCGCGATCGATATCAGCGCTTTAAATTTGCGCATGCTAGTgTAAggcatgcaagcttgatat<br>caagcttatcgataatcaacctctggattacaaaatttgtgaaagattgactggtattcttaactatgttgctccttttacgctatgtg<br>gatacgctgctttaatgcctttgtatcatgctattgcttcccgtatggctttcattttctcctccttgtataaatcctggttgctgtctctt<br>tatgaggagttgtggcccgttgtcaggcaacgtggcgtggtgtgcactgtgtttgctgacgcaacccccactggttggggcattgcc<br>accacctgtcagctcctttccgggactttcgctttccccctccctattgccacggcggaactcatcgccgcctgccttgcccgctgctg<br>gacaggggctcggctgttgggcactgacaattccgtggtgttgtcggggaaatcatcgtcctttccttggctgctcgcctgtgttgcc<br>acctggattctgcgcgggacgtccttctgctacgtcccttcggccctcaatccagcggaccttccttcccgcggcctgctgccggctc<br>tgcggcctcttccgcgtcttcgccttcgccctcagacgagtcggatctccctttgggccgcctccccgcatcgataccgtcgacctcg<br>agggaattaattcgagctcggtacctttaagaccaatgacttacaaggcagctgtagatcttagccactttttaaaagaaaagggg<br>ggactggaagggctaattcactcccaacgaagacaagatctgctttttgcttgtactgggtctctctggttagaccagatctgagcc<br>tgggagctctctggctaactagggaacccactgcttaagcctcaataaagcttgccttgagtgcttcaagtagtgtgtgcccgtctg<br>ttgtgtgactctggtaactagagatccctcagacccttttagtcagtgtggaaaatctctagcagcatctagaattaattccgtgtat<br>t |
| N6: pHR Tre3G Flank N6 240xCAG 12xMS2 WPRE | ctatagtgtcacctaaatcgtatgtgtatgatacataaggttatgtattaattgtagccgcgttctaacgacaatatgtacaagccta<br>attgtgtagcatctggcttactgaagcagaccctatcatctctctcgtaaactgccgtcagagtcggtttggttggacgaaccttctg<br>agtttctggtaacgccgtcccgcacccggaaatggtcagcgaaccaatcagcagggtcatcgctagccagatcctctacgccgga<br>cgcatcgtggccggcatcaccggcgccacaggtgcggttgctggcgcctatatcgccgacatcaccgatggggaagatcgggctc<br>gccacttcgggctcatgagcgcttgtttcggcgtgggtatggtggcaggccccgtggccgggggactgttgggcgccatctccttgc<br>atgcaccattccttgcggcggcggtgctcaacggcctcaacctactactgggctgcttcctaatgcaggagtcgcataagggagag<br>cgtcgaatggtgcactctcagtacaatctgctctgatgccgcatagttaagccagccccgacacccgccaacacccgctgacgcgc<br>cctgacgggcttgtctgctcccggcatccgcttacagacaagctgtgaccgtctccgggagctgcatgtgtcagaggttttcaccgt<br>catcaccgaaacgcgcgagacgaaagggcctcgtgatacgcctatttttataggttaatgtcatgataataatggtttcttagacgt<br>caggtggcacttttcggggaaatgtgcgcggaacccctatttgtttatttttctaaatacattcaaatatgtatccgctcatgagaca<br>ataaccctgataaatgcttcaataatattgaaaaaggaagagtatgagtattcaacatttccgtgtcgcccttattcccttttttgcg<br>gcattttgccttcctgtttttgctcacccagaaacgctggtgaaagtaaaagatgctgaagatcagttgggtgcacgagtgggttac<br>atcgaactggatctcaacagcggtaagatccttgagagttttcgccccgaagaacgttttccaatgatgagcacttttaaagttctg<br>ctatgtggcgcggtattatcccgtattgacgccgggcaagagcaactcggtcgccgcatacactattctcagaatgacttggttga<br>gtactcaccagtcacagaaaagcatcttacggatggcatgacagtaagagaattatgcagtgctgccataaccatgagtgataac<br>actgcggccaacttacttctgacaacgatcggaggaccgaaggagctaaccgcttttttgcacaacatgggggatcatgtaactcg<br>ccttgatcgttgggaaccggagctgaatgaagccataccaaacgacgagcgtgacaccacgatgcctgtagcaatggcaacaac<br>gttgcgcaaactattaactggcgaactacttactctagcttcccggcaacaattaatagactggatggaggcggataaagttgcag<br>gaccacttctgcgctcggcccttccggctggctggtttattgctgataaatctggagccggtgagcgtgggtctcgcggtatcattgc<br>agcactggggccagatggtaagccctcccgtatcgtagttatctacacgacggggagtcaggcaactatggatgaacgaaatag<br>acagatcgctgagataggtgcctcactgattaagcattggtaactgtcagaccaagtttactcatatatactttagattgatttaaa<br>acttcatttttaatttaaaaggatctaggtgaagatcctttttgataatctcatgaccaaaatcccttaacgtgagttttcgttccact<br>gagcgtcagaccccgtagaaaagatcaaaggatcttcttgagatcctttttttctgcgcgtaatctgctgcttgcaaacaaaaaaa<br>ccaccgctaccagcggtggtttgtttgccggatcaagagctaccaactctttttccgaaggtaactggcttcagcagagcgcagat<br>accaaatactgtctttctagtgtagccgtagttaggccaccacttcaagaactctgtagcaccgcctacatacctcgctctgctaatc<br>ctgttaccagtggctgctgccagtggcgataagtcgtgtcttaccgggttggactcaagacgatagttaccggataaggcgcagcg<br>gtcgggctgaacggggggttcgtgcacacagcccagcttggagcgaacgacctacaccgaactgagatacctacagcgtgagct<br>atgagaaagcgccacgcttcccgaagggagaaaggcggacaggtatccggtaagcggcagggtcggaacaggagagcgcacg<br>agggagcttccagggggaaacgcctggtatctttatagtcctgtcgggtttcgccacctctgacttgagcgtcgatttttgtgatgct<br>cgtcaggggggcggagcctatggaaaaacgccagcaacgcggcctttttacggttcctggccttttgctggccttttgctcacatgt<br>tctttcctgcgttatcccctgattctgtggataaccgtattaccgcctttgagtgagctgataccgctcgccgcagccgaacgaccga<br>gcgcagcgagtcagtgagcgaggaagcggaagagcgcccaatacgcaaaccgcctctccccgcgcgttggccgattcattaatg<br>cagctgtggaatgtgtgtcagttagggtgtggaaagtccccaggctccccagcaggcagaagtatgcaaagcatgcatctcaatt<br>agtcagcaaccaggtgtggaaagtccccaggctccccagcaggcagaagtatgcaaagcatgcatctcaattagtcagcaacca |

| | |
|---|---|
| | tagtcccgcccctaactccgcccatcccgcccctaactccgcccagttccgcccattctccgccccatggctgactaattttttttattt<br>atgcagaggccgaggccgcctcggcctctgagctattccagaagtagtgaggaggcttttttggaggcctaggcttttgcaaaaag<br>cttggacacaagacaggcttgcgagatatgtttgagaataccactttatcccgcgtcagggagaggcagtgcgtaaaaagacgc<br>ggactcatgtgaaatactggtttttagtgcgccagatctctataatctcgcgcaacctattttcccctcgaacactttttaagccgtag<br>ataaacaggctgggacacttcacatgagcgaaaaatacatcgtcacctgggacatgttgcagatccatgcacgtaaactcgcaa<br>gccgactgatgccttctgaacaatggaaaggcattattgccgtaagccgtggcggtctgtaccgggtgcgttactggcgcgtgaac<br>tgggtattcgtcatgtcgataccgtttgtatttccagctacgatcacgacaaccagcgcgagcttaaagtgctgaaacgcgcagaa<br>ggcgatggcgaaggcttcatcgttattgatgacctggtggataccggtggtactgcggttgcgattcgtgaaatgtatccaaaagc<br>gcactttgtcaccatcttcgcaaaaccggctggtcgtccgctggttgatgactatgttgttgatatcccgcaagatacctggattgaa<br>cagccgtgggatatgggcgtcgtattcgtcccgccaatctccggtcgctaatcttttcaacgcctggcactgccgggcgttgttctttt<br>taacttcaggcgggttacaatagtttccagtaagtattctggaggctgcatccatgacacaggcaaacctgagcgaaaccctgttc<br>aaaccccgctttaaacatcctgaaacctcgacgctagtccgccgctttaatcacggcgcacaaccgcctgtgcagtcggcccttga<br>tggtaaaaccatccctcactggtatcgcatgattaaccgtctgatgtggatctggcgcggcattgacccacgcgaaatcctcgacg<br>tccaggcacgtattgtgatgagcgatgccgaacgtaccgacgatgatttatacgatacggtgattggctaccgtggcggcaactgg<br>atttatgagtgggccccggatctttgtgaaggaaccttacttctgtggtgtgacataattggacaaactacctacagagatttaaag<br>ctctaaggtaaatataaaatttttaagtgtataatgtgttaaactactgattctaattgtttgtgtattttagattccaacctatggaa<br>ctgatgaatgggagcagtggtggaatgcctttaatgaggaaaacctgttttgctcagaagaaatgccatctagtgatgatgaggct<br>actgctgactctcaacattctactcctccaaaaaagaagagaaaggtagaagaccccaaggactttccttcagaattgctaagttt<br>tttgagtcatgctgtgtttagtaatagaactcttgcttgctttgctatttacaccacaaaggaaaaagctgcactgctatacaagaa<br>aattatggaaaaatattctgtaacctttataagtaggcataacagttataatcataacatactgttttttcttactccacacaggcat<br>agagtgtctgctattaataactatgctcaaaaattgtgtacctttagctttttaatttgtaaaggggttaataaggaatatttgatgta<br>tagtgccttgactagagatcataatcagccataccacatttgtagaggttttacttgctttaaaaaacctcccacacctccccctga<br>acctgaaacataaaatgaatgcaattgttgttgttaacttgtttattgcagcttataatggttacaaataaagcaatagcatcacaa<br>atttcacaaataaagcatttttttcactgcattctagttgtggtttgtccaaactcatcaatgtatcttatcatgtctggatcaactgga<br>taactcaagctaaccaaaatcatcccaaacttcccaccccataccctattaccactgccaattacctagtggtttcatttactctaaa<br>cctgtgattcctctgaattattttcattttaaagaaattgtatttgttaaatatgtactacaaacttagtagttggaagggctaattca<br>ctcccaaagaagacaagatatccttgatctgtggatctaccacacacaaggctacttccctgattagcagaactacacaccaggg<br>ccaggggtcagatatccactgacctttggatggtgctacaagctagtaccagttgagccagataaggtagaagaggccaataaa<br>ggagagaacaccagcttgttacaccctgtgagcctgcatgggatggatgacccggagagagaagtgttagagtggaggtttgac<br>agccgcctagcatttcatcacgtggcccgagagctgcatccggagtacttcaagaactgctgatatcgagcttgctacaagggact<br>ttccgctggggactttccagggaggcgtggcctgggcgggactggggagtggcgagccctcagatcctgcatataagcagctgct<br>ttttgcctgtactgggtctctctggttagaccagatctgagcctgggagctctctggctaactagggaacccactgcttaagcctcaa<br>taaagcttgccttgagtgcttcaagtagtgtgtgcccgtctgttgtgtgactctggtaactagagatccctcagacccttttagtcagt<br>gtggaaaatctctagcagtggcgcccgaacagggacttgaaagcgaaagggaaaccagaggagctctctcgacgcaggactcg<br>gcttgctgaagcgcgcacggcaagaggcgaggggcggcgactggtgagtacgccaaaaattttgactagcggaggctagaagg<br>agagagatgggtgcgagagcgtcagtattaagcgggggagaattagatcgcgatgggaaaaaattcggttaaggccaggggga<br>aagaaaaaatataaattaaaacatatagtatgggcaagcagggagctagaacgattcgcagttaatcctggcctgttagaaaca<br>tcagaaggctgtagacaaatactgggacagctacaaccatcccttcagacaggatcagaagaacttagatcattatataatacag<br>tagcaaccctctattgtgtgcatcaaaggatagagataaaagacaccaaggaagctttagacaagatagaggaagagcaaaac<br>aaaagtaagaccaccgcacagcaagcggccggccgctgatcttcagacctggaggaggagatatgagggacaattggagaagt<br>gaattatataaatataaagtagtaaaaattgaaccattaggagtagcacccaccaaggcaaagagaagagtggtgcagagaga<br>aaaaagagcagtgggaataggagctttgttccttgggttcttgggagcagcaggaagcactatgggcgcagcgtcaatgacgctg<br>acggtacaggccagacaattattgtctggtatagtgcagcagcagaacaatttgctgagggctattgaggcgcaacagcatctgtt<br>gcaactcacagtctggggcatcaagcagctccaggcaagaatcctggctgtgaaagatacctaaaggatcaacagctcctgggg<br>atttggggttgctctggaaaactcatttgcaccactgctgtgccttggaatgctagttggagtaataaatctctggaacagatttgg<br>aatcacacgacctggatggagtgggacagagaaattaacaattacacaagcttaatacactccttaattgaagaatcgcaaaac<br>cagcaagaaaagaatgaacaagaattattggaattagataaatgggcaagtttgtggaattggtttaacataacaaattggctgt<br>ggtatataaaattattcataatgatagtaggaggcttggtaggtttaagaatagtttttgctgtactttctatagtgaatagagttag |

| | |
|---|---|
| | gcagggatattcaccattatcgtttcagacccacctcccaaccccgaggggacccgacaggcccgaaggaatagaagaagaag<br>gtggagagagagacagagacagatccattcgattagtgaacggatctcgacggtatcgccaaatggcagtattcatccacaattt<br>taaaagaaaaggggggattggggggtacagtgcaggggaaagaatagtagacataatagcaacagacatacaaactaaagaa<br>ttacaaaaacaaattacaaaaattcaaaattttcgggtttattacagggacagcagagatccagtttatcgatgaggccctttcgt<br>cttcactcgagtttactccctatcagtgatagagaacgtatgaagagtttactccctatcagtgatagagaacgtatgcagacttta<br>ctccctatcagtgatagagaacgtataaggagtttactccctatcagtgatagagaacgtatgaccagtttactccctatcagtgat<br>agagaacgtatctacagtttactccctatcagtgatagagaacgtatatccagtttactccctatcagtgatagagaacgtatgtcg<br>aggtaggcgtgtacggtgggcgcctataaaagcagagctcgtttagtgaaccgtcagatcgcctggagcaattccacaacactttt<br>gtcttatacttACGCGTGTCGACcatatgACATAGTGTCTAAGAAGTAATCCTAAATTAATCTATGAACAA<br>GAGCGTATTAAAATTATAAATGGCTCATAATTTTCTCTACGAGTGCAAACATATATAAAACATAAG<br>GGTCTATGGATTCTGAGTAAGTAACATACTTCTTATTATAATCTTCCGATGGTTCAAGCAAGAAGAA<br>GTTTTCTCAACTGAAAGGAAACTTTATTAATTATCTTAATGAGGAGTGAGGTACAGGGGAGATAAT<br>AATAATGAAATatgGGTAATTATAACCCGGGCCCTATATATTAGGCTAACTAGCTAACTAGGAATTC<br>ggtctcaCAGCAGCAGCAGCAGCAGCAGCAGCAGCAGCAGCAGCAGCACAGCAGCAGCAGCA<br>GCAGCAGCAGCAGCAGCAGCAGCAGCAGCAGCAGCAGCAGCAGCAGCAGCAGCAGCAGCA<br>GCAGCAGCAGCAGCAGCACAGCAGCAGCAGCAGCAGCAGCAGCAGCAGCAGCAGCAGCAG<br>CAGCAGCAGCAGCAGCAGCAGCAGCAGCAGCAGCAGCAGCAGCAGCACAGCAGCAGCAGC<br>AGCAGCAGCAGCAGCAGCAGCAGCAGCAGCAGCAGCAGCAGCAGCAGCAGCAGCAGCAGC<br>AGCAGCAGCAGCAGCACAGCAGCAGCAGCAGCAGCAGCAGCAGCAGCAGCAGCAGCAGCA<br>GCAGCAGCAGCAGCAGCAGCAGCAGCAGCAGCAGCAGCAGCAGCACAGCAGCAGCAGCAG<br>CAGCAGCAGCAGCAGCAGCAGCAGCAGCAGCAGCAGCAGCAGCAGCAGCAGCAGCAGCAG<br>CAGCAGCAGCAGCAGCACAGCAGCAGCAGCAGCAGCAGCAGCAGCAGCAGCAGCAGCAGC<br>AGCAGCAGCAGCAGCAGCAGCAGCACAGCAGCAGCAGCAGCAGCAGCAGCAGCAGCAGCA<br>GCAGCAGCAGCAGCAGCAGCAGCAGCAGCAGCAGCAGCAGCAGCAGCAGCAGCAGCACAG<br>CAGCAGCAGCAGCAGCAGCAGCAGCAGCAGCAGCAGCAgagacgGAGCTCgtcgacGCggccgca<br>agcttggtaccgagctcGGATCCTAAGGTACCTAATTGCCTAGAAAACATGAGGATCACCCATGTCTGCA<br>GGTCGACTCTAGAAAACATGAGGATCACCCATGTCTGCAGTATTCCCGGGTTCATTAGATCCTAAG<br>GTACCTAATTGCCTAGAAAACATGAGGATCACCCATGTCTGCAGGTCGACTCTAGAAAACATGAGG<br>ATCACCCATGTCTGCAGTATTCCCGGGTTCATTAGATCCTAAGGTACCTAATTGCCTAGAAAACATG<br>AGGATCACCCATGTCTGCAGGTCGACTCTAGAAAACATGAGGATCACCCATGTCTGCAGTATTCCC<br>GGGTTCATTAGATCCTAAGGTACCTAATTGCCTAGAAAACATGAGGATCACCCATGTCTGCAGGTC<br>GACTCTAGAAAACATGAGGATCACCCATGTCTGCAGTATTCCCGGGTTCATTAGATCCTAAGGTAC<br>CTAATTGCCTAGAAAACATGAGGATCACCCATGTCTGCAGGTCGACTCTAGAAAACATGAGGATCA<br>CCCATGTCTGCAGTATTCCCGGGTTCATTAGATCCTAAGGTACCTAATTGCCTAGAAAACATGAGG<br>ATCACCCATGTCTGCAGGTCGACTCTAGAAAACATGAGGATCACCCATGTCTGCAGTATTCCCGGG<br>TTCATTAGATCTGCGCGCGATCGATATCAGCGCTTTAAATTTGCGCATGCTAGTgTAAggcatgcaagc<br>ttgatatcaagcttatcgataatcaacctctggattacaaaatttgtgaaagattgactggtattcttaactatgttgctccttttacg<br>ctatgtggatacgctgctttaatgcctttgtatcatgctattgcttcccgtatggctttcattttctcctccttgtataaatcctggttgct<br>gtctctttatgaggagttgtggcccgttgtcaggcaacgtggcgtggtgtgcactgtgtttgctgacgcaacccccactggttgggg<br>cattgccaccacctgtcagctcctttccgggactttcgctttccccctccctattgccacggcggaactcatcgccgcctgccttgccc<br>gctgctggacaggggctcggctgttgggcactgacaattccgtggtgttgtcggggaaatcatcgtcctttccttggctgctcgcctg<br>tgttgccacctggattctgcgcgggacgtccttctgctacgtcccttcggccctcaatccagcggaccttccttcccgcggcctgctg<br>ccggctctgcggcctcttccgcgtcttcgccttcgccctcagacgagtcggatctccctttgggccgcctccccgcatcgataccgtc<br>gacctcgagggaattaattcgagctcggtacctttaagaccaatgacttacaaggcagctgtagatcttagccactttttaaaaga<br>aaaggggggactggaagggctaattcactcccaacgaagacaagatctgctttttgcttgtactgggtctctctggttagaccaga<br>tctgagcctgggagctctctggctaactagggaacccactgcttaagcctcaataaagcttgccttgagtgcttcaagtagtgtgtg<br>cccgtctgttgtgtgactctggtaactagagatccctcagacccttttagtcagtgtggaaaatctctagcagcatctagaattaatt<br>ccgtgtatt |

| | |
|---|---|
| N7: pHR Tre3G Flank N7 240xCAG 12xMS2 WPRE | ctatagtgtcacctaaatcgtatgtgtatgatacataaggttatgtattaattgtagccgcgttctaacgacaatatgtacaagccta<br>attgtgtagcatctggcttactgaagcagaccctatcatctctctcgtaaactgccgtcagagtcggtttggttggacgaaccttctg<br>agtttctggtaacgccgtcccgcacccggaaatggtcagcgaaccaatcagcagggtcatcgctagccagatcctctacgccgga<br>cgcatcgtggccggcatcaccggcgccacaggtgcggttgctggcgcctatatcgccgacatcaccgatggggaagatcgggctc<br>gccacttcgggctcatgagcgcttgtttcggcgtgggtatggtggcaggccccgtggccgggggactgttgggcgccatctccttgc<br>atgcaccattccttgcggcggcggtgctcaacggcctcaacctactactgggctgcttcctaatgcaggagtcgcataagggagag<br>cgtcgaatggtgcactctcagtacaatctgctctgatgccgcatagttaagccagccccgacacccgccaacacccgctgacgcgc<br>cctgacgggcttgtctgctcccggcatccgcttacagacaagctgtgaccgtctccgggagctgcatgtgtcagaggttttcaccgt<br>catcaccgaaacgcgcgagacgaaagggcctcgtgatacgcctatttttataggttaatgtcatgataataatggtttcttagacgt<br>caggtggcacttttcggggaaatgtgcgcggaacccctatttgtttatttttctaaatacattcaaatatgtatccgctcatgagaca<br>ataaccctgataaatgcttcaataatattgaaaaaggaagagtatgagtattcaacatttccgtgtcgcccttattcccttttttgcg<br>gcattttgccttcctgtttttgctcacccagaaacgctggtgaaagtaaaagatgctgaagatcagttgggtgcacgagtgggttac<br>atcgaactggatctcaacagcggtaagatccttgagagttttcgccccgaagaacgttttccaatgatgagcacttttaaagttctg<br>ctatgtggcgcggtattatcccgtattgacgccgggcaagagcaactcggtcgccgcatacactattctcagaatgacttggttga<br>gtactcaccagtcacagaaaagcatcttacggatggcatgacagtaagagaattatgcagtgctgccataaccatgagtgataac<br>actgcggccaacttacttctgacaacgatcggaggaccgaaggagctaaccgcttttttgcacaacatgggggatcatgtaactcg<br>ccttgatcgttgggaaccggagctgaatgaagccataccaaacgacgagcgtgacaccacgatgcctgtagcaatggcaacaac<br>gttgcgcaaactattaactggcgaactacttactctagcttcccggcaacaattaatagactggatggaggcggataaagttgcag<br>gaccacttctgcgctcggcccttccggctggctggtttattgctgataaatctggagccggtgagcgtgggtctcgcggtatcattgc<br>agcactggggccagatggtaagccctcccgtatcgtagttatctacacgacggggagtcaggcaactatggatgaacgaaatag<br>acagatcgctgagataggtgcctcactgattaagcattggtaactgtcagaccaagtttactcatatatactttagattgatttaaa<br>acttcatttttaatttaaaaggatctaggtgaagatcctttttgataatctcatgaccaaaatcccttaacgtgagttttcgttccact<br>gagcgtcagaccccgtagaaaagatcaaaggatcttcttgagatcctttttttctgcgcgtaatctgctgcttgcaaacaaaaaaa<br>ccaccgctaccagcggtggtttgtttgccggatcaagagctaccaactctttttccgaaggtaactggcttcagcagagcgcagat<br>accaaatactgtctttctagtgtagccgtagttaggccaccacttcaagaactctgtagcaccgcctacatacctcgctctgctaatc<br>ctgttaccagtggctgctgccagtggcgataagtcgtgtcttaccgggttggactcaagacgatagttaccggataaggcgcagcg<br>gtcgggctgaacggggggttcgtgcacacagcccagcttggagcgaacgacctacaccgaactgagatacctacagcgtgagct<br>atgagaaagcgccacgcttcccgaagggagaaaggcggacaggtatccggtaagcggcagggtcggaacaggagagcgcacg<br>agggagcttccagggggaaacgcctggtatctttatagtcctgtcgggtttcgccacctctgacttgagcgtcgatttttgtgatgct<br>cgtcaggggggcggagcctatggaaaaacgccagcaacgcggcctttttacggttcctggccttttgctggccttttgctcacatgt<br>tctttcctgcgttatcccctgattctgtggataaccgtattaccgcctttgagtgagctgataccgctcgccgcagccgaacgaccga<br>gcgcagcgagtcagtgagcgaggaagcggaagagcgcccaatacgcaaaccgcctctccccgcgcgttggccgattcattaatg<br>cagctgtggaatgtgtgtcagttagggtgtggaaagtccccaggctccccagcaggcagaagtatgcaaagcatgcatctcaatt<br>agtcagcaaccaggtgtggaaagtccccaggctccccagcaggcagaagtatgcaaagcatgcatctcaattagtcagcaacca<br>tagtcccgcccctaactccgcccatcccgcccctaactccgcccagttccgcccattctccgccccatggctgactaattttttttattt<br>atgcagaggccgaggccgcctcggcctctgagctattccagaagtagtgaggaggcttttttggaggcctaggcttttgcaaaaag<br>cttggacacaagacaggcttgcgagatatgtttgagaataccactttatcccgcgtcagggagaggcagtgcgtaaaaagacgc<br>ggactcatgtgaaatactggtttttagtgcgccagatctctataatctcgcgcaacctattttcccctcgaacactttttaagccgtag<br>ataaacaggctgggacacttcacatgagcgaaaaatacatcgtcacctgggacatgttgcagatccatgcacgtaaactcgcaa<br>gccgactgatgccttctgaacaatggaaaggcattattgccgtaagccgtggcggtctgtaccgggtgcgttactggcgcgtgaac<br>tgggtattcgtcatgtcgataccgtttgtatttccagctacgatcacgacaaccagcgcgagcttaaagtgctgaaacgcgcagaa<br>ggcgatggcgaaggcttcatcgttattgatgacctggtggataccggtggtactgcggttgcgattcgtgaaatgtatccaaaagc<br>gcactttgtcaccatcttcgcaaaaccggctggtcgtccgctggttgatgactatgttgttgatatcccgcaagatacctggattgaa<br>cagccgtgggatatgggcgtcgtattcgtcccgccaatctccggtcgctaatcttttcaacgcctggcactgccgggcgttgttctttt<br>taacttcaggcgggttacaatagtttccagtaagtattctggaggctgcatccatgacacaggcaaacctgagcgaaaccctgttc<br>aaaccccgctttaaacatcctgaaacctcgacgctagtccgccgctttaatcacggcgcacaaccgcctgtgcagtcggcccttga<br>tggtaaaaccatccctcactggtatcgcatgattaaccgtctgatgtggatctggcgcggcattgacccacgcgaaatcctcgacg |

| | |
|---|---|
| | tccaggcacgtattgtgatgagcgatgccgaacgtaccgacgatgatttatacgatacggtgattggctaccgtggcggcaactgg<br>atttatgagtgggccccggatctttgtgaaggaaccttacttctgtggtgtgacataattggacaaactacctacagagatttaaag<br>ctctaaggtaaatataaaatttttaagtgtataatgtgttaaactactgattctaattgtttgtgtattttagattccaacctatggaa<br>ctgatgaatgggagcagtggtggaatgcctttaatgaggaaaacctgttttgctcagaagaaatgccatctagtgatgatgaggct<br>actgctgactctcaacattctactcctccaaaaaagaagagaaaggtagaagaccccaaggactttccttcagaattgctaagttt<br>tttgagtcatgctgtgtttagtaatagaactcttgcttgctttgctatttacaccacaaaggaaaaagctgcactgctatacaagaa<br>aattatggaaaaatattctgtaacctttataagtaggcataacagttataatcataacatactgttttttcttactccacacaggcat<br>agagtgtctgctattaataactatgctcaaaaattgtgtacctttagctttttaatttgtaaaggggttaataaggaatatttgatgta<br>tagtgccttgactagagatcataatcagccataccacatttgtagaggttttacttgctttaaaaaacctcccacacctccccctga<br>acctgaaacataaaatgaatgcaattgttgttgttaacttgtttattgcagcttataatggttacaaataaagcaatagcatcacaa<br>atttcacaaataaagcatttttttcactgcattctagttgtggtttgtccaaactcatcaatgtatcttatcatgtctggatcaactgga<br>taactcaagctaaccaaaatcatcccaaacttcccaccccataccctattaccactgccaattacctagtggtttcatttactctaaa<br>cctgtgattcctctgaattattttcattttaaagaaattgtatttgttaaatatgtactacaaacttagtagttggaagggctaattca<br>ctcccaaagaagacaagatatccttgatctgtggatctaccacacacaaggctacttccctgattagcagaactacacaccaggg<br>ccaggggtcagatatccactgacctttggatggtgctacaagctagtaccagttgagccagataaggtagaagaggccaataaa<br>ggagagaacaccagcttgttacaccctgtgagcctgcatgggatggatgacccggagagagaagtgttagagtggaggtttgac<br>agccgcctagcatttcatcacgtggcccgagagctgcatccggagtacttcaagaactgctgatatcgagcttgctacaagggact<br>ttccgctggggactttccagggaggcgtggcctgggcgggactggggagtggcgagccctcagatcctgcatataagcagctgct<br>ttttgcctgtactgggtctctctggttagaccagatctgagcctgggagctctctggctaactagggaacccactgcttaagcctcaa<br>taaagcttgccttgagtgcttcaagtagtgtgtgcccgtctgttgtgtgactctggtaactagagatccctcagacccttttagtcagt<br>gtggaaaatctctagcagtggcgcccgaacagggacttgaaagcgaaagggaaaccagaggagctctctcgacgcaggactcg<br>gcttgctgaagcgcgcacggcaagaggcgaggggcggcgactggtgagtacgccaaaaattttgactagcggaggctagaagg<br>agagagatgggtgcgagagcgtcagtattaagcgggggagaattagatcgcgatgggaaaaaattcggttaaggccagggggа<br>aagaaaaaatataaattaaaacatatagtatgggcaagcagggagctagaacgattcgcagttaatcctggcctgttagaaaca<br>tcagaaggctgtagacaaatactgggacagctacaaccatcccttcagacaggatcagaagaacttagatcattatataatacag<br>tagcaaccctctattgtgtgcatcaaaggatagagataaaagacaccaaggaagctttagacaagatagaggaagagcaaaac<br>aaaagtaagaccaccgcacagcaagcggccggccgctgatcttcagacctggaggaggagatatgagggacaattggagaagt<br>gaattatataaatataaagtagtaaaaattgaaccattaggagtagcacccaccaaggcaaagagaagagtggtgcagagaga<br>aaaaagagcagtgggaataggagctttgttccttgggttcttgggagcagcaggaagcactatgggcgcagcgtcaatgacgctg<br>acggtacaggccagacaattattgtctggtatagtgcagcagcagaacaatttgctgagggctattgaggcgcaacagcatctgtt<br>gcaactcacagtctggggcatcaagcagctccaggcaagaatcctggctgtgaaagatacctaaaggatcaacagctcctgggg<br>atttggggttgctctggaaaactcatttgcaccactgctgtgccttggaatgctagttggagtaataaatctctggaacagatttgg<br>aatcacacgacctggatggagtgggacagagaaattaacaattacacaagcttaatacactccttaattgaagaatcgcaaaac<br>cagcaagaaaagaatgaacaagaattattggaattagataaatgggcaagtttgtggaattggtttaacataacaaattggctgt<br>ggtatataaaattattcataatgatagtaggaggcttggtaggtttaagaatagtttttgctgtactttctatagtgaatagagttag<br>gcagggatattcaccattatcgtttcagacccacctcccaaccccgaggggacccgacaggcccgaaggaatagaagaagaag<br>gtggagagagagacagagacagatccattcgattagtgaacggatctcgacggtatcgccaaatggcagtattcatccacaattt<br>taaaagaaaaggggggattggggggtacagtgcaggggaaagaatagtagacataatagcaacagacatacaaactaaagaa<br>ttacaaaaacaaattacaaaaattcaaaattttcgggtttattacagggacagcagagatccagtttatcgatgaggccctttcgt<br>cttcactcgagtttactccctatcagtgatagagaacgtatgaagagtttactccctatcagtgatagagaacgtatgcagacttta<br>ctccctatcagtgatagagaacgtataaggagtttactccctatcagtgatagagaacgtatgaccagtttactccctatcagtgat<br>agagaacgtatctacagtttactccctatcagtgatagagaacgtatatccagtttactccctatcagtgatagagaacgtatgtcg<br>aggtaggcgtgtacggtgggcgcctataaaagcagagctcgtttagtgaaccgtcagatcgcctggagcaattccacaacactttt<br>gtcttatacttACGCGTGTCGACcatatgATCACCAGGCAAGTGCTGCAGTATAACTAGGTACTACGTCA<br>GGTGCTAAGGTTAAGAGAGTATTTTCCTTCACTGACTCCTCACTCCGAGAATCCATTTTACAGCTTC<br>ATTGGTTTGGGTTATTCCAATTTTTTGATGTGAGTAAATAAATGACTTCTATTTGCCCAAAATAAAG<br>CTTATATAGGCCTTATAACCATGCAAATGTGTCCATTAAGTTGGACTTGGAATGAGTGAATGAGTA<br>TTACTGCCAGTatgGGTAATTATAACCCGGGCCCTATATATTAGGCTAACTAGCTAACTAGGAATTC |

| | |
|---|---|
| | ggtctcaCAGCAGCAGCAGCAGCAGCAGCAGCAGCAGCAGCAGCAGCACAGCAGCAGCAGCA<br>GCAGCAGCAGCAGCAGCAGCAGCAGCAGCAGCAGCAGCAGCAGCAGCAGCAGCAGCAGCA<br>GCAGCAGCAGCAGCAGCACAGCAGCAGCAGCAGCAGCAGCAGCAGCAGCAGCAGCAGCAG<br>CAGCAGCAGCAGCAGCAGCAGCAGCAGCAGCAGCAGCAGCAGCAGCACAGCAGCAGCAGC<br>AGCAGCAGCAGCAGCAGCAGCAGCAGCAGCAGCAGCAGCAGCAGCAGCAGCAGCAGCAGC<br>AGCAGCAGCAGCAGCACAGCAGCAGCAGCAGCAGCAGCAGCAGCAGCAGCAGCAGCAGCA<br>GCAGCAGCAGCAGCAGCAGCAGCAGCAGCAGCAGCAGCAGCAGCACAGCAGCAGCAGCAG<br>CAGCAGCAGCAGCAGCAGCAGCAGCAGCAGCAGCAGCAGCAGCAGCAGCAGCAGCAGCAG<br>CAGCAGCAGCAGCAGCACAGCAGCAGCAGCAGCAGCAGCAGCAGCAGCAGCAGCAGCAGC<br>AGCAGCAGCAGCAGCAGCAGCAGCAGCACAGCAGCAGCAGCAGCAGCAGCAGCAGCAGCA<br>GCAGCAGCAGCAGCAGCAGCAGCAGCAGCAGCAGCAGCAGCAGCAGCAGCAGCAGCACAG<br>CAGCAGCAGCAGCAGCAGCAGCAGCAGCAGCAGCAGCAGCAgagacgGAGCTCgtcgacGCggccgca<br>agcttggtaccgagctcGGATCCTAAGGTACCTAATTGCCTAGAAAACATGAGGATCACCCATGTCTGCA<br>GGTCGACTCTAGAAAACATGAGGATCACCCATGTCTGCAGTATTCCCGGGTTCATTAGATCCTAAG<br>GTACCTAATTGCCTAGAAAACATGAGGATCACCCATGTCTGCAGGTCGACTCTAGAAAACATGAGG<br>ATCACCCATGTCTGCAGTATTCCCGGGTTCATTAGATCCTAAGGTACCTAATTGCCTAGAAAACATG<br>AGGATCACCCATGTCTGCAGGTCGACTCTAGAAAACATGAGGATCACCCATGTCTGCAGTATTCCC<br>GGGTTCATTAGATCCTAAGGTACCTAATTGCCTAGAAAACATGAGGATCACCCATGTCTGCAGGTC<br>GACTCTAGAAAACATGAGGATCACCCATGTCTGCAGTATTCCCGGGTTCATTAGATCCTAAGGTAC<br>CTAATTGCCTAGAAAACATGAGGATCACCCATGTCTGCAGGTCGACTCTAGAAAACATGAGGATCA<br>CCCATGTCTGCAGTATTCCCGGGTTCATTAGATCCTAAGGTACCTAATTGCCTAGAAAACATGAGG<br>ATCACCCATGTCTGCAGGTCGACTCTAGAAAACATGAGGATCACCCATGTCTGCAGTATTCCCGGG<br>TTCATTAGATCTGCGCGCGATCGATATCAGCGCTTTAAATTTGCGCATGCTAGTgTAAggcatgcaagc<br>ttgatatcaagcttatcgataatcaacctctggattacaaaatttgtgaaagattgactggtattcttaactatgttgctccttttacg<br>ctatgtggatacgctgctttaatgcctttgtatcatgctattgcttcccgtatggctttcattttctcctccttgtataaatcctggttgct<br>gtctctttatgaggagttgtggcccgttgtcaggcaacgtggcgtggtgtgcactgtgtttgctgacgcaacccccactggttgggg<br>cattgccaccacctgtcagctcctttccgggactttcgctttccccctccctattgccacggcggaactcatcgccgcctgccttgccc<br>gctgctggacaggggctcggctgttgggcactgacaattccgtggtgttgtcggggaaatcatcgtcctttccttggctgctcgcctg<br>tgttgccacctggattctgcgcgggacgtccttctgctacgtcccttcggccctcaatccagcggaccttccttcccgcggcctgctg<br>ccggctctgcggcctcttccgcgtcttcgccttcgccctcagacgagtcggatctccctttgggccgcctccccgcatcgataccgtc<br>gacctcgagggaattaattcgagctcggtacctttaagaccaatgacttacaaggcagctgtagatcttagccactttttaaaaga<br>aaaggggggactggaagggctaattcactcccaacgaagacaagatctgctttttgcttgtactgggtctctctggttagaccaga<br>tctgagcctgggagctctctggctaactagggaacccactgcttaagcctcaataaagcttgccttgagtgcttcaagtagtgtgtg<br>cccgtctgttgtgtgactctggtaactagagatccctcagacccttttagtcagtgtggaaaatctctagcagcatctagaattaatt<br>ccgtgtatt |
| C1: pHR Tre3G Flank C1 240xCAG 12xMS2 WPRE | ctatagtgtcacctaaatcgtatgtgtatgatacataaggttatgtattaattgtagccgcgttctaacgacaatatgtacaagccta<br>attgtgtagcatctggcttactgaagcagaccctatcatctctctcgtaaactgccgtcagagtcggtttggttggacgaaccttctg<br>agtttctggtaacgccgtcccgcacccggaaatggtcagcgaaccaatcagcagggtcatcgctagccagatcctctacgccgga<br>cgcatcgtggccggcatcaccggcgccacaggtgcggttgctggcgcctatatcgccgacatcaccgatggggaagatcgggctc<br>gccacttcgggctcatgagcgcttgtttcggcgtgggtatggtggcaggccccgtggccgggggactgttgggcgccatctccttgc<br>atgcaccattccttgcggcggcggtgctcaacggcctcaacctactactgggctgcttcctaatgcaggagtcgcataagggagag<br>cgtcgaatggtgcactctcagtacaatctgctctgatgccgcatagttaagccagccccgacacccgccaacacccgctgacgcgc<br>cctgacgggcttgtctgctcccggcatccgcttacagacaagctgtgaccgtctccgggagctgcatgtgtcagaggttttcaccgt<br>catcaccgaaacgcgcgagacgaaagggcctcgtgatacgcctatttttataggttaatgtcatgataataatggtttcttagacgt<br>caggtggcacttttcggggaaatgtgcgcggaacccctatttgtttatttttctaaatacattcaaatatgtatccgctcatgagaca<br>ataaccctgataaatgcttcaataatattgaaaaaggaagagtatgagtattcaacatttccgtgtcgcccttattcccttttttgcg<br>gcattttgccttcctgtttttgctcacccagaaacgctggtgaaagtaaaagatgctgaagatcagttgggtgcacgagtgggttac |

| | |
|---|---|
| | atcgaactggatctcaacagcggtaagatccttgagagttttcgccccgaagaacgttttccaatgatgagcacttttaaagttctg<br>ctatgtggcgcggtattatcccgtattgacgccgggcaagagcaactcggtcgccgcatacactattctcagaatgacttggttga<br>gtactcaccagtcacagaaaagcatcttacggatggcatgacagtaagagaattatgcagtgctgccataaccatgagtgataac<br>actgcggccaacttacttctgacaacgatcggaggaccgaaggagctaaccgcttttttgcacaacatgggggatcatgtaactcg<br>ccttgatcgttgggaaccggagctgaatgaagccataccaaacgacgagcgtgacaccacgatgcctgtagcaatggcaacaac<br>gttgcgcaaactattaactggcgaactacttactctagcttcccggcaacaattaatagactggatggaggcggataaagttgcag<br>gaccacttctgcgctcggcccttccggctggctggtttattgctgataaatctggagccggtgagcgtgggtctcgcggtatcattgc<br>agcactggggccagatggtaagccctcccgtatcgtagttatctacacgacggggagtcaggcaactatggatgaacgaaatag<br>acagatcgctgagataggtgcctcactgattaagcattggtaactgtcagaccaagtttactcatatatactttagattgatttaaa<br>acttcatttttaatttaaaaggatctaggtgaagatcctttttgataatctcatgaccaaaatcccttaacgtgagttttcgttccact<br>gagcgtcagaccccgtagaaaagatcaaaggatcttcttgagatcctttttttctgcgcgtaatctgctgcttgcaaacaaaaaaa<br>ccaccgctaccagcggtggtttgtttgccggatcaagagctaccaactctttttccgaaggtaactggcttcagcagagcgcagat<br>accaaatactgtctttctagtgtagccgtagttaggccaccacttcaagaactctgtagcaccgcctacatacctcgctctgctaatc<br>ctgttaccagtggctgctgccagtggcgataagtcgtgtcttaccgggttggactcaagacgatagttaccggataaggcgcagcg<br>gtcgggctgaacggggggttcgtgcacacagcccagcttggagcgaacgacctacaccgaactgagatacctacagcgtgagct<br>atgagaaagcgccacgcttcccgaagggagaaaggcggacaggtatccggtaagcggcagggtcggaacaggagagcgcacg<br>agggagcttccagggggaaacgcctggtatctttatagtcctgtcgggtttcgccacctctgacttgagcgtcgatttttgtgatgct<br>cgtcaggggggcggagcctatggaaaaacgccagcaacgcggcctttttacggttcctggccttttgctggccttttgctcacatgt<br>tctttcctgcgttatcccctgattctgtggataaccgtattaccgcctttgagtgagctgataccgctcgccgcagccgaacgaccga<br>gcgcagcgagtcagtgagcgaggaagcggaagagcgcccaatacgcaaaccgcctctccccgcgcgttggccgattcattaatg<br>cagctgtggaatgtgtgtcagttagggtgtggaaagtccccaggctccccagcaggcagaagtatgcaaagcatgcatctcaatt<br>agtcagcaaccaggtgtggaaagtccccaggctccccagcaggcagaagtatgcaaagcatgcatctcaattagtcagcaacca<br>tagtcccgcccctaactccgcccatcccgcccctaactccgcccagttccgcccattctccgccccatggctgactaattttttttattt<br>atgcagaggccgaggccgcctcggcctctgagctattccagaagtagtgaggaggcttttttggaggcctaggcttttgcaaaaag<br>cttggacacaagacaggcttgcgagatatgtttgagaataccactttatcccgcgtcagggagaggcagtgcgtaaaaagacgc<br>ggactcatgtgaaatactggtttttagtgcgccagatctctataatctcgcgcaacctattttcccctcgaacactttttaagccgtag<br>ataaacaggctgggacacttcacatgagcgaaaaatacatcgtcacctgggacatgttgcagatccatgcacgtaaactcgcaa<br>gccgactgatgccttctgaacaatggaaaggcattattgccgtaagccgtggcggtctgtaccgggtgcgttactggcgcgtgaac<br>tgggtattcgtcatgtcgataccgtttgtatttccagctacgatcacgacaaccagcgcgagcttaaagtgctgaaacgcgcagaa<br>ggcgatggcgaaggcttcatcgttattgatgacctggtggataccggtggtactgcggttgcgattcgtgaaatgtatccaaaagc<br>gcactttgtcaccatcttcgcaaaaccggctggtcgtccgctggttgatgactatgttgttgatatcccgcaagatacctggattgaa<br>cagccgtgggatatgggcgtcgtattcgtcccgccaatctccggtcgctaatcttttcaacgcctggcactgccgggcgttgttctttt<br>taacttcaggcgggttacaatagtttccagtaagtattctggaggctgcatccatgacacaggcaaacctgagcgaaaccctgttc<br>aaaccccgctttaaacatcctgaaacctcgacgctagtccgccgctttaatcacggcgcacaaccgcctgtgcagtcggcccttga<br>tggtaaaaccatccctcactggtatcgcatgattaaccgtctgatgtggatctggcgcggcattgacccacgcgaaatcctcgacg<br>tccaggcacgtattgtgatgagcgatgccgaacgtaccgacgatgatttatacgatacggtgattggctaccgtggcggcaactgg<br>atttatgagtgggccccggatctttgtgaaggaaccttacttctgtggtgtgacataattggacaaactacctacagagatttaaag<br>ctctaaggtaaatataaaatttttaagtgtataatgtgttaaactactgattctaattgtttgtgtattttagattccaacctatggaa<br>ctgatgaatgggagcagtggtggaatgcctttaatgaggaaaacctgttttgctcagaagaaatgccatctagtgatgatgaggct<br>actgctgactctcaacattctactcctccaaaaaagaagagaaaggtagaagaccccaaggactttccttcagaattgctaagttt<br>tttgagtcatgctgtgtttagtaatagaactcttgcttgctttgctatttacaccacaaaggaaaaagctgcactgctatacaagaa<br>aattatggaaaaatattctgtaacctttataagtaggcataacagttataatcataacatactgttttttcttactccacacaggcat<br>agagtgtctgctattaataactatgctcaaaaattgtgtacctttagctttttaatttgtaaaggggttaataaggaatatttgatgta<br>tagtgccttgactagagatcataatcagccataccacatttgtagaggttttacttgctttaaaaaacctcccacacctccccctga<br>acctgaaacataaaatgaatgcaattgttgttgttaacttgtttattgcagcttataatggttacaaataaagcaatagcatcacaa<br>atttcacaaataaagcatttttttcactgcattctagttgtggtttgtccaaactcatcaatgtatcttatcatgtctggatcaactgga<br>taactcaagctaaccaaaatcatcccaaacttcccaccccataccctattaccactgccaattacctagtggtttcatttactctaaa<br>cctgtgattcctctgaattattttcattttaaagaaattgtatttgttaaatatgtactacaaacttagtagttggaagggctaattca |

| | |
|---|---|
| | ctcccaaagaagacaagatatccttgatctgtggatctaccacacacaaggctacttccctgattagcagaactacacaccaggg<br>ccaggggtcagatatccactgacctttggatggtgctacaagctagtaccagttgagccagataaggtagaagaggccaataaa<br>ggagagaacaccagcttgttacaccctgtgagcctgcatgggatggatgacccggagagagaagtgttagagtggaggtttgac<br>agccgcctagcatttcatcacgtggcccgagagctgcatccggagtacttcaagaactgctgatatcgagcttgctacaagggact<br>ttccgctggggactttccagggaggcgtggcctgggcgggactggggagtggcgagccctcagatcctgcatataagcagctgct<br>ttttgcctgtactgggtctctctggttagaccagatctgagcctgggagctctctggctaactagggaacccactgcttaagcctcaa<br>taaagcttgccttgagtgcttcaagtagtgtgtgcccgtctgttgtgtgactctggtaactagagatccctcagacccttttagtcagt<br>gtggaaaatctctagcagtggcgcccgaacagggacttgaaagcgaaagggaaaccagaggagctctctcgacgcaggactcg<br>gcttgctgaagcgcgcacggcaagaggcgaggggcggcgactggtgagtacgccaaaaattttgactagcggaggctagaagg<br>agagagatgggtgcgagagcgtcagtattaagcgggggagaattagatcgcgatgggaaaaaattcggttaaggccagggggа<br>aagaaaaaatataaattaaaacatatagtatgggcaagcagggagctagaacgattcgcagttaatcctggcctgttagaaaca<br>tcagaaggctgtagacaaatactgggacagctacaaccatcccttcagacaggatcagaagaacttagatcattatataatacag<br>tagcaaccctctattgtgtgcatcaaaggatagagataaaagacaccaaggaagctttagacaagatagaggaagagcaaaac<br>aaaagtaagaccaccgcacagcaagcggccggccgctgatcttcagacctggaggaggagatatgagggacaattggagaagt<br>gaattatataaatataaagtagtaaaaattgaaccattaggagtagcacccaccaaggcaaagagaagagtggtgcagagaga<br>aaaaagagcagtgggaataggagctttgttccttgggttcttgggagcagcaggaagcactatgggcgcagcgtcaatgacgctg<br>acggtacaggccagacaattattgtctggtatagtgcagcagcagaacaatttgctgagggctattgaggcgcaacagcatctgtt<br>gcaactcacagtctggggcatcaagcagctccaggcaagaatcctggctgtgaaagatacctaaaggatcaacagctcctgggg<br>atttggggttgctctggaaaactcatttgcaccactgctgtgccttggaatgctagttggagtaataaatctctggaacagatttgg<br>aatcacacgacctggatggagtgggacagagaaattaacaattacacaagcttaatacactccttaattgaagaatcgcaaaac<br>cagcaagaaaagaatgaacaagaattattggaattagataaatgggcaagtttgtggaattggtttaacataacaaattggctgt<br>ggtatataaaattattcataatgatagtaggaggcttggtaggtttaagaatagtttttgctgtactttctatagtgaatagagttag<br>gcagggatattcaccattatcgtttcagacccacctcccaaccccgaggggacccgacaggcccgaaggaatagaagaagaag<br>gtggagagagagacagagacagatccattcgattagtgaacggatctcgacggtatcgccaaatggcagtattcatccacaattt<br>taaaagaaaaggggggattggggggtacagtgcaggggaaagaatagtagacataatagcaacagacatacaaactaaagaa<br>ttacaaaaacaaattacaaaaattcaaaattttcgggtttattacagggacagcagagatccagtttatcgatgaggccctttcgt<br>cttcactcgagtttactccctatcagtgatagagaacgtatgaagagtttactccctatcagtgatagagaacgtatgcagacttta<br>ctccctatcagtgatagagaacgtataaggagtttactccctatcagtgatagagaacgtatgaccagtttactccctatcagtgat<br>agagaacgtatctacagtttactccctatcagtgatagagaacgtatatccagtttactccctatcagtgatagagaacgtatgtcg<br>aggtaggcgtgtacggtgggcgcctataaaagcagagctcgtttagtgaaccgtcagatcgcctggagcaattccacaacactttt<br>gtcttatacttACGCGTGTCGACcatatgAGGTTCCTGGCCACCGTCGGCGTCTCGCCCGACCACCAGGG<br>CAAGGGTCTGGGCAGCGCCGTCGTGCTCCCCGGAGTGGAGGCGGCCGAGCGCGCCGGGGTGCCC<br>GCCTTCCTGGAGACCTCCGCGCCCCGCAACCTCCCCTTCTACGAGCGGCTCGGCTTCACCGTCACCG<br>CCGACGTCGAGGTGCCCGAAGGACCGCGCACCTGGTGCAcGACCCGCAAGCCCGGTGCCGGATCG<br>GGAGAGGGCAGAGGTatgGGTAATTATAACCCGGGCCCTATATATTAGGCTAACTAGCTAACTAGG<br>AATTCggtctcaCAGCAGCAGCAGCAGCAGCAGCAGCAGCAGCAGCAGCAGCAGCACAGCAGCAGC<br>AGCAGCAGCAGCAGCAGCAGCAGCAGCAGCAGCAGCAGCAGCAGCAGCAGCAGCAGCAGCAGC<br>AGCAGCAGCAGCAGCAGCACAGCAGCAGCAGCAGCAGCAGCAGCAGCAGCAGCAGCAGCAGCA<br>GCAGCAGCAGCAGCAGCAGCAGCAGCAGCAGCAGCAGCAGCAGCAGCAGCAGCACAGCAGCAG<br>CAGCAGCAGCAGCAGCAGCAGCAGCAGCAGCAGCAGCAGCAGCAGCAGCAGCAGCAGCAGCAG<br>CAGCAGCAGCAGCAGCAGCACAGCAGCAGCAGCAGCAGCAGCAGCAGCAGCAGCAGCAGCAGC<br>AGCAGCAGCAGCAGCAGCAGCAGCAGCAGCAGCAGCAGCAGCAGCAGCAGCACAGCAGCAGCA<br>GCAGCAGCAGCAGCAGCAGCAGCAGCAGCAGCAGCAGCAGCAGCAGCAGCAGCAGCAGCAGCA<br>GCAGCAGCAGCAGCAGCACAGCAGCAGCAGCAGCAGCAGCAGCAGCAGCAGCAGCAGCAGCAG<br>CAGCAGCAGCAGCAGCAGCAGCAGCAGCAGCACAGCAGCAGCAGCAGCAGCAGCAGCAGCAGC<br>AGCAGCAGCAGCAGCAGCAGCAGCAGCAGCAGCAGCAGCAGCAGCAGCAGCAGCAGCAGCAGC<br>ACAGCAGCAGCAGCAGCAGCAGCAGCAGCAGCAGCAGCAGCAGCAgagacgGAGCTCgtcgacGCgg<br>ccgcaagcttggtaccgagctcGGATCCTAAGGTACCTAATTGCCTAGAAAACATGAGGATCACCCATGTC |

| | |
|---|---|
| | TGCAGGTCGACTCTAGAAAACATGAGGATCACCCATGTCTGCAGTATTCCCGGGTTCATTAGATCC<br>TAAGGTACCTAATTGCCTAGAAAACATGAGGATCACCCATGTCTGCAGGTCGACTCTAGAAAACAT<br>GAGGATCACCCATGTCTGCAGTATTCCCGGGTTCATTAGATCCTAAGGTACCTAATTGCCTAGAAA<br>ACATGAGGATCACCCATGTCTGCAGGTCGACTCTAGAAAACATGAGGATCACCCATGTCTGCAGTA<br>TTCCCGGGTTCATTAGATCCTAAGGTACCTAATTGCCTAGAAAACATGAGGATCACCCATGTCTGCA<br>GGTCGACTCTAGAAAACATGAGGATCACCCATGTCTGCAGTATTCCCGGGTTCATTAGATCCTAAG<br>GTACCTAATTGCCTAGAAAACATGAGGATCACCCATGTCTGCAGGTCGACTCTAGAAAACATGAGG<br>ATCACCCATGTCTGCAGTATTCCCGGGTTCATTAGATCCTAAGGTACCTAATTGCCTAGAAAACATG<br>AGGATCACCCATGTCTGCAGGTCGACTCTAGAAAACATGAGGATCACCCATGTCTGCAGTATTCCC<br>GGGTTCATTAGATCTGCGCGCGATCGATATCAGCGCTTTAAATTTGCGCATGCTAGTgTAAggcatgc<br>aagcttgatatcaagcttatcgataatcaacctctggattacaaaatttgtgaaagattgactggtattcttaactatgttgctcctttt<br>tacgctatgtggatacgctgctttaatgcctttgtatcatgctattgcttcccgtatggctttcattttctcctccttgtataaatcctgg<br>ttgctgtctctttatgaggagttgtggcccgttgtcaggcaacgtggcgtggtgtgcactgtgtttgctgacgcaacccccactggtt<br>ggggcattgccaccacctgtcagctcctttccgggactttcgctttccccctccctattgccacggcggaactcatcgccgcctgcct<br>tgcccgctgctggacaggggctcggctgttgggcactgacaattccgtggtgttgtcggggaaatcatcgtcctttccttggctgctc<br>gcctgtgttgccacctggattctgcgcgggacgtccttctgctacgtcccttcggccctcaatccagcggaccttccttcccgcggcc<br>tgctgccggctctgcggcctcttccgcgtcttcgccttcgccctcagacgagtcggatctccctttgggccgcctccccgcatcgata<br>ccgtcgacctcgagggaattaattcgagctcggtacctttaagaccaatgacttacaaggcagctgtagatcttagccactttttaa<br>aagaaaaggggggactggaagggctaattcactcccaacgaagacaagatctgctttttgcttgtactgggtctctctggttagac<br>cagatctgagcctgggagctctctggctaactagggaacccactgcttaagcctcaataaagcttgccttgagtgcttcaagtagt<br>gtgtgcccgtctgttgtgtgactctggtaactagagatccctcagacccttttagtcagtgtggaaaatctctagcagcatctagaat<br>taattccgtgtatt |
| C2: pHR Tre3G Flank C2 240xCAG 12xMS2 WPRE | ctatagtgtcacctaaatcgtatgtgtatgatacataaggttatgtattaattgtagccgcgttctaacgacaatatgtacaagccta<br>attgtgtagcatctggcttactgaagcagaccctatcatctctctcgtaaactgccgtcagagtcggtttggttggacgaaccttctg<br>agtttctggtaacgccgtcccgcacccggaaatggtcagcgaaccaatcagcagggtcatcgctagccagatcctctacgccgga<br>cgcatcgtggccggcatcaccggcgccacaggtgcggttgctggcgcctatatcgccgacatcaccgatggggaagatcgggctc<br>gccacttcgggctcatgagcgcttgtttcggcgtgggtatggtggcaggccccgtggccgggggactgttgggcgccatctccttgc<br>atgcaccattccttgcggcggcggtgctcaacggcctcaacctactactgggctgcttcctaatgcaggagtcgcataagggagag<br>cgtcgaatggtgcactctcagtacaatctgctctgatgccgcatagttaagccagccccgacacccgccaacacccgctgacgcgc<br>cctgacgggcttgtctgctcccggcatccgcttacagacaagctgtgaccgtctccgggagctgcatgtgtcagaggttttcaccgt<br>catcaccgaaacgcgcgagacgaaagggcctcgtgatacgcctatttttataggttaatgtcatgataataatggtttcttagacgt<br>caggtggcacttttcggggaaatgtgcgcggaacccctatttgtttatttttctaaatacattcaaatatgtatccgctcatgagaca<br>ataaccctgataaatgcttcaataatattgaaaaaggaagagtatgagtattcaacatttccgtgtcgcccttattcccttttttgcg<br>gcattttgccttcctgtttttgctcacccagaaacgctggtgaaagtaaaagatgctgaagatcagttgggtgcacgagtgggttac<br>atcgaactggatctcaacagcggtaagatccttgagagttttcgccccgaagaacgttttccaatgatgagcacttttaaagttctg<br>ctatgtggcgcggtattatcccgtattgacgccgggcaagagcaactcggtcgccgcatacactattctcagaatgacttggttga<br>gtactcaccagtcacagaaaagcatcttacggatggcatgacagtaagagaattatgcagtgctgccataaccatgagtgataac<br>actgcggccaacttacttctgacaacgatcggaggaccgaaggagctaaccgcttttttgcacaacatgggggatcatgtaactcg<br>ccttgatcgttgggaaccggagctgaatgaagccataccaaacgacgagcgtgacaccacgatgcctgtagcaatggcaacaac<br>gttgcgcaaactattaactggcgaactacttactctagcttcccggcaacaattaatagactggatggaggcggataaagttgcag<br>gaccacttctgcgctcggcccttccggctggctggtttattgctgataaatctggagccggtgagcgtgggtctcgcggtatcattgc<br>agcactggggccagatggtaagccctcccgtatcgtagttatctacacgacggggagtcaggcaactatggatgaacgaaatag<br>acagatcgctgagataggtgcctcactgattaagcattggtaactgtcagaccaagtttactcatatatactttagattgatttaaa<br>acttcatttttaatttaaaaggatctaggtgaagatcctttttgataatctcatgaccaaaatcccttaacgtgagttttcgttccact<br>gagcgtcagaccccgtagaaaagatcaaaggatcttcttgagatcctttttttctgcgcgtaatctgctgcttgcaaacaaaaaaa<br>ccaccgctaccagcggtggtttgtttgccggatcaagagctaccaactctttttccgaaggtaactggcttcagcagagcgcagat<br>accaaatactgtctttctagtgtagccgtagttaggccaccacttcaagaactctgtagcaccgcctacatacctcgctctgctaatc |

| | |
|---|---|
| | ctgttaccagtggctgctgccagtggcgataagtcgtgtcttaccgggttggactcaagacgatagttaccggataaggcgcagcg<br>gtcgggctgaacggggggttcgtgcacacagcccagcttggagcgaacgacctacaccgaactgagatacctacagcgtgagct<br>atgagaaagcgccacgcttcccgaagggagaaaggcggacaggtatccggtaagcggcagggtcggaacaggagagcgcacg<br>agggagcttccagggggaaacgcctggtatctttatagtcctgtcgggtttcgccacctctgacttgagcgtcgatttttgtgatgct<br>cgtcaggggggcggagcctatggaaaaacgccagcaacgcggcctttttacggttcctggccttttgctggccttttgctcacatgt<br>tctttcctgcgttatcccctgattctgtggataaccgtattaccgcctttgagtgagctgataccgctcgccgcagccgaacgaccga<br>gcgcagcgagtcagtgagcgaggaagcggaagagcgcccaatacgcaaaccgcctctccccgcgcgttggccgattcattaatg<br>cagctgtggaatgtgtgtcagttagggtgtggaaagtccccaggctccccagcaggcagaagtatgcaaagcatgcatctcaatt<br>agtcagcaaccaggtgtggaaagtccccaggctccccagcaggcagaagtatgcaaagcatgcatctcaattagtcagcaacca<br>tagtcccgcccctaactccgcccatcccgcccctaactccgcccagttccgcccattctccgccccatggctgactaattttttttattt<br>atgcagaggccgaggccgcctcggcctctgagctattccagaagtagtgaggaggcttttttggaggcctaggcttttgcaaaaag<br>cttggacacaagacaggcttgcgagatatgtttgagaataccactttatcccgcgtcagggagaggcagtgcgtaaaaagacgc<br>ggactcatgtgaaatactggtttttagtgcgccagatctctataatctcgcgcaacctattttcccctcgaacactttttaagccgtag<br>ataaacaggctgggacacttcacatgagcgaaaaatacatcgtcacctgggacatgttgcagatccatgcacgtaaactcgcaa<br>gccgactgatgccttctgaacaatggaaaggcattattgccgtaagccgtggcggtctgtaccgggtgcgttactggcgcgtgaac<br>tgggtattcgtcatgtcgataccgtttgtatttccagctacgatcacgacaaccagcgcgagcttaaagtgctgaaacgcgcagaa<br>ggcgatggcgaaggcttcatcgttattgatgacctggtggataccggtggtactgcggttgcgattcgtgaaatgtatccaaaagc<br>gcactttgtcaccatcttcgcaaaaccggctggtcgtccgctggttgatgactatgttgttgatatcccgcaagatacctggattgaa<br>cagccgtgggatatgggcgtcgtattcgtcccgccaatctccggtcgctaatcttttcaacgcctggcactgccgggcgttgttctttt<br>taacttcaggcgggttacaatagtttccagtaagtattctggaggctgcatccatgacacaggcaaacctgagcgaaaccctgttc<br>aaaccccgctttaaacatcctgaaacctcgacgctagtccgccgctttaatcacggcgcacaaccgcctgtgcagtcggcccttga<br>tggtaaaaccatccctcactggtatcgcatgattaaccgtctgatgtggatctggcgcggcattgacccacgcgaaatcctcgacg<br>tccaggcacgtattgtgatgagcgatgccgaacgtaccgacgatgatttatacgatacggtgattggctaccgtggcggcaactgg<br>atttatgagtgggccccggatctttgtgaaggaaccttacttctgtggtgtgacataattggacaaactacctacagagatttaaag<br>ctctaaggtaaatataaaatttttaagtgtataatgtgttaaactactgattctaattgtttgtgtattttagattccaacctatggaa<br>ctgatgaatgggagcagtggtggaatgcctttaatgaggaaaacctgttttgctcagaagaaatgccatctagtgatgatgaggct<br>actgctgactctcaacattctactcctccaaaaaagaagagaaaggtagaagaccccaaggactttccttcagaattgctaagttt<br>tttgagtcatgctgtgtttagtaatagaactcttgcttgctttgctatttacaccacaaaggaaaaagctgcactgctatacaagaa<br>aattatggaaaaatattctgtaacctttataagtaggcataacagttataatcataacatactgttttttcttactccacacaggcat<br>agagtgtctgctattaataactatgctcaaaaattgtgtacctttagctttttaatttgtaaaggggttaataaggaatatttgatgta<br>tagtgccttgactagagatcataatcagccataccacatttgtagaggttttacttgctttaaaaaacctcccacacctccccctga<br>acctgaaacataaaatgaatgcaattgttgttgttaacttgtttattgcagcttataatggttacaaataaagcaatagcatcacaa<br>atttcacaaataaagcatttttttcactgcattctagttgtggtttgtccaaactcatcaatgtatcttatcatgtctggatcaactgga<br>taactcaagctaaccaaaatcatcccaaacttcccaccccataccctattaccactgccaattacctagtggtttcatttactctaaa<br>cctgtgattcctctgaattattttcattttaaagaaattgtatttgttaaatatgtactacaaacttagtagttggaagggctaattca<br>ctcccaaagaagacaagatatccttgatctgtggatctaccacacacaaggctacttccctgattagcagaactacacaccaggg<br>ccaggggtcagatatccactgacctttggatggtgctacaagctagtaccagttgagccagataaggtagaagaggccaataaa<br>ggagagaacaccagcttgttacaccctgtgagcctgcatgggatggatgacccggagagagaagtgttagagtggaggtttgac<br>agccgcctagcatttcatcacgtggcccgagagctgcatccggagtacttcaagaactgctgatatcgagcttgctacaagggact<br>ttccgctggggactttccagggaggcgtggcctgggcgggactggggagtggcgagccctcagatcctgcatataagcagctgct<br>ttttgcctgtactgggtctctctggttagaccagatctgagcctgggagctctctggctaactagggaacccactgcttaagcctcaa<br>taaagcttgccttgagtgcttcaagtagtgtgtgcccgtctgttgtgtgactctggtaactagagatccctcagacccttttagtcagt<br>gtggaaaatctctagcagtggcgcccgaacagggacttgaaagcgaaagggaaaccagaggagctctctcgacgcaggactcg<br>gcttgctgaagcgcgcacggcaagaggcgaggggcggcgactggtgagtacgccaaaaattttgactagcggaggctagaagg<br>agagagatgggtgcgagagcgtcagtattaagcgggggagaattagatcgcgatgggaaaaaattcggttaaggccagggggа<br>aagaaaaaatataaattaaaacatatagtatgggcaagcagggagctagaacgattcgcagttaatcctggcctgttagaaaca<br>tcagaaggctgtagacaaatactgggacagctacaaccatcccttcagacaggatcagaagaacttagatcattatataatacag<br>tagcaaccctctattgtgtgcatcaaaggatagagataaaagacaccaaggaagctttagacaagatagaggaagagcaaaac |

| | |
|---|---|
| | aaaagtaagaccaccgcacagcaagcggccggccgctgatcttcagacctggaggaggagatatgagggacaattggagaagt<br>gaattatataaatataaagtagtaaaaattgaaccattaggagtagcacccaccaaggcaaagagaagagtggtgcagagaga<br>aaaaagagcagtgggaataggagctttgttccttgggttcttgggagcagcaggaagcactatgggcgcagcgtcaatgacgctg<br>acggtacaggccagacaattattgtctggtatagtgcagcagcagaacaatttgctgagggctattgaggcgcaacagcatctgtt<br>gcaactcacagtctggggcatcaagcagctccaggcaagaatcctggctgtgaaagatacctaaaggatcaacagctcctgggg<br>atttggggttgctctggaaaactcatttgcaccactgctgtgccttggaatgctagttggagtaataaatctctggaacagatttgg<br>aatcacacgacctggatggagtgggacagagaaattaacaattacacaagcttaatacactccttaattgaagaatcgcaaaac<br>cagcaagaaaagaatgaacaagaattattggaattagataaatgggcaagtttgtggaattggtttaacataacaaattggctgt<br>ggtatataaaattattcataatgatagtaggaggcttggtaggtttaagaatagtttttgctgtactttctatagtgaatagagttag<br>gcagggatattcaccattatcgtttcagacccacctcccaaccccgaggggacccgacaggcccgaaggaatagaagaagaag<br>gtggagagagagacagagacagatccattcgattagtgaacggatctcgacggtatcgccaaatggcagtattcatccacaattt<br>taaaagaaaaggggggattggggggtacagtgcaggggaaagaatagtagacataatagcaacagacatacaaactaaagaa<br>ttacaaaaacaaattacaaaaattcaaaattttcgggtttattacagggacagcagagatccagtttatcgatgaggccctttcgt<br>cttcactcgagtttactccctatcagtgatagagaacgtatgaagagtttactccctatcagtgatagagaacgtatgcagacttta<br>ctccctatcagtgatagagaacgtataaggagtttactccctatcagtgatagagaacgtatgaccagtttactccctatcagtgat<br>agagaacgtatctacagtttactccctatcagtgatagagaacgtatatccagtttactccctatcagtgatagagaacgtatgtcg<br>aggtaggcgtgtacggtgggcgcctataaaagcagagctcgtttagtgaaccgtcagatcgcctggagcaattccacaacactttt<br>gtcttatacttACGCGTGTCGACcatatgATTACAGACTATCCGGGTCGCTAGCCGGTACCAGGCACTTC<br>GTGGGATCGCCTTGCGAGCGGCACGCGGAGGTCCACCCCTCATGTCGATGTACCGACGCAGCTCC<br>CGATGTTCTGCCCAGAATATTGCAGGGGGTCCGAACTCCACGCGGACTAGGCAGAGGGCCTGGAA<br>TGCGGCCGGCGCATTCCCGGGGGTGGTCCTGAGGCTGCGGACGGCGGGGTATTGGCGAATCGGC<br>GAGGCGGATTGTGAGATatgGGTAATTATAACCCGGGCCCTATATATTAGGCTAACTAGCTAACTA<br>GGAATTCggtctcaCAGCAGCAGCAGCAGCAGCAGCAGCAGCAGCAGCAGCAGCACAGCAGCA<br>GCAGCAGCAGCAGCAGCAGCAGCAGCAGCAGCAGCAGCAGCAGCAGCAGCAGCAGCAGCA<br>GCAGCAGCAGCAGCAGCAGCACAGCAGCAGCAGCAGCAGCAGCAGCAGCAGCAGCAGCAG<br>CAGCAGCAGCAGCAGCAGCAGCAGCAGCAGCAGCAGCAGCAGCAGCAGCAGCACAGCAGC<br>AGCAGCAGCAGCAGCAGCAGCAGCAGCAGCAGCAGCAGCAGCAGCAGCAGCAGCAGCAGC<br>AGCAGCAGCAGCAGCAGCACAGCAGCAGCAGCAGCAGCAGCAGCAGCAGCAGCAGCAGCA<br>GCAGCAGCAGCAGCAGCAGCAGCAGCAGCAGCAGCAGCAGCAGCAGCACAGCAGCAG<br>CAGCAGCAGCAGCAGCAGCAGCAGCAGCAGCAGCAGCAGCAGCAGCAGCAGCAGCAGCAG<br>CAGCAGCAGCAGCAGCAGCAGCACAGCAGCAGCAGCAGCAGCAGCAGCAGCAGCAGCAGC<br>AGCAGCAGCAGCAGCAGCAGCAGCAGCAGCAGCACAGCAGCAGCAGCAGCAGCAGCAGCA<br>GCAGCAGCAGCAGCAGCAGCAGCAGCAGCAGCAGCAGCAGCAGCAGCAGCAGCAGCAGCA<br>GCACAGCAGCAGCAGCAGCAGCAGCAGCAGCAGCAGCAGCAGCAgagacgGAGCTCgtcgacG<br>CggccgcaagcttggtaccgagctcGGATCCTAAGGTACCTAATTGCCTAGAAAACATGAGGATCACCCAT<br>GTCTGCAGGTCGACTCTAGAAAACATGAGGATCACCCATGTCTGCAGTATTCCCGGGTTCATTAGA<br>TCCTAAGGTACCTAATTGCCTAGAAAACATGAGGATCACCCATGTCTGCAGGTCGACTCTAGAAAA<br>CATGAGGATCACCCATGTCTGCAGTATTCCCGGGTTCATTAGATCCTAAGGTACCTAATTGCCTAGA<br>AAACATGAGGATCACCCATGTCTGCAGGTCGACTCTAGAAAACATGAGGATCACCCATGTCTGCAG<br>TATTCCCGGGTTCATTAGATCCTAAGGTACCTAATTGCCTAGAAAACATGAGGATCACCCATGTCTG<br>CAGGTCGACTCTAGAAAACATGAGGATCACCCATGTCTGCAGTATTCCCGGGTTCATTAGATCCTA<br>AGGTACCTAATTGCCTAGAAAACATGAGGATCACCCATGTCTGCAGGTCGACTCTAGAAAACATGA<br>GGATCACCCATGTCTGCAGTATTCCCGGGTTCATTAGATCCTAAGGTACCTAATTGCCTAGAAAACA<br>TGAGGATCACCCATGTCTGCAGGTCGACTCTAGAAAACATGAGGATCACCCATGTCTGCAGTATTC<br>CCGGGTTCATTAGATCTGCGCGCGATCGATATCAGCGCTTTAAATTTGCGCATGCTAGTgTAAggcat<br>gcaagcttgatatcaagcttatcgataatcaacctctggattacaaaatttgtgaaagattgactggtattcttaactatgttgctcc<br>ttttacgctatgtggatacgctgctttaatgcctttgtatcatgctattgcttcccgtatggctttcattttctcctccttgtataaatcct<br>ggttgctgtctctttatgaggagttgtggcccgttgtcaggcaacgtggcgtggtgtgcactgtgtttgctgacgcaacccccactgg |

| | |
|---|---|
| | ttggggcattgccaccacctgtcagctcctttccgggactttcgctttccccctccctattgccacggcggaactcatcgccgcctgc<br>cttgcccgctgctggacaggggctcggctgttgggcactgacaattccgtggtgttgtcggggaaatcatcgtcctttccttggctgc<br>tcgcctgtgttgccacctggattctgcgcgggacgtccttctgctacgtcccttcggccctcaatccagcggaccttccttcccgcgg<br>cctgctgccggctctgcggcctcttccgcgtcttcgccttcgccctcagacgagtcggatctccctttgggccgcctccccgcatcga<br>taccgtcgacctcgagggaattaattcgagctcggtacctttaagaccaatgacttacaaggcagctgtagatcttagccactttt t<br>aaaagaaaaggggggactggaagggctaattcactcccaacgaagacaagatctgctttttgcttgtactgggtctctctggttag<br>accagatctgagcctgggagctctctggctaactagggaacccactgcttaagcctcaataaagcttgccttgagtgcttcaagta<br>gtgtgtgcccgtctgttgtgtgactctggtaactagagatccctcagacccttttagtcagtgtggaaaatctctagcagcatctaga<br>attaattccgtgtatt |
| C3: pHR Tre3G Flank C3 240xCA G 12xMS2 WPRE | ctatagtgtcacctaaatcgtatgtgtatgatacataaggttatgtattaattgtagccgcgttctaacgacaatatgtacaagccta<br>attgtgtagcatctggcttactgaagcagaccctatcatctctctcgtaaactgccgtcagagtcggtttggttggacgaaccttctg<br>agtttctggtaacgccgtcccgcacccggaaatggtcagcgaaccaatcagcagggtcatcgctagccagatcctctacgccgga<br>cgcatcgtggccggcatcaccggcgccacaggtgcggttgctggcgcctatatcgccgacatcaccgatggggaagatcgggctc<br>gccacttcgggctcatgagcgcttgtttcggcgtgggtatggtggcaggccccgtggccgggggactgttgggcgccatctccttgc<br>atgcaccattccttgcggcggcggtgctcaacggcctcaacctactactgggctgcttcctaatgcaggagtcgcataagggagag<br>cgtcgaatggtgcactctcagtacaatctgctctgatgccgcatagttaagccagccccgacacccgccaacacccgctgacgcgc<br>cctgacgggcttgtctgctcccggcatccgcttacagacaagctgtgaccgtctccgggagctgcatgtgtcagaggttttcaccgt<br>catcaccgaaacgcgcgagacgaaagggcctcgtgatacgcctatttttataggttaatgtcatgataataatggtttcttagacgt<br>caggtggcacttttcggggaaatgtgcgcggaacccctatttgtttatttttctaaatacattcaaatatgtatccgctcatgagaca<br>ataaccctgataaatgcttcaataatattgaaaaaggaagagtatgagtattcaacatttccgtgtcgcccttattcccttttttgcg<br>gcattttgccttcctgtttttgctcacccagaaacgctggtgaaagtaaaagatgctgaagatcagttgggtgcacgagtgggttac<br>atcgaactggatctcaacagcggtaagatccttgagagttttcgccccgaagaacgttttccaatgatgagcacttttaaagttctg<br>ctatgtggcgcggtattatcccgtattgacgccgggcaagagcaactcggtcgccgcatacactattctcagaatgacttggttga<br>gtactcaccagtcacagaaaagcatcttacggatggcatgacagtaagagaattatgcagtgctgccataaccatgagtgataac<br>actgcggccaacttacttctgacaacgatcggaggaccgaaggagctaaccgcttttttgcacaacatgggggatcatgtaactcg<br>ccttgatcgttgggaaccggagctgaatgaagccataccaaacgacgagcgtgacaccacgatgcctgtagcaatggcaacaac<br>gttgcgcaaactattaactggcgaactacttactctagcttcccggcaacaattaatagactggatggaggcggataaagttgcag<br>gaccacttctgcgctcggcccttccggctggctggtttattgctgataaatctggagccggtgagcgtgggtctcgcggtatcattgc<br>agcactggggccagatggtaagccctcccgtatcgtagttatctacacgacggggagtcaggcaactatggatgaacgaaatag<br>acagatcgctgagataggtgcctcactgattaagcattggtaactgtcagaccaagtttactcatatatactttagattgatttaaa<br>acttcatttttaatttaaaaggatctaggtgaagatcctttttgataatctcatgaccaaaatcccttaacgtgagttttcgttccact<br>gagcgtcagaccccgtagaaaagatcaaaggatcttcttgagatcctttttttctgcgcgtaatctgctgcttgcaaacaaaaaa<br>ccaccgctaccagcggtggtttgtttgccggatcaagagctaccaactctttttccgaaggtaactggcttcagcagagcgcagat<br>accaaatactgtctttctagtgtagccgtagttaggccaccacttcaagaactctgtagcaccgcctacatacctcgctctgctaatc<br>ctgttaccagtggctgctgccagtggcgataagtcgtgtcttaccgggttggactcaagacgatagttaccggataaggcgcagcg<br>gtcgggctgaacggggggttcgtgcacacagcccagcttggagcgaacgacctacaccgaactgagatacctacagcgtgagct<br>atgagaaagcgccacgcttcccgaagggagaaaggcggacaggtatccggtaagcggcagggtcggaacaggagagcgcacg<br>agggagcttccagggggaaacgcctggtatctttatagtcctgtcgggtttcgccacctctgacttgagcgtcgatttttgtgatgct<br>cgtcaggggggcggagcctatggaaaaacgccagcaacgcggcctttttacggttcctggccttttgctggccttttgctcacatgt<br>tctttcctgcgttatcccctgattctgtggataaccgtattaccgcctttgagtgagctgataccgctcgccgcagccgaacgaccga<br>gcgcagcgagtcagtgagcgaggaagcggaagagcgcccaatacgcaaaccgcctctccccgcgcgttggccgattcattaatg<br>cagctgtggaatgtgtgtcagttagggtgtggaaagtccccaggctccccagcaggcagaagtatgcaaagcatgcatctcaatt<br>agtcagcaaccaggtgtggaaagtccccaggctccccagcaggcagaagtatgcaaagcatgcatctcaattagtcagcaacca<br>tagtcccgcccctaactccgcccatcccgcccctaactccgcccagttccgcccattctccgccccatggctgactaattttttttattt<br>atgcagaggccgaggccgcctcggcctctgagctattccagaagtagtgaggaggcttttttggaggcctaggcttttgcaaaaag<br>cttggacacaagacaggcttgcgagatatgtttgagaataccactttatcccgcgtcagggagaggcagtgcgtaaaagacgc<br>ggactcatgtgaaatactggtttttagtgcgccagatctctataatctcgcgcaacctattttcccctcgaacacttttaagccgtag |

| | |
|---|---|
| | ataaacaggctgggacacttcacatgagcgaaaaatacatcgtcacctgggacatgttgcagatccatgcacgtaaactcgcaa<br>gccgactgatgccttctgaacaatggaaaggcattattgccgtaagccgtggcggtctgtaccgggtgcgttactggcgcgtgaac<br>tgggtattcgtcatgtcgataccgtttgtatttccagctacgatcacgacaaccagcgcgagcttaaagtgctgaaacgcgcagaa<br>ggcgatggcgaaggcttcatcgttattgatgacctggtggataccggtggtactgcggttgcgattcgtgaaatgtatccaaaagc<br>gcactttgtcaccatcttcgcaaaaccggctggtcgtccgctggttgatgactatgttgttgatatcccgcaagatacctggattgaa<br>cagccgtgggatatgggcgtcgtattcgtcccgccaatctccggtcgctaatcttttcaacgcctggcactgccgggcgttgttctttt<br>taacttcaggcgggttacaatagtttccagtaagtattctggaggctgcatccatgacacaggcaaacctgagcgaaaccctgttc<br>aaaccccgctttaaacatcctgaaacctcgacgctagtccgccgctttaatcacggcgcacaaccgcctgtgcagtcggcccttga<br>tggtaaaaccatccctcactggtatcgcatgattaaccgtctgatgtggatctggcgcggcattgacccacgcgaaatcctcgacg<br>tccaggcacgtattgtgatgagcgatgccgaacgtaccgacgatgatttatacgatacggtgattggctaccgtggcggcaactgg<br>atttatgagtgggccccggatctttgtgaaggaaccttacttctgtggtgtgacataattggacaaactacctacagagatttaaag<br>ctctaaggtaaatataaaatttttaagtgtataatgtgttaaactactgattctaattgtttgtgtattttagattccaacctatggaa<br>ctgatgaatgggagcagtggtggaatgcctttaatgaggaaaacctgttttgctcagaagaaatgccatctagtgatgatgaggct<br>actgctgactctcaacattctactcctccaaaaaagaagagaaaggtagaagaccccaaggactttccttcagaattgctaagttt<br>tttgagtcatgctgtgtttagtaatagaactcttgcttgctttgctatttacaccacaaaggaaaaagctgcactgctatacaagaa<br>aattatggaaaaatattctgtaacctttataagtaggcataacagttataatcataacatactgttttttcttactccacacaggcat<br>agagtgtctgctattaataactatgctcaaaaattgtgtacctttagctttttaatttgtaaaggggttaataaggaatatttgatgta<br>tagtgccttgactagagatcataatcagccataccacatttgtagaggttttacttgctttaaaaaacctcccacacctccccctga<br>acctgaaacataaaatgaatgcaattgttgttgttaacttgtttattgcagcttataatggttacaaataaagcaatagcatcacaa<br>atttcacaaataaagcatttttttcactgcattctagttgtggtttgtccaaactcatcaatgtatcttatcatgtctggatcaactgga<br>taactcaagctaaccaaaatcatcccaaacttcccaccccataccctattaccactgccaattacctagtggtttcatttactctaaa<br>cctgtgattcctctgaattattttcattttaaagaaattgtatttgttaaatatgtactacaaacttagtagttggaagggctaattca<br>ctcccaaagaagacaagatatccttgatctgtggatctaccacacacaaggctacttccctgattagcagaactacacaccaggg<br>ccaggggtcagatatccactgacctttggatggtgctacaagctagtaccagttgagccagataaggtagaagaggccaataaa<br>ggagagaacaccagcttgttacaccctgtgagcctgcatgggatggatgacccggagagagaagtgttagagtggaggtttgac<br>agccgcctagcatttcatcacgtggcccgagagctgcatccggagtacttcaagaactgctgatatcgagcttgctacaagggact<br>ttccgctggggactttccagggaggcgtggcctgggcgggactggggagtggcgagccctcagatcctgcatataagcagctgct<br>ttttgcctgtactgggtctctctggttagaccagatctgagcctgggagctctctggctaactagggaacccactgcttaagcctcaa<br>taaagcttgccttgagtgcttcaagtagtgtgtgcccgtctgttgtgtgactctggtaactagagatccctcagacccttttagtcagt<br>gtggaaaatctctagcagtggcgcccgaacagggacttgaaagcgaaagggaaaccagaggagctctctcgacgcaggactcg<br>gcttgctgaagcgcgcacggcaagaggcgaggggcggcgactggtgagtacgccaaaaattttgactagcggaggctagaagg<br>agagagatgggtgcgagagcgtcagtattaagcgggggagaattagatcgcgatgggaaaaaattcggttaaggccagggggа<br>aagaaaaaatataaattaaaacatatagtatgggcaagcagggagctagaacgattcgcagttaatcctggcctgttagaaaca<br>tcagaaggctgtagacaaatactgggacagctacaaccatcccttcagacaggatcagaagaacttagatcattatataatacag<br>tagcaaccctctattgtgtgcatcaaaggatagagataaaagacaccaaggaagctttagacaagatagaggaagagcaaaac<br>aaaagtaagaccaccgcacagcaagcggccggccgctgatcttcagacctggaggaggagatatgagggacaattggagaagt<br>gaattatataaatataaagtagtaaaaattgaaccattaggagtagcacccaccaaggcaaagagaagagtggtgcagagaga<br>aaaaagagcagtgggaataggagctttgttccttgggttcttgggagcagcaggaagcactatgggcgcagcgtcaatgacgctg<br>acggtacaggccagacaattattgtctggtatagtgcagcagcagaacaatttgctgagggctattgaggcgcaacagcatctgtt<br>gcaactcacagtctggggcatcaagcagctccaggcaagaatcctggctgtgaaagatacctaaaggatcaacagctcctgggg<br>atttggggttgctctggaaaactcatttgcaccactgctgtgccttggaatgctagttggagtaataaatctctggaacagatttgg<br>aatcacacgacctggatggagtgggacagagaaattaacaattacacaagcttaatacactccttaattgaagaatcgcaaaac<br>cagcaagaaaagaatgaacaagaattattggaattagataaatgggcaagtttgtggaattggtttaacataacaaattggctgt<br>ggtatataaaattattcataatgatagtaggaggcttggtaggtttaagaatagtttttgctgtactttctatagtgaatagagttag<br>gcagggatattcaccattatcgtttcagacccacctcccaaccccgaggggacccgacaggcccgaaggaatagaagaagaag<br>gtggagagagagacagagacagatccattcgattagtgaacggatctcgacggtatcgccaaatggcagtattcatccacaattt<br>taaaagaaaaggggggattggggggtacagtgcaggggaaagaatagtagacataatagcaacagacatacaaactaaagaa<br>ttacaaaaacaaattacaaaaattcaaaattttcgggtttattacagggacagcagagatccagtttatcgatgaggccctttcgt |

| | |
|---|---|
| | cttcactcgagtttactccctatcagtgatagagaacgtatgaagagtttactccctatcagtgatagagaacgtatgcagacttta<br>ctccctatcagtgatagagaacgtataaggagtttactccctatcagtgatagagaacgtatgaccagtttactccctatcagtgat<br>agagaacgtatctacagtttactccctatcagtgatagagaacgtatatccagtttactccctatcagtgatagagaacgtatgtcg<br>aggtaggcgtgtacggtgggcgcctataaaagcagagctcgtttagtgaaccgtcagatcgcctggagcaattccacaacactttt<br>gtcttatacttACGCGTGTCGACcatatgATTGGGTCTGGGCTCATAGGGTGTTCACATCTGGGGAGTAT<br>GGTCGGATCTAATACGTCGTTTGGGAAATCAAGAGTCGGCAGCGCAGGTCCCACCAACTACTCAT<br>GTGATTCGCACTTACTATTCCTACTCGATGGAGAGGATCTAGGGTTAGCAGCCAAGCCGGAGCCTC<br>ACATATCACCGTCACTAGTCGGTAATATCCTGCCTATGCAAAGCACGATGAAGCATAGATTCATTCT<br>CGTCATTCGGGATatgGGTAATTATAACCCGGGCCCTATATATTAGGCTAACTAGCTAACTAGGAAT<br>TCggtctcaCAGCAGCAGCAGCAGCAGCAGCAGCAGCAGCAGCAGCAGCACAGCAGCAGCAGC<br>AGCAGCAGCAGCAGCAGCAGCAGCAGCAGCAGCAGCAGCAGCAGCAGCAGCAGCAGCAGC<br>AGCAGCAGCAGCAGCAGCACAGCAGCAGCAGCAGCAGCAGCAGCAGCAGCAGCAGCAGCAGCA<br>GCAGCAGCAGCAGCAGCAGCAGCAGCAGCAGCAGCAGCAGCAGCAGCACAGCAGCAGCAG<br>CAGCAGCAGCAGCAGCAGCAGCAGCAGCAGCAGCAGCAGCAGCAGCAGCAGCAGCAGCAG<br>CAGCAGCAGCAGCAGCACAGCAGCAGCAGCAGCAGCAGCAGCAGCAGCAGCAGCAGCAGC<br>AGCAGCAGCAGCAGCAGCAGCAGCAGCAGCAGCAGCAGCAGCAGCACAGCAGCAGCAGCA<br>GCAGCAGCAGCAGCAGCAGCAGCAGCAGCAGCAGCAGCAGCAGCAGCAGCAGCAGCAGCA<br>GCAGCAGCAGCAGCAGCACAGCAGCAGCAGCAGCAGCAGCAGCAGCAGCAGCAGCAGCAG<br>CAGCAGCAGCAGCAGCAGCAGCAGCAGCACAGCAGCAGCAGCAGCAGCAGCAGCAGCAGC<br>AGCAGCAGCAGCAGCAGCAGCAGCAGCAGCAGCAGCAGCAGCAGCAGCAGCAGCAGCACA<br>GCAGCAGCAGCAGCAGCAGCAGCAGCAGCAGCAGCAGCAgagacgGAGCTCgtcgacGCggccgc<br>aagcttggtaccgagctcGGATCCTAAGGTACCTAATTGCCTAGAAAACATGAGGATCACCCATGTCTGC<br>AGGTCGACTCTAGAAAACATGAGGATCACCCATGTCTGCAGTATTCCCGGGTTCATTAGATCCTAA<br>GGTACCTAATTGCCTAGAAAACATGAGGATCACCCATGTCTGCAGGTCGACTCTAGAAAACATGAG<br>GATCACCCATGTCTGCAGTATTCCCGGGTTCATTAGATCCTAAGGTACCTAATTGCCTAGAAAACAT<br>GAGGATCACCCATGTCTGCAGGTCGACTCTAGAAAACATGAGGATCACCCATGTCTGCAGTATTCC<br>CGGGTTCATTAGATCCTAAGGTACCTAATTGCCTAGAAAACATGAGGATCACCCATGTCTGCAGGT<br>CGACTCTAGAAAACATGAGGATCACCCATGTCTGCAGTATTCCCGGGTTCATTAGATCCTAAGGTA<br>CCTAATTGCCTAGAAAACATGAGGATCACCCATGTCTGCAGGTCGACTCTAGAAAACATGAGGATC<br>ACCCATGTCTGCAGTATTCCCGGGTTCATTAGATCCTAAGGTACCTAATTGCCTAGAAAACATGAG<br>GATCACCCATGTCTGCAGGTCGACTCTAGAAAACATGAGGATCACCCATGTCTGCAGTATTCCCGG<br>GTTCATTAGATCTGCGCGCGATCGATATCAGCGCTTTAAATTTGCGCATGCTAGTgTAAggcatgcaag<br>cttgatatcaagcttatcgataatcaacctctggattacaaaatttgtgaaagattgactggtattcttaactatgttgctccttttac<br>gctatgtggatacgctgctttaatgcctttgtatcatgctattgcttcccgtatggctttcattttctcctccttgtataaatcctggttg<br>ctgtctctttatgaggagttgtggcccgttgtcaggcaacgtggcgtggtgtgcactgtgtttgctgacgcaacccccactggttggg<br>gcattgccaccacctgtcagctcctttccgggactttcgctttccccctccctattgccacggcggaactcatcgccgcctgccttgcc<br>cgctgctggacaggggctcggctgttgggcactgacaattccgtggtgttgtcggggaaatcatcgtcctttccttggctgctcgcct<br>gtgttgccacctggattctgcgcgggacgtccttctgctacgtcccttcggccctcaatccagcggaccttccttcccgcggcctgct<br>gccggctctgcggcctcttccgcgtcttcgccttcgccctcagacgagtcggatctccctttgggccgcctccccgcatcgataccgt<br>cgacctcgagggaattaattcgagctcggtacctttaagaccaatgacttacaaggcagctgtagatcttagccactttttaaaag<br>aaaagggggggactggaagggctaattcactcccaacgaagacaagatctgctttttgcttgtactgggtctctctggttagaccag<br>atctgagcctgggagctctctggctaactagggaacccactgcttaagcctcaataaagcttgccttgagtgcttcaagtagtgtgt<br>gcccgtctgttgtgtgactctggtaactagagatccctcagacccttttagtcagtgtggaaaatctctagcagcatctagaattaat<br>tccgtgtatt |
| C4: pHR Tre3G Flank | ctatagtgtcacctaaatcgtatgtgtatgatacataaggttatgtattaattgtagccgcgttctaacgacaatatgtacaagccta<br>attgtgtagcatctggcttactgaagcagaccctatcatctctctcgtaaactgccgtcagagtcggtttggttggacgaaccttctg<br>agtttctggtaacgccgtcccgcacccggaaatggtcagcgaaccaatcagcagggtcatcgctagccagatcctctacgccgga |

| | |
|---|---|
| C4<br>240xCA<br>G<br>12xMS2<br>WPRE | cgcatcgtggccggcatcaccggcgccacaggtgcggttgctggcgcctatatcgccgacatcaccgatggggaagatcgggctc<br>gccacttcgggctcatgagcgcttgtttcggcgtgggtatggtggcaggccccgtggccgggggactgttgggcgccatctccttgc<br>atgcaccattccttgcggcggcggtgctcaacggcctcaacctactactgggctgcttcctaatgcaggagtcgcataagggagag<br>cgtcgaatggtgcactctcagtacaatctgctctgatgccgcatagttaagccagccccgacacccgccaacacccgctgacgcgc<br>cctgacgggcttgtctgctcccggcatccgcttacagacaagctgtgaccgtctccgggagctgcatgtgtcagaggttttcaccgt<br>catcaccgaaacgcgcgagacgaaagggcctcgtgatacgcctatttttataggttaatgtcatgataataatggtttcttagacgt<br>caggtggcacttttcggggaaatgtgcgcggaacccctatttgtttatttttctaaatacattcaaatatgtatccgctcatgagaca<br>ataaccctgataaatgcttcaataatattgaaaaaggaagagtatgagtattcaacatttccgtgtcgcccttattcccttttttgcg<br>gcattttgccttcctgtttttgctcacccagaaacgctggtgaaagtaaaagatgctgaagatcagttgggtgcacgagtgggttac<br>atcgaactggatctcaacagcggtaagatccttgagagttttcgccccgaagaacgttttccaatgatgagcacttttaaagttctg<br>ctatgtggcgcggtattatcccgtattgacgccgggcaagagcaactcggtcgccgcatacactattctcagaatgacttggttga<br>gtactcaccagtcacagaaaagcatcttacggatggcatgacagtaagagaattatgcagtgctgccataaccatgagtgataac<br>actgcggccaacttacttctgacaacgatcggaggaccgaaggagctaaccgcttttttgcacaacatgggggatcatgtaactcg<br>ccttgatcgttgggaaccggagctgaatgaagccataccaaacgacgagcgtgacaccacgatgcctgtagcaatggcaacaac<br>gttgcgcaaactattaactggcgaactacttactctagcttcccggcaacaattaatagactggatggaggcggataaagttgcag<br>gaccacttctgcgctcggcccttccggctggctggtttattgctgataaatctggagccggtgagcgtgggtctcgcggtatcattgc<br>agcactggggccagatggtaagccctcccgtatcgtagttatctacacgacggggagtcaggcaactatggatgaacgaaatag<br>acagatcgctgagataggtgcctcactgattaagcattggtaactgtcagaccaagtttactcatatatactttagattgatttaaa<br>acttcatttttaatttaaaaggatctaggtgaagatcctttttgataatctcatgaccaaaatcccttaacgtgagttttcgttccact<br>gagcgtcagaccccgtagaaaagatcaaaggatcttcttgagatcctttttttctgcgcgtaatctgctgcttgcaaacaaaaaaa<br>ccaccgctaccagcggtggtttgtttgccggatcaagagctaccaactctttttccgaaggtaactggcttcagcagagcgcagat<br>accaaatactgtctttctagtgtagccgtagttaggccaccacttcaagaactctgtagcaccgcctacatacctcgctctgctaatc<br>ctgttaccagtggctgctgccagtggcgataagtcgtgtcttaccgggttggactcaagacgatagttaccggataaggcgcagcg<br>gtcgggctgaacggggggttcgtgcacacagcccagcttggagcgaacgacctacaccgaactgagatacctacagcgtgagct<br>atgagaaagcgccacgcttcccgaagggagaaaggcggacaggtatccggtaagcggcagggtcggaacaggagagcgcacg<br>agggagcttccagggggaaacgcctggtatctttatagtcctgtcgggtttcgccacctctgacttgagcgtcgatttttgtgatgct<br>cgtcaggggggcggagcctatggaaaaacgccagcaacgcggcctttttacggttcctggccttttgctggccttttgctcacatgt<br>tctttcctgcgttatcccctgattctgtggataaccgtattaccgcctttgagtgagctgataccgctcgccgcagccgaacgaccga<br>gcgcagcgagtcagtgagcgaggaagcggaagagcgcccaatacgcaaaccgcctctccccgcgcgttggccgattcattaatg<br>cagctgtggaatgtgtgtcagttagggtgtggaaagtccccaggctccccagcaggcagaagtatgcaaagcatgcatctcaatt<br>agtcagcaaccaggtgtggaaagtccccaggctccccagcaggcagaagtatgcaaagcatgcatctcaattagtcagcaacca<br>tagtcccgcccctaactccgcccatcccgcccctaactccgcccagttccgcccattctccgccccatggctgactaattttttttattt<br>atgcagaggccgaggccgcctcggcctctgagctattccagaagtagtgaggaggcttttttggaggcctaggcttttgcaaaaag<br>cttggacacaagacaggcttgcgagatatgtttgagaataccactttatcccgcgtcagggagaggcagtgcgtaaaaagacgc<br>ggactcatgtgaaatactggtttttagtgcgccagatctctataatctcgcgcaacctattttcccctcgaacactttttaagccgtag<br>ataaacaggctgggacacttcacatgagcgaaaaatacatcgtcacctgggacatgttgcagatccatgcacgtaaactcgcaa<br>gccgactgatgccttctgaacaatggaaaggcattattgccgtaagccgtggcggtctgtaccgggtgcgttactggcgcgtgaac<br>tgggtattcgtcatgtcgataccgtttgtatttccagctacgatcacgacaaccagcgcgagcttaaagtgctgaaacgcgcagaa<br>ggcgatggcgaaggcttcatcgttattgatgacctggtggataccggtggtactgcggttgcgattcgtgaaatgtatccaaaagc<br>gcactttgtcaccatcttcgcaaaaccggctggtcgtccgctggttgatgactatgttgttgatatcccgcaagatacctggattgaa<br>cagccgtgggatatgggcgtcgtattcgtcccgccaatctccggtcgctaatcttttcaacgcctggcactgccgggcgttgttctttt<br>taacttcaggcgggttacaatagtttccagtaagtattctggaggctgcatccatgacacaggcaaacctgagcgaaaccctgttc<br>aaaccccgctttaaacatcctgaaacctcgacgctagtccgccgctttaatcacggcgcacaaccgcctgtgcagtcggcccttga<br>tggtaaaaccatccctcactggtatcgcatgattaaccgtctgatgtggatctggcgcggcattgacccacgcgaaatcctcgacg<br>tccaggcacgtattgtgatgagcgatgccgaacgtaccgacgatgatttatacgatacggtgattggctaccgtggcggcaactgg<br>atttatgagtgggccccggatctttgtgaaggaaccttacttctgtggtgtgacataattggacaaactacctacagagatttaaag<br>ctctaaggtaaatataaaatttttaagtgtataatgtgttaaactactgattctaattgtttgtgtattttagattccaacctatggaa<br>ctgatgaatgggagcagtggtggaatgcctttaatgaggaaaacctgttttgctcagaagaaatgccatctagtgatgatgaggct |

| | |
|---|---|
| | actgctgactctcaacattctactcctccaaaaaagaagagaaaggtagaagaccccaaggactttccttcagaattgctaagttt<br>tttgagtcatgctgtgtttagtaatagaactcttgcttgctttgctatttacaccacaaaggaaaaagctgcactgctatacaagaa<br>aattatggaaaaatattctgtaacctttataagtaggcataacagttataatcataacatactgtttttcttactccacacaggcat<br>agagtgtctgctattaataactatgctcaaaaattgtgtacctttagctttttaatttgtaaaggggttaataaggaatatttgatgta<br>tagtgccttgactagagatcataatcagccataccacatttgtagaggttttacttgctttaaaaaacctcccacacctccccctga<br>acctgaaacataaaatgaatgcaattgttgttgttaacttgtttattgcagcttataatggttacaaataaagcaatagcatcacaa<br>atttcacaaataaagcatttttttcactgcattctagttgtggtttgtccaaactcatcaatgtatcttatcatgtctggatcaactgga<br>taactcaagctaaccaaaatcatcccaaacttcccaccccataccctattaccactgccaattacctagtggtttcatttactctaaa<br>cctgtgattcctctgaattattttcattttaaagaaattgtatttgttaaatatgtactacaaacttagtagttggaagggctaattca<br>ctcccaaagaagacaagatatccttgatctgtggatctaccacacacaaggctacttccctgattagcagaactacacaccaggg<br>ccaggggtcagatatccactgacctttggatggtgctacaagctagtaccagttgagccagataaggtagaagaggccaataaa<br>ggagagaacaccagcttgttacaccctgtgagcctgcatgggatggatgacccggagagagaagtgttagagtggaggtttgac<br>agccgcctagcatttcatcacgtggcccgagagctgcatccggagtacttcaagaactgctgatatcgagcttgctacaagggact<br>ttccgctggggactttccagggaggcgtggcctgggcgggactggggagtggcgagccctcagatcctgcatataagcagctgct<br>ttttgcctgtactgggtctctctggttagaccagatctgagcctgggagctctctggctaactagggaacccactgcttaagcctcaa<br>taaagcttgccttgagtgcttcaagtagtgtgtgcccgtctgttgtgtgactctggtaactagagatccctcagacccttttagtcagt<br>gtggaaaatctctagcagtggcgcccgaacagggacttgaaagcgaaagggaaaccagaggagctctctcgacgcaggactcg<br>gcttgctgaagcgcgcacggcaagaggcgaggggcggcgactggtgagtacgccaaaaattttgactagcggaggctagaagg<br>agagagatgggtgcgagagcgtcagtattaagcgggggagaattagatcgcgatgggaaaaaattcggttaaggccagggggа<br>aagaaaaaatataaattaaaacatatagtatgggcaagcagggagctagaacgattcgcagttaatcctggcctgttagaaaca<br>tcagaaggctgtagacaaatactgggacagctacaaccatcccttcagacaggatcagaagaacttagatcattatataatacag<br>tagcaaccctctattgtgtgcatcaaaggatagagataaaagacaccaaggaagctttagacaagatagaggaagagcaaaac<br>aaaagtaagaccaccgcacagcaagcggccggccgctgatcttcagacctggaggaggagatatgagggacaattggagaagt<br>gaattatataaatataaagtagtaaaaattgaaccattaggagtagcacccaccaaggcaaagagaagagtggtgcagagaga<br>aaaaagagcagtgggaataggagctttgttccttgggttcttgggagcagcaggaagcactatgggcgcagcgtcaatgacgctg<br>acggtacaggccagacaattattgtctggtatagtgcagcagcagaacaatttgctgagggctattgaggcgcaacagcatctgtt<br>gcaactcacagtctggggcatcaagcagctccaggcaagaatcctggctgtgaaagatacctaaaggatcaacagctcctgggg<br>atttggggttgctctggaaaactcatttgcaccactgctgtgccttggaatgctagttggagtaataaatctctggaacagatttgg<br>aatcacacgacctggatggagtgggacagagaaattaacaattacacaagcttaatacactccttaattgaagaatcgcaaaac<br>cagcaagaaaagaatgaacaagaattattggaattagataaatgggcaagtttgtggaattggtttaacataacaaattggctgt<br>ggtatataaaattattcataatgatagtaggaggcttggtaggtttaagaatagtttttgctgtactttctatagtgaatagagttag<br>gcagggatattcaccattatcgtttcagacccacctcccaaccccgaggggacccgacaggcccgaaggaatagaagaagaag<br>gtggagagagagacagagacagatccattcgattagtgaacggatctcgacggtatcgccaaatggcagtattcatccacaattt<br>taaaagaaaaggggggattggggggtacagtgcaggggaaagaatagtagacataatagcaacagacatacaaactaaagaa<br>ttacaaaaacaaattacaaaaattcaaaattttcgggtttattacagggacagcagagatccagtttatcgatgaggccctttcgt<br>cttcactcgagtttactccctatcagtgatagagaacgtatgaagagtttactccctatcagtgatagagaacgtatgcagacttta<br>ctccctatcagtgatagagaacgtataaggagtttactccctatcagtgatagagaacgtatgaccagtttactccctatcagtgat<br>agagaacgtatctacagtttactccctatcagtgatagagaacgtatatccagtttactccctatcagtgatagagaacgtatgtcg<br>aggtaggcgtgtacggtgggcgcctataaaagcagagctcgtttagtgaaccgtcagatcgcctggagcaattccacaacactttt<br>gtcttatacttACGCGTGTCGACcatatgACACGGGGTACCGGCCGTCGGACGCCGCGTTCGCCCAACTG<br>GCACAGTACCCCGGCACTGCCCGCGCAGGCCTCGCGCGCGCCAGGACCTGCCCTGCGCGAGCGGG<br>TACCGCGTACGATGCGGCCTCGGCGCCCAAGCGCGCACGGGCCCGTGGCCCAATTCCAGCGTGGG<br>GTGAGCGAGAGGCGGGACCCGCCGCTCTGCCTAGGCCACCTCGACGCGCAAACGTTGGTGGTCG<br>CGGCTACCTCCGTCCCTatgGGTAATTATAACCCGGGCCCTATATATTAGGCTAACTAGCTAACTAG<br>GAATTCggtctcaCAGCAGCAGCAGCAGCAGCAGCAGCAGCAGCAGCAGCAGCACAGCAGCAG<br>CAGCAGCAGCAGCAGCAGCAGCAGCAGCAGCAGCAGCAGCAGCAGCAGCAGCAGCAGCAG<br>CAGCAGCAGCAGCAGCAGCAGCACAGCAGCAGCAGCAGCAGCAGCAGCAGCAGCAGCAGCAGC<br>AGCAGCAGCAGCAGCAGCAGCAGCAGCAGCAGCAGCAGCAGCAGCAGCAGCAGCACAGCAGCA |

| | |
|---|---|
| | GCAGCAGCAGCAGCAGCAGCAGCAGCAGCAGCAGCAGCAGCAGCAGCAGCAGCAGCAGCA<br>GCAGCAGCAGCAGCAGCACAGCAGCAGCAGCAGCAGCAGCAGCAGCAGCAGCAGCAG<br>CAGCAGCAGCAGCAGCAGCAGCAGCAGCAGCAGCAGCAGCAGCAGCACAGCAGCAGC<br>AGCAGCAGCAGCAGCAGCAGCAGCAGCAGCAGCAGCAGCAGCAGCAGCAGCAGCAGC<br>AGCAGCAGCAGCAGCAGCACAGCAGCAGCAGCAGCAGCAGCAGCAGCAGCAGCAGCA<br>GCAGCAGCAGCAGCAGCAGCAGCAGCAGCACAGCAGCAGCAGCAGCAGCAGCAGCAG<br>CAGCAGCAGCAGCAGCAGCAGCAGCAGCAGCAGCAGCAGCAGCAGCAGCAGCAGCAG<br>CACAGCAGCAGCAGCAGCAGCAGCAGCAGCAGCAGCAGCAGCAGCAgagacgGAGCTCgtcgacGC<br>ggccgcaagcttggtaccgagctcGGATCCTAAGGTACCTAATTGCCTAGAAAACATGAGGATCACCCATG<br>TCTGCAGGTCGACTCTAGAAAACATGAGGATCACCCATGTCTGCAGTATTCCCGGGTTCATTAGAT<br>CCTAAGGTACCTAATTGCCTAGAAAACATGAGGATCACCCATGTCTGCAGGTCGACTCTAGAAAAC<br>ATGAGGATCACCCATGTCTGCAGTATTCCCGGGTTCATTAGATCCTAAGGTACCTAATTGCCTAGAA<br>AACATGAGGATCACCCATGTCTGCAGGTCGACTCTAGAAAACATGAGGATCACCCATGTCTGCAGT<br>ATTCCCGGGTTCATTAGATCCTAAGGTACCTAATTGCCTAGAAAACATGAGGATCACCCATGTCTGC<br>AGGTCGACTCTAGAAAACATGAGGATCACCCATGTCTGCAGTATTCCCGGGTTCATTAGATCCTAA<br>GGTACCTAATTGCCTAGAAAACATGAGGATCACCCATGTCTGCAGGTCGACTCTAGAAAACATGAG<br>GATCACCCATGTCTGCAGTATTCCCGGGTTCATTAGATCCTAAGGTACCTAATTGCCTAGAAAACAT<br>GAGGATCACCCATGTCTGCAGGTCGACTCTAGAAAACATGAGGATCACCCATGTCTGCAGTATTCC<br>CGGGTTCATTAGATCTGCGCGCGATCGATATCAGCGCTTTAAATTTGCGCATGCTAGTgTAAggcatg<br>caagcttgatatcaagcttatcgataatcaacctctggattacaaaatttgtgaaagattgactggtattcttaactatgttgctcctt<br>ttacgctatgtggatacgctgctttaatgcctttgtatcatgctattgcttcccgtatggctttcattttctcctccttgtataaatcctg<br>gttgctgtctctttatgaggagttgtggcccgttgtcaggcaacgtggcgtggtgtgcactgtgtttgctgacgcaacccccactggt<br>tggggcattgccaccacctgtcagctcctttccgggactttcgctttccccctccctattgccacggcggaactcatcgccgcctgcc<br>ttgcccgctgctggacaggggctcggctgttgggcactgacaattccgtggtgttgtcggggaaatcatcgtcctttccttggctgct<br>cgcctgtgttgccacctggattctgcgcgggacgtccttctgctacgtcccttcggccctcaatccagcggaccttccttcccgcggc<br>ctgctgccggctctgcggcctcttccgcgtcttcgccttcgccctcagacgagtcggatctccctttgggccgcctccccgcatcgat<br>accgtcgacctcgagggaattaattcgagctcggtacctttaagaccaatgacttacaaggcagctgtagatcttagccactttttta<br>aaagaaaaggggggactggaagggctaattcactcccaacgaagacaagatctgctttttgcttgtactgggtctctctggttaga<br>ccagatctgagcctgggagctctctggctaactagggaacccactgcttaagcctcaataaagcttgccttgagtgcttcaagtagt<br>gtgtgcccgtctgttgtgtgactctggtaactagagatccctcagacccttttagtcagtgtggaaaatctctagcagcatctagaat<br>taattccgtgtatt |
| C5: pHR Tre3G Flank C5 240xCAG 12xMS2 WPRE | ctatagtgtcacctaaatcgtatgtgtatgatacataaggttatgtattaattgtagccgcgttctaacgacaatatgtacaagccta<br>attgtgtagcatctggcttactgaagcagaccctatcatctctctcgtaaactgccgtcagagtcggtttggttggacgaaccttctg<br>agtttctggtaacgccgtcccgcacccggaaatggtcagcgaaccaatcagcagggtcatcgctagccagatcctctacgccgga<br>cgcatcgtggccggcatcaccggcgccacaggtgcggttgctggcgcctatatcgccgacatcaccgatggggaagatcgggctc<br>gccacttcgggctcatgagcgcttgtttcggcgtgggtatggtggcaggccccgtggccgggggactgttgggcgccatctccttgc<br>atgcaccattccttgcggcggcggtgctcaacggcctcaacctactactgggctgcttcctaatgcaggagtcgcataagggagag<br>cgtcgaatggtgcactctcagtacaatctgctctgatgccgcatagttaagccagccccgacacccgccaacacccgctgacgcgc<br>cctgacgggcttgtctgctcccggcatccgcttacagacaagctgtgaccgtctccgggagctgcatgtgtcagaggttttcaccgt<br>catcaccgaaacgcgcgagacgaaagggcctcgtgatacgcctatttttataggttaatgtcatgataataatggtttcttagacgt<br>caggtggcacttttcggggaaatgtgcgcggaacccctatttgtttatttttctaaatacattcaaatatgtatccgctcatgagaca<br>ataaccctgataaatgcttcaataatattgaaaaaggaagagtatgagtattcaacatttccgtgtcgcccttattcccttttttgcg<br>gcattttgccttcctgtttttgctcacccagaaacgctggtgaaagtaaaagatgctgaagatcagttgggtgcacgagtgggttac<br>atcgaactggatctcaacagcggtaagatccttgagagttttcgccccgaagaacgttttccaatgatgagcacttttaaagttctg<br>ctatgtggcgcggtattatcccgtattgacgccgggcaagagcaactcggtcgccgcatacactattctcagaatgacttggttga<br>gtactcaccagtcacagaaaagcatcttacggatggcatgacagtaagagaattatgcagtgctgccataaccatgagtgataac<br>actgcggccaacttacttctgacaacgatcggaggaccgaaggagctaaccgcttttttgcacaacatgggggatcatgtaactcg |

| | |
|---|---|
| | ccttgatcgttgggaaccggagctgaatgaagccataccaaacgacgagcgtgacaccacgatgcctgtagcaatggcaacaac<br>gttgcgcaaactattaactggcgaactacttactctagcttcccggcaacaattaatagactggatggaggcggataaagttgcag<br>gaccacttctgcgctcggcccttccggctggctggtttattgctgataaatctggagccggtgagcgtgggtctcgcggtatcattgc<br>agcactggggccagatggtaagccctcccgtatcgtagttatctacacgacggggagtcaggcaactatggatgaacgaaatag<br>acagatcgctgagataggtgcctcactgattaagcattggtaactgtcagaccaagtttactcatatatactttagattgatttaaa<br>acttcatttttaatttaaaaggatctaggtgaagatcctttttgataatctcatgaccaaaatcccttaacgtgagttttcgttccact<br>gagcgtcagaccccgtagaaaagatcaaaggatcttcttgagatcctttttttctgcgcgtaatctgctgcttgcaaacaaaaaaa<br>ccaccgctaccagcggtggtttgtttgccggatcaagagctaccaactctttttccgaaggtaactggcttcagcagagcgcagat<br>accaaatactgtctttctagtgtagccgtagttaggccaccacttcaagaactctgtagcaccgcctacatacctcgctctgctaatc<br>ctgttaccagtggctgctgccagtggcgataagtcgtgtcttaccgggttggactcaagacgatagttaccggataaggcgcagcg<br>gtcgggctgaacggggggttcgtgcacacagcccagcttggagcgaacgacctacaccgaactgagatacctacagcgtgagct<br>atgagaaagcgccacgcttcccgaagggagaaaggcggacaggtatccggtaagcggcagggtcggaacaggagagcgcacg<br>agggagcttccagggggaaacgcctggtatctttatagtcctgtcgggtttcgccacctctgacttgagcgtcgatttttgtgatgct<br>cgtcaggggggcggagcctatggaaaaacgccagcaacgcggcctttttacggttcctggccttttgctggccttttgctcacatgt<br>tctttcctgcgttatcccctgattctgtggataaccgtattaccgcctttgagtgagctgataccgctcgccgcagccgaacgaccga<br>gcgcagcgagtcagtgagcgaggaagcggaagagcgcccaatacgcaaaccgcctctccccgcgcgttggccgattcattaatg<br>cagctgtggaatgtgtgtcagttagggtgtggaaagtccccaggctccccagcaggcagaagtatgcaaagcatgcatctcaatt<br>agtcagcaaccaggtgtggaaagtccccaggctccccagcaggcagaagtatgcaaagcatgcatctcaattagtcagcaacca<br>tagtcccgcccctaactccgcccatcccgcccctaactccgcccagttccgcccattctccgccccatggctgactaattttttttattt<br>atgcagaggccgaggccgcctcggcctctgagctattccagaagtagtgaggaggcttttttggaggcctaggcttttgcaaaaag<br>cttggacacaagacaggcttgcgagatatgtttgagaataccactttatcccgcgtcagggagaggcagtgcgtaaaaagacgc<br>ggactcatgtgaaatactggtttttagtgcgccagatctctataatctcgcgcaacctattttcccctcgaacactttttaagccgtag<br>ataaacaggctgggacacttcacatgagcgaaaaatacatcgtcacctgggacatgttgcagatccatgcacgtaaactcgcaa<br>gccgactgatgccttctgaacaatggaaaggcattattgccgtaagccgtggcggtctgtaccgggtgcgttactggcgcgtgaac<br>tgggtattcgtcatgtcgataccgtttgtatttccagctacgatcacgacaaccagcgcgagcttaaagtgctgaaacgcgcagaa<br>ggcgatggcgaaggcttcatcgttattgatgacctggtggataccggtggtactgcggttgcgattcgtgaaatgtatccaaaagc<br>gcactttgtcaccatcttcgcaaaaccggctggtcgtccgctggttgatgactatgttgttgatatcccgcaagatacctggattgaa<br>cagccgtgggatatgggcgtcgtattcgtcccgccaatctccggtcgctaatcttttcaacgcctggcactgccgggcgttgttctttt<br>taacttcaggcgggttacaatagtttccagtaagtattctggaggctgcatccatgacacaggcaaacctgagcgaaaccctgttc<br>aaaccccgctttaaacatcctgaaacctcgacgctagtccgccgctttaatcacggcgcacaaccgcctgtgcagtcggcccttga<br>tggtaaaaccatccctcactggtatcgcatgattaaccgtctgatgtggatctggcgcggcattgacccacgcgaaatcctcgacg<br>tccaggcacgtattgtgatgagcgatgccgaacgtaccgacgatgatttatacgatacggtgattggctaccgtggcggcaactgg<br>atttatgagtgggccccggatctttgtgaaggaaccttacttctgtggtgtgacataattggacaaactacctacagagatttaaag<br>ctctaaggtaaatataaaatttttaagtgtataatgtgttaaactactgattctaattgtttgtgtattttagattccaacctatggaa<br>ctgatgaatgggagcagtggtggaatgcctttaatgaggaaaacctgttttgctcagaagaaatgccatctagtgatgatgaggct<br>actgctgactctcaacattctactcctccaaaaaagaagagaaaggtagaagaccccaaggactttccttcagaattgctaagttt<br>tttgagtcatgctgtgtttagtaatagaactcttgcttgctttgctatttacaccacaaaggaaaaagctgcactgctatacaagaa<br>aattatggaaaaatattctgtaacctttataagtaggcataacagttataatcataacatactgttttttcttactccacacaggcat<br>agagtgtctgctattaataactatgctcaaaaattgtgtacctttagctttttaatttgtaaaggggttaataaggaatatttgatgta<br>tagtgccttgactagagatcataatcagccataccacatttgtagaggttttacttgctttaaaaaacctcccacacctccccctga<br>acctgaaacataaaatgaatgcaattgttgttgttaacttgtttattgcagcttataatggttacaaataaagcaatagcatcacaa<br>atttcacaaataaagcatttttttcactgcattctagttgtggtttgtccaaactcatcaatgtatcttatcatgtctggatcaactgga<br>taactcaagctaaccaaaatcatcccaaacttcccaccccataccctattaccactgccaattacctagtggtttcatttactctaaa<br>cctgtgattcctctgaattattttcattttaaagaaattgtatttgttaaatatgtactacaaacttagtagttggaagggctaattca<br>ctcccaaagaagacaagatatccttgatctgtggatctaccacacacaaggctacttccctgattagcagaactacacaccaggg<br>ccaggggtcagatatccactgacctttggatggtgctacaagctagtaccagttgagccagataaggtagaagaggccaataaa<br>ggagagaacaccagcttgttacaccctgtgagcctgcatgggatggatgacccggagagagaagtgttagagtggaggtttgac<br>agccgcctagcatttcatcacgtggcccgagagctgcatccggagtacttcaagaactgctgatatcgagcttgctacaagggact |

| | |
|---|---|
| | ttccgctggggactttccagggaggcgtggcctgggcgggactggggagtggcgagccctcagatcctgcatataagcagctgct<br>ttttgcctgtactgggtctctctggttagaccagatctgagcctgggagctctctggctaactagggaacccactgcttaagcctcaa<br>taaagcttgccttgagtgcttcaagtagtgtgtgcccgtctgttgtgtgactctggtaactagagatccctcagacccttttagtcagt<br>gtggaaaatctctagcagtggcgcccgaacagggacttgaaagcgaaagggaaaccagaggagctctctcgacgcaggactcg<br>gcttgctgaagcgcgcacggcaagaggcgaggggcggcgactggtgagtacgccaaaaattttgactagcggaggctagaagg<br>agagagatgggtgcgagagcgtcagtattaagcgggggagaattagatcgcgatgggaaaaaattcggttaaggccaggggga<br>aagaaaaaatataaattaaaacatatagtatgggcaagcagggagctagaacgattcgcagttaatcctggcctgttagaaaca<br>tcagaaggctgtagacaaatactgggacagctacaaccatcccttcagacaggatcagaagaacttagatcattatataatacag<br>tagcaaccctctattgtgtgcatcaaaggatagagataaaagacaccaaggaagctttagacaagatagaggaagagcaaaac<br>aaaagtaagaccaccgcacagcaagcggccggccgctgatcttcagacctggaggaggagatatgagggacaattggagaagt<br>gaattatataaatataaagtagtaaaaattgaaccattaggagtagcacccaccaaggcaaagagaagagtggtgcagagaga<br>aaaaagagcagtgggaataggagctttgttccttgggttcttgggagcagcaggaagcactatgggcgcagcgtcaatgacgctg<br>acggtacaggccagacaattattgtctggtatagtgcagcagcagaacaatttgctgagggctattgaggcgcaacagcatctgtt<br>gcaactcacagtctggggcatcaagcagctccaggcaagaatcctggctgtgaaagatacctaaaggatcaacagctcctgggg<br>atttggggttgctctggaaaactcatttgcaccactgctgtgccttggaatgctagttggagtaataaatctctggaacagatttgg<br>aatcacacgacctggatggagtgggacagagaaattaacaattacacaagcttaatacactccttaattgaagaatcgcaaaac<br>cagcaagaaaagaatgaacaagaattattggaattagataaatgggcaagtttgtggaattggtttaacataacaaattggctgt<br>ggtatataaaattattcataatgatagtaggaggcttggtaggtttaagaatagtttttgctgtactttctatagtgaatagagttag<br>gcagggatattcaccattatcgtttcagacccacctcccaaccccgaggggacccgacaggcccgaaggaatagaagaagaag<br>gtggagagagagacagagacagatccattcgattagtgaacggatctcgacggtatcgccaaatggcagtattcatccacaattt<br>taaaagaaaaggggggattggggggtacagtgcaggggaaagaatagtagacataatagcaacagacatacaaactaaagaa<br>ttacaaaaacaaattacaaaaattcaaaattttcgggtttattacagggacagcagagatccagtttatcgatgaggccctttcgt<br>cttcactcgagtttactccctatcagtgatagagaacgtatgaagagtttactccctatcagtgatagagaacgtatgcagacttta<br>ctccctatcagtgatagagaacgtataaggagtttactccctatcagtgatagagaacgtatgaccagtttactccctatcagtgat<br>agagaacgtatctacagtttactccctatcagtgatagagaacgtatatccagtttactccctatcagtgatagagaacgtatgtcg<br>aggtaggcgtgtacggtgggcgcctataaaagcagagctcgtttagtgaaccgtcagatcgcctggagcaattccacaacactttt<br>gtcttatacttACGCGTGTCGACcatatgATGGCACCGCGAGGTGGCACCCCGGCGCCGAGACCCTCGTG<br>TCTCAGTCCTGGCCCGCAACACGCCCTGCAAGTGCGCGCGATCAGGAACCGACCCCCGCGAGCCC<br>CGCTGAGCCCGTACTCCGGCCCACCCCGCGTGGCCCCGTCCCCTATCGCGGTGGGTGGGCGCCAC<br>AGGCGAGTCCAGGGGGGTGGTGGGCTGTTCGGCTGGCCCGGGCCTTGGTGGGGCCGAGGGAGC<br>CGGCTGATGGTCGAACCTatgGGTAATTATAACCCGGGCCCTATATATTAGGCTAACTAGCTAACTA<br>GGAATTCggtctcaCAGCAGCAGCAGCAGCAGCAGCAGCAGCAGCAGCAGCAGCAGCACAGCAGCA<br>GCAGCAGCAGCAGCAGCAGCAGCAGCAGCAGCAGCAGCAGCAGCAGCAGCAGCAGCAGCAGCA<br>GCAGCAGCAGCAGCAGCAGCAGCACAGCAGCAGCAGCAGCAGCAGCAGCAGCAGCAGCAGCAG<br>CAGCAGCAGCAGCAGCAGCAGCAGCAGCAGCAGCAGCAGCAGCAGCAGCAGCAGCACAGCAGC<br>AGCAGCAGCAGCAGCAGCAGCAGCAGCAGCAGCAGCAGCAGCAGCAGCAGCAGCAGCAGCAGC<br>AGCAGCAGCAGCAGCAGCAGCACAGCAGCAGCAGCAGCAGCAGCAGCAGCAGCAGCAGCAGCA<br>GCAGCAGCAGCAGCAGCAGCAGCAGCAGCAGCAGCAGCAGCAGCAGCAGCAGCACAGCAGCAG<br>CAGCAGCAGCAGCAGCAGCAGCAGCAGCAGCAGCAGCAGCAGCAGCAGCAGCAGCAGCAGCAG<br>CAGCAGCAGCAGCAGCAGCAGCACAGCAGCAGCAGCAGCAGCAGCAGCAGCAGCAGCAGCAGC<br>AGCAGCAGCAGCAGCAGCAGCAGCAGCAGCAGCACAGCAGCAGCAGCAGCAGCAGCAGCAGCA<br>GCAGCAGCAGCAGCAGCAGCAGCAGCAGCAGCAGCAGCAGCAGCAGCAGCAGCAGCAGCAGCA<br>GCACAGCAGCAGCAGCAGCAGCAGCAGCAGCAGCAGCAGCAGCAGCAgagacgGAGCTCgtcgacG<br>CggccgcaagcttggtaccgagctcGGATCCTAAGGTACCTAATTGCCTAGAAAACATGAGGATCACCCAT<br>GTCTGCAGGTCGACTCTAGAAAACATGAGGATCACCCATGTCTGCAGTATTCCCGGGTTCATTAGA<br>TCCTAAGGTACCTAATTGCCTAGAAAACATGAGGATCACCCATGTCTGCAGGTCGACTCTAGAAAA<br>CATGAGGATCACCCATGTCTGCAGTATTCCCGGGTTCATTAGATCCTAAGGTACCTAATTGCCTAGA<br>AAACATGAGGATCACCCATGTCTGCAGGTCGACTCTAGAAAACATGAGGATCACCCATGTCTGCAG |

| | |
|---|---|
| | TATTCCCGGGTTCATTAGATCCTAAGGTACCTAATTGCCTAGAAAACATGAGGATCACCCATGTCTG<br>CAGGTCGACTCTAGAAAACATGAGGATCACCCATGTCTGCAGTATTCCCGGGTTCATTAGATCCTA<br>AGGTACCTAATTGCCTAGAAAACATGAGGATCACCCATGTCTGCAGGTCGACTCTAGAAAACATGA<br>GGATCACCCATGTCTGCAGTATTCCCGGGTTCATTAGATCCTAAGGTACCTAATTGCCTAGAAAACA<br>TGAGGATCACCCATGTCTGCAGGTCGACTCTAGAAAACATGAGGATCACCCATGTCTGCAGTATTC<br>CCGGGTTCATTAGATCTGCGCGCGATCGATATCAGCGCTTTAAATTTGCGCATGCTAGTgTAAggcat<br>gcaagcttgatatcaagcttatcgataatcaacctctggattacaaaatttgtgaaagattgactggtattcttaactatgttgctcc<br>ttttacgctatgtggatacgctgctttaatgcctttgtatcatgctattgcttcccgtatggctttcattttctcctccttgtataaatcct<br>ggttgctgtctctttatgaggagttgtggcccgttgtcaggcaacgtggcgtggtgtgcactgtgtttgctgacgcaacccccactgg<br>ttggggcattgccaccacctgtcagctcctttccgggactttcgctttccccctccctattgccacggcggaactcatcgccgcctgc<br>cttgcccgctgctggacaggggctcggctgttgggcactgacaattccgtggtgttgtcggggaaatcatcgtcctttccttggctgc<br>tcgcctgtgttgccacctggattctgcgcgggacgtccttctgctacgtcccttcggccctcaatccagcggaccttccttcccgcgg<br>cctgctgccggctctgcggcctcttccgcgtcttcgccttcgccctcagacgagtcggatctccctttgggccgcctccccgcatcga<br>taccgtcgacctcgagggaattaattcgagctcggtacctttaagaccaatgacttacaaggcagctgtagatcttagccactttttt<br>aaaagaaaaggggggactggaagggctaattcactcccaacgaagacaagatctgctttttgcttgtactgggtctctctggttag<br>accagatctgagcctgggagctctctggctaactagggaacccactgcttaagcctcaataaagcttgccttgagtgcttcaagta<br>gtgtgtgcccgtctgttgtgtgactctggtaactagagatccctcagacccttttagtcagtgtggaaaatctctagcagcatctaga<br>attaattccgtgtatt |
| Delta MS2: pHR Tre3G Flank C1 240xCAG WPRE | ctatagtgtcacctaaatcgtatgtgtatgatacataaggttatgtattaattgtagccgcgttctaacgacaatatgtacaagccta<br>attgtgtagcatctggcttactgaagcagaccctatcatctctctcgtaaactgccgtcagagtcggtttggttggacgaaccttctg<br>agtttctggtaacgccgtcccgcacccggaaatggtcagcgaaccaatcagcagggtcatcgctagccagatcctctacgccgga<br>cgcatcgtggccggcatcaccggcgccacaggtgcggttgctggcgcctatatcgccgacatcaccgatggggaagatcgggctc<br>gccacttcgggctcatgagcgcttgtttcggcgtgggtatggtggcaggccccgtggccgggggactgttgggcgccatctccttgc<br>atgcaccattccttgcggcggcggtgctcaacggcctcaacctactactgggctgcttcctaatgcaggagtcgcataagggagag<br>cgtcgaatggtgcactctcagtacaatctgctctgatgccgcatagttaagccagccccgacacccgccaacacccgctgacgcgc<br>cctgacgggcttgtctgctcccggcatccgcttacagacaagctgtgaccgtctccgggagctgcatgtgtcagaggttttcaccgt<br>catcaccgaaacgcgcgagacgaaagggcctcgtgatacgcctatttttataggttaatgtcatgataataatggtttcttagacgt<br>caggtggcacttttcggggaaatgtgcgcggaacccctatttgtttatttttctaaatacattcaaatatgtatccgctcatgagaca<br>ataaccctgataaatgcttcaataatattgaaaaaggaagagtatgagtattcaacatttccgtgtcgcccttattcccttttttgcg<br>gcattttgccttcctgtttttgctcacccagaaacgctggtgaaagtaaaagatgctgaagatcagttgggtgcacgagtgggttac<br>atcgaactggatctcaacagcggtaagatccttgagagttttcgccccgaagaacgttttccaatgatgagcacttttaaagttctg<br>ctatgtggcgcggtattatcccgtattgacgccgggcaagagcaactcggtcgccgcatacactattctcagaatgacttggttga<br>gtactcaccagtcacagaaaagcatcttacggatggcatgacagtaagagaattatgcagtgctgccataaccatgagtgataac<br>actgcggccaacttacttctgacaacgatcggaggaccgaaggagctaaccgcttttttgcacaacatgggggatcatgtaactcg<br>ccttgatcgttgggaaccggagctgaatgaagccataccaaacgacgagcgtgacaccacgatgcctgtagcaatggcaacaac<br>gttgcgcaaactattaactggcgaactacttactctagcttcccggcaacaattaatagactggatggaggcggataaagttgcag<br>gaccacttctgcgctcggcccttccggctggctggtttattgctgataaatctggagccggtgagcgtgggtctcgcggtatcattgc<br>agcactggggccagatggtaagccctcccgtatcgtagttatctacacgacggggagtcaggcaactatggatgaacgaaatag<br>acagatcgctgagataggtgcctcactgattaagcattggtaactgtcagaccaagtttactcatatatactttagattgatttaaa<br>acttcatttttaatttaaaaggatctaggtgaagatcctttttgataatctcatgaccaaaatcccttaacgtgagttttcgttccact<br>gagcgtcagaccccgtagaaaagatcaaaggatcttcttgagatcctttttttctgcgcgtaatctgctgcttgcaaacaaaaaaa<br>ccaccgctaccagcggtggtttgtttgccggatcaagagctaccaactctttttccgaaggtaactggcttcagcagagcgcagat<br>accaaatactgtctttctagtgtagccgtagttaggccaccacttcaagaactctgtagcaccgcctacatacctcgctctgctaatc<br>ctgttaccagtggctgctgccagtggcgataagtcgtgtcttaccgggttggactcaagacgatagttaccggataaggcgcagcg<br>gtcgggctgaacggggggttcgtgcacacagcccagcttggagcgaacgacctacaccgaactgagatacctacagcgtgagct<br>atgagaaagcgccacgcttcccgaagggagaaaggcggacaggtatccggtaagcggcagggtcggaacaggagagcgcacg<br>agggagcttccagggggaaacgcctggtatctttatagtcctgtcgggtttcgccacctctgacttgagcgtcgatttttgtgatgct |

cgtcaggggggcggagcctatggaaaaacgccagcaacgcggcctttttacggttcctggccttttgctggccttttgctcacatgt
tctttcctgcgttatcccctgattctgtggataaccgtattaccgcctttgagtgagctgataccgctcgccgcagccgaacgaccga
gcgcagcgagtcagtgagcgaggaagcggaagagcgcccaatacgcaaaccgcctctccccgcgcgttggccgattcattaatg
cagctgtggaatgtgtgtcagttagggtgtggaaagtccccaggctccccagcaggcagaagtatgcaaagcatgcatctcaatt
agtcagcaaccaggtgtggaaagtccccaggctccccagcaggcagaagtatgcaaagcatgcatctcaattagtcagcaacca
tagtcccgcccctaactccgcccatcccgcccctaactccgcccagttccgcccattctccgccccatggctgactaattttttttattt
atgcagaggccgaggccgcctcggcctctgagctattccagaagtagtgaggaggcttttttggaggcctaggcttttgcaaaaag
cttggacacaagacaggcttgcgagatatgtttgagaataccactttatcccgcgtcagggagaggcagtgcgtaaaaagacgc
ggactcatgtgaaatactggtttttagtgcgccagatctctataatctcgcgcaacctattttcccctcgaacactttttaagccgtag
ataaacaggctgggacacttcacatgagcgaaaaatacatcgtcacctgggacatgttgcagatccatgcacgtaaactcgcaa
gccgactgatgccttctgaacaatggaaaggcattattgccgtaagccgtggcggtctgtaccgggtgcgttactggcgcgtgaac
tgggtattcgtcatgtcgataccgtttgtatttccagctacgatcacgacaaccagcgcgagcttaaagtgctgaaacgcgcagaa
ggcgatggcgaaggcttcatcgttattgatgacctggtggataccggtggtactgcggttgcgattcgtgaaatgtatccaaaagc
gcactttgtcaccatcttcgcaaaaccggctggtcgtccgctggttgatgactatgttgttgatatcccgcaagatacctggattgaa
cagccgtgggatatgggcgtcgtattcgtcccgccaatctccggtcgctaatcttttcaacgcctggcactgccgggcgttgttctttt
taacttcaggcgggttacaatagtttccagtaagtattctggaggctgcatccatgacacaggcaaacctgagcgaaaccctgttc
aaaccccgctttaaacatcctgaaacctcgacgctagtccgccgctttaatcacggcgcacaaccgcctgtgcagtcggcccttga
tggtaaaaccatccctcactggtatcgcatgattaaccgtctgatgtggatctggcgcggcattgacccacgcgaaatcctcgacg
tccaggcacgtattgtgatgagcgatgccgaacgtaccgacgatgatttatacgatacggtgattggctaccgtggcggcaactgg
atttatgagtgggccccggatctttgtgaaggaaccttacttctgtggtgtgacataattggacaaactacctacagagatttaaag
ctctaaggtaaatataaaatttttaagtgtataatgtgttaaactactgattctaattgtttgtgtattttagattccaacctatggaa
ctgatgaatgggagcagtggtggaatgcctttaatgaggaaaacctgttttgctcagaagaaatgccatctagtgatgatgaggct
actgctgactctcaacattctactcctccaaaaaagaagagaaaggtagaagaccccaaggactttccttcagaattgctaagttt
tttgagtcatgctgtgtttagtaatagaactcttgcttgctttgctatttacaccacaaaggaaaaagctgcactgctatacaagaa
aattatggaaaaatattctgtaacctttataagtaggcataacagttataatcataacatactgttttttcttactccacacaggcat
agagtgtctgctattaataactatgctcaaaaattgtgtacctttagctttttaatttgtaaaggggttaataaggaatatttgatgta
tagtgccttgactagagatcataatcagccataccacatttgtagaggttttacttgctttaaaaaacctcccacacctccccctga
acctgaaacataaaatgaatgcaattgttgttgttaacttgtttattgcagcttataatggttacaaataaagcaatagcatcacaa
atttcacaaataaagcatttttttcactgcattctagttgtggtttgtccaaactcatcaatgtatcttatcatgtctggatcaactgga
taactcaagctaaccaaaatcatcccaaacttcccaccccataccctattaccactgccaattacctagtggtttcatttactctaaa
cctgtgattcctctgaattattttcattttaaagaaattgtatttgttaaatatgtactacaaacttagtagttggaagggctaattca
ctcccaaagaagacaagatatccttgatctgtggatctaccacacacaaggctacttccctgattagcagaactacacaccaggg
ccaggggtcagatatccactgacctttggatggtgctacaagctagtaccagttgagccagataaggtagaagaggccaataaa
ggagagaacaccagcttgttacaccctgtgagcctgcatgggatggatgacccggagagagaagtgttagagtggaggtttgac
agccgcctagcatttcatcacgtggcccgagagctgcatccggagtacttcaagaactgctgatatcgagcttgctacaagggact
ttccgctggggactttccagggaggcgtggcctgggcgggactggggagtggcgagccctcagatcctgcatataagcagctgct
ttttgcctgtactgggtctctctggttagaccagatctgagcctgggagctctctggctaactagggaacccactgcttaagcctcaa
taaagcttgccttgagtgcttcaagtagtgtgtgcccgtctgttgtgtgactctggtaactagagatccctcagacccttttagtcagt
gtggaaaatctctagcagtggcgcccgaacagggacttgaaagcgaaagggaaaccagaggagctctctcgacgcaggactcg
gcttgctgaagcgcgcacggcaagaggcgaggggcggcgactggtgagtacgccaaaaattttgactagcggaggctagaagg
agagagatgggtgcgagagcgtcagtattaagcgggggagaattagatcgcgatgggaaaaaattcggttaaggccagggggа
aagaaaaaatataaattaaaacatatagtatgggcaagcagggagctagaacgattcgcagttaatcctggcctgttagaaaca
tcagaaggctgtagacaaatactgggacagctacaaccatcccttcagacaggatcagaagaacttagatcattatataatacag
tagcaaccctctattgtgtgcatcaaaggatagagataaaagacaccaaggaagctttagacaagatagaggaagagcaaaac
aaaagtaagaccaccgcacagcaagcggccggccgctgatcttcagacctggaggaggagatatgagggacaattggagaagt
gaattatataaatataaagtagtaaaaattgaaccattaggagtagcacccaccaaggcaaagagaagagtggtgcagagaga
aaaaagagcagtgggaataggagctttgttccttgggttcttgggagcagcaggaagcactatgggcgcagcgtcaatgacgctg
acggtacaggccagacaattattgtctggtatagtgcagcagcagaacaatttgctgagggctattgaggcgcaacagcatctgtt

| | |
|---|---|
| | gcaactcacagtctggggcatcaagcagctccaggcaagaatcctggctgtgaaagatacctaaaggatcaacagctcctgggg<br>atttggggttgctctggaaaactcatttgcaccactgctgtgccttggaatgctagttggagtaataaatctctggaacagatttgg<br>aatcacacgacctggatggagtgggacagagaaattaacaattacacaagcttaatacactccttaattgaagaatcgcaaaac<br>cagcaagaaaagaatgaacaagaattattggaattagataaatgggcaagtttgtggaattggtttaacataacaaattggctgt<br>ggtatataaaattattcataatgatagtaggaggcttggtaggtttaagaatagtttttgctgtactttctatagtgaatagagttag<br>gcagggatattcaccattatcgtttcagacccacctcccaaccccgaggggacccgacaggcccgaaggaatagaagaagaag<br>gtggagagagagacagagacagatccattcgattagtgaacggatctcgacggtatcgccaaatggcagtattcatccacaattt<br>taaaagaaaaggggggattggggggtacagtgcaggggaaagaatagtagacataatagcaacagacatacaaactaaagaa<br>ttacaaaaacaaattacaaaaattcaaaattttcgggtttattacagggacagcagagatccagtttatcgatgaggccctttcgt<br>cttcactcgagtttactccctatcagtgatagagaacgtatgaagagtttactccctatcagtgatagagaacgtatgcagacttta<br>ctccctatcagtgatagagaacgtataaggagtttactccctatcagtgatagagaacgtatgaccagtttactccctatcagtgat<br>agagaacgtatctacagtttactccctatcagtgatagagaacgtatatccagtttactccctatcagtgatagagaacgtatgtcg<br>aggtaggcgtgtacggtgggcgcctataaaagcagagctcgtttagtgaaccgtcagatcgcctggagcaattccacaacactttt<br>gtcttatacttacgcgtgtcgaccatatgaggttcctggccaccgtcggcgtctcgcccgaccaccagggcaagggtctgggcagc<br>gccgtcgtgctccccggagtggaggcggccgagcgcgccggggtgcccgccttcctggagacctccgcgccccgcaacctcccct<br>tctacgagcggctcggcttcaccgtcaccgccgacgtcgaggtgcccgaaggaccgcgcacctggtgcacgacccgcaagcccg<br>gtgccggatcgggagagggcagaggtatgggtaattataacccgggccctatatattaggctaactagctaactaggaattcggt<br>ctcacagcagcagcagcagcagcagcagcagcagcagcagcagcagcacagcagcagcagcagcagcagcagcagcagcag<br>cagcagcagcagcagcagcagcagcagcagcagcagcagcagcagcagcagcagcagcacagcagcagcagcagcagc<br>agcagcagcagcagcagcagcagcagcagcagcagcagcagcagcagcagcagcagcagcagcagcagcagcagcacagc<br>agcagcagcagcagcagcagcagcagcagcagcagcagcagcagcagcagcagcagcagcagcagcagcagcagcagcag<br>cagcagcacagcagcagcagcagcagcagcagcagcagcagcagcagcagcagcagcagcagcagcagcagcagcagcagc<br>agcagcagcagcagcagcagcacagcagcagcagcagcagcagcagcagcagcagcagcagcagcagcagcagcagcagc<br>agcagcagcagcagcagcagcagcagcagcagcagcacagcagcagcagcagcagcagcagcagcagcagcagcagcagc<br>agcagcagcagcagcagcagcagcagcagcacagcagcagcagcagcagcagcagcagcagcagcagcagcagcagcagc<br>agcagcagcagcagcagcagcagcagcagcagcagcagcagcacagcagcagcagcagcagcagcagcagcagcagcagc<br>agcagcagcagagacggagctcgtcgacgcggccgcaagcttggtacctttaagaccaatgacttacaaggcagctgtagatctt<br>agccacttttaaaagaaaaggggggactggaagggctaattcactcccaacgaagacaagatctgctttttgcttgtactgggtc<br>tctctggttagaccagatctgagcctgggagctctctggctaactagggaacccactgcttaagcctcaataaagcttgccttgagt<br>gcttcaagtagtgtgtgcccgtctgttgtgtgactctggtaactagagatccctcagacccttttagtcagtgtggaaaatctctagc<br>agcatctagaattaattccgtgtatt |
| Delta MS2 + Delta WPRE:pHR Tre3G Flank C1 240xCAG | ctatagtgtcacctaaatcgtatgtgtatgatacataaggttatgtattaattgtagccgcgttctaacgacaatatgtacaagccta<br>attgtgtagcatctggcttactgaagcagaccctatcatctctctcgtaaactgccgtcagagtcggtttggttggacgaaccttctg<br>agtttctggtaacgccgtcccgcacccggaaatggtcagcgaaccaatcagcagggtcatcgctagccagatcctctacgccgga<br>cgcatcgtggccggcatcaccggcgccacaggtgcggttgctggcgcctatatcgccgacatcaccgatggggaagatcgggctc<br>gccacttcgggctcatgagcgcttgtttcggcgtgggtatggtggcaggccccgtggccgggggactgttgggcgccatctccttgc<br>atgcaccattccttgcggcggcggtgctcaacggcctcaacctactactgggctgcttcctaatgcaggagtcgcataagggagag<br>cgtcgaatggtgcactctcagtacaatctgctctgatgccgcatagttaagccagccccgacacccgccaacacccgctgacgcgc<br>cctgacgggcttgtctgctcccggcatccgcttacagacaagctgtgaccgtctccgggagctgcatgtgtcagaggttttcaccgt<br>catcaccgaaacgcgcgagacgaaagggcctcgtgatacgcctatttttataggttaatgtcatgataataatggtttcttagacgt<br>caggtggcacttttcggggaaatgtgcgcggaacccctatttgtttatttttctaaatacattcaaatatgtatccgctcatgagaca<br>ataaccctgataaatgcttcaataatattgaaaaaggaagagtatgagtattcaacatttccgtgtcgcccttattcccttttttgcg<br>gcattttgccttcctgtttttgctcacccagaaacgctggtgaaagtaaaagatgctgaagatcagttgggtgcacgagtgggttac<br>atcgaactggatctcaacagcggtaagatccttgagagttttcgccccgaagaacgttttccaatgatgagcacttttaaagttctg<br>ctatgtggcgcggtattatcccgtattgacgccgggcaagagcaactcggtcgccgcatacactattctcagaatgacttggttga<br>gtactcaccagtcacagaaaagcatcttacggatggcatgacagtaagagaattatgcagtgctgccataaccatgagtgataac<br>actgcggccaacttacttctgacaacgatcggaggaccgaaggagctaaccgcttttttgcacaacatgggggatcatgtaactcg |

| | |
|---|---|
| | ccttgatcgttgggaaccggagctgaatgaagccataccaaacgacgagcgtgacaccacgatgcctgtagcaatggcaacaac<br>gttgcgcaaactattaactggcgaactacttactctagcttcccggcaacaattaatagactggatggaggcggataaagttgcag<br>gaccacttctgcgctcggcccttccggctggctggtttattgctgataaatctggagccggtgagcgtgggtctcgcggtatcattgc<br>agcactggggccagatggtaagccctcccgtatcgtagttatctacacgacggggagtcaggcaactatggatgaacgaaatag<br>acagatcgctgagataggtgcctcactgattaagcattggtaactgtcagaccaagtttactcatatatactttagattgatttaaa<br>acttcatttttaatttaaaaggatctaggtgaagatcctttttgataatctcatgaccaaaatcccttaacgtgagttttcgttccact<br>gagcgtcagaccccgtagaaaagatcaaaggatcttcttgagatcctttttttctgcgcgtaatctgctgcttgcaaacaaaaaaa<br>ccaccgctaccagcggtggtttgtttgccggatcaagagctaccaactctttttccgaaggtaactggcttcagcagagcgcagat<br>accaaatactgtctttctagtgtagccgtagttaggccaccacttcaagaactctgtagcaccgcctacatacctcgctctgctaatc<br>ctgttaccagtggctgctgccagtggcgataagtcgtgtcttaccgggttggactcaagacgatagttaccggataaggcgcagcg<br>gtcgggctgaacggggggttcgtgcacacagcccagcttggagcgaacgacctacaccgaactgagatacctacagcgtgagct<br>atgagaaagcgccacgcttcccgaagggagaaaggcggacaggtatccggtaagcggcagggtcggaacaggagagcgcacg<br>agggagcttccagggggaaacgcctggtatctttatagtcctgtcgggtttcgccacctctgacttgagcgtcgatttttgtgatgct<br>cgtcaggggggcggagcctatggaaaaacgccagcaacgcggcctttttacggttcctggccttttgctggccttttgctcacatgt<br>tctttcctgcgttatcccctgattctgtggataaccgtattaccgcctttgagtgagctgataccgctcgccgcagccgaacgaccga<br>gcgcagcgagtcagtgagcgaggaagcggaagagcgcccaatacgcaaaccgcctctccccgcgcgttggccgattcattaatg<br>cagctgtggaatgtgtgtcagttagggtgtggaaagtccccaggctccccagcaggcagaagtatgcaaagcatgcatctcaatt<br>agtcagcaaccaggtgtggaaagtccccaggctccccagcaggcagaagtatgcaaagcatgcatctcaattagtcagcaacca<br>tagtcccgcccctaactccgcccatcccgcccctaactccgcccagttccgcccattctccgccccatggctgactaattttttttattt<br>atgcagaggccgaggccgcctcggcctctgagctattccagaagtagtgaggaggcttttttggaggcctaggcttttgcaaaaag<br>cttggacacaagacaggcttgcgagatatgtttgagaataccactttatcccgcgtcagggagaggcagtgcgtaaaaagacgc<br>ggactcatgtgaaatactggtttttagtgcgccagatctctataatctcgcgcaacctattttcccctcgaacactttttaagccgtag<br>ataaacaggctgggacacttcacatgagcgaaaaatacatcgtcacctgggacatgttgcagatccatgcacgtaaactcgcaa<br>gccgactgatgccttctgaacaatggaaaggcattattgccgtaagccgtggcggtctgtaccgggtgcgttactggcgcgtgaac<br>tgggtattcgtcatgtcgataccgtttgtatttccagctacgatcacgacaaccagcgcgagcttaaagtgctgaaacgcgcagaa<br>ggcgatggcgaaggcttcatcgttattgatgacctggtggataccggtggtactgcggttgcgattcgtgaaatgtatccaaaagc<br>gcactttgtcaccatcttcgcaaaaccggctggtcgtccgctggttgatgactatgttgttgatatcccgcaagatacctggattgaa<br>cagccgtgggatatgggcgtcgtattcgtcccgccaatctccggtcgctaatcttttcaacgcctggcactgccgggcgttgttctttt<br>taacttcaggcgggttacaatagtttccagtaagtattctggaggctgcatccatgacacaggcaaacctgagcgaaaccctgttc<br>aaaccccgctttaaacatcctgaaacctcgacgctagtccgccgctttaatcacggcgcacaaccgcctgtgcagtcggcccttga<br>tggtaaaaccatccctcactggtatcgcatgattaaccgtctgatgtggatctggcgcggcattgacccacgcgaaatcctcgacg<br>tccaggcacgtattgtgatgagcgatgccgaacgtaccgacgatgatttatacgatacggtgattggctaccgtggcggcaactgg<br>atttatgagtgggccccggatctttgtgaaggaaccttacttctgtggtgtgacataattggacaaactacctacagagatttaaag<br>ctctaaggtaaatataaaatttttaagtgtataatgtgttaaactactgattctaattgtttgtgtattttagattccaacctatggaa<br>ctgatgaatgggagcagtggtggaatgcctttaatgaggaaaacctgttttgctcagaagaaatgccatctagtgatgatgaggct<br>actgctgactctcaacattctactcctccaaaaaagaagagaaaggtagaagaccccaaggactttccttcagaattgctaagttt<br>tttgagtcatgctgtgtttagtaatagaactcttgcttgctttgctatttacaccacaaaggaaaaagctgcactgctatacaagaa<br>aattatggaaaaatattctgtaacctttataagtaggcataacagttataatcataacatactgttttttcttactccacacaggcat<br>agagtgtctgctattaataactatgctcaaaaattgtgtacctttagctttttaatttgtaaaggggttaataaggaatatttgatgta<br>tagtgccttgactagagatcataatcagccataccacatttgtagaggttttacttgctttaaaaaacctcccacacctccccctga<br>acctgaaacataaaatgaatgcaattgttgttgttaacttgtttattgcagcttataatggttacaaataaagcaatagcatcacaa<br>atttcacaaataaagcatttttttcactgcattctagttgtggtttgtccaaactcatcaatgtatcttatcatgtctggatcaactgga<br>taactcaagctaaccaaaatcatcccaaacttcccaccccataccctattaccactgccaattacctagtggtttcatttactctaaa<br>cctgtgattcctctgaattattttcattttaaagaaattgtatttgttaaatatgtactacaaacttagtagttggaagggctaattca<br>ctcccaaagaagacaagatatccttgatctgtggatctaccacacacaaggctacttccctgattagcagaactacacaccaggg<br>ccaggggtcagatatccactgacctttggatggtgctacaagctagtaccagttgagccagataaggtagaagaggccaataaa<br>ggagagaacaccagcttgttacaccctgtgagcctgcatgggatggatgacccggagagagaagtgttagagtggaggtttgac<br>agccgcctagcatttcatcacgtggcccgagagctgcatccggagtacttcaagaactgctgatatcgagcttgctacaagggact |

| | |
|---|---|
| | ttccgctggggactttccagggaggcgtggcctgggcgggactggggagtggcgagccctcagatcctgcatataagcagctgct<br>ttttgcctgtactgggtctctctggttagaccagatctgagcctgggagctctctggctaactagggaacccactgcttaagcctcaa<br>taaagcttgccttgagtgcttcaagtagtgtgtgcccgtctgttgtgtgactctggtaactagagatccctcagacccttttagtcagt<br>gtggaaaatctctagcagtggcgcccgaacagggacttgaaagcgaaagggaaaccagaggagctctctcgacgcaggactcg<br>gcttgctgaagcgcgcacggcaagaggcgaggggcggcgactggtgagtacgccaaaaattttgactagcggaggctagaagg<br>agagagatgggtgcgagagcgtcagtattaagcgggggagaattagatcgcgatgggaaaaaattcggttaaggccagggggа<br>aagaaaaaatataaattaaaacatatagtatgggcaagcagggagctagaacgattcgcagttaatcctggcctgttagaaaca<br>tcagaaggctgtagacaaatactgggacagctacaaccatcccttcagacaggatcagaagaacttagatcattatataatacag<br>tagcaaccctctattgtgtgcatcaaaggatagagataaaagacaccaaggaagctttagacaagatagaggaagagcaaaac<br>aaaagtaagaccaccgcacagcaagcggccggccgctgatcttcagacctggaggaggagatatgagggacaattggagaagt<br>gaattatataaatataaagtagtaaaaattgaaccattaggagtagcacccaccaaggcaaagagaagagtggtgcagagaga<br>aaaaagagcagtgggaataggagctttgttccttgggttcttgggagcagcaggaagcactatgggcgcagcgtcaatgacgctg<br>acggtacaggccagacaattattgtctggtatagtgcagcagcagaacaatttgctgagggctattgaggcgcaacagcatctgtt<br>gcaactcacagtctggggcatcaagcagctccaggcaagaatcctggctgtgaaagatacctaaaggatcaacagctcctgggg<br>atttggggttgctctggaaaactcatttgcaccactgctgtgccttggaatgctagttggagtaataaatctctggaacagatttgg<br>aatcacacgacctggatggagtgggacagagaaattaacaattacacaagcttaatacactccttaattgaagaatcgcaaaac<br>cagcaagaaaagaatgaacaagaattattggaattagataaatgggcaagtttgtggaattggtttaacataacaaattggctgt<br>ggtatataaaattattcataatgatagtaggaggcttggtaggtttaagaatagtttttgctgtactttctatagtgaatagagttag<br>gcagggatattcaccattatcgtttcagacccacctcccaaccccgaggggacccgacaggcccgaaggaatagaagaagaag<br>gtggagagagagacagagacagatccattcgattagtgaacggatctcgacggtatcgccaaatggcagtattcatccacaattt<br>taaaagaaaaggggggattggggggtacagtgcaggggaaagaatagtagacataatagcaacagacatacaaactaaagaa<br>ttacaaaaacaaattacaaaaattcaaaattttcgggtttattacagggacagcagagatccagtttatcgatgaggccctttcgt<br>cttcactcgagtttactccctatcagtgatagagaacgtatgaagagtttactccctatcagtgatagagaacgtatgcagacttta<br>ctccctatcagtgatagagaacgtataaggagtttactccctatcagtgatagagaacgtatgaccagtttactccctatcagtgat<br>agagaacgtatctacagtttactccctatcagtgatagagaacgtatatccagtttactccctatcagtgatagagaacgtatgtcg<br>aggtaggcgtgtacggtgggcgcctataaaagcagagctcgtttagtgaaccgtcagatcgcctggagcaattccacaacactttt<br>gtcttatacttacgcgtgtcgaccatatgaggttcctggccaccgtcggcgtctcgcccgaccaccagggcaagggtctgggcagc<br>gccgtcgtgctccccggagtggaggcggccgagcgcgccggggtgcccgccttcctggagacctccgcgccccgcaacctcccct<br>tctacgagcggctcggcttcaccgtcaccgccgacgtcgaggtgcccgaaggaccgcgcacctggtgcacgacccgcaagcccg<br>gtgccggatcgggagagggcagaggtatgggtaattataacccgggccctatatattaggctaactagctaactaggaattcggt<br>ctcacagcagcagcagcagcagcagcagcagcagcagcagcagcagcacagcagcagcagcagcagcagcagcagcagcagcagcag<br>cagcagcagcagcagcagcagcagcagcagcagcagcagcagcagcagcagcagcagcagcagcagcacagcagcagcagcagcagc<br>agcagcagcagcagcagcagcagcagcagcagcagcagcagcagcagcagcagcagcagcagcagcagcagcagcagcagcacagc<br>agcagcagcagcagcagcagcagcagcagcagcagcagcagcagcagcagcagcagcagcagcagcagcagcagcagcagcagcag<br>cagcagcacagcagcagcagcagcagcagcagcagcagcagcagcagcagcagcagcagcagcagcagcagcagcagcagcagcagc<br>agcagcagcagcagcagcagcacagcagcagcagcagcagcagcagcagcagcagcagcagcagcagcagcagcagcagcagcagc<br>agcagcagcagcagcagcagcagcagcagcagcagcacagcagcagcagcagcagcagcagcagcagcagcagcagcagcagcagc<br>agcagcagcagcagcagcagcagcagcagcagcagcagcagcacagcagcagcagcagcagcagcagcagcagcagcagcagc<br>agcagcagcagcagcagcagcagcagcagcagcagcagcagcagcagcacagcagcagcagcagcagcagcagcagcagcagc<br>agcagcagcagagacggagctcgtcgacgcggccgcaagcttggtacctttaagaccaatgacttacaaggcagctgtagatctt<br>agccactttttaaaagaaaaggggggactggaagggctaattcactcccaacgaagacaagatctgctttttgcttgtactgggtc<br>tctctggttagaccagatctgagcctgggagctctctggctaactagggaacccactgcttaagcctcaataaagcttgccttgagt<br>gcttcaagtagtgtgtgcccgtctgttgtgtgactctggtaactagagatccctcagacccttttagtcagtgtggaaaatctctagc<br>agcatctagaattaattccgtgtatt |
| piggyBac: pPB Tre3G | atgaaaaagcctgaactcaccgcgacgtctgtcgagaagtttctgatcgaaaagttcgacagcgtctccgacctgatgcagctctc<br>ggagggcgaagaatctcgtgctttcagcttcgatgtaggagggcgtggatatgtcctgcgggtaaatagctgcgccgatggtttct<br>acaaagatcgttatgtttatcggcactttgcatcggccgcgctcccgattccggaagtgcttgacattggggaatttagcgagagcc |

| | |
|---|---|
| FlankC1_240xCAG_12xMS2 | tgacctattgcatctcccgccgtgcacagggtgtcacgttgcaagacctgcctgaaaccgaactgcccgctgttctgcagccggtc<br>gcggaggccatggatgcgatcgctgcggccgatcttagccagacgagcgggttcggcccattcggaccgcaaggaatcggtcaa<br>tacactacatggcgtgatttcatatgcgcgattgctgatccccatgtgtatcactggcaaactgtgatggacgacaccgtcagtgcg<br>tccgtcgcgcaggctctcgatgagctgatgctttgggccgaggactgccccgaagtccggcacctcgtgcacgcggatttcggctc<br>caacaatgtcctgacggacaatggccgcataacagcggtcattgactggagcgaggcgatgttcggggattcccaatacgaggtc<br>gccaacatcttcttctggaggccgtggttggcttgtatggagcagcagacgcgctacttcgagcggaggcatccggagcttgcagg<br>atcgccgcggctccgggcgtatatgctccgcattggtcttgaccaactctatcagagcttggttgacggcaatttcgatgatgcagc<br>ttgggcgcagggtcgatgcgacgcaatcgtccgatccggagccgggactgtcgggcgtacacaaatcgcccgcagaagcgcggc<br>cgtctggaccgatggctgtgtagaagtactcgccgatagtggaaaccgacgccccagcactcgtccgagggcaaaggaatagcc<br>tctggattacaaaatttgtgggttaacttgtttattgcagcttataatggttacaaataaagcaatagcatcacaaatttcacaaat<br>aaagcatttttttcactgcattctagttgtggtttgtccaaactcatcaatgtatcttatcatgtctggaattgactcaaatgatgtca<br>attagtctatcagaagctatctggtctcccttccgggggacaagacatccctgtttaatatttaaacagcagtgttcccaaactggg<br>ttcttatatcccttgctctggtcaaccaggttgcagggtttcctgtcctcacaggaacgaagtccctaaagaaacagtggcagccag<br>gtttagccccggaattgactggattcctttttttagggcccattggtatggctttttccccgtatcccccaggtgtctgcaggctcaaa<br>gagcagcgagaagcgttcagaggaaagcgatcccgtgccaccttccccgtgcccgggctgtccccgcacgctgccggctcgggg<br>atgcggggggagcgccggaccggagcggagccccgggcggctcgctgctgccccctagcgggggagggacgtaattacatccct<br>gggggctttgggggggggctgtccctgatatctataacaagaaaatatatatataataagttatcacgtaagtagaacatgaaat<br>aacaatataattatcgtatgagttaaatcttaaaagtcacgtaaaagataatcatgcgtcattttgactcacgcggtcgttatagtt<br>caaaatcagtgacacttaccgcattgacaagcacgcctcacgggagctccaagcggcgactgagatgtcctaaatgcacagcga<br>cggattcgcgctatttagaaagagagagcaatatttcaagaatgcatgcgtcaattttacgcagactatctttctagggttaatcta<br>gctgcatcaggatcatatcgtcgggtcttttttccggctcagtcatcgcccaagctggcgctatctgggcatcggggaggaagaag<br>cccgtgccttttcccgcgaggttgaagcggcatggaaagagtttgccgaggatgactgctgctgcattgacgttgagcgaaaacg<br>cacgtttaccatgatgattcgggaaggtgtggccatgcacgcctttaacggtgaactgttcgttcaggccacctgggataccagttc<br>gtcgcggcttttccggacacagttccggatggtcagcccgaagcgcatcagcaacccgaacaataccggcgacagccggaactg<br>ccgtgccggtgtgcagattaatgacagcggtgcggcgctgggatattacgtcagcgaggacgggtatcctggctggatgccgcag<br>aaatggacatggataccccgtgagttacccggcgggcgcgcttggcgtaatcatggtcatagctgtttcctgtgtgaaattgttatc<br>cgctcacaattccacacaacatacgagccggaagcataaagtgtaaagcctggggtgcctaatgagtgagctaactcacattaat<br>tgcgttgcgctcactgcccgctttccagtcgggaaacctgtcgtgccagctgcattaatgaatcggccaacgcgcggggagaggc<br>ggtttgcgtattgggcgctcttccgcttcctcgctcactgactcgctgcgctcggtcgttcggctgcggcgagcggtatcagctcact<br>caaaggcggtaatacggttatccacagaatcaggggataacgcaggaaagaacatgtgagcaaaaggccagcaaaaggccag<br>gaaccgtaaaaaggccgcgttgctggcgtttttccataggctccgcccccctgacgagcatcacaaaaatcgacgctcaagtcag<br>aggtggcgaaacccgacaggactataaagataccaggcgtttccccctggaagctccctcgtgcgctctcctgttccgaccctgcc<br>gcttaccggatacctgtccgcctttctcccttcgggaagcgtggcgctttctcatagctcacgctgtaggtatctcagttcggtgtag<br>gtcgttcgctccaagctgggctgtgtgcacgaaccccccgttcagcccgaccgctgcgccttatccggtaactatcgtcttgagtcc<br>aacccggtaagacacgacttatcgccactggcagcagccactggtaacaggattagcagagcgaggtatgtaggcggtgctaca<br>gagttcttgaagtggtggcctaactacggctacactagaaggacagtatttggtatctgcgctctgctgaagccagttaccttcgga<br>aaaagagttggtagctcttgatccggcaaacaaaccaccgctggtagcggtggtttttttgtttgcaagcagcagattacgcgcag<br>aaaaaaaggatctcaagaagatcctttgatcttttctacggggtctgacgctcagtggaacgaaaactcacgttaagggattttgg<br>tcatgagattatcaaaaaggatcttcacctagatccttttaaattaaaaatgaagttttaaatcaatctaaagtatatatgagtaaa<br>cttggtctgacagttaccaatgcttaatcagtgaggcacctatctcagcgatctgtctatttcgttcatccatagttgcctgactcccc<br>gtcgtgtagataactacgatacgggagggcttaccatctggccccagtgctgcaatgataccgcgagacccacgctcaccggctc<br>cagatttatcagcaataaaccagccagccggaagggccgagcgcagaagtggtcctgcaactttatccgcctccatccagtctatt<br>aattgttgccgggaagctagagtaagtagttcgccagttaatagtttgcgcaacgttgttgccattgctacaggcatcgtggtgtca<br>cgctcgtcgtttggtatggcttcattcagctccggttcccaacgatcaaggcgagttacatgatcccccatgttgtgcaaaaaagcg<br>gttagctccttcggtcctccgatcgttgtcagaagtaagttggccgcagtgttatcactcatggttatggcagcactgcataattctc<br>ttactgtcatgccatccgtaagatgcttttctgtgactggtgagtactcaaccaagtcattctgagaatagtgtatgcggcgaccga<br>gttgctcttgcccggcgtcaatacgggataataccgcgccacatagcagaactttaaaagtgctcatcattggaaaacgttcttcg<br>gggcgaaaactctcaaggatcttaccgctgttgagatccagttcgatgtaacccactcgtgcacccaactgatcttcagcatctttt |

| | |
|---|---|
| | acttcaccagcgtttctgggtgagcaaaaacaggaaggcaaaatgccgcaaaaaagggaataagggcgacacggaaatgttg aatactcatactcttcctttttcaatattattgaagcatttatcagggttattgtctcatgagcggatacatatttgaatgtatttagaa aaataaacaaataggggttccgcgcacatttccccgaaaagtgccacctaaattgtaagcgttaatattttgttaaaattcgcgtta aatttttgttaaatcagctcattttttaaccaataggccgaaatcggcaaaatcccttataaatcaaaagaatagaccgagatagg gttgagtgttgttccagtttggaacaagagtccactattaaagaacgtggactccaacgtcaaagggcgaaaaaccgtctatcag ggcgatggcccactacgtgaaccatcaccctaatcaagttttttggggtcgaggtgccgtaaagcactaaatcggaaccctaaag ggagcccccgatttagagcttgacggggaaagccggcgaacgtggcgagaaaggaagggaagaaagcgaaaggagcgggcg ctagggcgctggcaagtgtagcggtcacgctgcgcgtaaccaccacacccgccgcgcttaatgcgccgctacagggcgcgtccca ttcgccattcaggctgcgcaactgttgggaagggcgatcggtgcgggcctcttcgctattacgccagctggcgaaagggggatgtg ctgcaaggcgattaagttgggtaacgccagggttttcccagtcacgacgttgtaaaacgacggccagtgagcgcgcctcgttcatt cacgtttttgaacccgtggaggacgggcagactcgcggtgcaaatgtgttttacagcgtgatggagcagatgaagatgctcgaca cgctgcagaacacgcagctagattaaccctagaaagataatcatattgtgacgtacgttaaagataatcatgtgtaaaattgacg catgtgttttatcggtctgtatatcgaggtttatttattaatttgaatagatattaagttttattatatttacacttacatactaataata aattcaacaaacaatttatttatgtttatttatttattaaaaaaaaacaaaaactcaaaatttcttctataaagtaacaaaacttttat gagggacagcccccccccaaagcccccagggatgtaattacgtccctcccccgctagggggcagcagcgagccgcccggggctc cgctccggtccggcgctccccccgcatccccgagccggcagcgtgcggggacagcccgggcacggggaaggtggcacgggatc gctttcctctgaacgcttctcgctgctctttgagcctgcagacacctgggggggatacggggaaaaggcctccacggccactagtttt actccctatcagtgatagagaacgtatgaagagtttactccctatcagtgatagagaacgtatgcagactttactccctatcagtga tagagaacgtataaggagtttactccctatcagtgatagagaacgtatgaccagtttactccctatcagtgatagagaacgtatct acagtttactccctatcagtgatagagaacgtatatccagtttactccctatcagtgatagagaacgtataagctttaggcgtgtac ggtgggcgcctataaaagcagagctcgtttagtgaaccgtcagatcgcctggagcaattccacaacacttttgtcttataccaactt tccgtaccacttcctaccctcgtaaaggtacgcgtgtcgaccatatgaggttcctggccaccgtcggcgtctcgcccgaccaccag ggcaagggtctgggcagcgccgtcgtgctccccggagtggaggcggccgagcgcgccggggtgcccgccttcctggagacctcc gcgccccgcaacctccccttctacgagcggctcggcttcaccgtcaccgccgacgtcgaggtgcccgaaggaccgcgcacctggt gcacgacccgcaagcccggtgccggatcgggagagggcagaggtatgggtaattataacccgggccctatatattaggctaact agctaactaggaattcggtctcacagcagcagcagcagcagcagcagcagcagcagcagcagcacagcagcagcagcag cagcagcagcagcagcagcagcagcagcagcagcagcagcagcagcagcagcagcagcagcagcagcagcagcagcagcac agcagcagcagcagcagcagcagcagcagcagcagcagcagcagcagcagcagcagcagcagcagcagcagcagcagcag cagcagcagcagcacagcagcagcagcagcagcagcagcagcagcagcagcagcagcagcagcagcagcagcagcagcagc agcagcagcagcagcagcagcagcacagcagcagcagcagcagcagcagcagcagcagcagcagcagcagcagcagcagc agcagcagcagcagcagcagcagcagcagcagcagcagcacagcagcagcagcagcagcagcagcagcagcagcagcagc agcagcagcagcagcagcagcagcagcagcagcagcagcagcagcagcagcacagcagcagcagcagcagcagcagcagc agcagcagcagcagcagcagcagcagcagcagcagcagcagcagcagcagcagcagcagcagcacagcagcagcagcagc agcagcagcagcagcagcagcagcagcagagacggagctcgtcgacgcggccgcaagcttggtaccgagctcggatcctaagg tacctaattgcctagaaaacatgaggatcacccatgtctgcaggtcgactctagaaaacatgaggatcacccatgtctgcagtatt cccgggttcattagatcctaaggtacctaattgcctagaaaacatgaggatcacccatgtctgcaggtcgactctagaaaacatga ggatcacccatgtctgcagtattcccgggttcattagatcctaaggtacctaattgcctagaaaacatgaggatcacccatgtctgc aggtcgactctagaaaacatgaggatcacccatgtctgcagtattcccgggttcattagatcctaaggtacctaattgcctagaaa acatgaggatcacccatgtctgcaggtcgactctagaaaacatgaggatcacccatgtctgcagtattcccgggttcattagatcc taaggtacctaattgcctagaaaacatgaggatcacccatgtctgcaggtcgactctagaaaacatgaggatcacccatgtctgc agtattcccgggttcattagatcctaaggtacctaattgcctagaaaacatgaggatcacccatgtctgcaggtcgactctagaaa acatgaggatcacccatgtctgcagtattcccgggttcattagatccgcgcgcatatataccggttagtaatgagtttaaacgggg gaggctaactgaaacacggaaggagacaataccggaaggaacccgcgctatgacggcaataaaaagacagaataaaacgca cgggtgttgggtcgtttgttcataaacgcggggttcggtcccagggctggcactctgtcgataccccaccgagaccccattggggc caatacgcccgcgtttcttccttttccccaccccaccccccaagttcgggtgaaggcccagggctcgcagccaacgtcggggcggc aggccctgccatagcagatctgcgaaatccagctgaatctaagtggatctgcgatcgctccggtgcccgtcagtggcagagcgc acatcgcccacagtccccgagaagttggggggaggggtcggcaattgaacgggtgcctagagaaggtggcgcggggtaaactg |

| | |
|---|---|
| | ggaaagtgatgtcgtgtactggctccgcctttttcccgagggtgggggagaaccgtatataagtgcagtagtcgccgtgaacgttc<br>tttttcgcaacgggtttgccgccagaacacagctgaagcttcgaggggctcgcatctctccttcacgcgcccgccgccctacctgag<br>gccgccatccacgccggttgagtcgcgttctgccgcctcccgcctgtggtgcctcctgaactgcgtccgccgtctaggtaagtttaa<br>agctcaggtcgagaccgggcctttgtccggcgctcccttggagcctacctagactcagccggctctccacgctttgcctgaccctgc<br>ttgctcaactctacgtctttgtttcgttttctgttctgcgccgttacagatccaagctgtgaccggcgcctactctagtgccgccatgtc<br>cagactggacaagagcaaagtcataaactctgctctggaattactcaatggagtcggtatcgaaggcctgacgacaaggaaact<br>cgctcaaaagctgggagttgagcagcctaccctgtactggcacgtgaagaacaagcgggccctgctcgatgccctgccaatcgag<br>atgctggacaggcatcatacccactcctgcccccctggaaggcgagtcatggcaagactttctgcggaacaacgccaagtcatacc<br>gctgtgctctcctctcacatcgcgacggggctaaagtgcatctcggcacccgcccaacagagaaacagtacgaaaccctggaaa<br>atcagctcgcgttcctgtgtcagcaaggcttctccctggagaacgcactgtacgctctgtccgccgtgggccactttacactgggct<br>gcgtattggaggaacaggagcatcaagtagcaaaagaggaaagagagacacctaccaccgattctatgcccccacttctgaaac<br>aagcaattgagctgttcgaccggcagggagccgaacctgccttccttttcggcctggaactaatcatatgtggcctggagaaacag<br>ctaaagtgcgaaagcggcgggccgaccgacgcccttgacgattttgacttagacatgctcccagccgatgcccttgacgactttga<br>ccttgatatgctgcctgctgacgctcttgacgattttgaccttgacatgctccccggggagggcagaggaagtcttctaacatgcgg<br>tgacgtggaggagaatcccggccctgaatcc |
| RAN translation reporter: pHR Tre3G Flank C1 240xCAG BFP 12xMS2 | ctatagtgtcacctaaatcgtatgtgtatgatacataaggttatgtattaattgtagccgcgttctaacgacaatatgtacaagccta<br>attgtgtagcatctggcttactgaagcagaccctatcatctctctcgtaaactgccgtcagagtcggtttggttggacgaaccttctg<br>agtttctggtaacgccgtcccgcacccggaaatggtcagcgaaccaatcagcagggtcatcgctagccagatcctctacgccgga<br>cgcatcgtggccggcatcaccggcgccacaggtgcggttgctggcgcctatatcgccgacatcaccgatggggaagatcgggctc<br>gccacttcgggctcatgagcgcttgtttcggcgtgggtatggtggcaggccccgtggccgggggactgttgggcgccatctccttgc<br>atgcaccattccttgcggcggcggtgctcaacggcctcaacctactactgggctgcttcctaatgcaggagtcgcataagggagag<br>cgtcgaatggtgcactctcagtacaatctgctctgatgccgcatagttaagccagccccgacacccgccaacacccgctgacgcgc<br>cctgacgggcttgtctgctcccggcatccgcttacagacaagctgtgaccgtctccgggagctgcatgtgtcagaggttttcaccgt<br>catcaccgaaacgcgcgagacgaaagggcctcgtgatacgcctatttttataggttaatgtcatgataataatggtttcttagacgt<br>caggtggcacttttcggggaaatgtgcgcggaacccctatttgtttatttttctaaatacattcaaatatgtatccgctcatgagaca<br>ataaccctgataaatgcttcaataatattgaaaaaggaagagtatgagtattcaacatttccgtgtcgcccttattcccttttttgcg<br>gcattttgccttcctgtttttgctcacccagaaacgctggtgaaagtaaaagatgctgaagatcagttgggtgcacgagtgggttac<br>atcgaactggatctcaacagcggtaagatccttgagagttttcgccccgaagaacgttttccaatgatgagcacttttaaagttctg<br>ctatgtggcgcggtattatcccgtattgacgccgggcaagagcaactcggtcgccgcatacactattctcagaatgacttggttga<br>gtactcaccagtcacagaaaagcatcttacggatggcatgacagtaagagaattatgcagtgctgccataaccatgagtgataac<br>actgcggccaacttacttctgacaacgatcggaggaccgaaggagctaaccgcttttttgcacaacatgggggatcatgtaactcg<br>ccttgatcgttgggaaccggagctgaatgaagccataccaaacgacgagcgtgacaccacgatgcctgtagcaatggcaacaac<br>gttgcgcaaactattaactggcgaactacttactctagcttcccggcaacaattaatagactggatggaggcggataaagttgcag<br>gaccacttctgcgctcggcccttccggctggctggtttattgctgataaatctggagccggtgagcgtgggtctcgcggtatcattgc<br>agcactggggccagatggtaagccctcccgtatcgtagttatctacacgacggggagtcaggcaactatggatgaacgaaatag<br>acagatcgctgagataggtgcctcactgattaagcattggtaactgtcagaccaagtttactcatatatactttagattgatttaaa<br>acttcatttttaatttaaaaggatctaggtgaagatcctttttgataatctcatgaccaaaatcccttaacgtgagttttcgttccact<br>gagcgtcagaccccgtagaaaagatcaaaggatcttcttgagatcctttttttctgcgcgtaatctgctgcttgcaaacaaaaaaa<br>ccaccgctaccagcggtggtttgtttgccggatcaagagctaccaactctttttccgaaggtaactggcttcagcagagcgcagat<br>accaaatactgtctttctagtgtagccgtagttaggccaccacttcaagaactctgtagcaccgcctacatacctcgctctgctaatc<br>ctgttaccagtggctgctgccagtggcgataagtcgtgtcttaccgggttggactcaagacgatagttaccggataaggcgcagcg<br>gtcgggctgaacggggggttcgtgcacacagcccagcttggagcgaacgacctacaccgaactgagatacctacagcgtgagct<br>atgagaaagcgccacgcttcccgaagggagaaaggcggacaggtatccggtaagcggcagggtcggaacaggagagcgcacg<br>agggagcttccagggggaaacgcctggtatctttatagtcctgtcgggtttcgccacctctgacttgagcgtcgatttttgtgatgct<br>cgtcaggggggcggagcctatggaaaaacgccagcaacgcggcctttttacggttcctggccttttgctggccttttgctcacatgt<br>tctttcctgcgttatcccctgattctgtggataaccgtattaccgcctttgagtgagctgataccgctcgccgcagccgaacgaccga<br>gcgcagcgagtcagtgagcgaggaagcggaagagcgcccaatacgcaaaccgcctctccccgcgcgttggccgattcattaatg |

| | |
|---|---|
| | cagctgtggaatgtgtgtcagttagggtgtggaaagtccccaggctccccagcaggcagaagtatgcaaagcatgcatctcaatt<br>agtcagcaaccaggtgtggaaagtccccaggctccccagcaggcagaagtatgcaaagcatgcatctcaattagtcagcaacca<br>tagtcccgcccctaactccgcccatcccgcccctaactccgcccagttccgcccattctccgccccatggctgactaattttttttattt<br>atgcagaggccgaggccgcctcggcctctgagctattccagaagtagtgaggaggcttttttggaggcctaggcttttgcaaaaag<br>cttggacacaagacaggcttgcgagatatgtttgagaataccactttatcccgcgtcagggagaggcagtgcgtaaaaagacgc<br>ggactcatgtgaaatactggtttttagtgcgccagatctctataatctcgcgcaacctattttcccctcgaacactttttaagccgtag<br>ataaacaggctgggacacttcacatgagcgaaaaatacatcgtcacctgggacatgttgcagatccatgcacgtaaactcgcaa<br>gccgactgatgccttctgaacaatggaaaggcattattgccgtaagccgtggcggtctgtaccgggtgcgttactggcgcgtgaac<br>tgggtattcgtcatgtcgataccgtttgtatttccagctacgatcacgacaaccagcgcgagcttaaagtgctgaaacgcgcagaa<br>ggcgatggcgaaggcttcatcgttattgatgacctggtggataccggtggtactgcggttgcgattcgtgaaatgtatccaaaagc<br>gcactttgtcaccatcttcgcaaaaccggctggtcgtccgctggttgatgactatgttgttgatatcccgcaagatacctggattgaa<br>cagccgtgggatatgggcgtcgtattcgtcccgccaatctccggtcgctaatcttttcaacgcctggcactgccgggcgttgttctttt<br>taacttcaggcgggttacaatagtttccagtaagtattctggaggctgcatccatgacacaggcaaacctgagcgaaaccctgttc<br>aaaccccgctttaaacatcctgaaacctcgacgctagtccgccgctttaatcacggcgcacaaccgcctgtgcagtcggcccttga<br>tggtaaaaccatccctcactggtatcgcatgattaaccgtctgatgtggatctggcgcggcattgacccacgcgaaatcctcgacg<br>tccaggcacgtattgtgatgagcgatgccgaacgtaccgacgatgatttatacgatacggtgattggctaccgtggcggcaactgg<br>atttatgagtgggccccggatctttgtgaaggaaccttacttctgtggtgtgacataattggacaaactacctacagagatttaaag<br>ctctaaggtaaatataaaatttttaagtgtataatgtgttaaactactgattctaattgtttgtgtattttagattccaacctatggaa<br>ctgatgaatgggagcagtggtggaatgcctttaatgaggaaaacctgttttgctcagaagaaatgccatctagtgatgatgaggct<br>actgctgactctcaacattctactcctccaaaaaagaagagaaaggtagaagaccccaaggactttccttcagaattgctaagttt<br>tttgagtcatgctgtgtttagtaatagaactcttgcttgctttgctatttacaccacaaaggaaaaagctgcactgctatacaagaa<br>aattatggaaaaatattctgtaacctttataagtaggcataacagttataatcataacatactgttttttcttactccacacaggcat<br>agagtgtctgctattaataactatgctcaaaaattgtgtacctttagctttttaatttgtaaaggggttaataaggaatatttgatgta<br>tagtgccttgactagagatcataatcagccataccacatttgtagaggttttacttgctttaaaaaacctcccacacctccccctga<br>acctgaaacataaaatgaatgcaattgttgttgttaacttgtttattgcagcttataatggttacaaataaagcaatagcatcacaa<br>atttcacaaataaagcatttttttcactgcattctagttgtggtttgtccaaactcatcaatgtatcttatcatgtctggatcaactgga<br>taactcaagctaaccaaaatcatcccaaacttcccaccccataccctattaccactgccaattacctagtggtttcatttactctaaa<br>cctgtgattcctctgaattattttcattttaaagaaattgtatttgttaaatatgtactacaaacttagtagttggaagggctaattca<br>ctcccaaagaagacaagatatccttgatctgtggatctaccacacacaaggctacttccctgattagcagaactacacaccaggg<br>ccaggggtcagatatccactgacctttggatggtgctacaagctagtaccagttgagccagataaggtagaagaggccaataaa<br>ggagagaacaccagcttgttacaccctgtgagcctgcatgggatggatgacccggagagagaagtgttagagtggaggtttgac<br>agccgcctagcatttcatcacgtggcccgagagctgcatccggagtacttcaagaactgctgatatcgagcttgctacaagggact<br>ttccgctggggactttccagggaggcgtggcctgggcgggactggggagtggcgagccctcagatcctgcatataagcagctgct<br>ttttgcctgtactgggtctctctggttagaccagatctgagcctgggagctctctggctaactagggaacccactgcttaagcctcaa<br>taaagcttgccttgagtgcttcaagtagtgtgtgcccgtctgttgtgtgactctggtaactagagatccctcagacccttttagtcagt<br>gtggaaaatctctagcagtggcgcccgaacagggacttgaaagcgaaagggaaaccagaggagctctctcgacgcaggactcg<br>gcttgctgaagcgcgcacggcaagaggcgaggggcggcgactggtgagtacgccaaaaattttgactagcggaggctagaagg<br>agagagatgggtgcgagagcgtcagtattaagcgggggagaattagatcgcgatgggaaaaaattcggttaaggccaggggga<br>aagaaaaaatataaattaaaacatatagtatgggcaagcagggagctagaacgattcgcagttaatcctggcctgttagaaaca<br>tcagaaggctgtagacaaatactgggacagctacaaccatcccttcagacaggatcagaagaacttagatcattatataatacag<br>tagcaaccctctattgtgtgcatcaaaggatagagataaaagacaccaaggaagctttagacaagatagaggaagagcaaaac<br>aaaagtaagaccaccgcacagcaagcggccggccgctgatcttcagacctggaggaggagatatgagggacaattggagaagt<br>gaattatataaatataaagtagtaaaaattgaaccattaggagtagcacccaccaaggcaaagagaagagtggtgcagagaga<br>aaaaagagcagtgggaataggagctttgttccttgggttcttgggagcagcaggaagcactatgggcgcagcgtcaatgacgctg<br>acggtacaggccagacaattattgtctggtatagtgcagcagcagaacaatttgctgagggctattgaggcgcaacagcatctgtt<br>gcaactcacagtctggggcatcaagcagctccaggcaagaatcctggctgtgaaagatacctaaaggatcaacagctcctgggg<br>atttggggttgctctggaaaactcatttgcaccactgctgtgccttggaatgctagttggagtaataaatctctggaacagatttgg<br>aatcacacgacctggatggagtgggacagagaaattaacaattacacaagcttaatacactccttaattgaagaatcgcaaaac |

| | |
|---|---|
| | cagcaagaaaagaatgaacaagaattattggaattagataaatgggcaagtttgtggaattggtttaacataacaaattggctgt<br>ggtatataaaattattcataatgatagtaggaggcttggtaggtttaagaatagtttttgctgtactttctatagtgaatagagttag<br>gcagggatattcaccattatcgtttcagacccacctcccaaccccgaggggacccgacaggcccgaaggaatagaagaagaag<br>gtggagagagagacagagacagatccattcgattagtgaacggatctcgacggtatcgccaaatggcagtattcatccacaattt<br>taaaagaaaaggggggattggggggtacagtgcaggggaaagaatagtagacataatagcaacagacatacaaactaaagaa<br>ttacaaaaacaaattacaaaaattcaaaattttcgggtttattacagggacagcagagatccagtttatcgatgaggccctttcgt<br>cttcactcgagtttactccctatcagtgatagagaacgtatgaagagtttactccctatcagtgatagagaacgtatgcagacttta<br>ctccctatcagtgatagagaacgtataaggagtttactccctatcagtgatagagaacgtatgaccagtttactccctatcagtgat<br>agagaacgtatctacagtttactccctatcagtgatagagaacgtatatccagtttactccctatcagtgatagagaacgtatgtcg<br>aggtaggcgtgtacggtgggcgcctataaaagcagagctcgtttagtgaaccgtcagatcgcctggagcaattccacaacactttt<br>gtcttatacttACGCGTGTCGACcatatgAGGTTCCTGGCCACCGTCGGCGTCTCGCCCGACCACCAGGG<br>CAAGGGTCTGGGCAGCGCCGTCGTGCTCCCCGGAGTGGAGGCGGCCGAGCGCGCCGGGGTGCCC<br>GCCTTCCTGGAGACCTCCGCGCCCCGCAACCTCCCCTTCTACGAGCGGCTCGGCTTCACCGTCACCG<br>CCGACGTCGAGGTGCCCGAAGGACCGCGCACCTGGTGCATGACCCGCAAGCCCGGTGCCGGATC<br>GGGAGAGGGCAGAGGTatgGGTAATTATAACCCGGGCCCTATATATTAGGCTAACTAGCTAACTAG<br>GAATTCggtctcaCAGCAGCAGCAGCAGCAGCAGCAGCAGCAGCAGCAGCAGCACAGCAGCAG<br>CAGCAGCAGCAGCAGCAGCAGCAGCAGCAGCAGCAGCAGCAGCAGCAGCAGCAGCAGCAG<br>CAGCAGCAGCAGCAGCAGCAGCACAGCAGCAGCAGCAGCAGCAGCAGCAGCAGCAGCAGCAGC<br>AGCAGCAGCAGCAGCAGCAGCAGCAGCAGCAGCAGCAGCAGCAGCAGCAGCACAGCAGCA<br>GCAGCAGCAGCAGCAGCAGCAGCAGCAGCAGCAGCAGCAGCAGCAGCAGCAGCAGCAGCA<br>GCAGCAGCAGCAGCAGCAGCACAGCAGCAGCAGCAGCAGCAGCAGCAGCAGCAGCAGCAG<br>CAGCAGCAGCAGCAGCAGCAGCAGCAGCAGCAGCAGCAGCAGCAGCAGCACAGCAGCAGC<br>AGCAGCAGCAGCAGCAGCAGCAGCAGCAGCAGCAGCAGCAGCAGCAGCAGCAGCAGCAGC<br>AGCAGCAGCAGCAGCAGCAGCACAGCAGCAGCAGCAGCAGCAGCAGCAGCAGCAGCAGCA<br>GCAGCAGCAGCAGCAGCAGCAGCAGCAGCAGCACAGCAGCAGCAGCAGCAGCAGCAGCAG<br>CAGCAGCAGCAGCAGCAGCAGCAGCAGCAGCAGCAGCAGCAGCAGCAGCAGCAGCAGCAG<br>CACAGCAGCAGCAGCAGCAGCAGCAGCAGCAGCAGCAGCAGCAGCAgagacgGAGCTCgtcgacGC<br>ggccgcGTGAGCAAGGGCGAGGAGCTGTTCACCGGGGTGGTGCCCATCCTGGTCGAGCTGGACGG<br>CGACGTAAACGGCCACAAGTTCAGCGTGAGGGGCGAGGGCGAGGGCGATGCCACCAACGGCAA<br>GCTGACCCTGAAGTTCATCTGCACCACCGGCAAGCTGCCCGTGCCCTGGCCCACCCTCGTGACCAC<br>CCTGAGCCACGGCGTGCAGTGCTTCGCCCGCTACCCCGACCACATGAAGCAGCACGACTTCTTCAA<br>GTCCGCCATGCCCGAAGGCTACGTCCAGGAGCGCACCATCTTCTTCAAGGACGACGGCACCTACAA<br>GACCCGCGCCGAGGTGAAGTTCGAGGGCGACACCCTGGTGAACCGCATCGAGCTGAAGGGCGTC<br>GACTTCAAGGAGGACGGCAACATCCTGGGGCACAAGCTGGAGTACAACTTCAACAGCCACAACAT<br>CTATATCATGGCCGTCAAGCAGAAGAACGGCATCAAGGTGAACTTCAAGATCCGCCACAACGTGG<br>AGGACGGCAGCGTGCAGCTCGCCGACCACTACCAGCAGAACACCCCCATCGGCGACGGCCCCGTG<br>CTGCTGCCCGACAGCCACTACCTGAGCACCCAGTCCGTGCTGAGCAAAGACCCCAACGAGAAGCG<br>CGATCACATGGTCCTGCTGGAGTTCCGCACCGCCGCCGGGATCACTCTCGGCATGGACGAGCTGT<br>ACAAGTAAGGATCCTAAGGTACCTAATTGCCTAGAAAACATGAGGATCACCCATGTCTGCAGGTCG<br>ACTCTAGAAAACATGAGGATCACCCATGTCTGCAGTATTCCCGGGTTCATTAGATCCTAAGGTACCT<br>AATTGCCTAGAAAACATGAGGATCACCCATGTCTGCAGGTCGACTCTAGAAAACATGAGGATCACC<br>CATGTCTGCAGTATTCCCGGGTTCATTAGATCCTAAGGTACCTAATTGCCTAGAAAACATGAGGAT<br>CACCCATGTCTGCAGGTCGACTCTAGAAAACATGAGGATCACCCATGTCTGCAGTATTCCCGGGTT<br>CATTAGATCCTAAGGTACCTAATTGCCTAGAAAACATGAGGATCACCCATGTCTGCAGGTCGACTC<br>TAGAAAACATGAGGATCACCCATGTCTGCAGTATTCCCGGGTTCATTAGATCCTAAGGTACCTAATT<br>GCCTAGAAAACATGAGGATCACCCATGTCTGCAGGTCGACTCTAGAAAACATGAGGATCACCCAT<br>GTCTGCAGTATTCCCGGGTTCATTAGATCCTAAGGTACCTAATTGCCTAGAAAACATGAGGATCAC<br>CCATGTCTGCAGGTCGACTCTAGAAAACATGAGGATCACCCATGTCTGCAGTATTCCCGGGTTCAT |

| | |
|---|---|
| | TAGATCTGCGCGCGATCGATATCAGCGCTTTAAATTTGCGCATGCTAGTgTAAggcatgcaagcttgatatcaagcttatcgataatcaacctctggattacaaaatttgtgaaagattgactggtattcttaactatgttgctccttttacgctatgtggatacgctgctttaatgcctttgtatcatgctattgcttcccgtatggctttcattttctcctccttgtataaatcctggttgctgtctctttatgaggagttgtggcccgttgtcaggcaacgtggcgtggtgtgcactgtgtttgctgacgcaacccccactggttggggcattgccaccacctgtcagctcctttccgggactttcgctttccccctccctattgccacggcggaactcatcgccgcctgccttgcccgctgctggacaggggctcggctgttgggcactgacaattccgtggtgttgtcggggaaatcatcgtcctttccttggctgctcgcctgtgttgccacctggattctgcgcgggacgtccttctgctacgtcccttcggccctcaatccagcggaccttccttcccgcggcctgctgccggctctgcggcctcttccgcgtcttcgccttcgccctcagacgagtcggatctccctttgggccgcctccccgcatcgataccgtcgacctcgagggaattaattcgagctcggtacctttaagaccaatgacttacaaggcagctgtagatcttagccactttttaaaagaaaaggggggactggaagggctaattcactcccaacgaagacaagatctgctttttgcttgtactgggtctctctggttagaccagatctgagcctgggagctctctggctaactagggaacccactgcttaagcctcaataaagcttgccttgagtgcttcaagtagtgtgtgcccgtctgttgtgtgactctggtaactagagatccctcagacccttttagtcagtgtggaaaatctctagcagcatctagaattaattccgtatt |
| RAN translation reporter with stops: pHR Tre3G Flank C1 240xCAG STOP BFP 12xMS2 | ctatagtgtcacctaaatcgtatgtgtatgatacataaggttatgtattaattgtagccgcgttctaacgacaatatgtacaagcctaattgtgtagcatctggcttactgaagcagaccctatcatctctctcgtaaactgccgtcagagtcggtttggttggacgaaccttctgagtttctggtaacgccgtcccgcacccggaaatggtcagcgaaccaatcagcagggtcatcgctagccagatcctctacgccggacgcatcgtggccggcatcaccggcgccacaggtgcggttgctggcgcctatatcgccgacatcaccgatggggaagatcgggctcgccacttcgggctcatgagcgcttgtttcggcgtgggtatggtggcaggccccgtggccgggggactgttgggcgccatctccttgcatgcaccattccttgcggcggcggtgctcaacggcctcaacctactactgggctgcttcctaatgcaggagtcgcataagggagagcgtcgaatggtgcactctcagtacaatctgctctgatgccgcatagttaagccagccccgacacccgccaacacccgctgacgcgccctgacgggcttgtctgctcccggcatccgcttacagacaagctgtgaccgtctccgggagctgcatgtgtcagaggttttcaccgtcatcaccgaaacgcgcgagacgaaagggcctcgtgatacgcctatttttataggttaatgtcatgataataatggtttcttagacgtcaggtggcacttttcggggaaatgtgcgcggaacccctatttgtttatttttctaaatacattcaaatatgtatccgctcatgagacaataaccctgataaatgcttcaataatattgaaaaaggaagagtatgagtattcaacatttccgtgtcgcccttattcccttttttgcggcattttgccttcctgtttttgctcacccagaaacgctggtgaaagtaaaagatgctgaagatcagttgggtgcacgagtgggttacatcgaactggatctcaacagcggtaagatccttgagagttttcgccccgaagaacgttttccaatgatgagcacttttaaagttctgctatgtggcgcggtattatcccgtattgacgccgggcaagagcaactcggtcgccgcatacactattctcagaatgacttggttgagtactcaccagtcacagaaaagcatcttacggatggcatgacagtaagagaattatgcagtgctgccataaccatgagtgataacactgcggccaacttacttctgacaacgatcggaggaccgaaggagctaaccgcttttttgcacaacatgggggatcatgtaactcgccttgatcgttgggaaccggagctgaatgaagccataccaaacgacgagcgtgacaccacgatgcctgtagcaatggcaacaacgttgcgcaaactattaactggcgaactacttactctagcttcccggcaacaattaatagactggatggaggcggataaagttgcaggaccacttctgcgctcggcccttccggctggctggtttattgctgataaatctggagccggtgagcgtgggtctcgcggtatcattgcagcactggggccagatggtaagccctcccgtatcgtagttatctacacgacggggagtcaggcaactatggatgaacgaaatagacagatcgctgagataggtgcctcactgattaagcattggtaactgtcagaccaagtttactcatatatactttagattgatttaaaacttcatttttaatttaaaaggatctaggtgaagatcctttttgataatctcatgaccaaaatcccttaacgtgagttttcgttccactgagcgtcagaccccgtagaaaagatcaaaggatcttcttgagatcctttttttctgcgcgtaatctgctgcttgcaaacaaaaaaaccaccgctaccagcggtggtttgtttgccggatcaagagctaccaactctttttccgaaggtaactggcttcagcagagcgcagataccaaatactgtctttctagtgtagccgtagttaggccaccacttcaagaactctgtagcaccgcctacatacctcgctctgctaatcctgttaccagtggctgctgccagtggcgataagtcgtgtcttaccgggttggactcaagacgatagttaccggataaggcgcagcggtcgggctgaacggggggttcgtgcacacagcccagcttggagcgaacgacctacaccgaactgagatacctacagcgtgagctatgagaaagcgccacgcttcccgaagggagaaaggcggacaggtatccggtaagcggcagggtcggaacaggagagcgcacgagggagcttccagggggaaacgcctggtatctttatagtcctgtcgggtttcgccacctctgacttgagcgtcgatttttgtgatgctcgtcaggggggcggagcctatggaaaaacgccagcaacgcggcctttttacggttcctggccttttgctggccttttgctcacatgttctttcctgcgttatcccctgattctgtggataaccgtattaccgcctttgagtgagctgataccgctcgccgcagccgaacgaccgagcgcagcgagtcagtgagcgaggaagcggaagagcgcccaatacgcaaaccgcctctccccgcgcgttggccgattcattaatgcagctgtggaatgtgtgtcagttagggtgtggaaagtccccaggctccccagcaggcagaagtatgcaaagcatgcatctcaattagtcagcaaccaggtgtggaaagtccccaggctccccagcaggcagaagtatgcaaagcatgcatctcaattagtcagcaacca |

| | |
|---|---|
| | tagtcccgcccctaactccgcccatcccgcccctaactccgcccagttccgcccattctccgccccatggctgactaattttttttattt<br>atgcagaggccgaggccgcctcggcctctgagctattccagaagtagtgaggaggcttttttggaggcctaggcttttgcaaaaag<br>cttggacacaagacaggcttgcgagatatgtttgagaataccactttatcccgcgtcagggagaggcagtgcgtaaaaagacgc<br>ggactcatgtgaaatactggtttttagtgcgccagatctctataatctcgcgcaacctattttcccctcgaacactttttaagccgtag<br>ataaacaggctgggacacttcacatgagcgaaaaatacatcgtcacctgggacatgttgcagatccatgcacgtaaactcgcaa<br>gccgactgatgccttctgaacaatggaaaggcattattgccgtaagccgtggcggtctgtaccgggtgcgttactggcgcgtgaac<br>tgggtattcgtcatgtcgataccgtttgtatttccagctacgatcacgacaaccagcgcgagcttaaagtgctgaaacgcgcagaa<br>ggcgatggcgaaggcttcatcgttattgatgacctggtggataccggtggtactgcggttgcgattcgtgaaatgtatccaaaagc<br>gcactttgtcaccatcttcgcaaaaccggctggtcgtccgctggttgatgactatgttgttgatatcccgcaagatacctggattgaa<br>cagccgtgggatatgggcgtcgtattcgtcccgccaatctccggtcgctaatcttttcaacgcctggcactgccgggcgttgttctttt<br>taacttcaggcgggttacaatagtttccagtaagtattctggaggctgcatccatgacacaggcaaacctgagcgaaaccctgttc<br>aaaccccgctttaaacatcctgaaacctcgacgctagtccgccgctttaatcacggcgcacaaccgcctgtgcagtcggcccttga<br>tggtaaaaccatccctcactggtatcgcatgattaaccgtctgatgtggatctggcgcggcattgacccacgcgaaatcctcgacg<br>tccaggcacgtattgtgatgagcgatgccgaacgtaccgacgatgatttatacgatacggtgattggctaccgtggcggcaactgg<br>atttatgagtgggccccggatctttgtgaaggaaccttacttctgtggtgtgacataattggacaaactacctacagagatttaaag<br>ctctaaggtaaatataaaatttttaagtgtataatgtgttaaactactgattctaattgtttgtgtattttagattccaacctatggaa<br>ctgatgaatgggagcagtggtggaatgcctttaatgaggaaaacctgttttgctcagaagaaatgccatctagtgatgatgaggct<br>actgctgactctcaacattctactcctccaaaaaagaagagaaaggtagaagaccccaaggactttccttcagaattgctaagttt<br>tttgagtcatgctgtgtttagtaatagaactcttgcttgctttgctatttacaccacaaaggaaaaagctgcactgctatacaagaa<br>aattatggaaaaatattctgtaacctttataagtaggcataacagttataatcataacatactgttttttcttactccacacaggcat<br>agagtgtctgctattaataactatgctcaaaaattgtgtacctttagctttttaatttgtaaaggggttaataaggaatatttgatgta<br>tagtgccttgactagagatcataatcagccataccacatttgtagaggttttacttgctttaaaaaacctcccacacctccccctga<br>acctgaaacataaaatgaatgcaattgttgttgttaacttgtttattgcagcttataatggttacaaataaagcaatagcatcacaa<br>atttcacaaataaagcatttttttcactgcattctagttgtggtttgtccaaactcatcaatgtatcttatcatgtctggatcaactgga<br>taactcaagctaaccaaaatcatcccaaacttcccaccccataccctattaccactgccaattacctagtggtttcatttactctaaa<br>cctgtgattcctctgaattattttcattttaaagaaattgtatttgttaaatatgtactacaaacttagtagttggaagggctaattca<br>ctcccaaagaagacaagatatccttgatctgtggatctaccacacacaaggctacttccctgattagcagaactacacaccaggg<br>ccaggggtcagatatccactgacctttggatggtgctacaagctagtaccagttgagccagataaggtagaagaggccaataaa<br>ggagagaacaccagcttgttacaccctgtgagcctgcatgggatggatgacccggagagagaagtgttagagtggaggtttgac<br>agccgcctagcatttcatcacgtggcccgagagctgcatccggagtacttcaagaactgctgatatcgagcttgctacaagggact<br>ttccgctggggactttccagggaggcgtggcctgggcgggactggggagtggcgagccctcagatcctgcatataagcagctgct<br>ttttgcctgtactgggtctctctggttagaccagatctgagcctgggagctctctggctaactagggaacccactgcttaagcctcaa<br>taaagcttgccttgagtgcttcaagtagtgtgtgcccgtctgttgtgtgactctggtaactagagatccctcagacccttttagtcagt<br>gtggaaaatctctagcagtggcgcccgaacagggacttgaaagcgaaagggaaaccagaggagctctctcgacgcaggactcg<br>gcttgctgaagcgcgcacggcaagaggcgaggggcggcgactggtgagtacgccaaaaattttgactagcggaggctagaagg<br>agagagatgggtgcgagagcgtcagtattaagcgggggagaattagatcgcgatgggaaaaaattcggttaaggccagggggа<br>aagaaaaaatataaattaaaacatatagtatgggcaagcagggagctagaacgattcgcagttaatcctggcctgttagaaaca<br>tcagaaggctgtagacaaatactgggacagctacaaccatcccttcagacaggatcagaagaacttagatcattatataatacag<br>tagcaaccctctattgtgtgcatcaaaggatagagataaaagacaccaaggaagctttagacaagatagaggaagagcaaaac<br>aaaagtaagaccaccgcacagcaagcggccggccgctgatcttcagacctggaggaggagatatgagggacaattggagaagt<br>gaattatataaatataaagtagtaaaaattgaaccattaggagtagcacccaccaaggcaaagagaagagtggtgcagagaga<br>aaaaagagcagtgggaataggagctttgttccttgggttcttgggagcagcaggaagcactatgggcgcagcgtcaatgacgctg<br>acggtacaggccagacaattattgtctggtatagtgcagcagcagaacaatttgctgagggctattgaggcgcaacagcatctgtt<br>gcaactcacagtctggggcatcaagcagctccaggcaagaatcctggctgtgaaagatacctaaaggatcaacagctcctgggg<br>atttggggttgctctggaaaactcatttgcaccactgctgtgccttggaatgctagttggagtaataaatctctggaacagatttgg<br>aatcacacgacctggatggagtgggacagagaaattaacaattacacaagcttaatacactccttaattgaagaatcgcaaaac<br>cagcaagaaaagaatgaacaagaattattggaattagataaatgggcaagtttgtggaattggtttaacataacaaattggctgt<br>ggtatataaaattattcataatgatagtaggaggcttggtaggtttaagaatagtttttgctgtactttctatagtgaatagagttag |

| | |
|---|---|
| | gcagggatattcaccattatcgtttcagacccacctcccaaccccgaggggacccgacaggcccgaaggaatagaagaagaag<br>gtggagagagagacagagacagatccattcgattagtgaacggatctcgacggtatcgccaaatggcagtattcatccacaattt<br>taaaagaaaaggggggattggggggtacagtgcaggggaaagaatagtagacataatagcaacagacatacaaactaaagaa<br>ttacaaaaacaaattacaaaaattcaaaattttcgggtttattacagggacagcagagatccagtttatcgatgaggccctttcgt<br>cttcactcgagtttactccctatcagtgatagagaacgtatgaagagtttactccctatcagtgatagagaacgtatgcagacttta<br>ctccctatcagtgatagagaacgtataaggagtttactccctatcagtgatagagaacgtatgaccagtttactccctatcagtgat<br>agagaacgtatctacagtttactccctatcagtgatagagaacgtatatccagtttactccctatcagtgatagagaacgtatgtcg<br>aggtaggcgtgtacggtgggcgcctataaaagcagagctcgtttagtgaaccgtcagatcgcctggagcaattccacaacactttt<br>gtcttatacttACGCGTGTCGACcatatgAGGTTCCTGGCCACCGTCGGCGTCTCGCCCGACCACCAGGG<br>CAAGGGTCTGGGCAGCGCCGTCGTGCTCCCCGGAGTGGAGGCGGCCGAGCGCGCCGGGGTGCCC<br>GCCTTCCTGGAGACCTCCGCGCCCCGCAACCTCCCCTTCTACGAGCGGCTCGGCTTCACCGTCACCG<br>CCGACGTCGAGGTGCCCGAAGGACCGCGCACCTGGTGCATGACCCGCAAGCCCGGTGCCGGATC<br>GGGAGAGGGCAGAGGTatgGGTAATTATAACCCGGGCCCTATATATTAGGCTAACTAGCTAACTAG<br>GAATTCggtctcaCAGCAGCAGCAGCAGCAGCAGCAGCAGCAGCAGCAGCAGCACAGCAGCAG<br>CAGCAGCAGCAGCAGCAGCAGCAGCAGCAGCAGCAGCAGCAGCAGCAGCAGCAGCAGCAGCAG<br>CAGCAGCAGCAGCAGCAGCAGCACAGCAGCAGCAGCAGCAGCAGCAGCAGCAGCAGCAGCAGC<br>AGCAGCAGCAGCAGCAGCAGCAGCAGCAGCAGCAGCAGCAGCAGCAGCAGCAGCACAGCAGCA<br>GCAGCAGCAGCAGCAGCAGCAGCAGCAGCAGCAGCAGCAGCAGCAGCAGCAGCAGCAGCAGCA<br>GCAGCAGCAGCAGCAGCAGCACAGCAGCAGCAGCAGCAGCAGCAGCAGCAGCAGCAGCAGCAG<br>CAGCAGCAGCAGCAGCAGCAGCAGCAGCAGCAGCAGCAGCAGCAGCAGCACAGCAGCAGC<br>AGCAGCAGCAGCAGCAGCAGCAGCAGCAGCAGCAGCAGCAGCAGCAGCAGCAGCAGCAGCAGC<br>AGCAGCAGCAGCAGCAGCAGCACAGCAGCAGCAGCAGCAGCAGCAGCAGCAGCAGCAGCAGCA<br>GCAGCAGCAGCAGCAGCAGCAGCAGCAGCACAGCAGCAGCAGCAGCAGCAGCAGCAGCAG<br>CAGCAGCAGCAGCAGCAGCAGCAGCAGCAGCAGCAGCAGCAGCAGCAGCAGCAGCAGCAGCAG<br>CACAGCAGCAGCAGCAGCAGCAGCAGCAGCAGCAGCAGCAGCAGCAgagacgGAGCTCgtcgacGC<br>ggccGCCTAGCTAGTAGATAGTAGATAGTAGGactagtGCGGCCGCGTGAGCAAGGGCGAGGAGCT<br>GTTCACCGGGGTGGTGCCCATCCTGGTCGAGCTGGACGGCGACGTAAACGGCCACAAGTTCAGCG<br>TGAGGGGCGAGGGCGAGGGCGATGCCACCAACGGCAAGCTGACCCTGAAGTTCATCTGCACCAC<br>CGGCAAGCTGCCCGTGCCCTGGCCCACCCTCGTGACCACCCTGAGCCACGGCGTGCatAGTGCTTC<br>GCCCGCTACCCCGACCACATGAAGCAGCACGACTTCTTCAAGTCCGCCATGCCCGAAGGCTACGTC<br>CAGGAGCGCACCATCTTCTTCAAGGACGACGGCACCTACAAGACCCGCGCCGAGGTGAAGTTCGA<br>GGGCGACACCCTGGTGAACCGCATCGAGCTGAAGGGCGTCGACTTCAAGGAGGACGGCAACATC<br>CTGGGGCACAAGCTGGAGTACAACTTCAACAGCCACAACATCTATATCATGGCCGTCAAGCAGAA<br>GAACGGCATCAAGGTGAACTTCAAGATCCGCCACAACGTGGAGGACGGCAGCGTGCAGCTCGCC<br>GACCACTACCAGCAGAACACCCCCATCGGCGACGGCCCCGTGCTGCTGCCCGACAGCCACTACCTG<br>AGCACCCAGTCCGTGCTGAGCAAAGACCCCAACGAGAAGCGCGATCACATGGTCCTGCTGGAGTT<br>CCGCACCGCCGCCGGGATCACTCTCGGCATGGACGAGCTGTACAAGTAAGGATCCTAAGGTACCT<br>AATTGCCTAGAAAACATGAGGATCACCCATGTCTGCAGGTCGACTCTAGAAAACATGAGGATCACC<br>CATGTCTGCAGTATTCCCGGGTTCATTAGATCCTAAGGTACCTAATTGCCTAGAAAACATGAGGAT<br>CACCCATGTCTGCAGGTCGACTCTAGAAAACATGAGGATCACCCATGTCTGCAGTATTCCCGGGTT<br>CATTAGATCCTAAGGTACCTAATTGCCTAGAAAACATGAGGATCACCCATGTCTGCAGGTCGACTC<br>TAGAAAACATGAGGATCACCCATGTCTGCAGTATTCCCGGGTTCATTAGATCCTAAGGTACCTAATT<br>GCCTAGAAAACATGAGGATCACCCATGTCTGCAGGTCGACTCTAGAAAACATGAGGATCACCCAT<br>GTCTGCAGTATTCCCGGGTTCATTAGATCCTAAGGTACCTAATTGCCTAGAAAACATGAGGATCAC<br>CCATGTCTGCAGGTCGACTCTAGAAAACATGAGGATCACCCATGTCTGCAGTATTCCCGGGTTCAT<br>TAGATCCTAAGGTACCTAATTGCCTAGAAAACATGAGGATCACCCATGTCTGCAGGTCGACTCTAG<br>AAAACATGAGGATCACCCATGTCTGCAGTATTCCCGGGTTCATTAGATCTGCGCGCGATCGATATC<br>AGCGCTTTAAATTTGCGCATGCTAGTgTAAggcatgcaagcttgatatcaagcttatcgataatcaacctctggatta |

| | |
|---|---|
| | caaaatttgtgaaagattgactggtattcttaactatgttgctccttttacgctatgtggatacgctgctttaatgcctttgtatcatgc<br>tattgcttcccgtatggctttcattttctcctccttgtataaatcctggttgctgtctctttatgaggagttgtggcccgttgtcaggcaa<br>cgtggcgtggtgtgcactgtgtttgctgacgcaacccccactggttggggcattgccaccacctgtcagctcctttccgggactttcg<br>ctttccccctccctattgccacggcggaactcatcgccgcctgccttgcccgctgctggacaggggctcggctgttgggcactgaca<br>attccgtggtgttgtcggggaaatcatcgtcctttccttggctgctcgcctgtgttgccacctggattctgcgcgggacgtccttctgc<br>tacgtcccttcggccctcaatccagcggaccttccttcccgcggcctgctgccggctctgcggcctcttccgcgtcttcgccttcgccc<br>tcagacgagtcggatctccctttgggccgcctccccgcatcgataccgtcgacctcgagggaattaattcgagctcggtacctttaa<br>gaccaatgacttacaaggcagctgtagatcttagccactttttaaaagaaaaggggggactggaagggctaattcactcccaacg<br>aagacaagatctgctttttgcttgtactgggtctctctggttagaccagatctgagcctgggagctctctggctaactagggaaccc<br>actgcttaagcctcaataaagcttgccttgagtgcttcaagtagtgtgtgcccgtctgttgtgtgactctggtaactagagatccctc<br>agacccttttagtcagtgtggaaaatctctagcagcatctagaattaattccgtgtatt |
