## Supplemental Legends and Discussion for "Flanking sequences regulate the toxicity, non-ATG translation, and aggregation of RNA with tandem CAG Repeats"

Supp. Figure 1. Flanking sequences alter localization of CAG repeat RNA inclusions. Related to Figure 1.

**A,** Scatterplot comparing the GC-content of the flanking sequence in a construct and the construct's toxicity. Each point represents a single construct and is the median of three wells. Trendline shows least-squared regression. Dotted lines correspond to 95% confidence interval. **B,** Quantification of transgene expression across cell lines. Expression levels are normalized to *Actin*. Each point represents a cell line which is the average from two separate RNA extractions. Control cells were uninduced cells that contain a construct capable of forming RNA aggregates upon induction. Data are summarized as mean ± s.d. Asterisks denote significance by Student's t-test. **C,** Same as B, except showing integration efficiencies. Control cells were not transduced with repeat constructs. Each point represents a cell line which is the average from two separate DNA extractions. **D,** Same as C, except that each point shows data from a separate DNA extraction. **E,** Quantification of cell death at varying timepoints after induction. Data are normalized to cell counts on day 0. Control cells were uninduced. Each point represents a separate well. Data are representative of two independent experiments. **F,** Micrograph of a cell line exhibiting nuclear RNA inclusions (left). Inset in left image denotes the ROI for the middle and right images. Binary image showing nuclear inclusions in white after applying intensity and size thresholds as described in methods (right). **G,** Representative fluorescence micrographs of cells expressing CAG repeats with variable flanking sequences. Arrow denotes representative cytoplasmic inclusion. Data are representative of ≥ 40 cells from ≥ 2 independent experiments. **H,** Quantification of percent of cells containing nuclear foci. Control cells express mCherry mRNAs or reverse-complement of mCherry RNAs, each fused to MS2 hairpins. Each point represents the median of ≥ 40 cells from ≥ 2 independent experiments. **I,** Quantification of the percent area of nucleus occupied by foci. Control cells as in G. Each point represents a single cell and data are summarized as the median ± interquartile range. **J,** Same as E, except that the binary image on the right shows cytoplasmic inclusions in white after thresholding. Arrow denotes representative cytoplasmic aggregate. **K,** Quantification of aggregate area in T1 cells transduced with varying viral titers of the repeat-containing construct (left). Each data point represents a single cell and data are summarized as median of each condition. Micrographs are representative of ≥ 25 cells. Asterisks denote significance by Mann-Whitney test. Fluorescence micrographs of cells transduced with varying viral titers (right, top). Fluorescence micrographs of a cytoplasmic aggregating CAG construct (right, middle) and a nuclear foci construct (right, bottom) induced with the stated concentrations of doxycycline. **L,** Micrographs showing cellular distribution of RNA after staining with RNA FISH probes targeting the MS2 region (left). Quantification of percent of cells with cytoplasmic aggregates (middle). Each point represents the median of ≥ 40 cells. Quantification of cytoplasmic aggregate area (right). Each point represents a single cell and data are summarized as median of each condition. **M,** Micrographs of cells stained with RNA FISH probes targeting the CAG repeats after deletion of the MS2 loop region (top) or deletion of both the MS2 loops and the WPRE (bottom). **N,** Fluorescence micrographs of cells expressing an aggregating CAG repeat construct that received the construct by transposase-mediated integration (left) or via transient transfection (middle) compared to an uninduced control (right). All scale bars, 10 µm. *: $p < 0.01$, n.s.: not significant ($p > 0.01$).

Supp. Figure 2. Cytoplasmic RNA inclusions co-localize with RNA-binding proteins. Related to Figure 3.

**A,** Immunofluorescence micrographs showing localization of 240xCAG RNA (MS2-YFP) and Tis11b in uninduced control cells (top) or induced cells that form cytoplasmic aggregates. **B,** Similar to B except immunofluorescence was performed on cells expressing nuclear foci CAG repeat RNAs (N1 and N2). Vermillion channel shows the stated epitopes in each row. Cells were counterstained with DAPI. Micrographs are representative of ≥ 40 cells. All scale bars, 10 μm.

Supp. Figure 3. Aggregating CAG repeat RNAs undergo RAN translation and co-aggregate with their RAN translation products. Related to Figure 4.

**A,** Immunofluorescence micrographs showing localization of 240xCAG RNA (MS2-YFP) and polyglutamine-containing proteins in cell lines with different flanking sequences. PolyQ channel is scaled independently for each image (i) and equally across all images (e). Nuclei were counterstained with DAPI. Micrographs are representative of ≥ 40 cells. **B,** Schematic of constructs containing BFP translation reporter with or without a stop codon in-frame with the BFP coding sequence (top). Micrographs showing localization of RNA (MS2-YFP) and BFP in cell lines described above (bottom). BFP images were scaled equally. Micrographs are representative of ≥ 40 cells. **C,** Quantification of aggregate area and BFP fluorescence in a single cell over time. Each point represents the single cell at different timepoints. Data are representative of ≥ 3 independent experiments. All scale bars, 10 μm.

Supp. Figure 4. Translation is required for CAG repeat RNA aggregation and toxicity. Related to Figure 5.

**A,** Scatterplot of aggregate area and BFP fluorescence in cells treated with doxycycline alone (red circles, control) or doxycycline and cycloheximide (black circles, cycloheximide). Each point represents a single cell. The median of each condition is represented by squares. **B,** Fluorescence in situ hybridization micrographs showing RNA localization and accumulation after induction with doxycycline alone or doxycycline and cycloheximide (left). Images are equally scaled. Quantification of RNA levels (right). Each point represents a single cell and data are summarized as the median ± interquartile range. Data are representative of 2 independent experiments with ≥ 40 cells. Asterisks denote significance by Mann-Whitney test. **C-D,** Similar to A-B except after treatment with doxycycline and either a control morpholino or 8xCTG morpholino. All scale bars, 10 µm. *: $p < 0.01$, n.s.: not significant ($p > 0.01$).

Supp. Figure 5. Formaldehyde fixation reduces detection of RNA at cytoplasmic aggregates.

**A,** Fluorescence micrographs of T1 cells fixed with formaldehyde and stained with RNA FISH probes (left). Intensity profile showing RNA intensities as detected by MS2-YFP (green) and by RNA FISH (vermillion) (right). Regions of the line corresponding to a nuclear focus and a cytoplasmic aggregate are labelled. **B,** Fluorescence micrographs of T1 cells fixed with a methanol solution and stained with RNA FISH probes. MS2-YFP is not shown as methanol denatures and prevents fluorescence detection of the protein. All scale bars, 10 µm. Data are representative of at least 2 independent experiments.

Supp. Movie 1. FRAP of nuclear foci. Related to Figure 2.

Time-lapse of a nuclear RNA focus bleached at t = 0 s. Cells transduced with construct N1 were induced for 24 h prior to bleaching. Arrowhead denotes photobleached region. Images are taken at 30 s intervals after bleaching. Scale bar, 10 µm.

Supp. Movie 2. FRAP of cytoplasmic RNA aggregates. Related to Figure 2.

Time-lapse of cytoplasmic RNA aggregate bleached at t = 0 s. Cells transduced with construct T1 were induced for 24 h prior to bleaching. Arrowheads denote photobleached regions. Images are taken at 30 s intervals after bleaching. Scale bar, 10 µm.

Supp. Movie 3. Real-time visualization of RAN translation and RNA aggregation. Related to Figure 4.

Time-lapse movie of cells co-transduced with construct T1 and BFP RAN translation reporter. Expression of T1 construct is induced at time t = 0. Images show composite (cyan) of MS2-YFP (green) and BFP (blue) channels. Images are taken at 30 min intervals after induction. Scale bar, 10 µm.

Supp. Table 1. Flanking sequences. Related to Fig. 1.

Supp. Table 2. Full plasmid sequences. Related to Figure 1.

Supp. Discussion

We observed cytoplasmic aggregation of expanded CAG repeat RNAs through the MS2 reporter system (RNA tagged with 12×MS2 hairpins and detected visualized via the co-expression of MS2CP-YFP protein) as well as via RNA FISH. Aggregation was more pronounced for CAG repeat-containing RNA tagged with the MS2 reporter as compared to the RNA without the MS2-hairpin tag. These differences may indicate that MS2 hairpin tag itself may have a propensity to aggregate. Our control RNAs (mCherry or its reverse complement) tagged with the MS2 reporter did not form appreciable inclusions. A second possibility is that the MS2CP-YFP protein may itself aggregate and exacerbate the RNA aggregation phenotype. We used a monomerized mutant of YFP (A206K) for all experiments, however, it may still have residual oligomerization propensity. Cells lacking MS2CP-YFP that were transduced with CAG repeat constructs lacking MS2 hairpins also exhibited RNA aggregation (Supp. Fig. 1M), demonstrating that the MS2 system is not required for RNA aggregation. Our interpretation of these data is that RNA aggregation may bring together MS2CP-YFP proteins at a high local concentration which may potentiate aggregation of MS2CP-YFP. These MS2CP-YFP aggregates may further recruit MS2-hairpin tagged RNA, and augment aggregation. These results highlight the importance of adequate controls when using fluorescent proteins to study large RNA-protein assemblies such as condensates and aggregates.
